# Quantitative data independent acquisition glycoproteomics of sparkling wine

**DOI:** 10.1101/2020.06.09.141226

**Authors:** Cassandra L. Pegg, Toan K. Phung, Christopher H. Caboche, Suchada Niamsuphap, Marshall Bern, Kate Howell, Benjamin L. Schulz

## Abstract

Sparkling wine is an alcoholic beverage enjoyed around the world. The sensory properties of sparkling wine depend on a complex interplay between the chemical and biochemical components in the final product. Glycoproteins have been linked to positive and negative qualities in sparkling wine, but the glycosylation profiles of sparkling wine have not been previously investigated in detail. We analysed the glyco/proteome of sparkling wines using protein- and glycopeptide-centric approaches. We developed an automated workflow that created ion libraries to analyse Sequential Window Acquisition of all THeoretical mass spectra (SWATH) Data Independent Acquisition (DIA) mass spectrometry data based on glycopeptides identified by Byonic. We applied our workflow to three pairs of experimental sparkling wines to assess the effects of aging on lees and of different yeast strains used in the Liqueur de Tirage for secondary fermentation. We found that aging a cuvée on lees for 24 months compared to 8 months led to a dramatic decrease in overall protein abundance and an enrichment in large glycans at specific sites in some proteins. Secondary fermentation of a Riesling wine with *Saccharomyces cerevisiae* yeast strain Siha4 produced more yeast proteins and glycoproteins than with *S. cerevisiae* yeast strain DV10. The abundance and glycosylation profiles of grape glycoproteins were also different between grape varieties. This work represents the first in-depth study into protein- and peptide-specific glycosylation in sparkling wines and describes a quantitative glycoproteomic SWATH/DIA workflow that is broadly applicable to other sample types.

## INTRODUCTION

Sparkling wine is enjoyed around the world in celebrations and festivities. Although sparkling wine represents only 9% of the wine market, it is the fastest growing grape-derived alcoholic beverage, both in sales and volume produced (1,2). One of the most important characteristics of sparkling wine is its effervescence: the formation of bubbles. The number, frequency, and longevity of bubbles, as well as other quality attributes, are affected by many factors during sparkling wine production, including the winemaking technique, the grape varieties used, the specific yeast strain and the fermentation conditions applied (3-5). Many complex biomolecules are extracted or produced during sparkling wine production, including proteins, polysaccharides, polyphenols and lipids, all of which can influence the texture, flavour, colour and foaming properties of the final product (3). Despite the low concentration (4-20 mg/L) (6) of proteins and glycoproteins in wine and sparkling wine, they are especially important for determining its sensory properties. They can alter the clarity and stability (7-9) and positively influence foaming (10). Foaming is a highly desirable quality in sparkling wines, and measurements such as foam height and stability are often used to assess quality (3). Glycoproteins are particularly important in controlling bubble formation and stability, as they surround and stabilise the gas bubbles of the foam in sparkling wine (11-15).

Glycoproteins are proteins that are post-translationally modified with complex oligosaccharides also known as glycans. The number, location, and structure of the attached glycans strongly affect the biological activities and biophysical properties of glycoproteins (16). Glycans are naturally heterogeneous due to their non-template driven biosynthetic pathways and the numerous possible configurations of monosaccharide topology and glycosidic linkages (17). The final glycan structures present on mature glycoproteins depend on the organism from which they are produced (18). The glycans found on yeast and plant proteins differ substantially. *Saccharomyces cerevisiae* yeast produce high mannose *N*-linked and oligomannose *O*-linked glycans attached to the side-chain amide of Asn and the hydroxyl groups of Ser or Thr, respectively (19). This oligomannose *O*-glycosylation has been observed on proteins in sparkling wine (14). The *N*-linked glycans of grape (*Vitis vinifera*) are usually high mannose or paucimannose structures, commonly with β(1,2)-linked xylose and core α(1,3)-linked fucose (20). *N*-linked glycans consistent with these structures have been observed on grape vacuolar invertase purified from grape must (20). *O*-linked glycans on grape glycoproteins are commonly attached through the hydroxyl group of hydroxyproline (Hyp) and contain predominately arabinose and/or galactose (21,22).

Sparkling wine can be produced with several different methods (3). In the Traditional Method (**Fig. 1**), grapes are picked and pressed to produce grape must, which is used for primary fermentation. The base wines produced from this primary fermentation can be blended to achieve different sensory attributes in the final product (23). The blended base wine then undergoes a second fermentation and aging on lees (3,24). The second fermentation is the key step in sparkling wine production. A mixture of sugar and yeast, known as the Liqueur de Tirage, is added to the blended base wine, and allows production of additional CO_2_ to carbonate the wine (3). The lees in sparkling wine are composed of yeast cells together with small amounts of co-adjuvants that facilitate sedimentation of the yeast at the end of active fermentation (24). Sparkling wines are deliberately left in contact with lees during the second fermentation in a process known as *sur lie* to improve the organoleptic qualities of the final wine (24). During this time autolysis of the yeast occurs, as well as degradation of lipids, proteins, and polysaccharides, which together modify the molecular complexity and properties of the sparkling wine (25). Both primary and secondary fermentations normally use pure starter strains, typically *S. cerevisiae*, to control fermentation and limit bottle-to-bottle variability. A large number of pure *S. cerevisiae* strains are available for commercial use. Despite the genetic similarity of these commercial wine strains (26), the strain of yeast used can have a substantial effect on the quality and flavour profile of the final sparkling wine product (27). The Traditional Method is set apart from other methods as the second fermentation takes place in the same sealed bottles from which the sparkling wine will eventually be consumed (3,24). Towards the end of their time on lees the bottles are rotated on an angle in a process called riddling, which collects the yeast sediment in the neck of the bottle before removal. After the sediment is removed, a *dosage* solution typically consisting of sugar and wine is added to refill the bottle (24). The winemaking process is then considered complete, and the wines are left to age before consumption.

**Figure 1.**
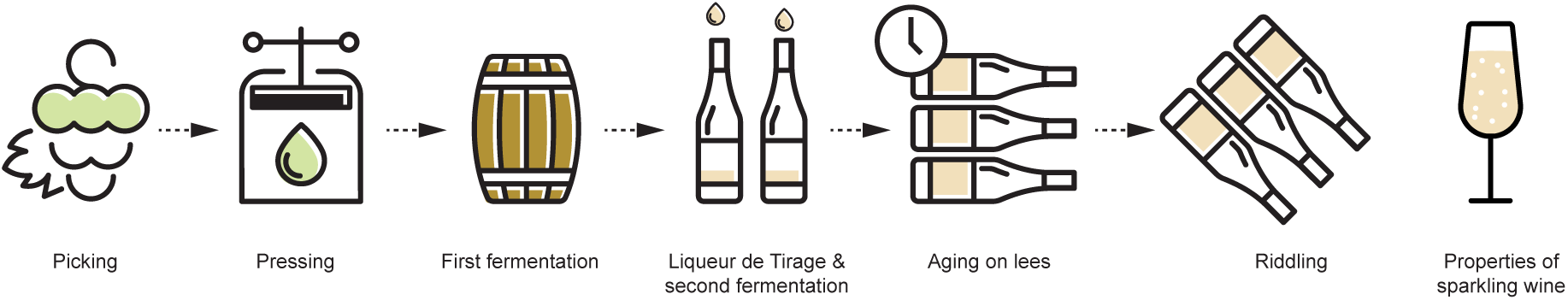
Overview of sparkling wine production using the Traditional Method.

Grape and yeast glycoproteins are released throughout the sparkling winemaking process, but are thought to be concentrated during the secondary fermentation and ageing on lees (28). Glycoproteins are an important class of macromolecule controlling foam production and foam quality in sparkling wine (11-15). The glycoproteins secreted by yeast and proteins from yeast autolysis are thought to be the main proteinaceous contributors to increased foaming (13,27). However, the combined presence of both yeast and grape glycoproteins yields the best foamability, highlighting the importance of glycoproteins from both species (13). Glycoproteins have been implicated in controlling the mouthfeel of both sparkling and still wines (29,30). Grape arabinogalactan proteins (7) and yeast mannoproteins (8,9) can also protect wine from haze formation, while other grape pathogenesis related glycoproteins such as thaumatin-like proteins and chitinases can cause haziness (31). Haze is an undesirable quality in wine, as aggregated proteins cause visible changes to clarity, considerably reducing the value of the product.

Despite the impact of glycoproteins on the quality of wine and sparkling wine, no high-throughput investigations of protein-specific glycosylation in wine have been previously conducted. Here, we developed a novel quantitative glycoproteomic and proteomic approach that integrated measurement of intact glycopeptides to a data independent acquisition (DIA) analytical workflow. Using this approach, we investigated the effects of aging on lees and the use of different yeast strains during the second fermentation on the molecular complexity and composition of sparkling wine.

## METHODS

### Experimental Design and Statistical Rationale

The purpose of this study was to measure the glycoproteome and proteome of sparkling wine to assess the effects of aging on lees and different yeast strains used during the second fermentation. We investigated the length of time on lees of a cuvée at 8 months or 24 months fermented with the yeast EC1118; the second fermentation of a Sauvignon blanc base wine with yeast strains DV10 and Zymaflore aged 16 months; and the second fermentation of a Riesling base wine with yeast strains DV10 and Siha4 aged 17 months. All six sparkling wine samples were prepared in technical triplicate (n=18). Samples were randomized using Microsoft Excel (For Mac v. 16.16.10) before MS analyses. Retention time standards were not used and MS1 data was acquired. A library was created (described in data analysis) using a sample measured with data-dependant acquisition (DDA) from each experimental condition (n=6). Statistical analyses for Sequential Window Acquisition of all THeoretical mass spectra (SWATH)-MS was performed using MSstats (v2.4) in R (32) with a significance threshold of P = 10^−5^ as described previously (33,34). Site-specificity of post-translational modifications was not a requirement for this study.

### Wine production

The wines were sourced from Sekt producer Schloss Vaux, in the Reingau, Germany. The wines were made at a commercial scale, with different experimental treatments. The sparkling wines were made with different base wines, made from Riesling grapes, Sauvignon blanc grapes or a cuvée (blend) from white wine grapes. The secondary fermentation was conducted with different commercial *S. cerevisiae* wine yeasts (Lalvin EC1118, Lalvin DV10, Laffort Zymaflore, or Siha4) and aged for different periods of time before riddling and stabilisation. The winemaking protocols were standard practice for this sparkling house, and were standardised between the treatments.

### Sample preparation

Proteins were prepared essentially as previously described (33). For each wine sample replicates of 250 µL were precipitated by addition of 1 mL methanol/acetone (1:1 v/v) and incubation overnight at −20 °C. The precipitated proteins were centrifuged for 10 min at 18,000 rcf and the protein pellets resuspended in 50 µL of 50 mM NH_4_HCO_3_ containing 10 mM dithiothreitol and incubated at room temperature for 10 min. A buffer of 50 mM NH_4_HCO_3_ containing 0.02 µg/µL of sequencing grade porcine trypsin (Sigma-Aldrich, MO, USA) was prepared and within 3 min of preparation 50 µL was added to the resuspended pellets. The samples were incubated for 16 h at 37 °C. Peptides were desalted and concentrated with a C18 ZipTip (10 µL pipette tip with a 0.6 µL resin bed; Millipore, MA, USA). Samples were dried and reconstituted in 100 µL of 0.1% formic acid.

### Mass spectrometry

Samples were analysed by LC-ESI-MS/MS using a Prominence nanoLC system (Shimadzu) coupled to a TripleTof 5600 instrument (SCIEX) using a Nanospray III interface essentially as previously described (35). Peptides and glycopeptides were separated with solvent A (1% CH_3_CN in 0.1% (v/v) aqueous formic acid) and solvent B (80% (v/v) CH_3_CN containing 0.1% (v/v) formic acid) with a gradient of 3-40% solvent B in 17 min. For DDA, 50 µL of the tryptic digests were injected (n=6). For SWATH/DIA analysis, 10 µL of the tryptic digests were injected (n=18). The mass spectrometer was operated in positive ion mode and gas and voltage settings were adjusted as required. Full MS scans were obtained with a range of *m/z* 350-1600 for both DDA and DIA with accumulation times of 0.5 s and 0.05 s, respectively. High sensitivity mode was used for DDA where the top 20 most intense precursors with charge states of 2-5 and intensities greater than 100 were selected for fragmentation with a collision energy (CE) of 40 V and a 15 V spread. An accumulation time of 0.05 s was used with a scan range of *m/z* 40-1600 and precursors were excluded for 5 s after two selections. High sensitivity mode was used for DIA where 34 windows were defined within a *m/z* range of 400-1250 with an isolation window of 26 *m/z* and overlap of 1 *m/z*. Automated rolling CE was used for each *m/z* window range with a 15 V spread and an accumulation time of 0.1 s.

### Data analysis

Peptide identification was performed with ProteinPilot 5.0.1 (SCIEX) using DDA files from one replicate of each sample. The files were searched against a combined protein database containing *Vitis vinifera* (grape) (NCBI RefSeq downloaded 02 August 2018 with 41,219 proteins and NCBI Accession RVX08988) (36), *S. cerevisiae* (yeast) (UniProt proteome UP000002311, downloaded 20 April 2018 with 6,049 proteins) and contaminants proteins (custom database created 11 November 2014 with 298 proteins). Standard search settings included: sample type, identification; digestion, trypsin; instrument, TripleTOF 5600; Cys alkylation, none; search effort, thorough. False discovery rate analysis using ProteinPilot was performed on all searches. Peptides identified with greater than 99% confidence and with a local false discovery rate of less than 1% were included for further analyses.

For glycopeptide analyses we used Byonic (Protein Metrics, v. 2.13.17) to search all DDA files. Two searches were conducted, one each against the yeast and grape protein databases described above. For yeast searches cleavage specificity was set as C-terminal to Arg/Lys and semi-specific (one terminus can disagree), a maximum of two missed cleavages were allowed, and mass tolerances of 20 ppm and 50 ppm were applied to precursor and fragment ions, respectively. Variable modifications set as “Common 1” allowed each modification to be present on a peptide once and included mono-oxidised Met and deamidated Asn. Dehydro Cys was set as “Common 2”, which allowed the modification to be present twice on a peptide. The setting “Rare 1”, which allowed each modification to be present once on a peptide, included the *N*-linked monosaccharide compositions HexNAc_1-2_ and HexNAc_2_Hex_1-15_ at the consensus sequence N-X-S/T and the *O*-linked monosaccharide compositions Hex_1_-Hex_10_ at any Ser or Thr residue (HexNAc, *N*-acetylhexosamine; Hex, hexose). A maximum of two common modifications and one rare modification were allowed per peptide.

For grape searches, cleavage specificity was set as semi-specific, a maximum of one missed cleavage event was allowed, and mass tolerances of 20 ppm and 50 ppm were applied to precursor and fragment ions, respectively. Oxidation of Pro was set as “Common 4”, which allowed each modification to be present four times on a peptide. The “Rare 1” setting included 52 plant *N*-glycans (**Supplementary Table S1**) with 17 monosaccharide compositions containing pentose (Pent) at the consensus sequence N-X-S/T. The *O*-linked monosaccharide compositions searched were Pent_1_-Pent_8_ + 15.9949 at any Pro residue. A maximum of four common modifications and one rare modification were allowed per peptide.

Unique glycopeptides identified in the Byonic searches were manually inspected and validated. For an assignment to be accepted, relevant oxonium ions had to be present in the spectrum to confirm the monosaccharide composition of the attached glycan. The exception to this was the presence of deoxyhexose (dHex) or Pent in an *N*-glycan. Where possible, monosaccharide presence was confirmed through glycopeptide Y ions. In addition, the glycopeptide Y0 ion had to be present for *O*-glycopeptides, and the Y1 ion for *N*-glycopeptides, as well as either *b* or *y* peptide ions (≥3). If peptide *b-* and *y-*ions were low in intensity, retention time information and elution of other glycoforms was used to validate identifications. If there were no Y0 or Y1 ions, then both *b* and *y* peptide ions had to be present (≥3 in total). Peptides from cleavage events at Lys/Arg-Pro or N-ragged cleavage were accepted only if and *b* and *y* peptide ions were present (≥3) in addition to Y0 for *O*-glycopeptides, or Y1 ions for *N*-glycopeptides. The best unique glycopeptide identification across all searches were used to create a glycopeptide library for SWATH/DIA analyses. The debug mode of Byonic was used to export details of the identified glycopeptides. We wrote a Python script that created a PeakView SWATH library from the exported details. An overview of the data used to populate each column of the PeakView library can be found in **Supplementary Table S2**. The Byonic debug report file is grouped by identified peptides. Our script matched scan numbers from the list of validated glycopeptides to those in the relevant report files and returned the corresponding precursor charge and theoretical precursor mass. The theoretical precursor *m/z* (Q1) was then calculated. Fragment ion information was also collected, including fragment type (Y0, Y1, Y2, b and y), fragment position within the peptide sequence (e.g. b1 or y6), theoretical fragment ion *m/z* (Q3), fragment ion charge, and fragment ion intensity. MS/MS spectra of glycopeptides are more complex than of peptides, owing to additional fragment ions from the attached glycan. To accommodate this complexity within the structure of a PeakView ion library, we assigned as “a” ions glycopeptide fragment ions Y0, Y1, and Y2 as well as b- and y-ions with HexNAc_1_ from position four and above within the peptide sequence (Table 1). This meant b- and y-ions with HexNAc_1_ at positions 1-3 within the peptide sequence were not included in the DIA library. Peptide b- and y-ions with the entire glycan still attached were assigned as “c” ions in the ion library (Table 1). It is important to note that these assignments did not represent actual designated a- and c-ions as defined by peptide fragmentation nomenclature (37). The combined debug file was merged with the information from the manually validated result file. The peptide-to-spectrum match information taken from the result file included the glycan monosaccharide composition, peptide sequence (plus the mass of the modification), retention time and protein name. Peptide modifications were removed from the peptide sequence to produce the stripped sequence. The library was then checked for duplication of transitions before being exported as a tab delimited .txt file compatible with import into PeakView.

**Table 1.**
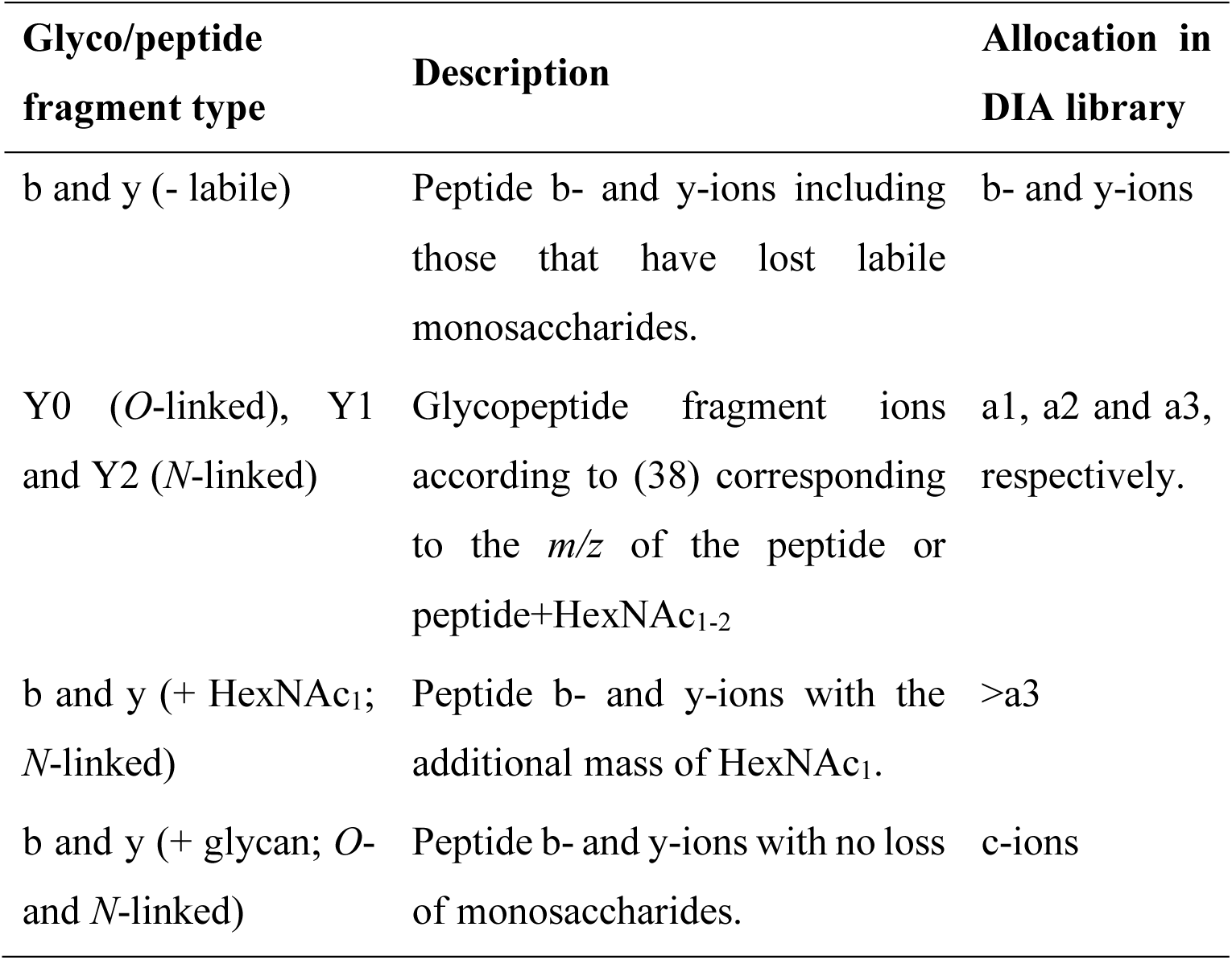
Allocation of ion-types for the glycopeptide SWATH/DIA library.

Identified peptides from the ProteinPilot search were combined with glycopeptides identified from the Byonic search to form one ion library. The library was used to measure peptide abundances in PeakView v2.2.0.11391 (SCIEX) using the SWATH Acquisition MicroApp. Settings included: number of peptides per protein, unlimited; transitions per peptide, 6; peptide confidence threshold, 99%; false discovery rate, 1%; shared peptides, allowed; retention time XIC window, 6 min; XIC width, 75 ppm. The extracted ion chromatograms for MS2 fragment ions for all glycopeptides were manually inspected.

For protein-centric analyses, changes in total protein abundance between samples was determined with: normalised protein intensities to the reference protein trypsin, as previously described (33); MSstats in R, as described previously (34); or principal component analysis (PCA) and clustered heatmaps with ClustVis (39) using protein abundances normalised to all yeast and grape protein abundances (34). For glycopeptide-centric analyses, glycoform abundances were normalised to the summed abundance of all detected forms of the same peptide (33). For both protein-centric and glycopeptide-centric analyses a peptide FDR cut-off of 1% was applied. Heatmaps were produced using PRISM v7.00 for Mac OS X (GraphPad Software, La Jolla California USA) and spectra for manual annotation were plotted using SciDAVis v1.25.

### Data Availability

The mass spectrometry proteomics data have been deposited to the ProteomeXchange Consortium via the PRIDE (40) partner repository with the dataset identifier PXD019572.

## RESULTS

Given the limited number of studies investigating the proteome of sparkling wine (41) and the likely importance of proteins in determining wine quality, we set out to conduct comprehensive, quantitative analyses of the proteomes of sparkling wines produced under different conditions. Unexpectedly, preliminary LC-MS/MS proteomic investigations of tryptic digests of sparkling wines revealed that the dominant components in these samples were glycopeptides (**Fig. 2**), with precursor masses separated by 162 Da (Hex) and 132 Da (Pent), and MS/MS spectra containing ions corresponding to oxonium ions for hexose and pentose. The presence of glycopeptides was not unexpected, as glycoproteins from grapes are released during pressing and secreted from yeast during primary and secondary fermentation (**Fig. 1**). However, the abundance of the glycopeptides was surprising, given that glycopeptide analysis generally requires enrichment during sample preparation (42-44). Glycoproteins are important for controlling the clarity of all wines (7-9) and are of particular importance in foam production specifically in sparkling wines [reviewed in (3)], but very little is known about protein-specific glycosylation and changes in glycoprotein content during wine and sparkling wine production. This prompted us to focus our analysis on the glycoproteome of sparkling wine.

**Figure 2.**
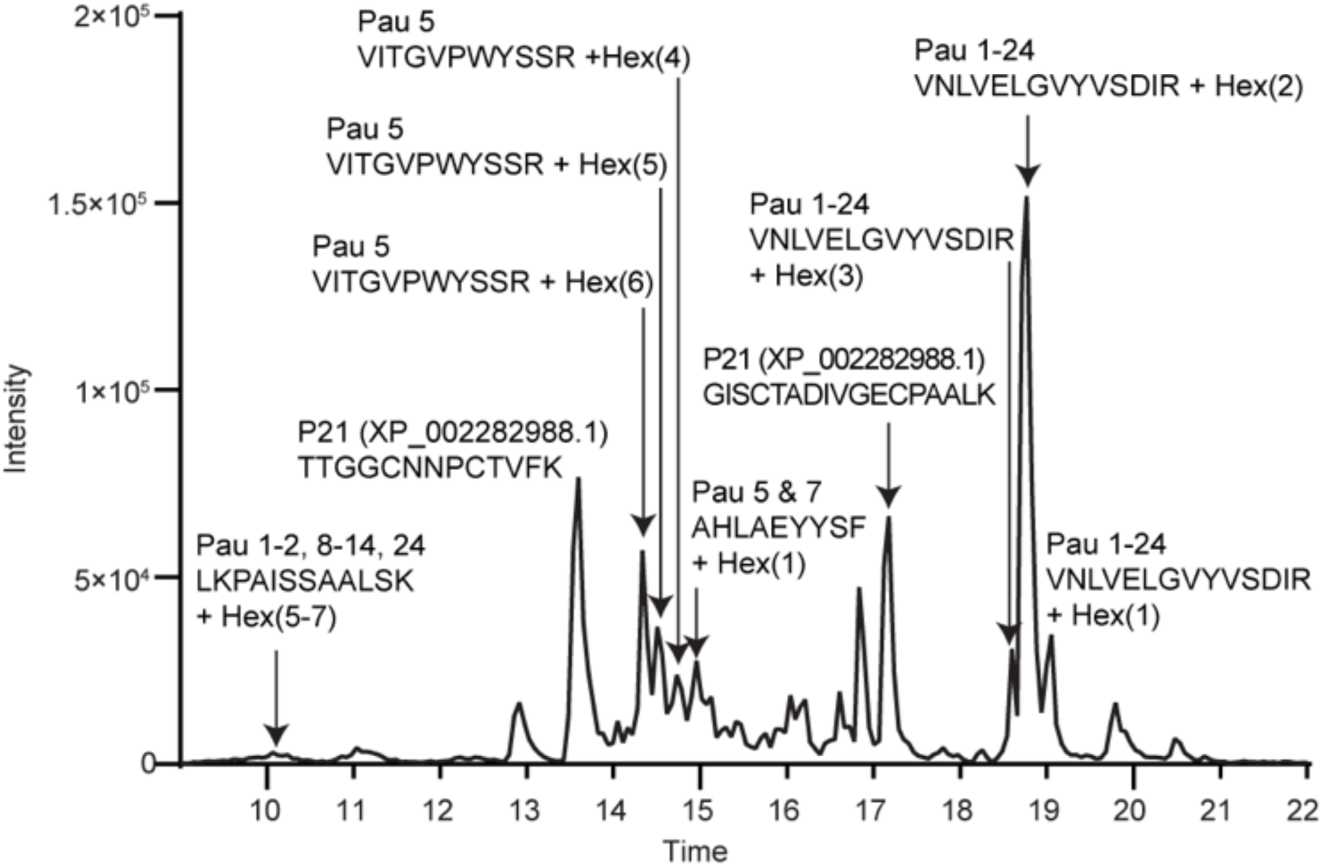
LC-MS/MS proteomics shows glycopeptides are abundant in sparkling wine. MS base peak chromatogram of a tryptic digest of cuvée aged 8 months depicting the retention time profile of the most intense peptides and glycopeptides eluting throughout the LC-MS/MS experiment. Pau, Seripauperin. Details of peptide and glycopeptide identifications are shown in Fig. 3, Supplementary Fig. S1, and Supplementary Table S4.

### Construction of a glycopeptide ion library

To allow efficient measurement of site-specific glycosylation with facile robust quantification, we developed a workflow integrating glycopeptide identification by Byonic with SWATH/DIA glycopeptide quantification. To create a SWATH/DIA library to allow measurement of glycopeptides by SWATH/DIA, we wrote a Python script to retrieve precursor and fragment ion data from the debug mode of Byonic (see methods for details). Tryptic digests of sparkling wine samples were measured with DDA LC-MS/MS, searched with Byonic, identified glycopeptides manually validated, and the data processed with our pipeline. The resulting SWATH/DIA library contained 142 Q1 values (precursors) and 2,849 transitions (fragment ions) for glycopeptides from 16 proteins (**Supplementary Table S3**). The 142 Q1 values represented glycopeptide precursors with different monosaccharide compositions, peptide sequences, and charge states (104 unique monosaccharide compositions and peptide sequences) (**Supplementary Table S4** and annotated spectra in **Supplementary Fig. S1**). Of the 104 unique glycopeptides identified, 93 were *O*-linked from yeast, and six *O*-linked and five *N*-linked from grape. Not unexpectedly, glycans from yeast and grape had distinct monosaccharide compositions. Glycoproteins from yeast have high mannose *N*-glycans and oligomannose *O*-glycans (19), while glycoproteins from grape have high mannose or paucimannose *N*-glycans (20) and in sparkling wine hydroxyproline linked arabinose and/or galactose *O*-glycans (21,45). The yeast *O*-linked glycopeptides we identified contained 1-9 Hex residues which were most likely mannose (representative spectra in **Fig. 3A and 3B**) while grape *O*-linked glycopeptides contained 1-6 Pent resides, most likely arabinose (representative spectrum in **Fig. 3C**). The detected grape *N*-linked glycopeptides contained a single HexNAc residue or the monosaccharide compositions HexNAc_2_Hex_3_dHex_1_Pent_1_ or HexNAc_3_Hex_3_dHex_1_Pent_1,_ (**Supplementary Fig. S1 slides 132-136**). Many of the peptides and glycopeptides we detected were semi-tryptic, suggesting that other yeast or grape proteases were active during sparkling wine production or storage.

**Figure 3.**
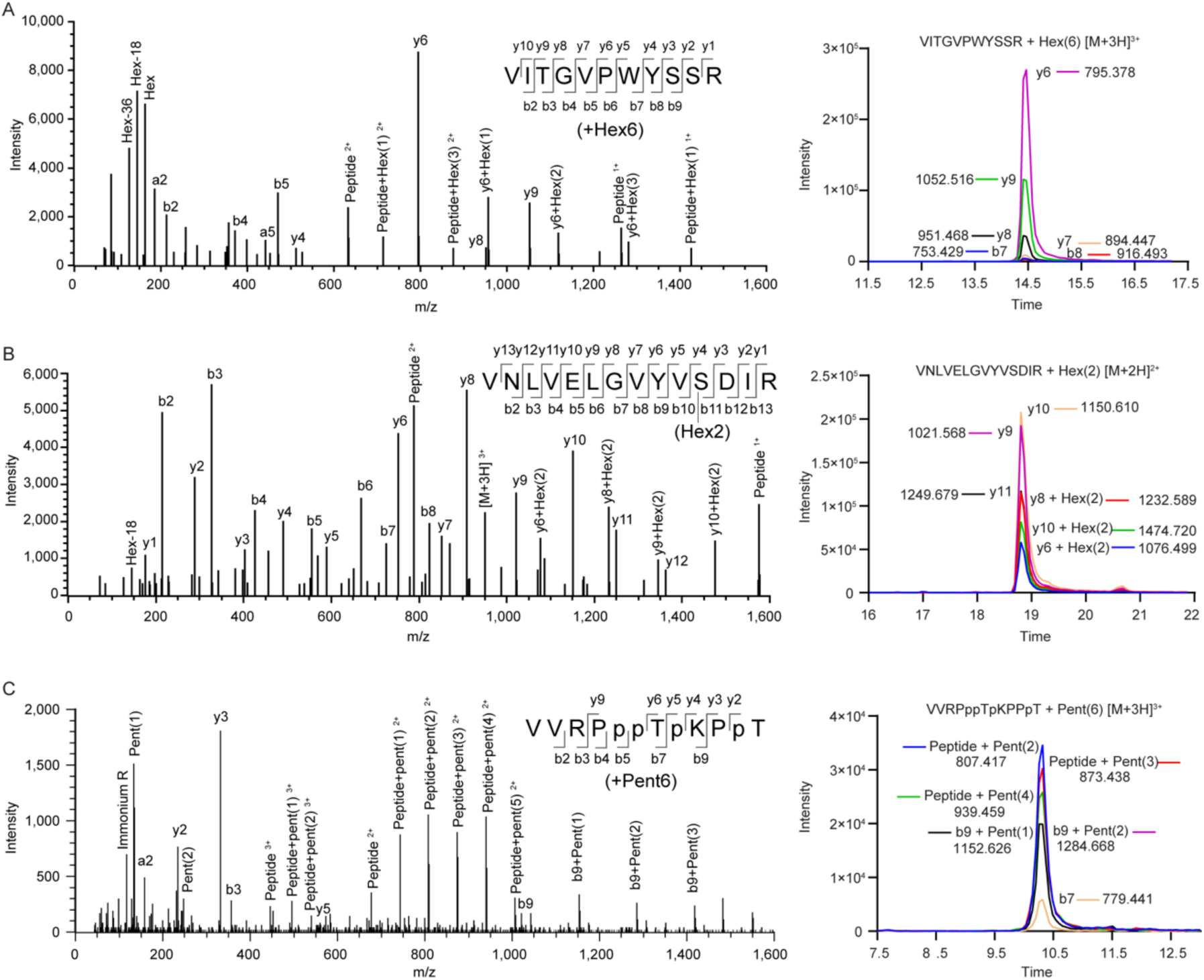
Representative data-dependent MS/MS spectra of yeast and grape *O*-glycopeptides in sparkling wine. Each panel contains a schematic of the fragmentation pattern of the glycopeptides; not all ions have been labelled in the spectra for ease of interpretation. Each panel also includes a representative extracted ion chromatogram of the MS2 fragment ions used for quantification by DIA. (**A**) VITGVPWYSSR from Pau5 (UniProt ID P43575) with six hexoses attached at *m/z* 746.3205 (3+) with a -17.15 ppm precursor mass error. (**B**) Peptide VNLVELGVYVSDIR from Seripauperins 1-24 with two hexoses attached at *m/z* 950.4798 (3+) with a -14.42 ppm precursor mass error. Spectra for the *O*-linked peptides from A & B were from the analysis of sparkling cuvée aged 8 months. (**C**) Peptide VVRPppTpKPpT from a putative proline-rich cell wall protein (NCBI Reference NP_001268136.1) found in Sauvignon blanc fermented with the yeast DV10 aged 16 months. The glycopeptide with six pentoses attached was observed at *m/z* 714.6610 (3+) with a -12.74 ppm precursor mass error. Lowercase prolines represent likely sites of hydroxyproline.

### Changes in protein and glycoprotein abundances

To investigate how the glyco/proteome of sparkling wine was affected by the style of production, we used our analytical workflow to investigate three pairs of commercial scale experimental sparkling wines: a cuvée that was aged for 8 or 24 months on lees; a Sauvignon blanc that underwent a second fermentation with DV10 or Zymaflore yeasts; and a Riesling that underwent a second fermentation with DV10 or Siha4 yeasts. To assess differences in both protein and glycoprotein abundances we combined our Byonic-derived glycopeptide library with a proteomic library from ProteinPilot searches (**Supplementary Table S5 and Supplementary Fig. S2**). We identified 35 proteins across all six experimental conditions. Of the 35 proteins, 17 were from *V. vinifera* and 18 were from *S. cerevisiae* and 12 proteins were identified through glycopeptides only (11 of these belonging to yeast). Using the combined library, 130 glycopeptide Q1 values and 219 peptide Q1 values were reliably measured in PeakView and used to calculate the abundance of 34 yeast and grape proteins.

We used the abundance data for the 34 yeast and grape proteins to investigate total protein abundance in each sample (**Fig. 4A**) and statistically significant log-fold-changes in protein abundance between wines (**Fig. 4B**). Seripauperins were the most abundant yeast proteins in all samples. The five seripauperin (Pau) proteins observed were all identified with at least one unique peptide. However, several peptides are common to all seripauperins, so unambiguous abundance information was not able to be obtained for individual gene products. We first investigated proteomic differences in cuvée wine aged for 8 or 24 months. This analysis showed a dramatic decrease in total protein abundance at 24 months compared to only 8 months on lees (**Fig. 4A**), consistent with a general loss of solubility of all proteins during this extended aging process. PCA analysis revealed clustering of the technical replicates and clear separation of samples from 8 and 24 months on lees (**Fig. 4C**). We next compared Sauvignon blanc base wine that underwent a second fermentation with either DV10 or Zymaflore yeasts. This comparison showed that there was little change in total protein content (**Fig. 4A**), and that relatively few proteins were differentially abundant depending on the yeast used for secondary fermentation (**Fig. 4B**). PCA analysis (**Fig. 4C**) confirmed the close association of the proteomes. Finally, we compared Riesling base wine that underwent a second fermentation with either DV10 or Siha4 yeasts. This analysis showed yeast proteins were generally lower in abundance relative to grape proteins after fermentation with DV10 compared with Siha4 (**Fig. 4A and 4B**). PCA (**Fig. 4C**) confirmed that the proteome of the Riesling fermented with DV10 was distinct from the Riesling fermented with Siha4 and was in fact more closely associated with the proteome of the cuvée aged 24 months.

**Figure 4.**
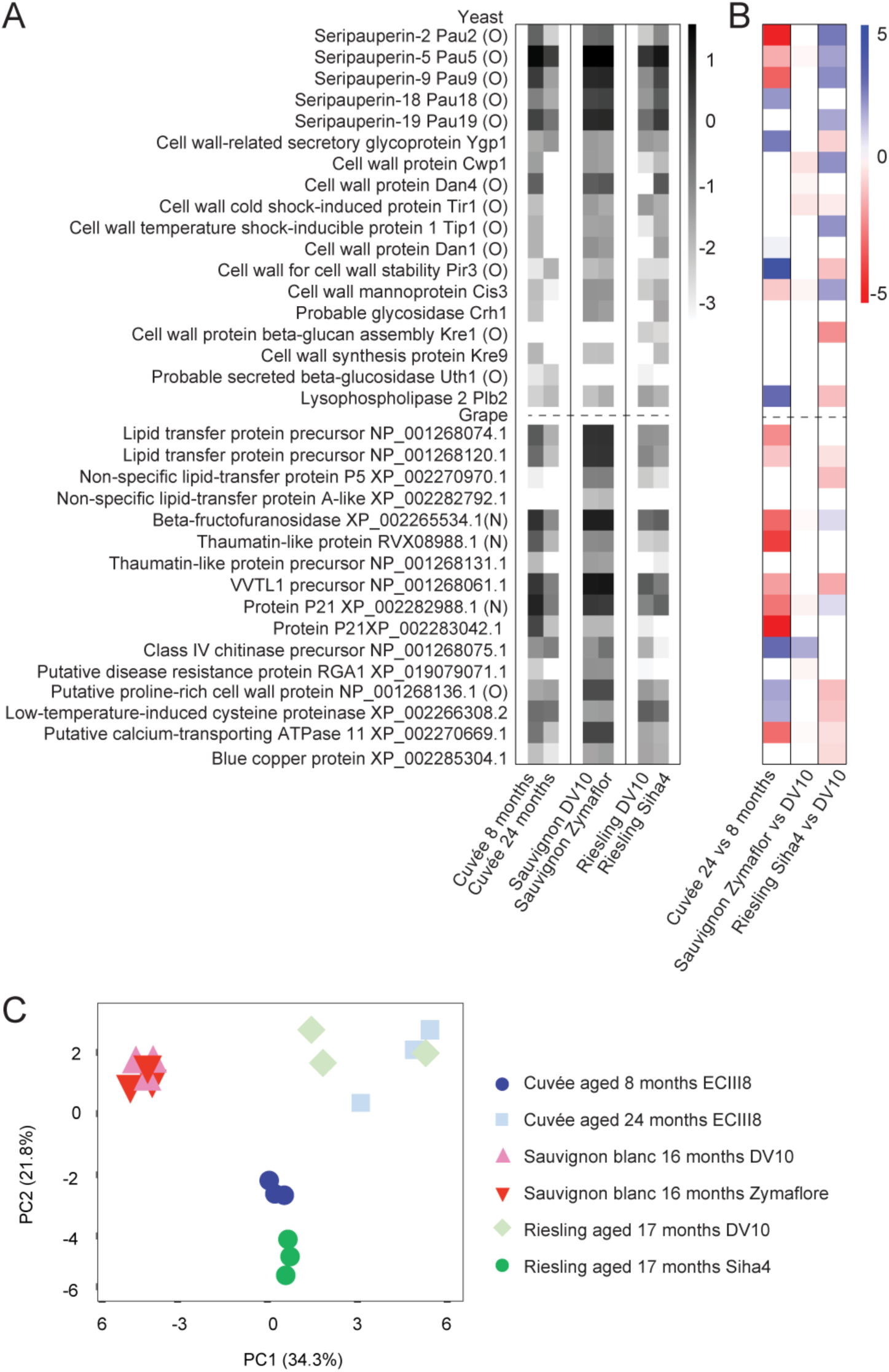
Protein-level analyses of sparkling wine. (**A**) Heatmap of the abundance of proteins normalised to trypsin. Values are the log of the mean of technical triplicates. (**B**) Heatmap of differentially abundant proteins (p < 10^−5^) between experimental groups. Values show the log_2_ fold difference in cuvée aged 24 months relative to 8 months, Sauvignon blanc aged 16 months fermented with Zymaflore relative to DV10 and Riesling aged 17 months fermented with Siha4 relative to DV10. Proteins names are listed in (A). Yeast proteins are displayed above the dotted line and grape below. Proteins labelled with (O) or (N) were observed with *O*-linked and *N*-linked glycans, respectively. (**C**) Principal component analysis (PCA) of sparkling wine proteomic and glycoproteomic profiles. Each point represents a technical replicate (n=3) from the experimental groups. The graph shows two principal components (PC1 and PC2) with the variance associated with each component in parentheses.

### Changes in glycoform abundances

We next made use of our rich glycoproteomic dataset to investigate any differences in the occupancy or composition of site-specific glycoforms between wines. We calculated glycoform abundances from high-confidence (glyco)peptide-level data (FDR cut-off of 1%) and calculated the relative abundance of glycoforms by the fraction of the summed intensity of all forms of peptides or glycopeptides in all charge states containing the site(s) of glycosylation. Similar to the protein-level profiles, a clustered heatmap (**Fig. 5A**) and PCA (**Fig. 5B**) confirmed the glycoforms of the cuvée aged 24 months clustered separately from the cuvée aged 8 months. This global glycoproteomic analysis identified nine glycoforms that were significantly different in abundance in the cuvée aged 24 months relative to 8 months, with the largest differences apparent between glycoforms of the VITGVPWYSSR/L peptide from Pau5 (**Fig. 5C**). Close inspection of these data confirmed an enrichment in the more highly glycosylated glycoforms of this peptide in the aged cuvée (**Fig. 6A**). In contrast, little difference was observed in the abundance of glycoforms from the glycopeptides ER/VN/LVELGVYVSDIR shared in all 24 seripauperins (**Fig. 6D**). Comparison of the glycoproteomes of Sauvignon blanc fermented with DV10 or Zymaflore yeasts by heatmap and PCA revealed the glycoforms of both conditions were closely associated (**Fig. 5A and 5B**) and there were no significant differences in the abundance of any individual glycoforms (**Fig. 5C and Fig. 6B and 6E)**. Consistent with protein-level analysis, clustered heatmap (**Fig. 5A**) and PCA (**Fig. 5B**) confirmed glycoforms of the Riesling fermented with DV10 were distinct from those derived from fermentation with Siha4 and clustered more closely with the cuvée aged 24 months. Nine glycoforms were significantly different in the Riesling fermented with Shia4 relative to DV10 (**Fig. 5C**). We observed more highly glycosylated species of the glycopeptides VITGVPWYSSR/L (Hex_2-9_) in the Riesling fermented with DV10 (**Fig. 6C**) and little difference was observed in the abundance of glycoforms from the glycopeptides ER/VN/LVELGVYVSDIR (**Fig. 6F**). We also observed differences in site-specific glycosylation between sparkling wine from different grape varieties. The glycoforms of the grape glycopeptide VVRPPPTPKPPT were more heterogeneous in the Sauvignon blanc samples compared to cuvée and Riesling (**Fig. 6G**). In summary, this glycoform abundance analysis showed that larger *O*-glycoforms on some yeast proteins were enriched in aged cuvée, there were limited differences in the glycoproteomes of Sauvignon blanc fermented with DV10 or Zymaflore yeasts, fermentation of Riesling with DV10 or Shia4 yeasts resulted in different yeast O-glycosylation profiles, and glycosylation of grape proteins was distinct in Sauvignon blanc compared to cuvée and Riesling.

**Figure 5.**
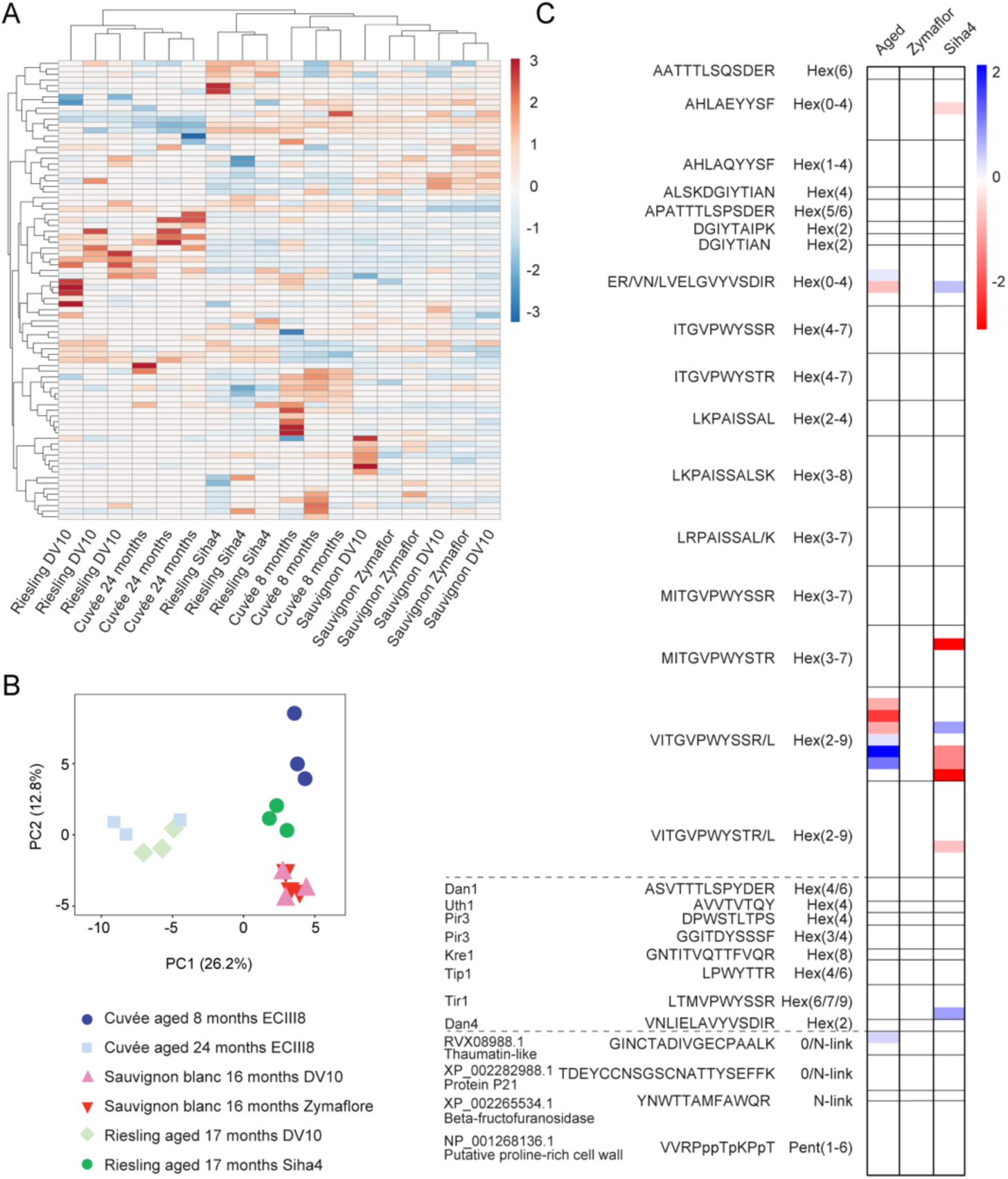
Sparkling wines show diverse glycoproteomes. (**A**) Clustered heatmap of the relative abundance of glycoforms in all replicates. Both rows and columns were clustered using correlation distance and average linkage. The full clustered heatmap with labelled rows (glycopeptide glycoforms) can be found in **Supplementary Figure S3**. (**B**) Principal component analysis (PCA) of sparkling wine glycoform abundances. Each point represents a technical replicate (n=3) from the experimental groups. The graph shows two principal components (PC1 and PC2) with the variance associated with each component in parentheses. (**C**) Heatmap of differentially abundant glycoforms (two-tailed t-test, P<0.05). Values show the log_2_ fold change in cuvée aged 24 months relative to 8 months, Sauvignon blanc aged 16 months fermented with Zymaflore relative to DV10 and Riesling aged 17 months fermented with Siha4 relative to DV10. Yeast seripauperin glycoforms are displayed above the first dotted line, the remaining yeast glycoforms are between the dotted lines, and grape glycoforms are below the lower dotted line.

**Figure 6.**
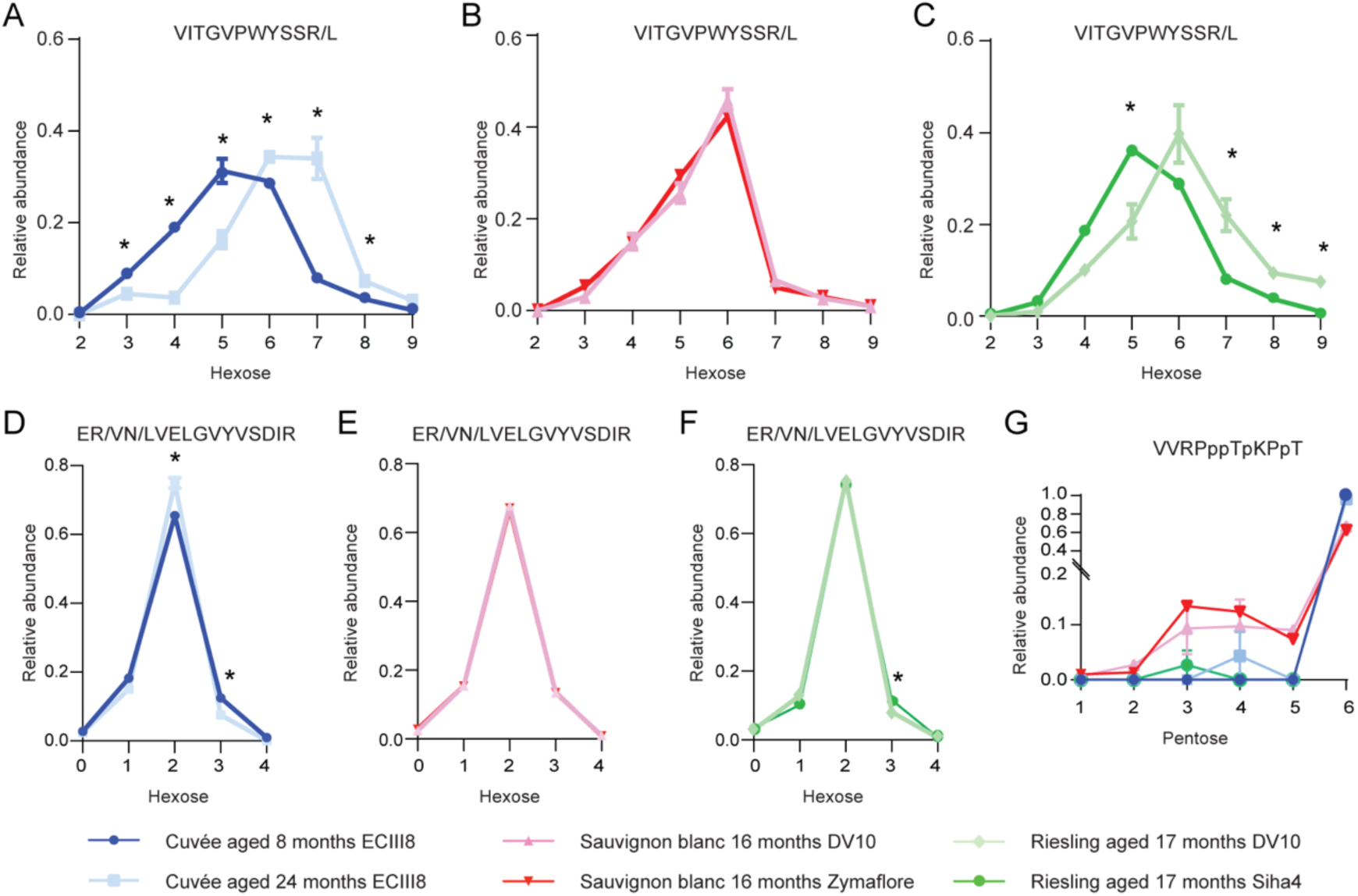
Site-specific glycosylation profiles differ between experimental sparkling wines. Relative abundances of Hex_2-9_ glycoforms of VITGVPWYSSR/L peptides from yeast Pau5 (UniProt ID P43575) in (**A**) Cuvée aged 8 or 24 months on lees, (**B**) Sauvignon blanc fermented with DV10 or Zymaflore, and (**C**) Riesling fermented with DV10 or Siha4. Relative abundances of unmodified and Hex_1-4_ glycoforms of ER/VN/LVELGVYVSDIR peptides from yeast Seripauperins 1-24 in (**D**) Cuvée aged 8 or 24 months on lees, (**E**) Sauvignon blanc fermented with DV10 or Zymaflore, and (**F**) Riesling fermented with DV10 or Siha4. (**G**) Relative abundances of Pent_1-6_ glycoforms of VVRPppTpKPpT peptide from grape proline-rich cell wall protein-like precursor (NCBI NP_001268136.1). Lowercase proline represents the likely sites of hydroxyproline (typical spectrum shown in Figure 3C). Abundances are represented as a fraction of the summed intensity of all forms of peptides or glycopeptides in all charge states containing the site(s) of glycosylation. Values represent the mean of technical triplicates. Error bars show standard deviation and significance is indicated by * (two-tailed t-test, P<0.05).

## DISCUSSION

Sparkling wine producers are continually investigating production and analytical methods to improve the quality and robustness of the final product. An improved understanding of how changes in the production of sparkling wines affect its molecular composition could identify key proteins and their modifications that promote desired qualities such as clarity, aroma, and foamability. Here, we describe a robust mass spectrometry based method that enables quantification of proteins, glycoproteins, and their glycosylation profiles from small amounts of sparkling wine. We developed an automated workflow that created a library of transitions from glycopeptides identified in Byonic searches for SWATH/DIA analysis. To exemplify the use of this method, we applied it to three experimental wines, investigating the effect of ageing on lees and how different yeast strains in the Liqueur de Tirage alter the glycoproteomic and proteomic profile of mature sparkling wines. We investigated a cuvée that was aged for 24 months on lees compared to 8 months. A Sauvignon blanc that underwent a secondary fermentation with either DV10 or Zymaflore yeasts, and a Riesling that underwent a secondary fermentation with either DV10 or Siha4 yeasts.

In this study we identified 35 proteins (17 from *V. vinifera* and 18 from *S. cerevisiae*) using 250 µL of sparkling wine for each replicate. Although the number of identified proteins is low, it is important to note that only a small amount of protein is found in sparkling wine (4-20 mg/L) (6). Other qualitative proteomic studies investigating base wine and sparkling wine implemented protein concentration methods yet still only identified a limited number of proteins (3,41). For example, studies of untreated champagne base wine (200 mL) or Chardonnay still wine (15 mL) identified 9-l3 grape proteins and up to 15 yeast proteins (46,47). Another study of untreated Recioto wine using combinatorial peptide ligand libraries (CPLLs) with four pH conditions used 750 mL of starting material for each pH condition (48). This study identified 106 *V. vinifera* and other viridiplantae proteins and 11 yeast proteins. A separate in-depth study of sparkling wine using CPLLs with 750 mL of starting material identified 12 grape proteins and 7 yeast proteins (41). Our results are therefore consistent with the low concentration and complexity of the sparkling wine proteome, and demonstrate that in-depth analysis of the wine glyco/proteome is possible with small 250 µL sample volumes.

As glycopeptides were abundant in sparkling wine (**Fig. 2**), we included measurement of peptides and glycopeptides in our protein-centric analyses. This is a substantial improvement on previous studies of wine and sparkling wine, which neglected the substantial contribution of glycopeptides to the overall proteome. This protein-level analysis revealed a decrease in total protein abundance in cuvée aged 24 months compared to 8 months (**Fig. 4A**). Yeast cell viability and protein secretion decrease after the first few months of the second fermentation in sparkling wine production (28), and after 9 months of aging typically no viable yeast cells remain (1). The further decrease in protein content we observed at 24 months is likely due to ongoing protein aggregation (31) or degradation by yeast enzymes (24,49,50). For example, grape thaumatin-like proteins and chitinases are particularly susceptible to hydrolysis by proteolytic activity by *S. cerevisiae* Pir1 (51). Surprisingly, even after extended aging on lees we only identified bona fide secreted yeast proteins, rather than intracellular proteins, suggesting that the impact of yeast autolysis on the aged sparkling wine proteome is not as extensive as previously thought. Our protein-centric analyses also revealed very little change in protein abundances in Sauvignon blanc fermented with the different yeast strains DV10 and Zymaflore, and detected a decrease in yeast proteins in Riesling fermented with DV10 compared to Siha4. During the secondary fermentation, different yeast strains vary in growth kinetics, viability, and autolytic properties which can alter the protein and free monosaccharide content in sparkling wine (28). Such differences in yeast properties and activity may account for the lower total yeast protein content we observed in Riesling fermented with DV10.

We found seripauperins were the most abundant yeast glycoproteins present in sparkling wine (**Fig. 4A**). *PAU5* expression is induced at low temperatures and with low oxygen, similar to conditions used during winemaking (52). Pau5 has also been identified as a glycoprotein that may reduce spontaneous over-foaming of sparkling wine (12,14), a phenomenon known as gushing, which is perceived negatively by consumers. The absence or undetectable levels of Pau5 is associated with a high probability of gushing (14) while the addition of purified native Pau5 from non-gushing sparkling wine stabilises foam formation (12).

In addition to protein-centric abundances we also quantified the relative abundance of glycoforms by measuring the intensities of glycopeptides (**Fig. 5 and Fig. 6**). To do this, we created a SWATH/DIA library of transitions for glycopeptides after identifying glycopeptides by searching DDA files using Byonic software. To our knowledge, our study is the most in-depth investigation of the glycosylation profile of proteins in wine or sparkling wine (12,14,20,46). Other groups have investigated protein glycosylation after removal of the glycans in non-sparkling Chardonnay, identifying 44 *N*-linked sites (46). We identified one of these sites in our work (site N184, protein P21 or thaumatin-like protein, XP_002282988.1) with two alternate glycoforms attached. Another group studied purified Pau5 from white sparkling wine (12,14). Similar to our results at a glycopeptide level, the authors noted modifications of 162 Da (hexose) to the protein after intact analysis (14). Finally, *N*-linked glycosylation of purified grape vacuolar invertase isolated from grape must (produced after pressing the grapes) has been investigated (20). The study identified 10 sites after deglycosylation, one of which was also identified in our work with one glycoform attached (site N101, beta-fructofuranosidase, XP_002265534.1).

We identified more glycopeptides from yeast origin than from grape, 93 compared to 11, suggesting the dominant glycoproteins present in sparkling wine originate from yeast. This also highlights the importance of secondary fermentation in providing carbonation and flavour compounds to sparkling wine, but also how ageing can contribute more complex biomolecules that are likely to affect its complex sensory features. This is consistent with research identifying mannose as the main free monosaccharide present in sparkling wine (28). Our glycopeptide-centric data revealed some glycopeptides with larger glycans are enriched in aged cuvée. This effect could be due to increased solubility provided by the larger glycans reducing the propensity of the protein to aggregate, or protection from low levels of protease activity (**Fig. 6A-C**). However, large glycans were not enriched at all sites after aging, suggesting the precise sites of glycosylation in proteins are critical for determining their function in affecting stability during wine aging (**Fig. 6D-F**).

In summary, we have developed a sensitive method that can be used to identify and measure glycoproteins and their site-specific modifications to investigate their importance to the organoleptic properties of sparkling wine. The development of a protein- and glycoform-specific quantitative method from small volumes of sparkling wine will be highly useful and complementary to other tools that measure the sensory properties of sparkling wine to pinpoint specific proteins that correlate with desirable outcomes. The small-scale quantitative workflow that we have developed could also be applied to still or base wines to investigate the features of their proteomes and glycoproteomes that contribute to wine quality. More broadly, the workflow we have developed for SWATH/DIA glycoproteomics will be applicable to the study of other diverse complex glycoproteomes.

## ACKNOWLEDGMENTS

We thank Dr Amanda Nouwens and Peter Josh at The University of Queensland, School of Chemistry and Molecular Biosciences Mass Spectrometry Facility for their assistance and expertise. The authors sincerely thank the Schloss Vaux winery for providing the experimental wines and encouraging publication of these results and Professor Dr Mark Strobl and Shraddha More from Hochschule Geisenheim University, Germany who enabled access to the wines.

## Notes

### Competing Interest Statement

The authors have declared no competing interest.

