## Supplementary Figure S1 for "Quantitative data independent acquisition glycoproteomics of sparkling wine"

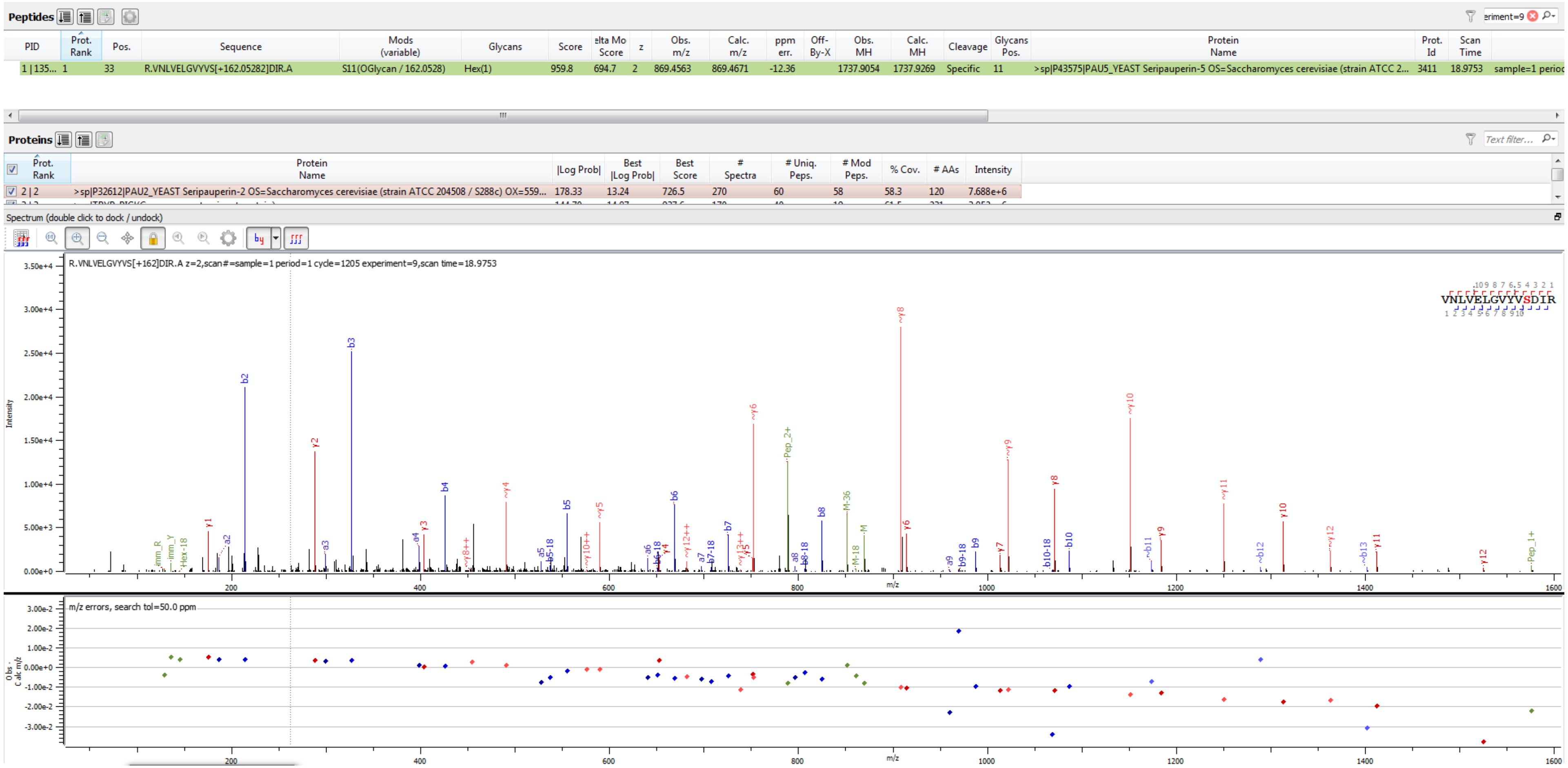

Peptides

| PID | Prot. Rank | Pos. | Sequence | Mods (variable) | Glycans | Score | elta Mo Score | z | Obs. m/z | Calc. m/z | ppm err. | Off-By-X | Obs. MH | Calc. MH | Cleavage | Glycans Pos. | Protein Name | Prot. Id | Scan Time | |
|---|---|---|---|---|---|---|---|---|---|---|---|---|---|---|---|---|---|---|---|---|
| 1 \| 135... | 1 | 33 | R.VNLVELGVYVS[+162.05282]DIR.A | S11(OGlycan / 162.0528) | Hex(1) | 959.8 | 694.7 | 2 | 869.4563 | 869.4671 | -12.36 | | 1737.9054 | 1737.9269 | Specific | 11 | >sp\|P43575\|PAU5_YEAST Seripauperin-5 OS=Saccharomyces cerevisiae (strain ATCC 2... | 3411 | 18.9753 | sample=1 perioc |

Proteins

| Prot. Rank | Protein Name | \|Log Prob\| | Best \|Log Prob\| | Best Score | # Spectra | # Uniq. Peps. | # Mod Peps. | % Cov. | # AAs | Intensity |
|---|---|---|---|---|---|---|---|---|---|---|
| 2 \| 2 | >sp\|P32612\|PAU2_YEAST Seripauperin-2 OS=Saccharomyces cerevisiae (strain ATCC 204508 / S288c) OX=559... | 178.33 | 13.24 | 726.5 | 270 | 60 | 58 | 58.3 | 120 | 7.688e+6 |



Sample S2 R.VNLVELGVYVS[+162.053]DIR.A

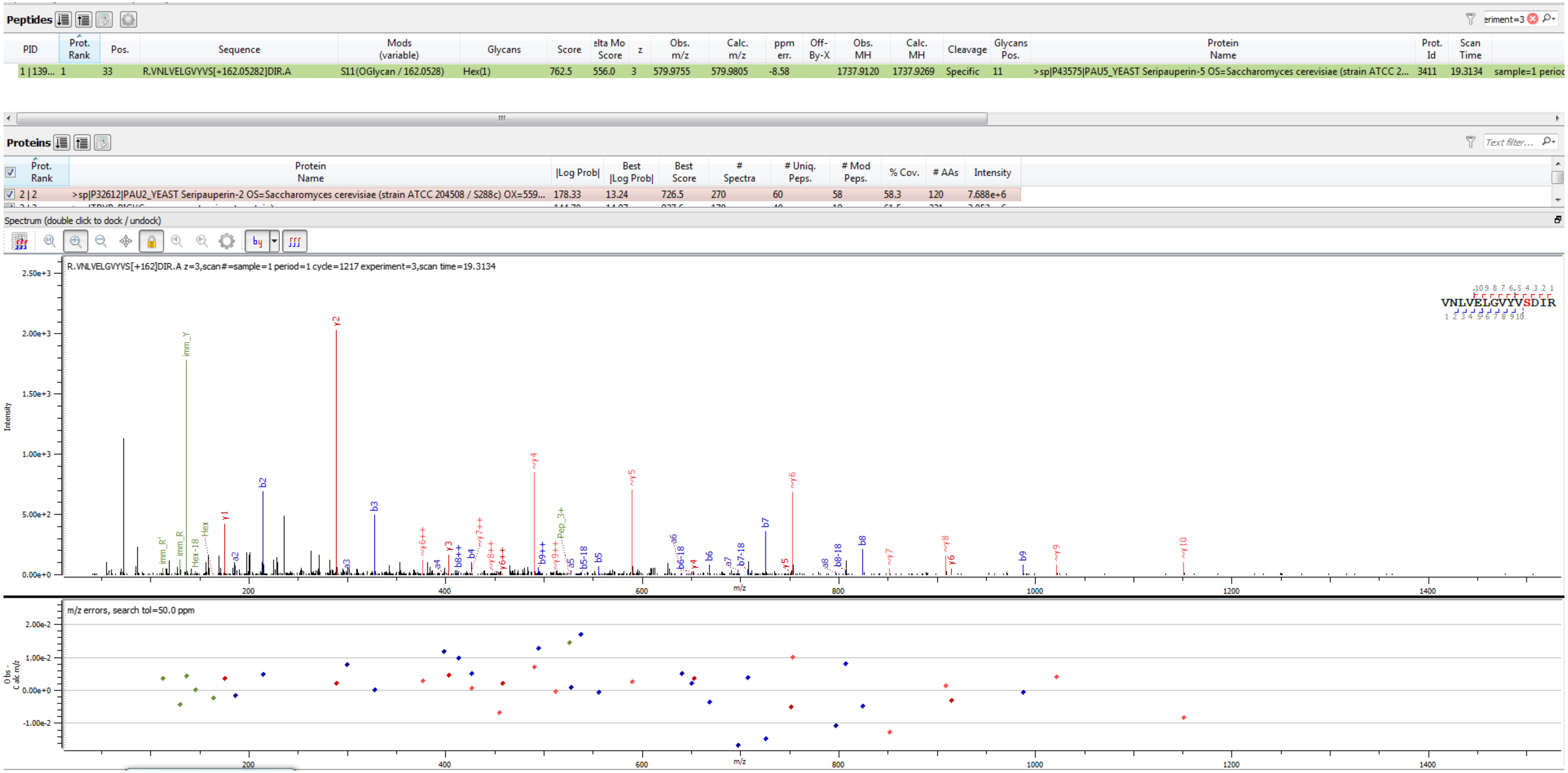


Sample S2 R.VNLVELGVYVS[+162.053]DIR.A

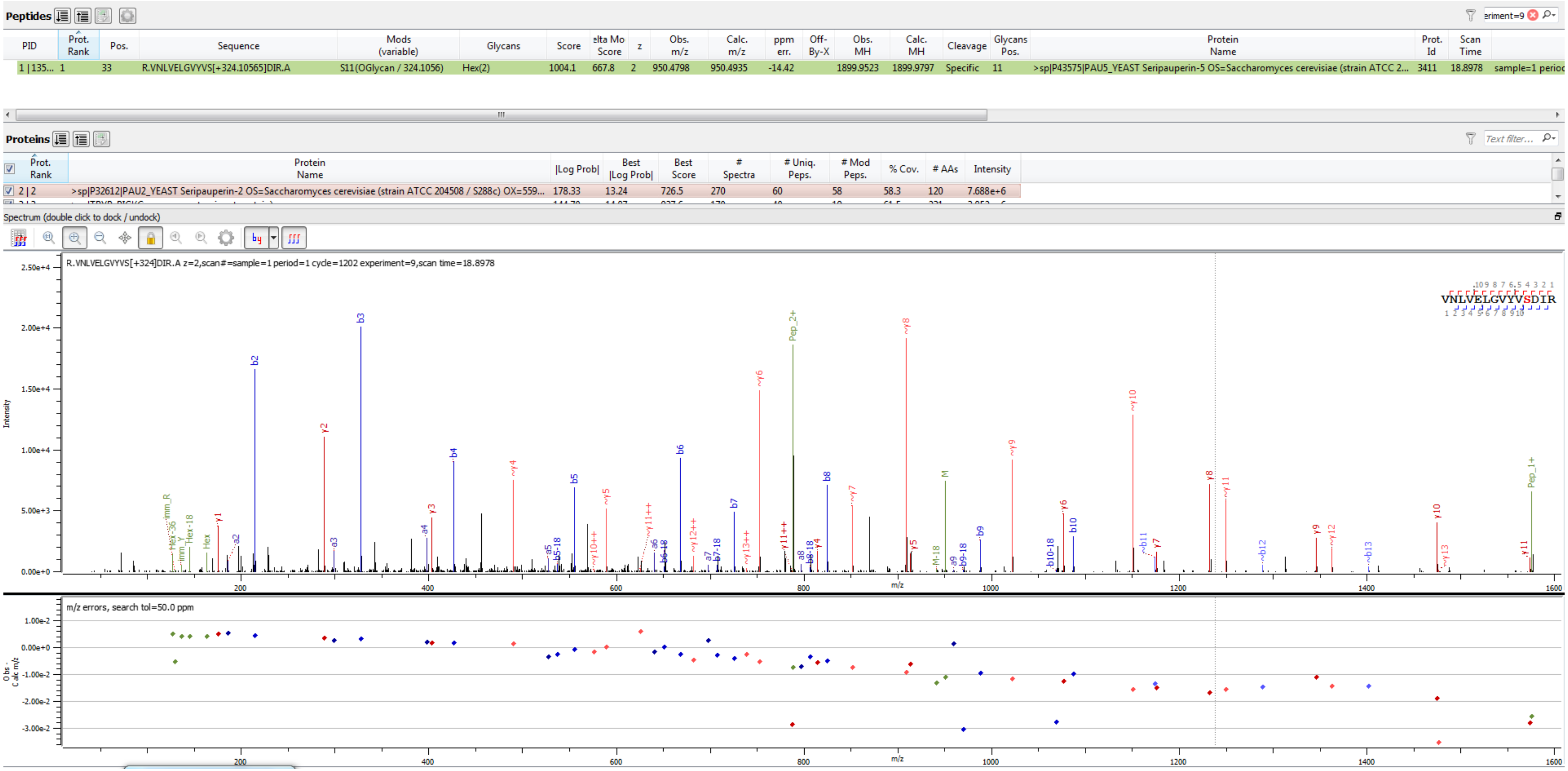


Sample S1 R.VNLVELGVYVS[+324.106]DIR.A

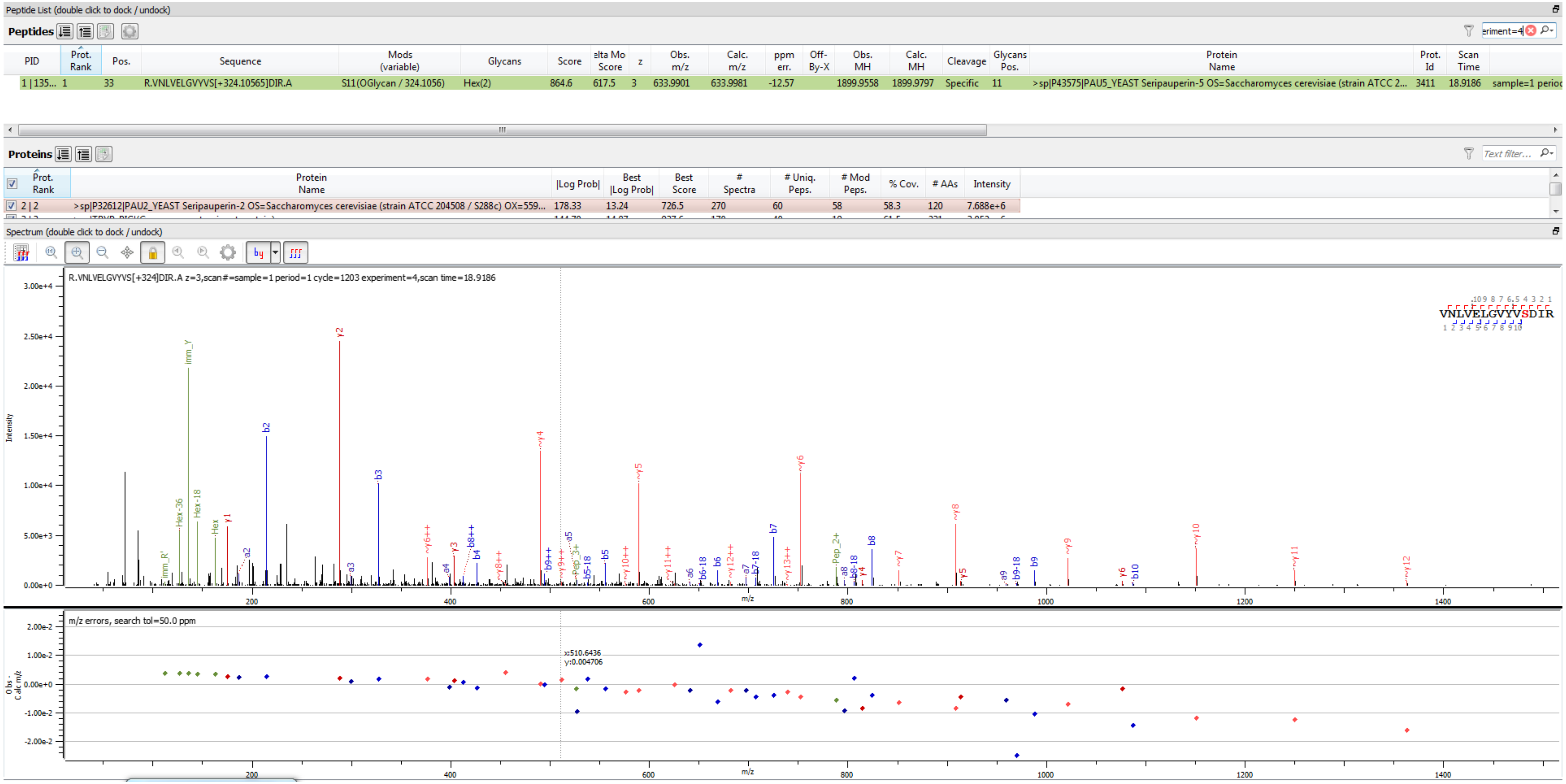

Sample S1 R.VNLVELGVYVS[+324.106]DIR.A

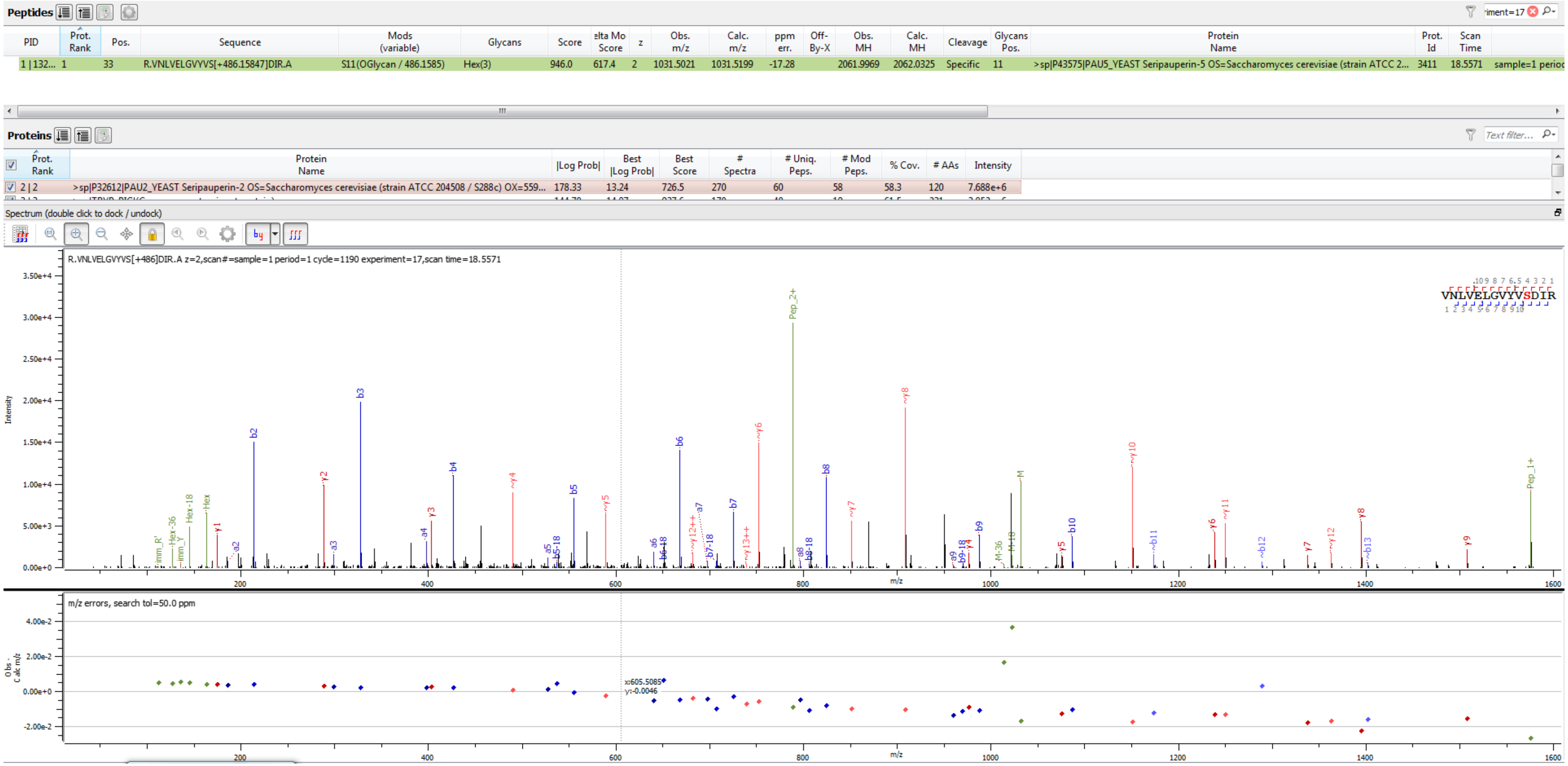


Sample S1 R.VNLVELGVYVS[+486.158]DIR.A

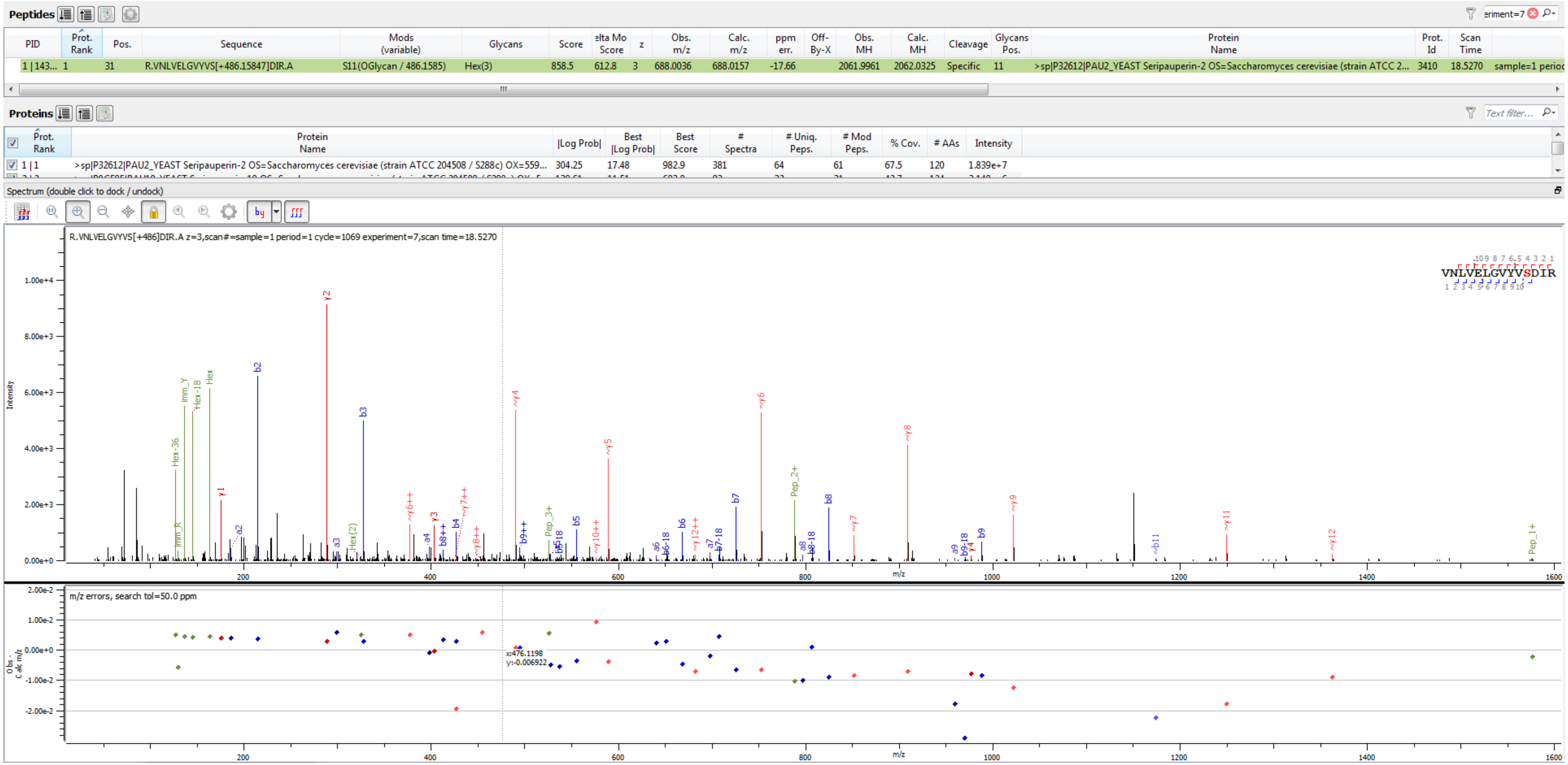


Sample S4 R.VNLVELGVYVS[+486.158]DIR.A

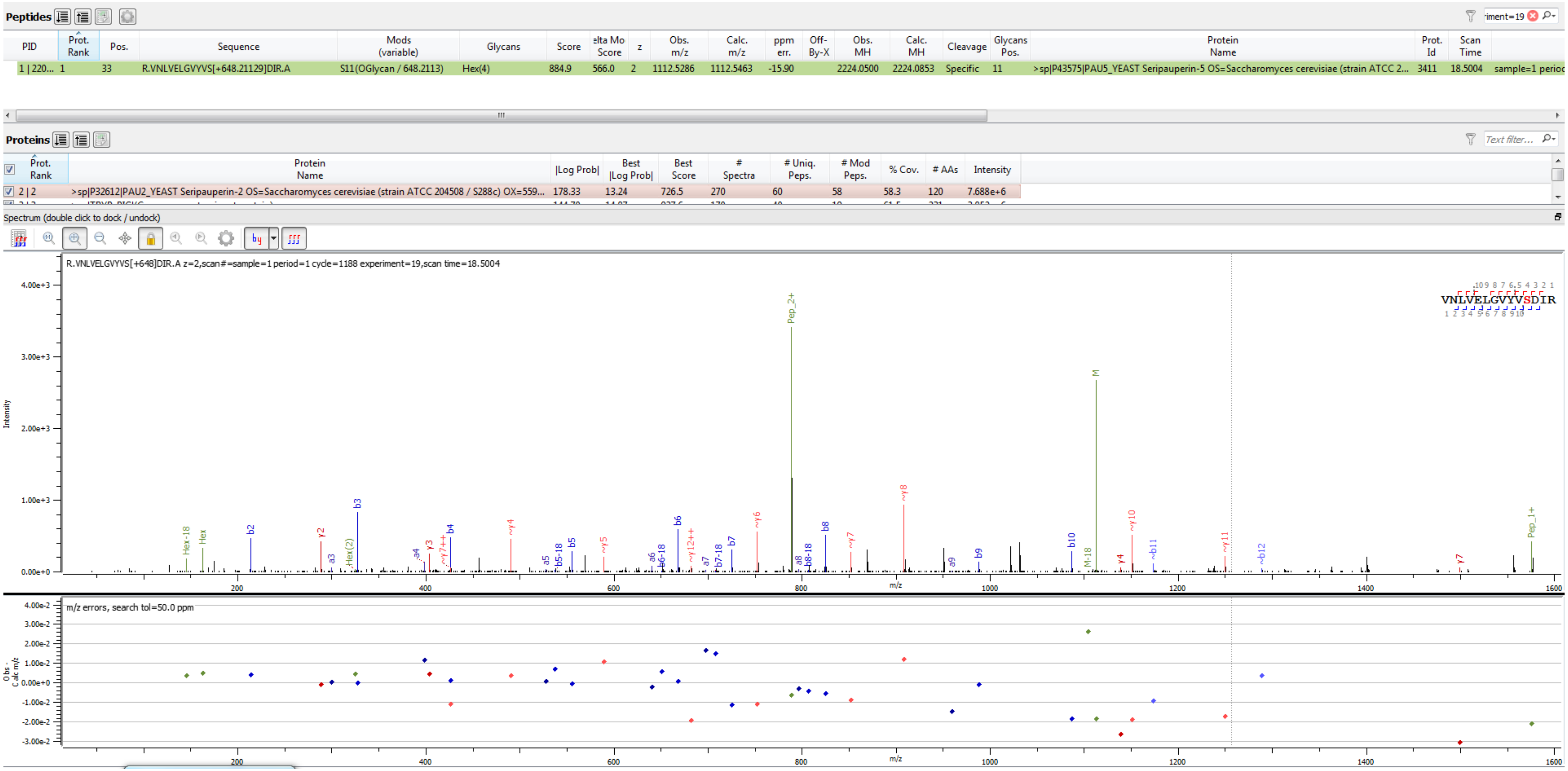


Sample S1 R.VNLVELGVYVS[+648.211]DIR.A

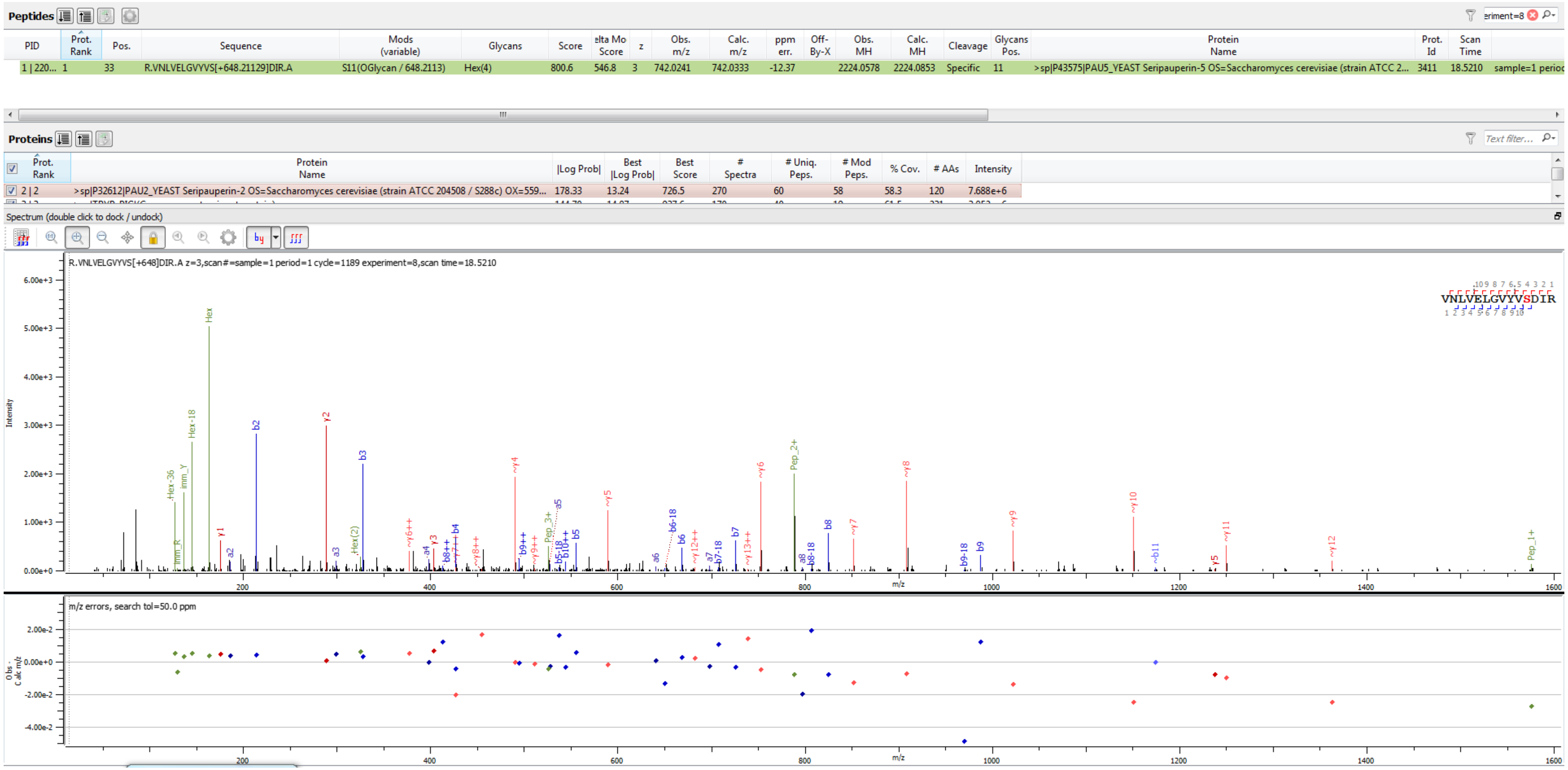

Sample S1 R.VNLVELGVYVS[+648.211]DIR.A

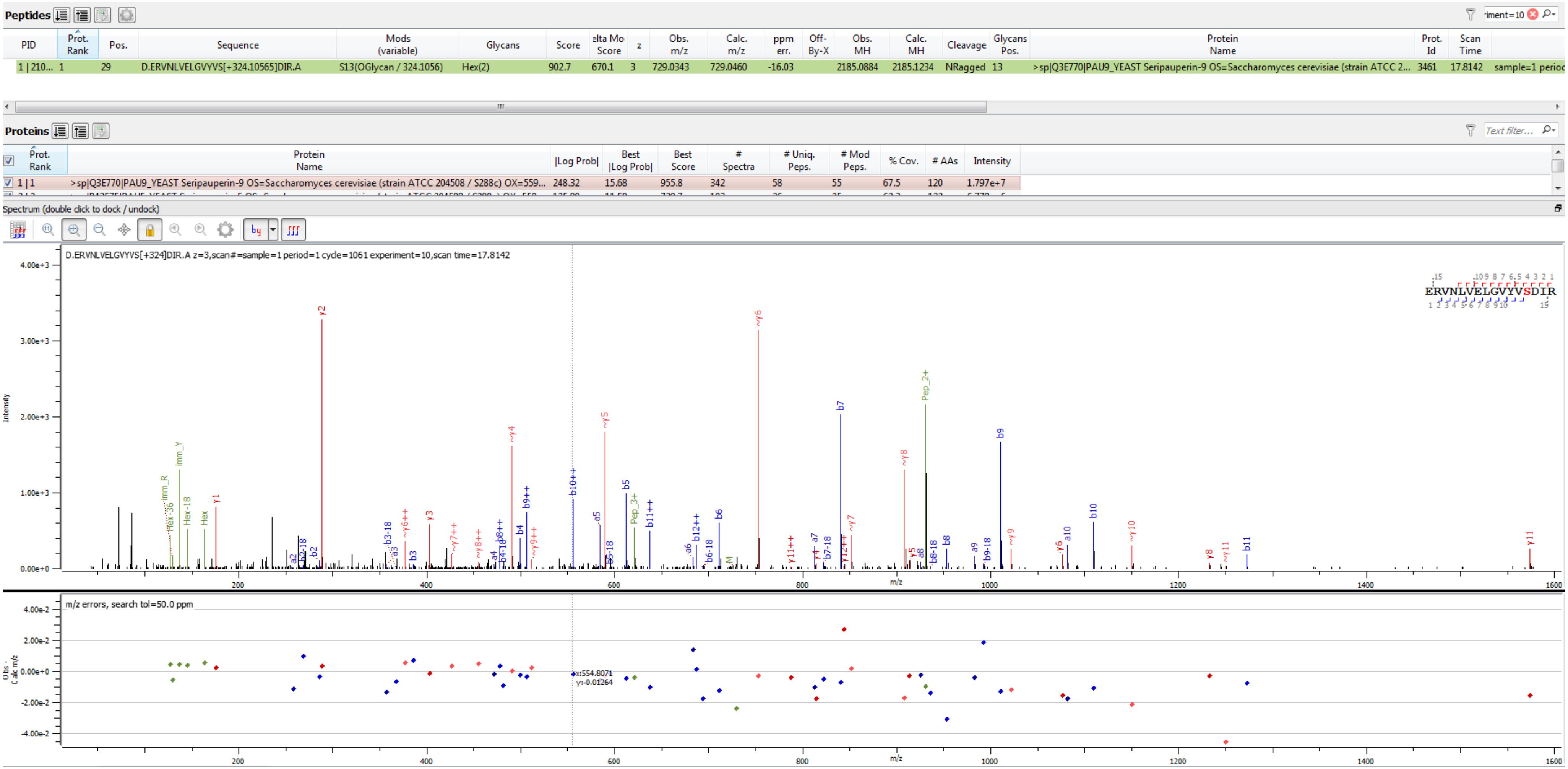


Sample S3 D.ERVNLVELGVYYVS[+324.106]DIR.A

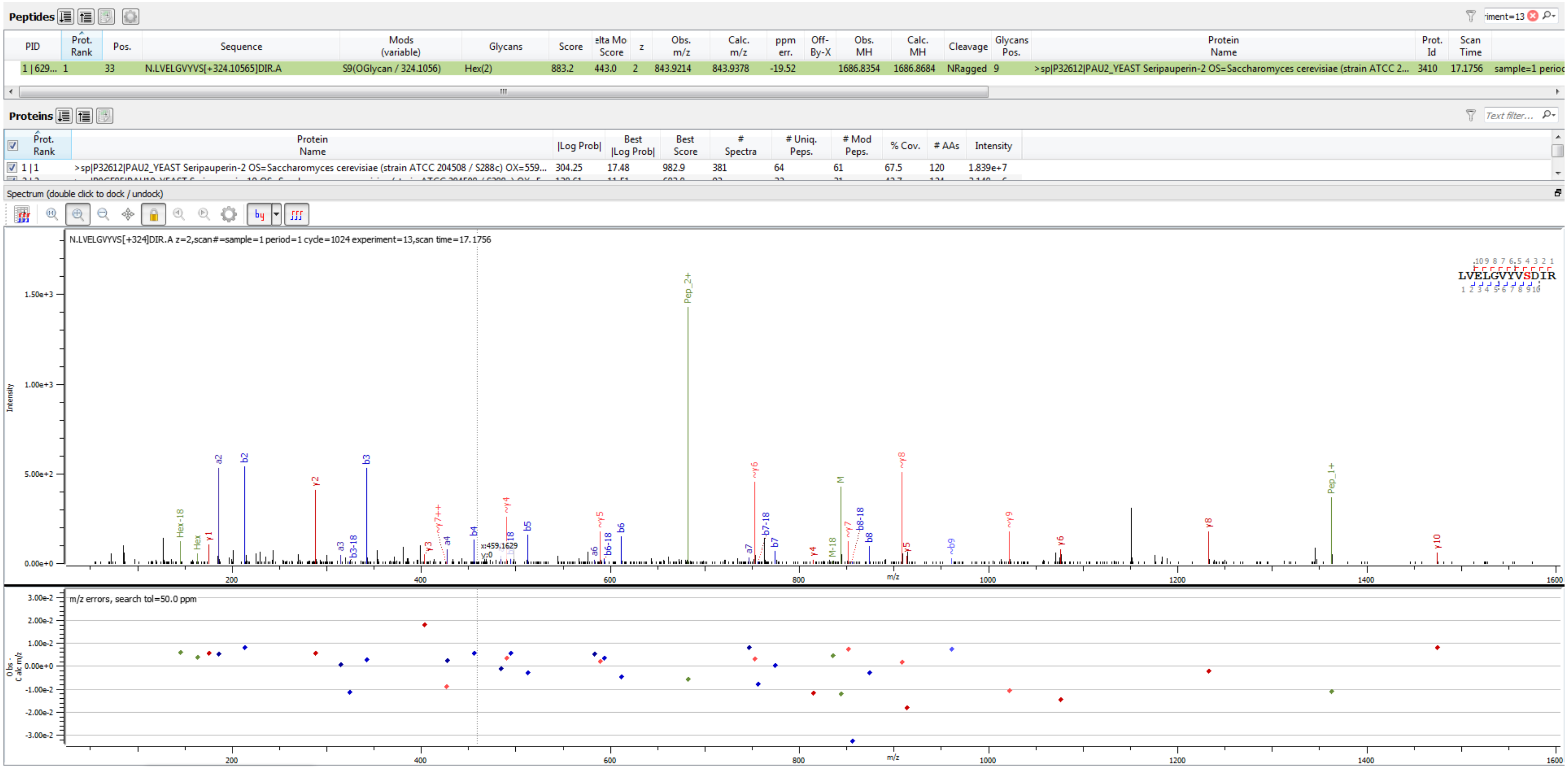


Sample S4 N.LVELGVYVS[+324.106]DIR.A

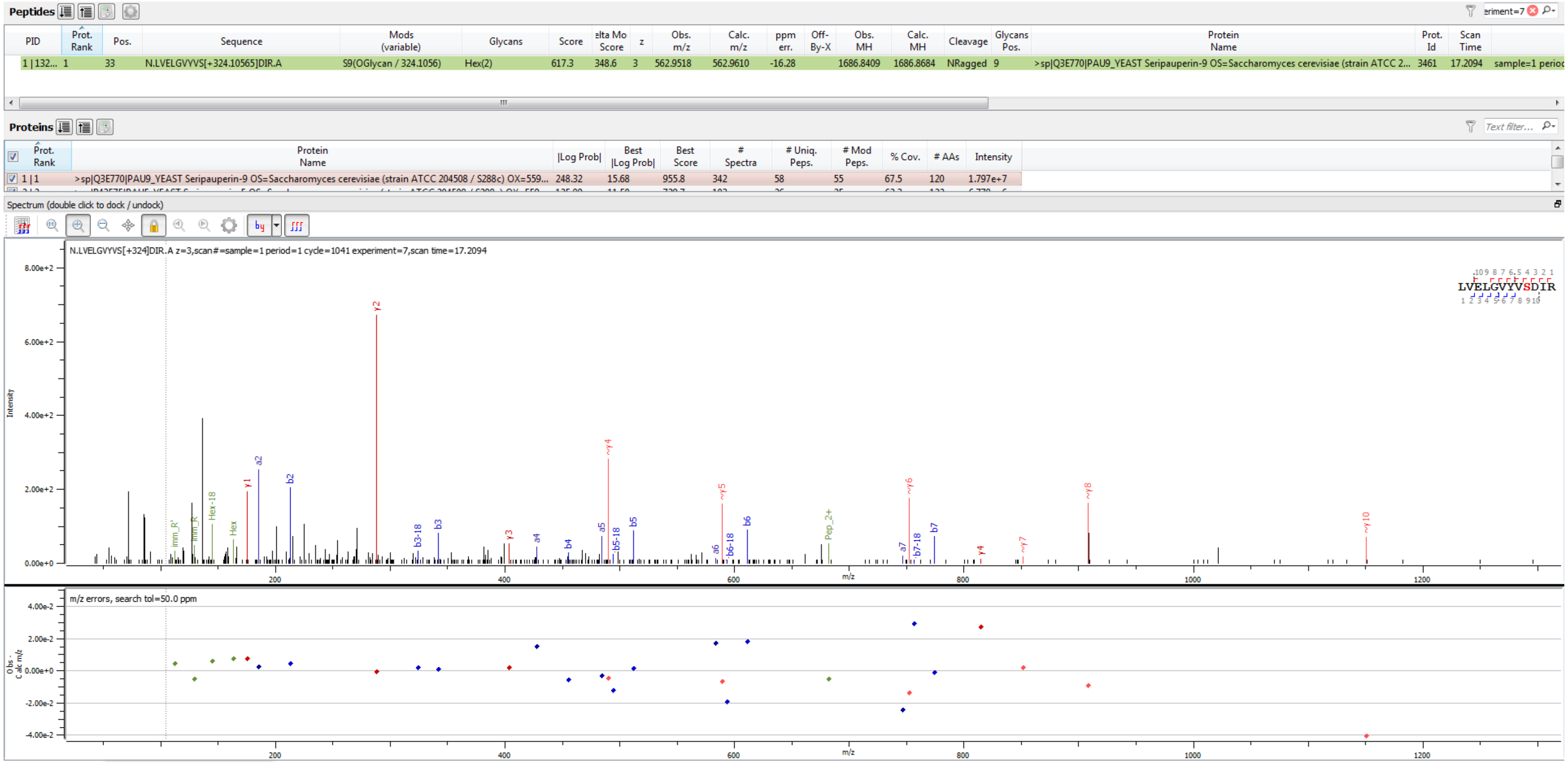


Sample S3 N.LVELGVYVS[+324.106]DIR.A

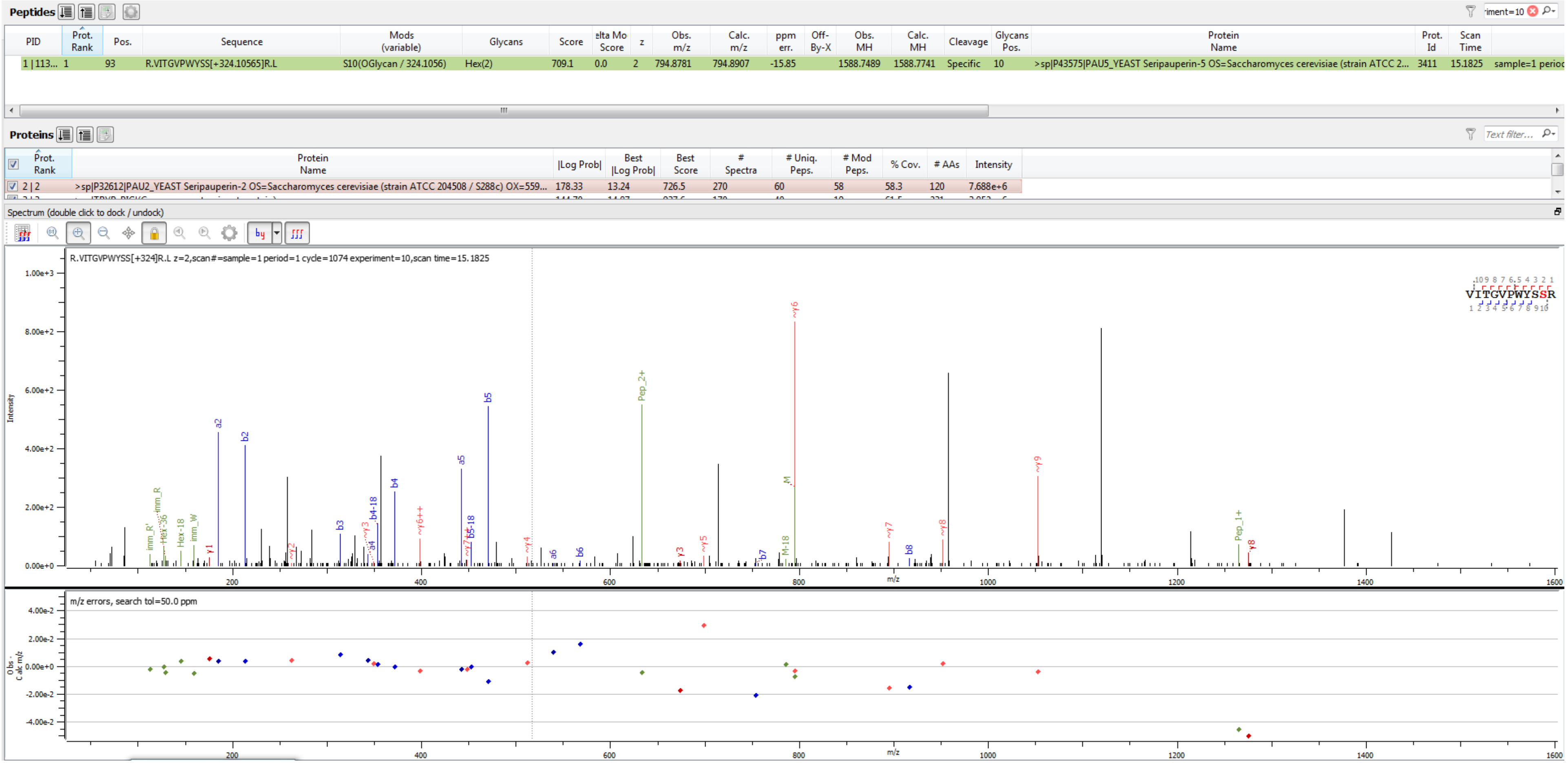


Sample S1 R.VITGVPWYS[+324.106]SR.L

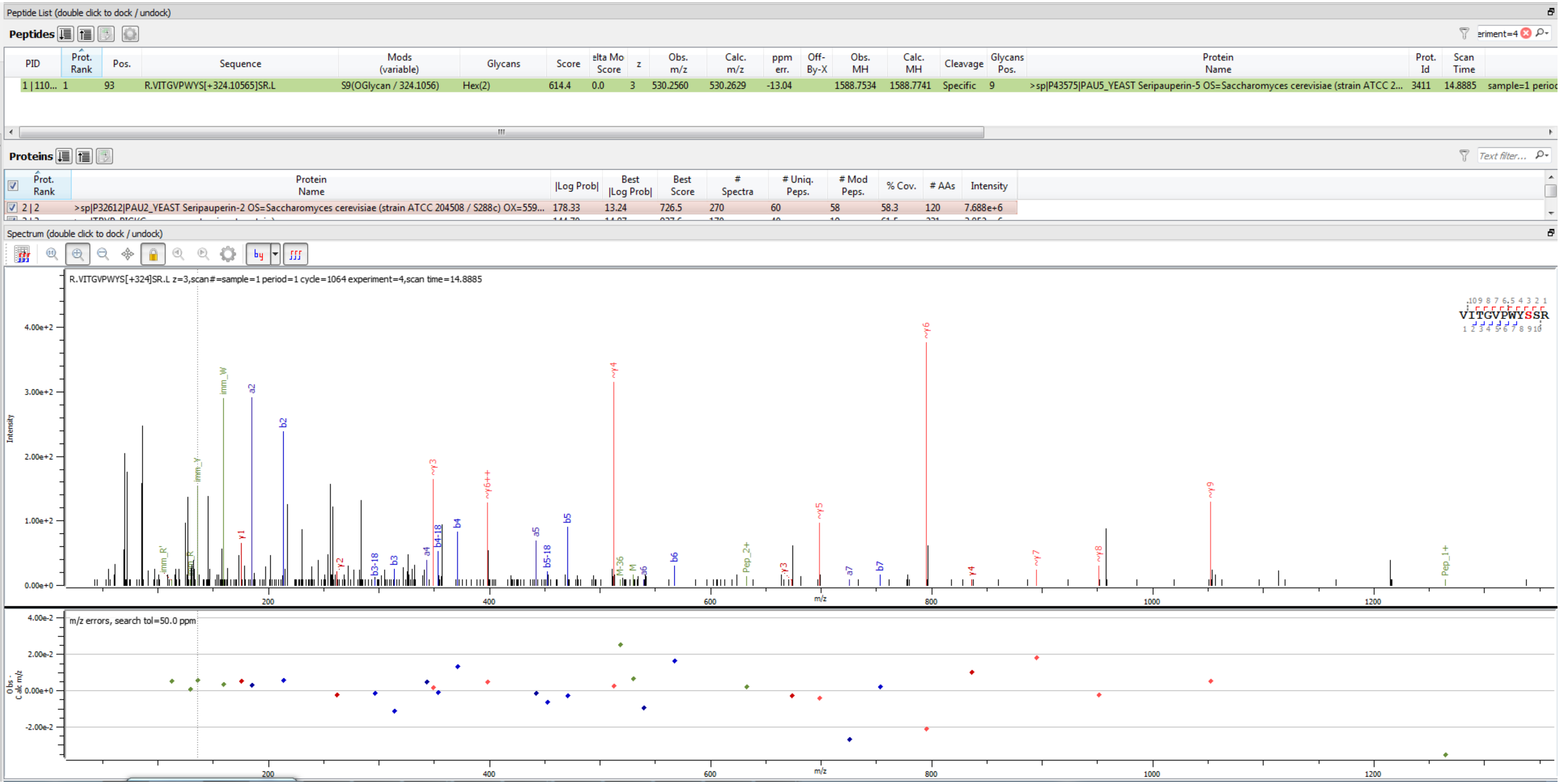


Sample S1 R.VITGVPWYS[+324.106]SR.L

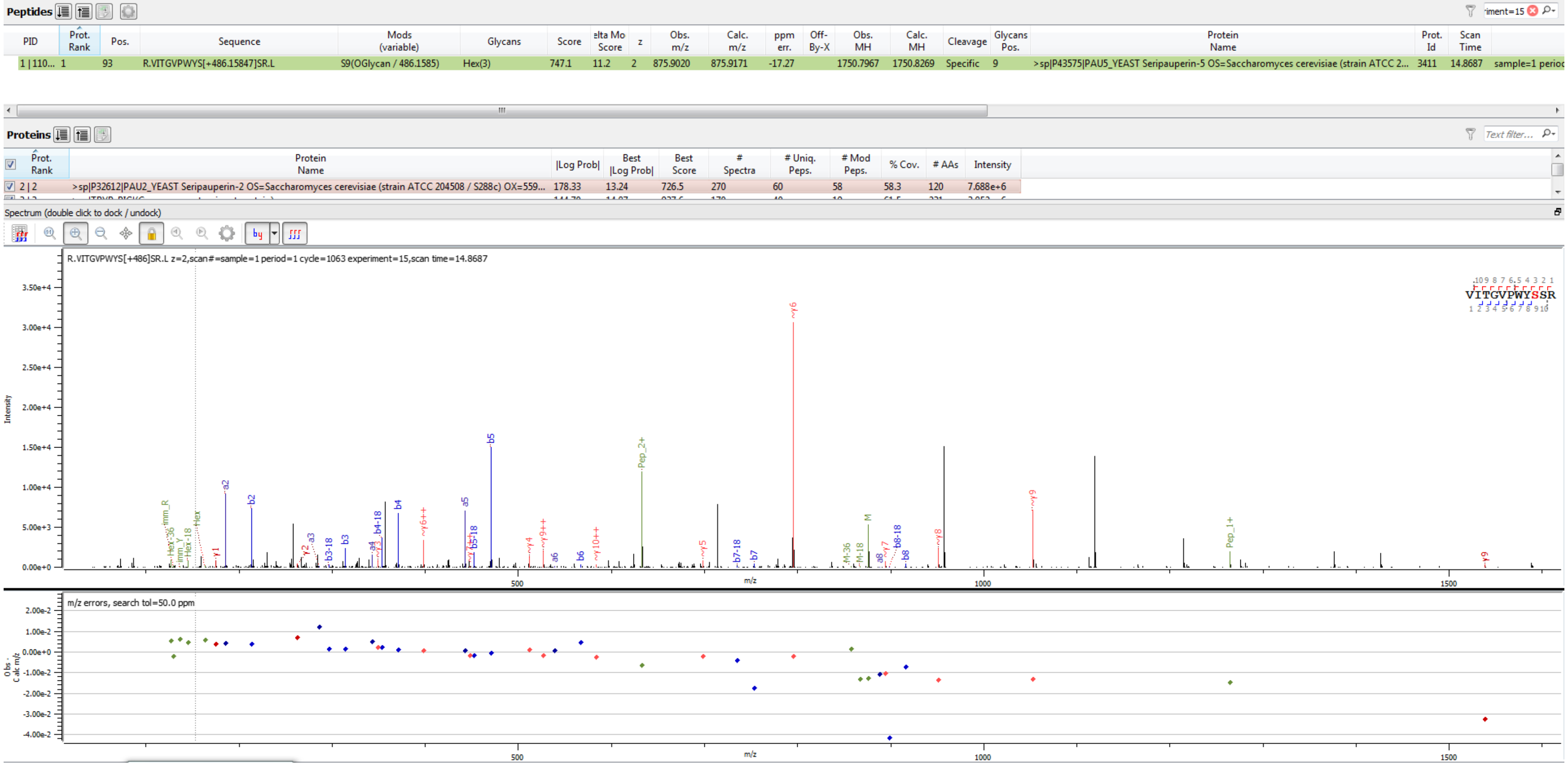


Sample S1 R.VITGVPWYS[+486.158]SR.L

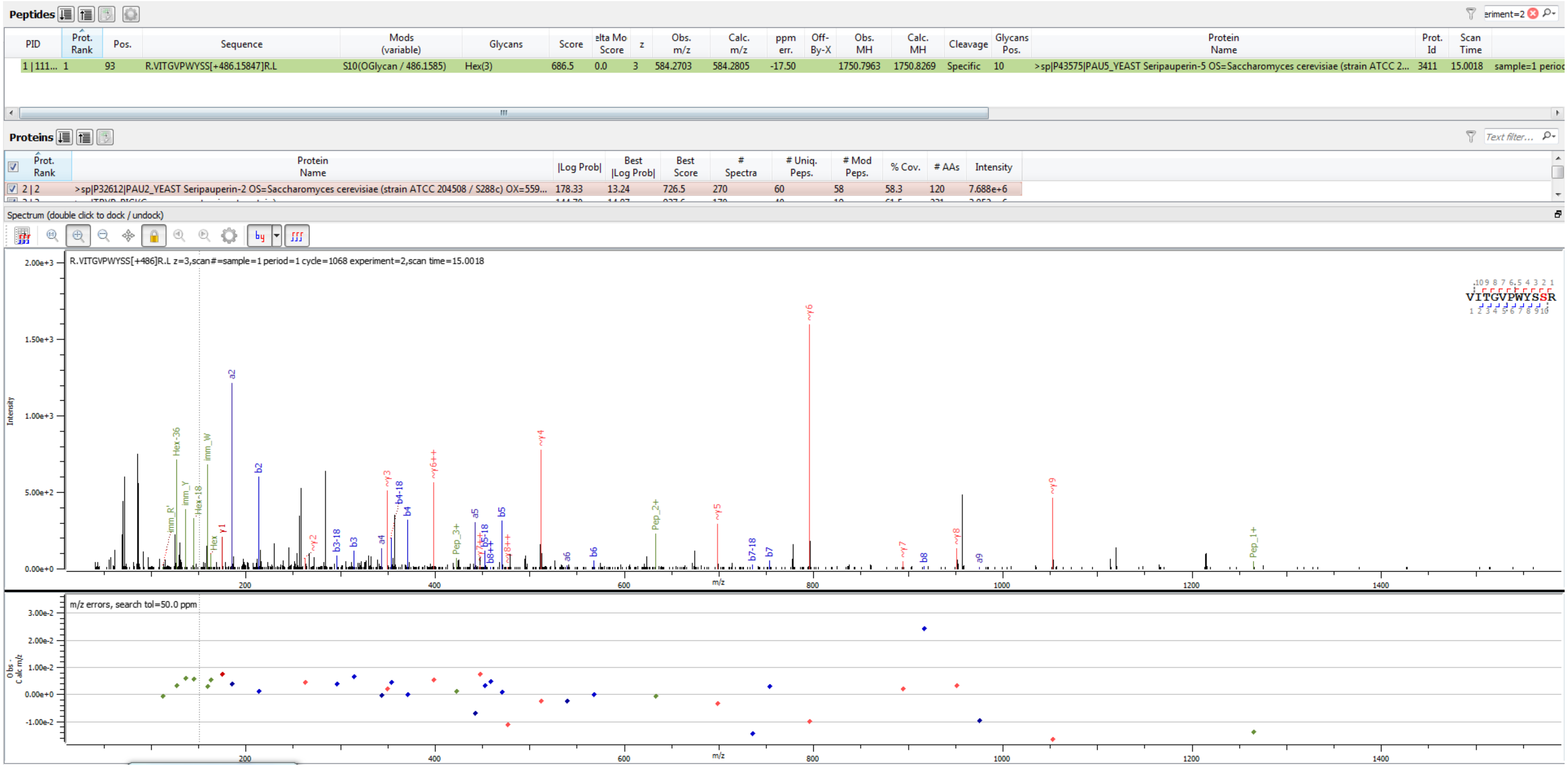


Sample S1 R.VITGVPWYS[+486.158]SR.L

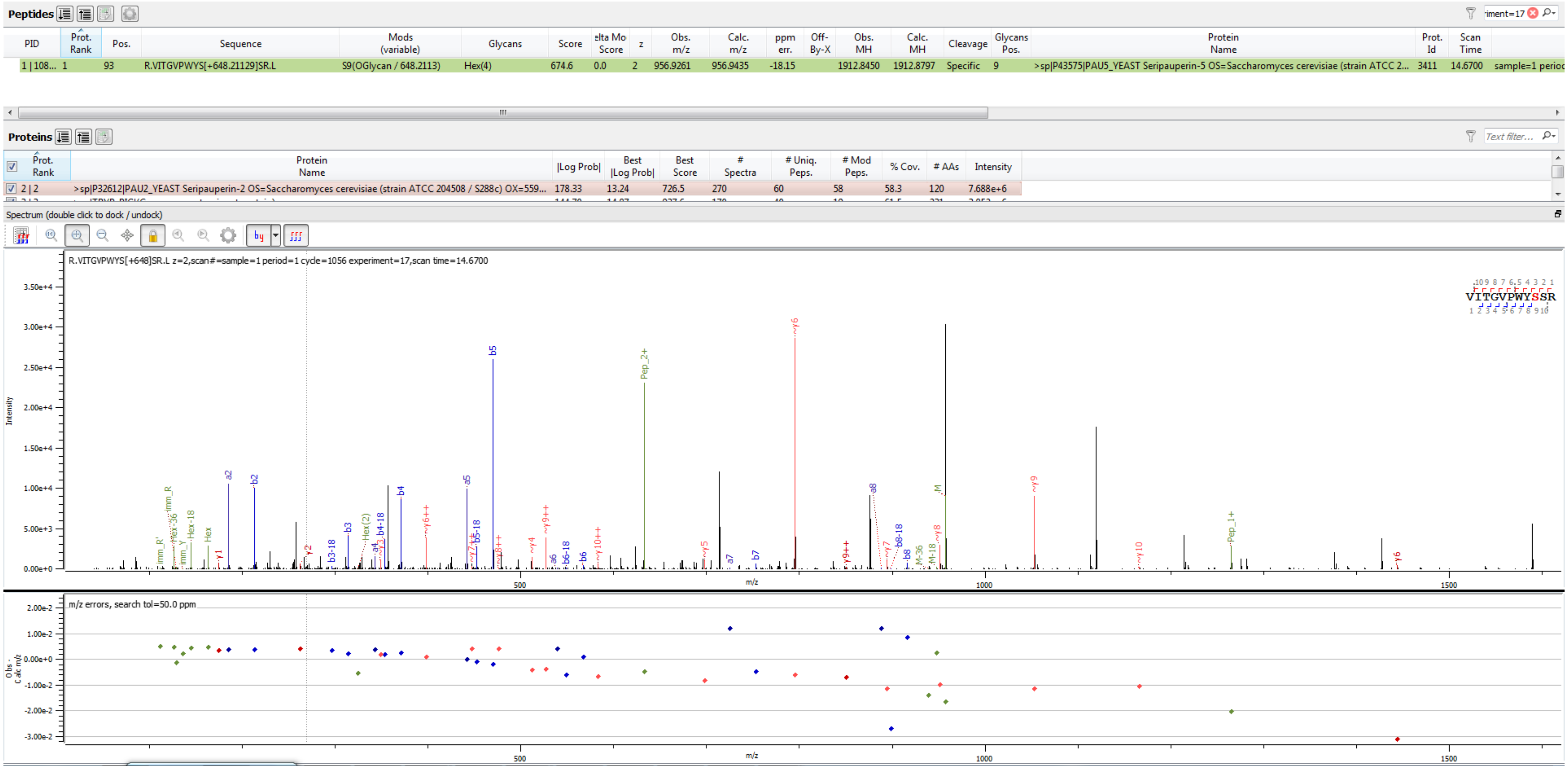

Sample S1 R.VITGVPWYS[+648.211]SR.L

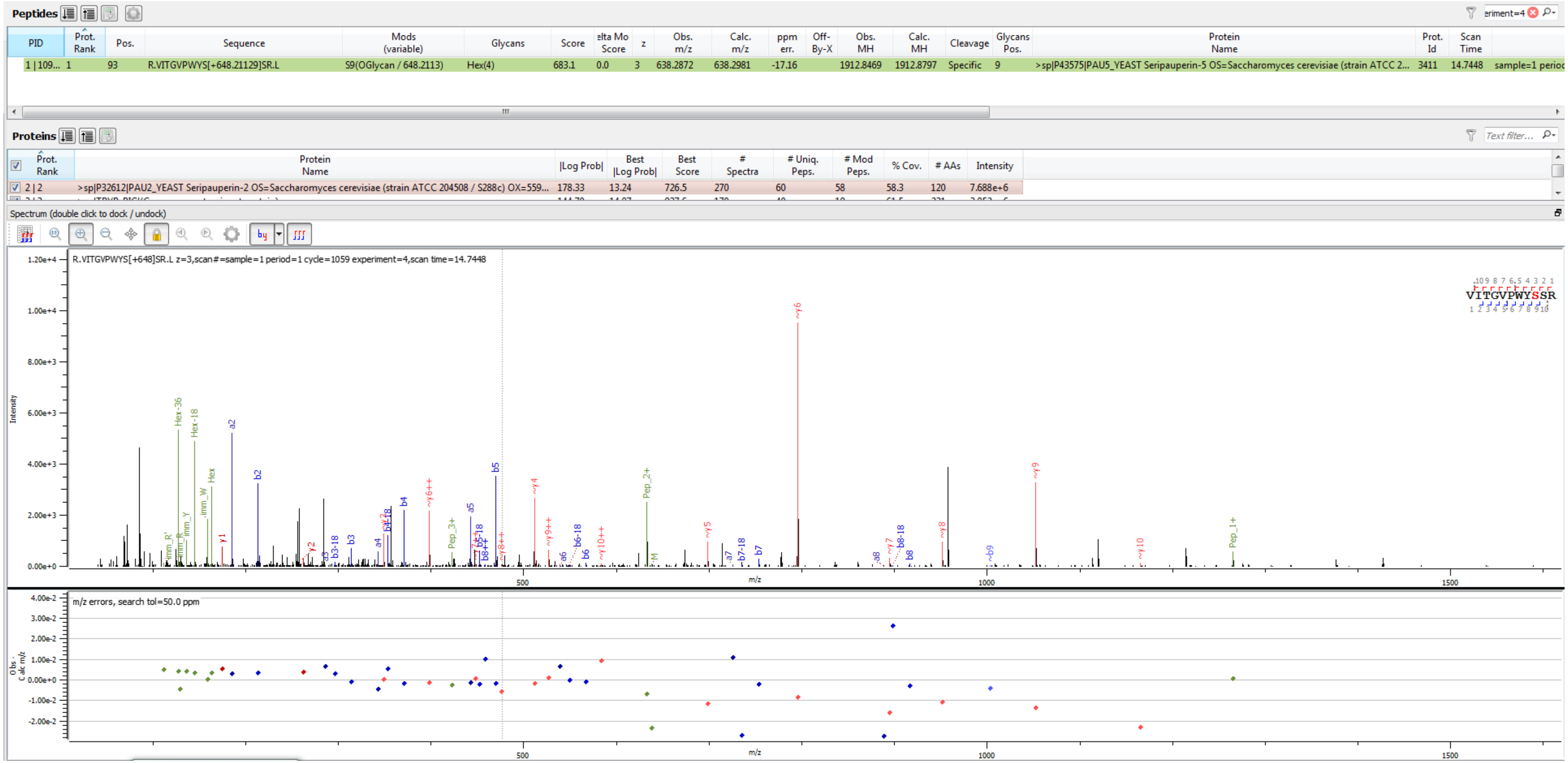


Sample S1 R.VITGVPWYS[+648.211]SR.L

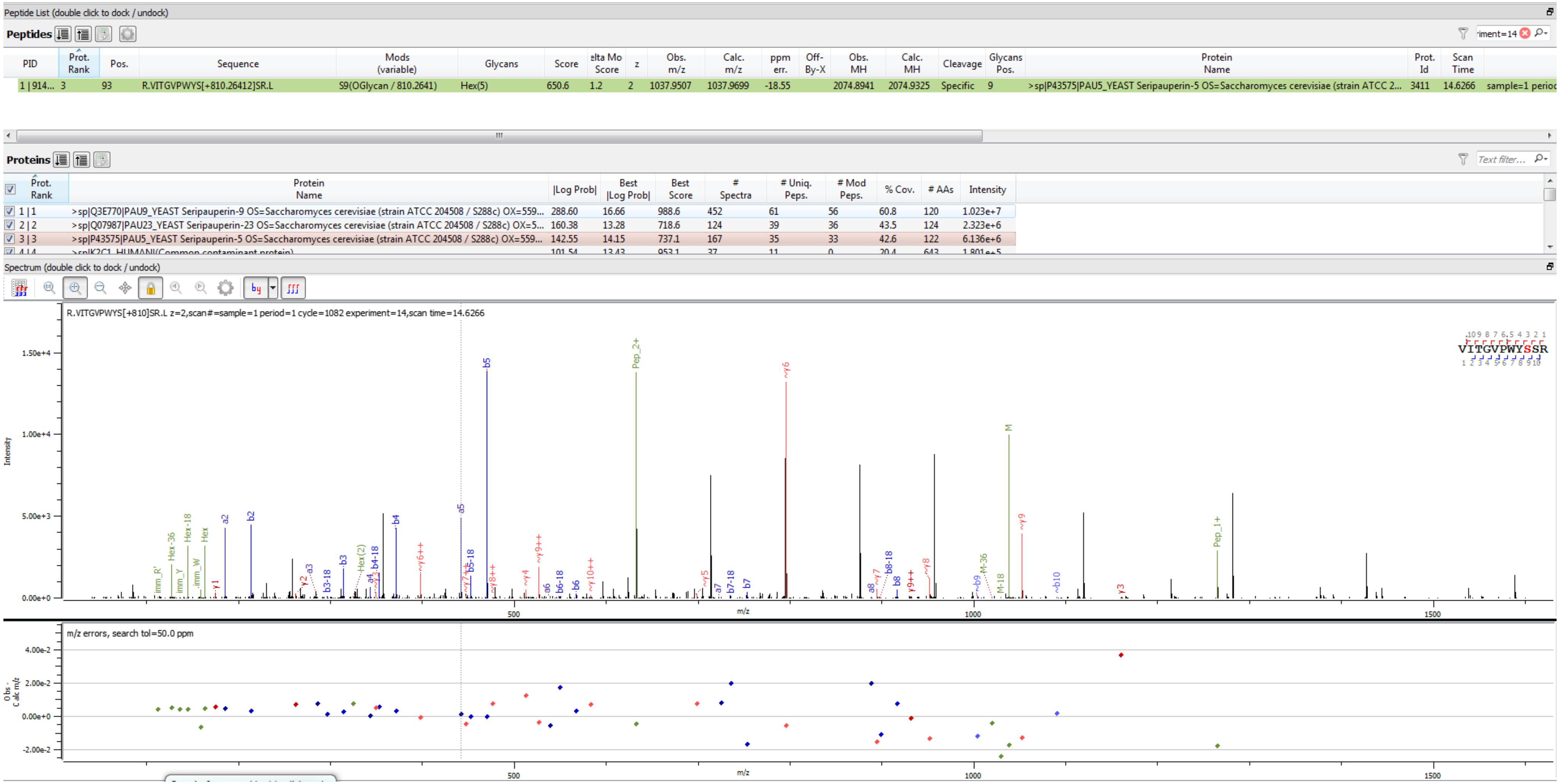


Sample S6 R.VITGVPWYS[+810.264]SR.L

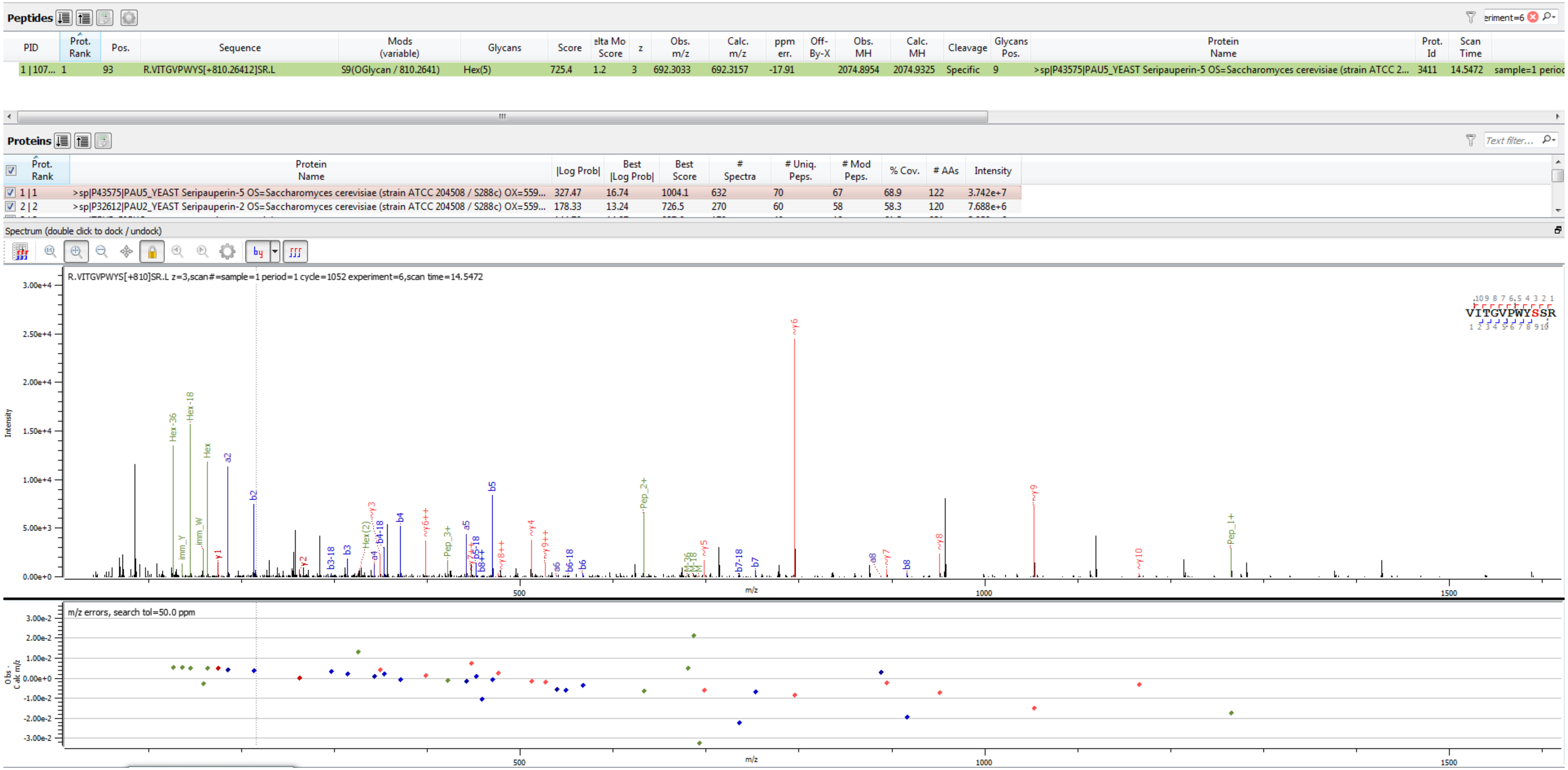


Sample S1 R.VITGVPWYS[+810.264]SR.L

Peptide List (double click to dock / undock)

**Peptides**

| PID | Prot. Rank | Pos. | Sequence | Mods (variable) | Glycans | Score | elta Mo Score | z | Obs. m/z | Calc. m/z | ppm err. | Off-By-X | Obs. MH | Calc. MH | Cleavage | Glycans Pos. | Protein Name | Prot. Id | Scan Time | |
|---|---|---|---|---|---|---|---|---|---|---|---|---|---|---|---|---|---|---|---|---|
| 1 \| 130... | 3 | 93 | R.VITGVPWYS[+972.31694]SR.L | S9(OGlycan / 972.3169) | Hex(6) | 654.1 | 1.2 | 2 | 1118.9854 | 1118.9963 | -9.79 | | 2236.9635 | 2236.9854 | Specific | 9 | >sp\|P43575\|PAU5_YEAST Seripauperin-5 OS=Saccharomyces cerevisiae (strain ATCC 2... | 3411 | 14.3331 | sample=1 perioc |

**Proteins**

| Prot. Rank | Protein Name | \|Log Prob\| | Best \|Log Prob\| | Best Score | # Spectra | # Uniq. Peps. | # Mod Peps. | % Cov. | # AAs | Intensity |
|---|---|---|---|---|---|---|---|---|---|---|
| 1 \| 1 | >sp\|TRYP_PIG\|(Common contaminant protein) | 201.90 | 15.17 | 1032.1 | 433 | 52 | 23 | 59.3 | 231 | 4.729e+6 |
| 2 \| 2 | >sp\|K2C1_HUMAN\|(Common contaminant protein) | 164.15 | 12.91 | 1037.3 | 98 | 28 | 4 | 35.8 | 643 | 4.850e+5 |
| 3 \| 3 | >sp\|P43575\|PAU5_YEAST Seripauperin-5 OS=Saccharomyces cerevisiae (strain ATCC 204508 / S288c) OX=559... | 117.08 | 14.43 | 972.1 | 214 | 40 | 39 | 34.4 | 122 | 3.360e+6 |
| 4 \| 4 | >sp\|K1C9_HUMAN\|(Common contaminant protein) | 66.04 | 18.99 | 1150.7 | 49 | 14 | 3 | 24.9 | 623 | 1.617e+5 |

Spectrum (double click to dock / undock)

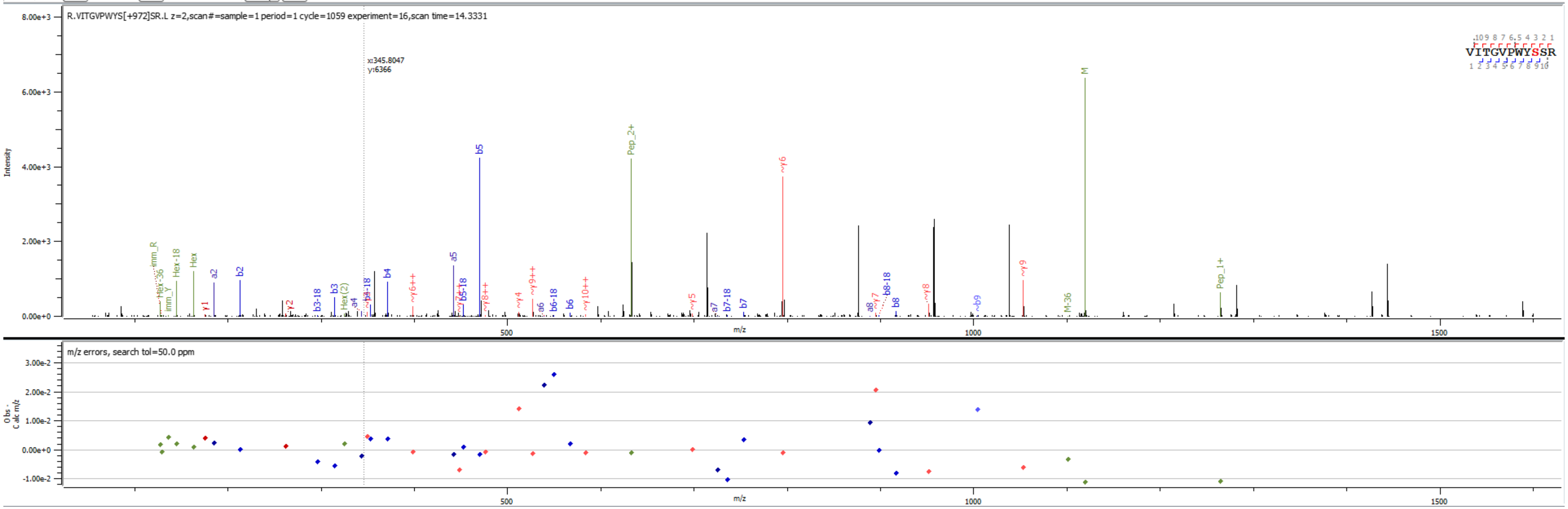


Sample S2 R.VITGVPWYS[+972.317]SR.L

**Peptides**

| PID | Prot. Rank | Pos. | Sequence | Mods (variable) | Glycans | Score | elta Mo Score | z | Obs. m/z | Calc. m/z | ppm err. | Off-By-X | Obs. MH | Calc. MH | Cleavage | Glycans Pos. | Protein Name | Prot. Id | Scan Time | |
|---|---|---|---|---|---|---|---|---|---|---|---|---|---|---|---|---|---|---|---|---|
| 1 \| 180... | 1 | 93 | R.VITGVPWYS[+972.31694]SR.L | S9(OGlycan / 972.3169) | Hex(6) | 712.2 | 1.2 | 3 | 746.3205 | 746.3333 | -17.15 | | 2236.9470 | 2236.9854 | Specific | 9 | >sp\|P43575\|PAU5_YEAST Seripauperin-5 OS=Saccharomyces cerevisiae (strain ATCC 2... | 3411 | 14.2101 | sample=1 perioc |

**Proteins**

| Prot. Rank | Protein Name | \|Log Prob\| | Best \|Log Prob\| | Best Score | # Spectra | # Uniq. Peps. | # Mod Peps. | % Cov. | # AAs | Intensity |
|---|---|---|---|---|---|---|---|---|---|---|
| 1 \| 1 | >sp\|P43575\|PAU5_YEAST Seripauperin-5 OS=Saccharomyces cerevisiae (strain ATCC 204508 / S288c) OX=559... | 327.47 | 16.74 | 1004.1 | 632 | 70 | 67 | 68.9 | 122 | 3.742e+7 |
| 2 \| 2 | >sp\|P32612\|PAU2_YEAST Seripauperin-2 OS=Saccharomyces cerevisiae (strain ATCC 204508 / S288c) OX=559... | 178.33 | 13.24 | 726.5 | 270 | 60 | 58 | 58.3 | 120 | 7.688e+6 |

Spectrum (double click to dock / undock)

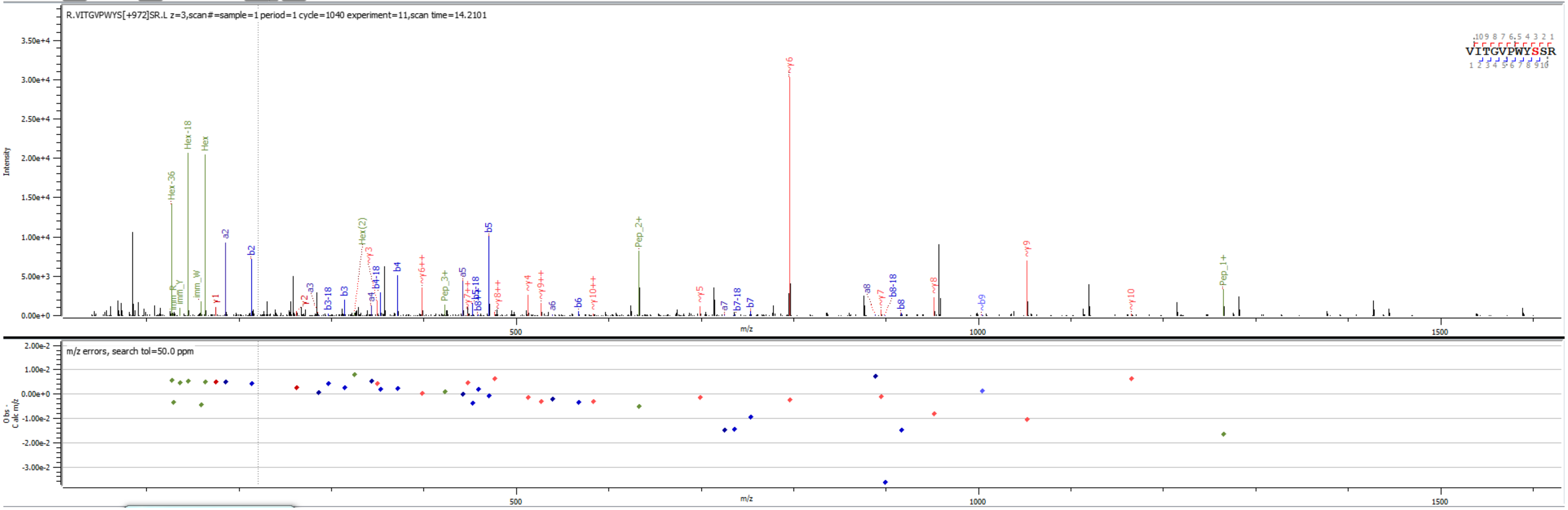


Sample S1 R.VITGVPWYS[+972.317]SR.L

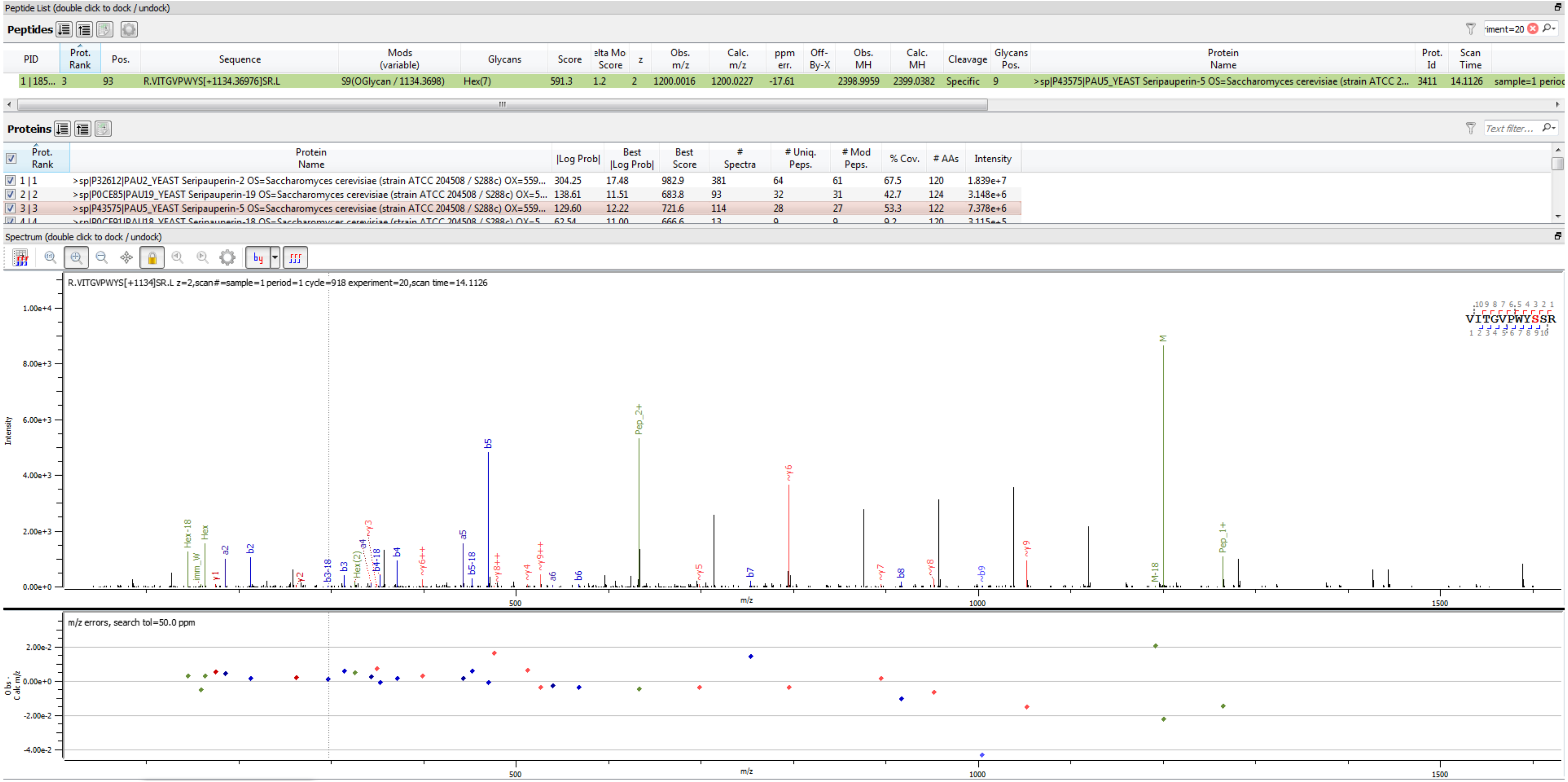


Sample S4 R.VITGVPWYS[+1134.370]SR.L

**Peptides**

| PID | Prot. Rank | Pos. | Sequence | Mods (variable) | Glycans | Score | elta Mo Score | z | Obs. m/z | Calc. m/z | ppm err. | Off-By-X | Obs. MH | Calc. MH | Cleavage | Glycans Pos. | Protein Name | Prot. Id | Scan Time | |
|---|---|---|---|---|---|---|---|---|---|---|---|---|---|---|---|---|---|---|---|---|
| 1 \| 180... | 1 | 93 | R.VITGVPWYS[+1134.36976]SR.L | S9(OGlycan / 1134.3698) | Hex(7) | 673.9 | 1.2 | 3 | 800.3351 | 800.3509 | -19.80 | | 2398.9907 | 2399.0382 | Specific | 9 | >sp\|P43575\|PAU5_YEAST Seripauperin-5 OS=Saccharomyces cerevisiae (strain ATCC 2... | 3411 | 14.2126 | sample=1 perioc |

**Proteins**

| Prot. Rank | Protein Name | \|Log Prob\| | Best \|Log Prob\| | Best Score | # Spectra | # Uniq. Peps. | # Mod Peps. | % Cov. | # AAs | Intensity |
|---|---|---|---|---|---|---|---|---|---|---|
| 1 \| 1 | >sp\|P43575\|PAU5_YEAST Seripauperin-5 OS=Saccharomyces cerevisiae (strain ATCC 204508 / S288c) OX=559... | 327.47 | 16.74 | 1004.1 | 632 | 70 | 67 | 68.9 | 122 | 3.742e+7 |
| 2 \| 2 | >sp\|P32612\|PAU2_YEAST Seripauperin-2 OS=Saccharomyces cerevisiae (strain ATCC 204508 / S288c) OX=559... | 178.33 | 13.24 | 726.5 | 270 | 60 | 58 | 58.3 | 120 | 7.688e+6 |

Spectrum (double click to dock / undock)

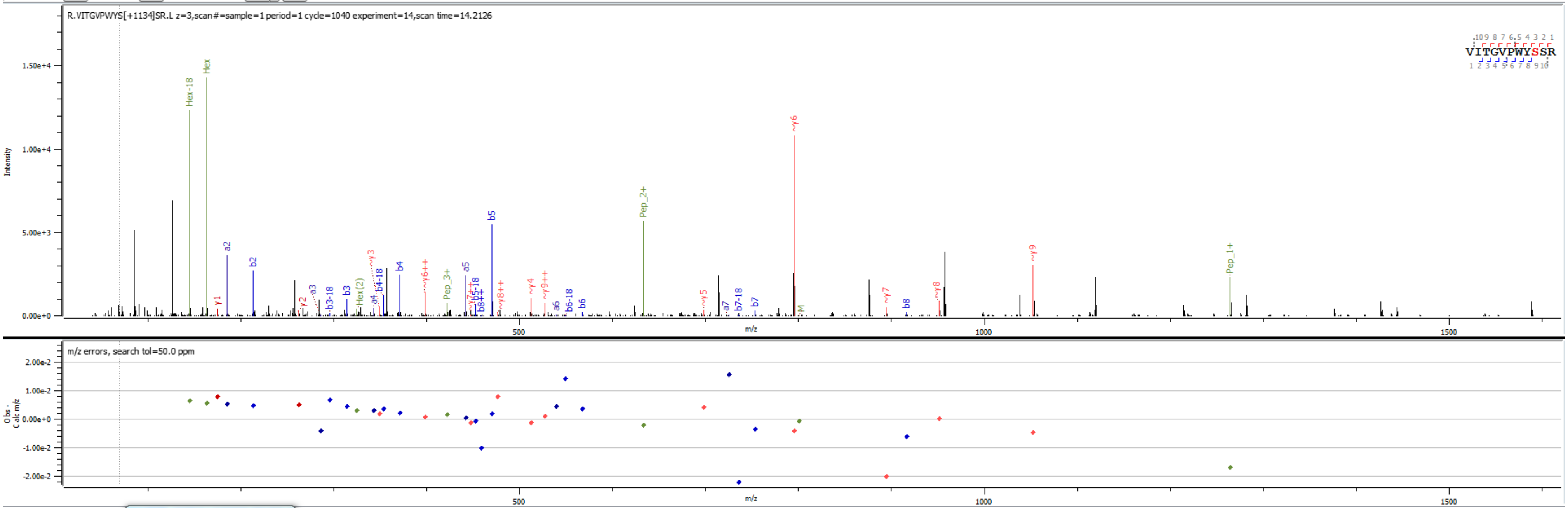


Sample S1 R.VITGVPWYS[+1134.370]SR.L

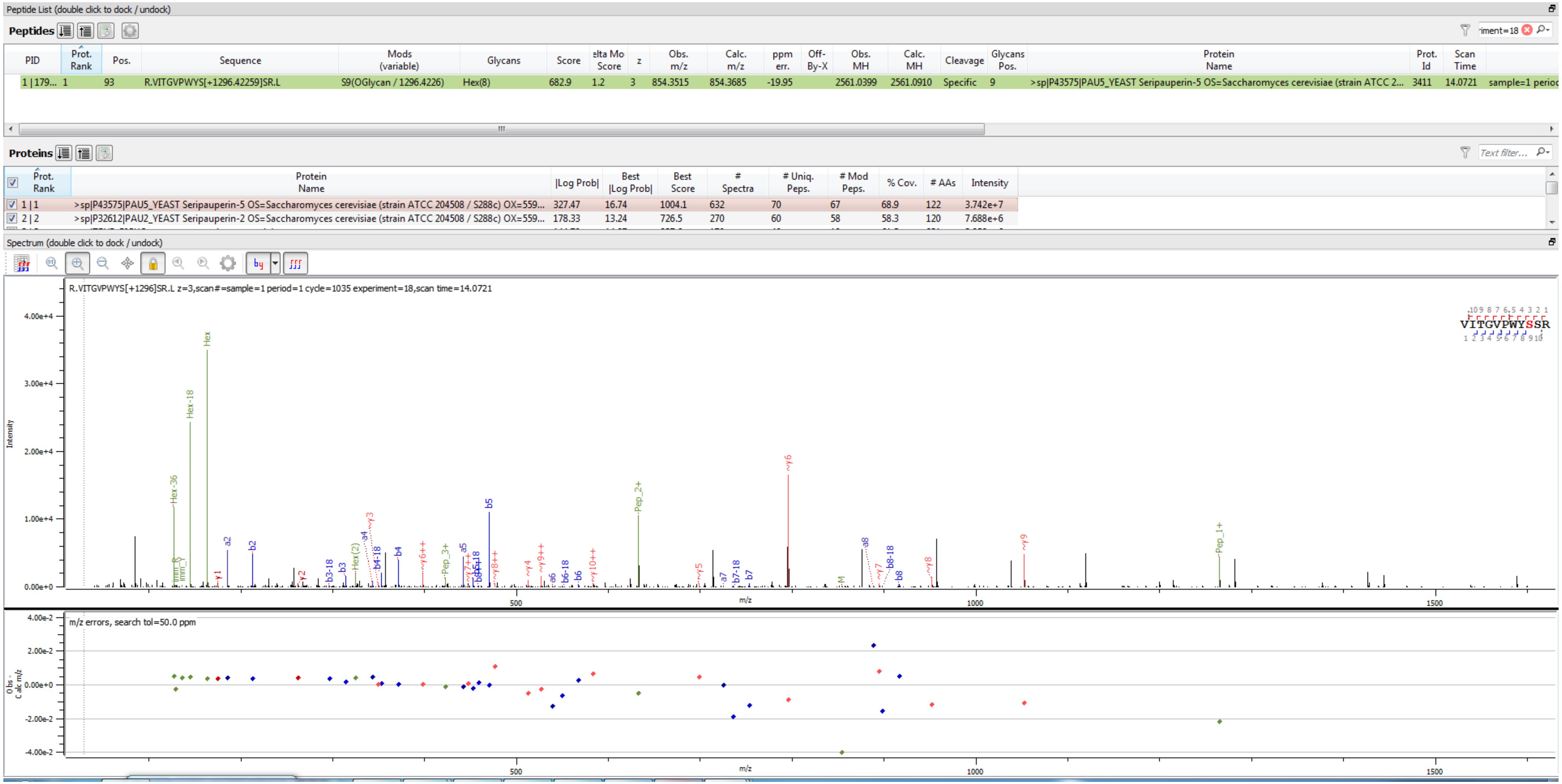


Sample S1 R.VITGVPWYS[+1296.423]SR.L

**Peptides**

| PID | Prot. Rank | Pos. | Sequence | Mods (variable) | Glycans | Score | elta Mo Score | z | Obs. m/z | Calc. m/z | ppm err. | Off-By-X | Obs. MH | Calc. MH | Cleavage | Glycans Pos. | Protein Name | Prot. Id | Scan Time | |
|---|---|---|---|---|---|---|---|---|---|---|---|---|---|---|---|---|---|---|---|---|
| 1 \| 178... | 1 | 93 | R.VITGVPWYS[+1458.47541]SR.L | S9(OGlycan / 1458.4754) | Hex(9) | 613.5 | 1.2 | 3 | 908.3717 | 908.3861 | -15.89 | | 2723.1006 | 2723.1438 | Specific | 9 | >sp\|P43575\|PAU5_YEAST Seripauperin-5 OS=Saccharomyces cerevisiae (strain ATCC 2... | 3411 | 13.9516 | sample=1 perioc |

**Proteins**

| Prot. Rank | Protein Name | \|Log Prob\| | Best \|Log Prob\| | Best Score | # Spectra | # Uniq. Peps. | # Mod Peps. | % Cov. | # AAs | Intensity |
|---|---|---|---|---|---|---|---|---|---|---|
| 1 \| 1 | >sp\|P43575\|PAU5_YEAST Seripauperin-5 OS=Saccharomyces cerevisiae (strain ATCC 204508 / S288c) OX=559... | 327.47 | 16.74 | 1004.1 | 632 | 70 | 67 | 68.9 | 122 | 3.742e+7 |
| 2 \| 2 | >sp\|P32612\|PAU2_YEAST Seripauperin-2 OS=Saccharomyces cerevisiae (strain ATCC 204508 / S288c) OX=559... | 178.33 | 13.24 | 726.5 | 270 | 60 | 58 | 58.3 | 120 | 7.688e+6 |

Spectrum (double click to dock / undock)

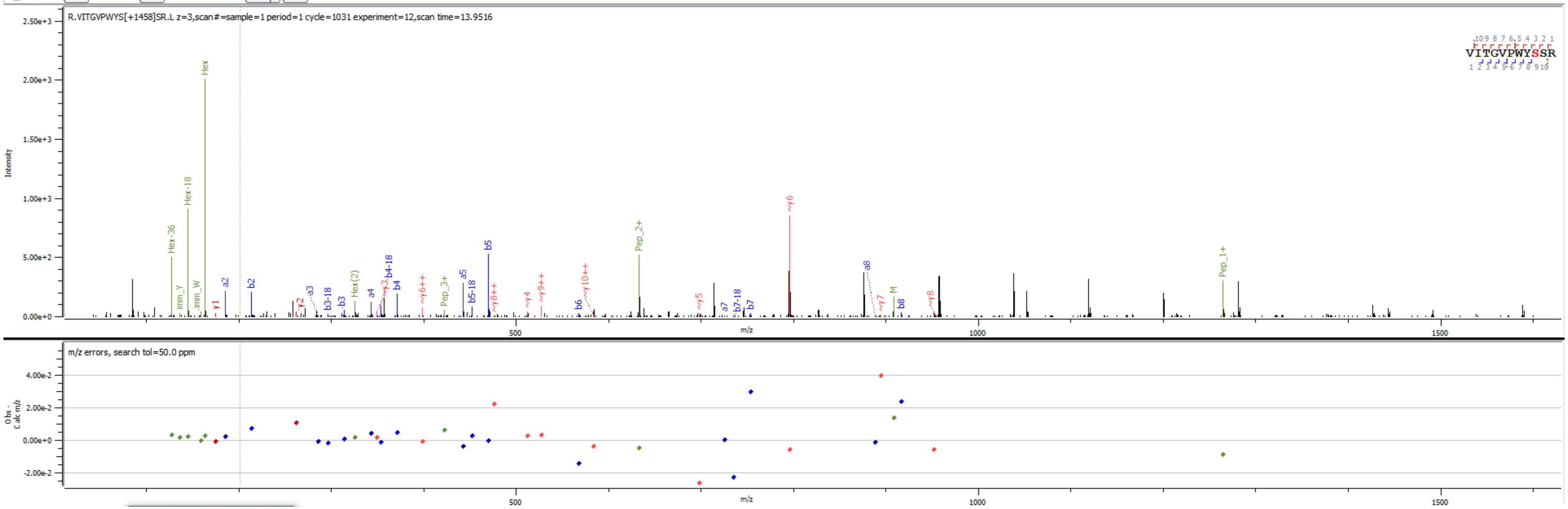


Sample S1 R.VITGVPWYS[+1458.475]SR.L

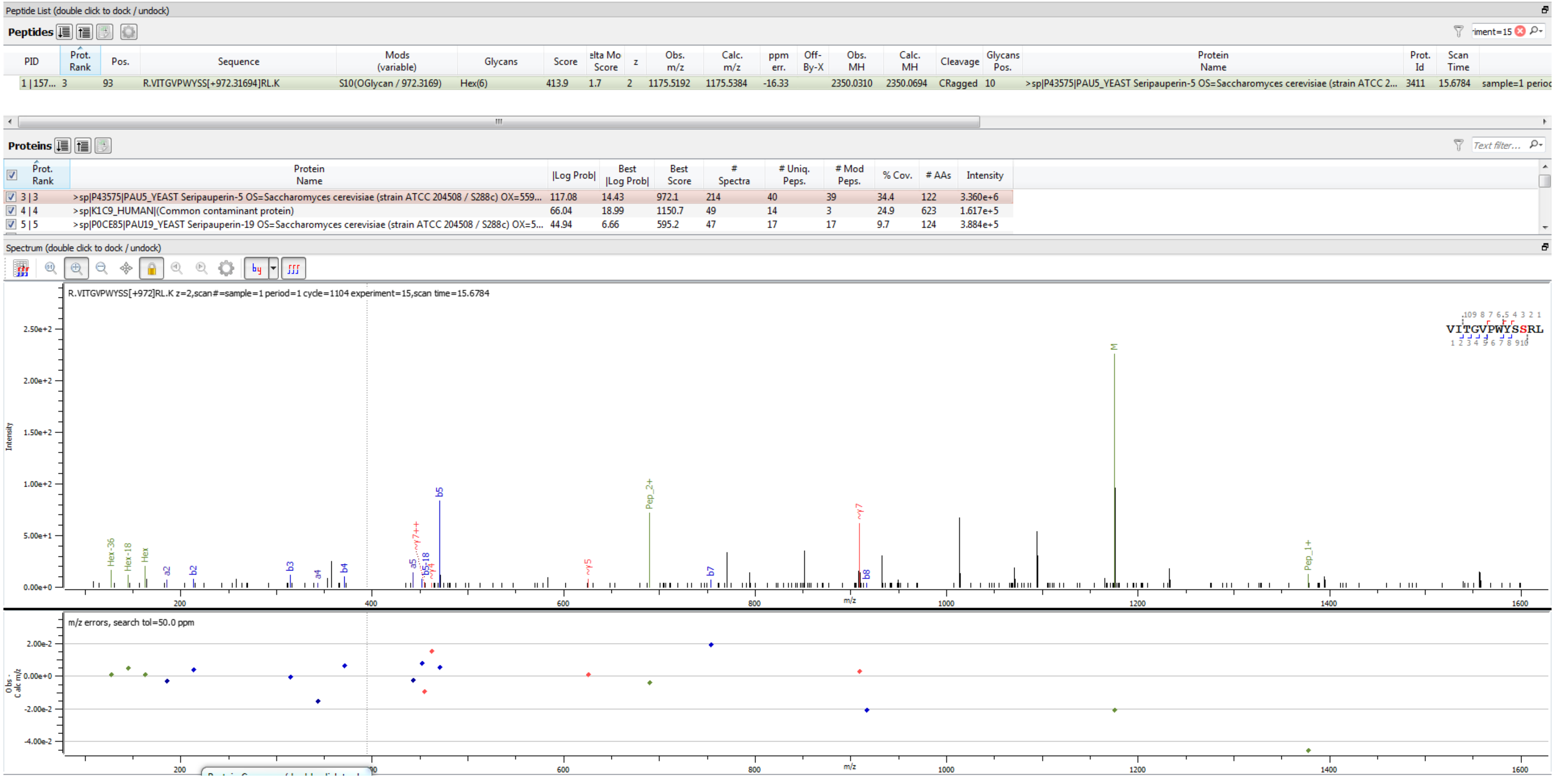


Sample S2 R.VITGVPWYSS[+972.317]RL.K

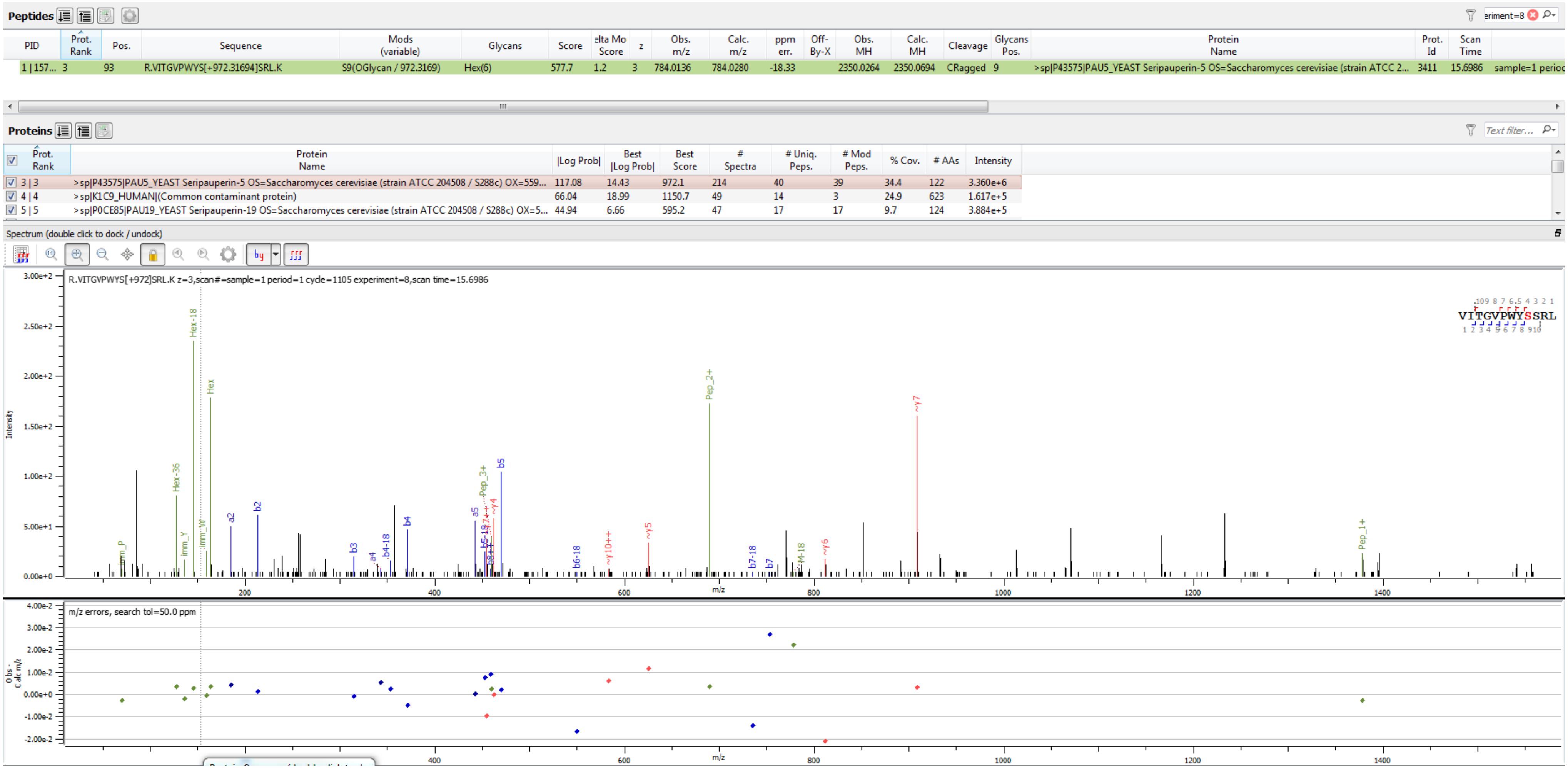


Sample S2 R.VITGVPWYS[+972.317]SRL.K

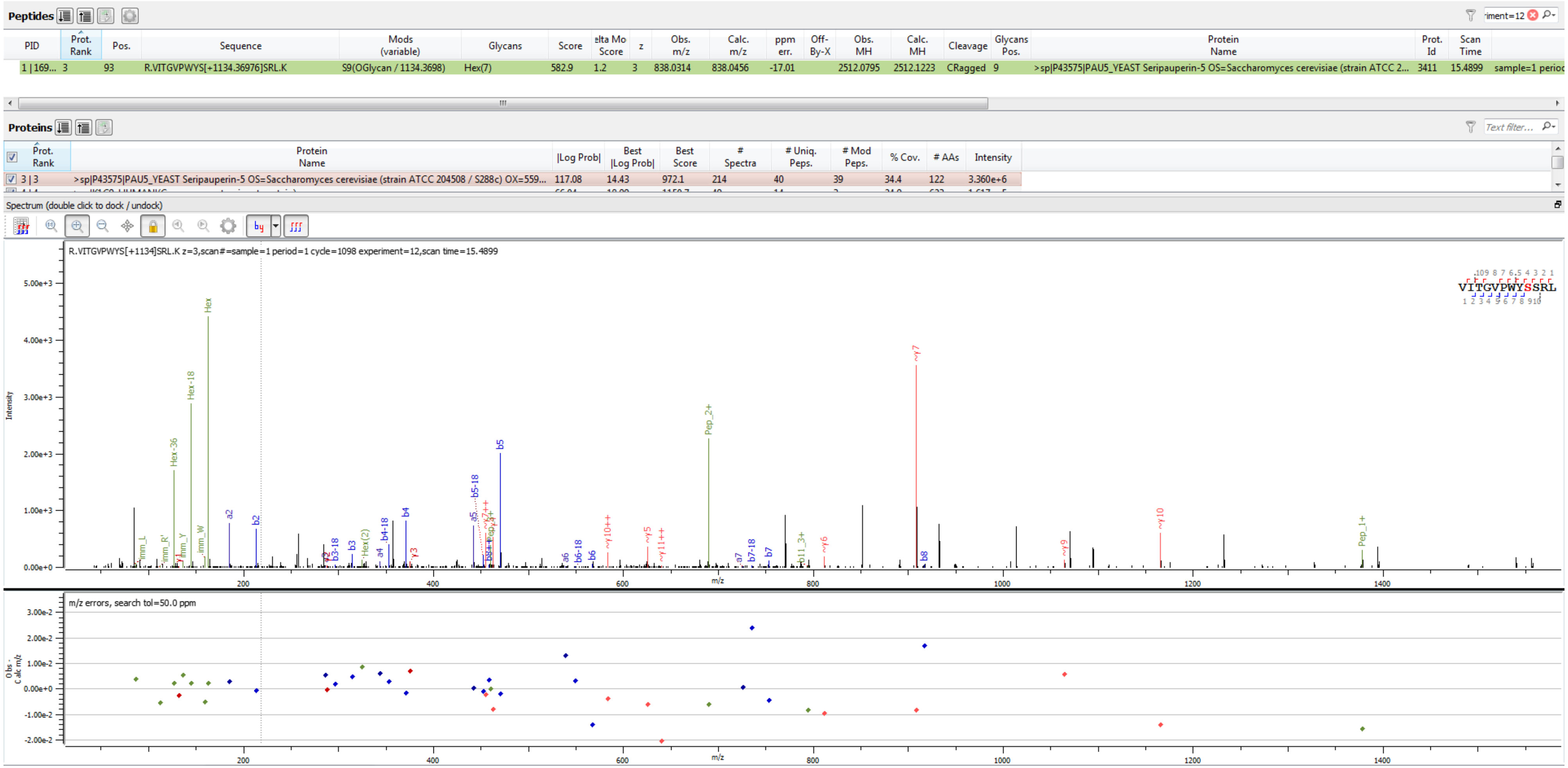


Sample S2 R.VITGVPWYS[+1134.370]SRL.K

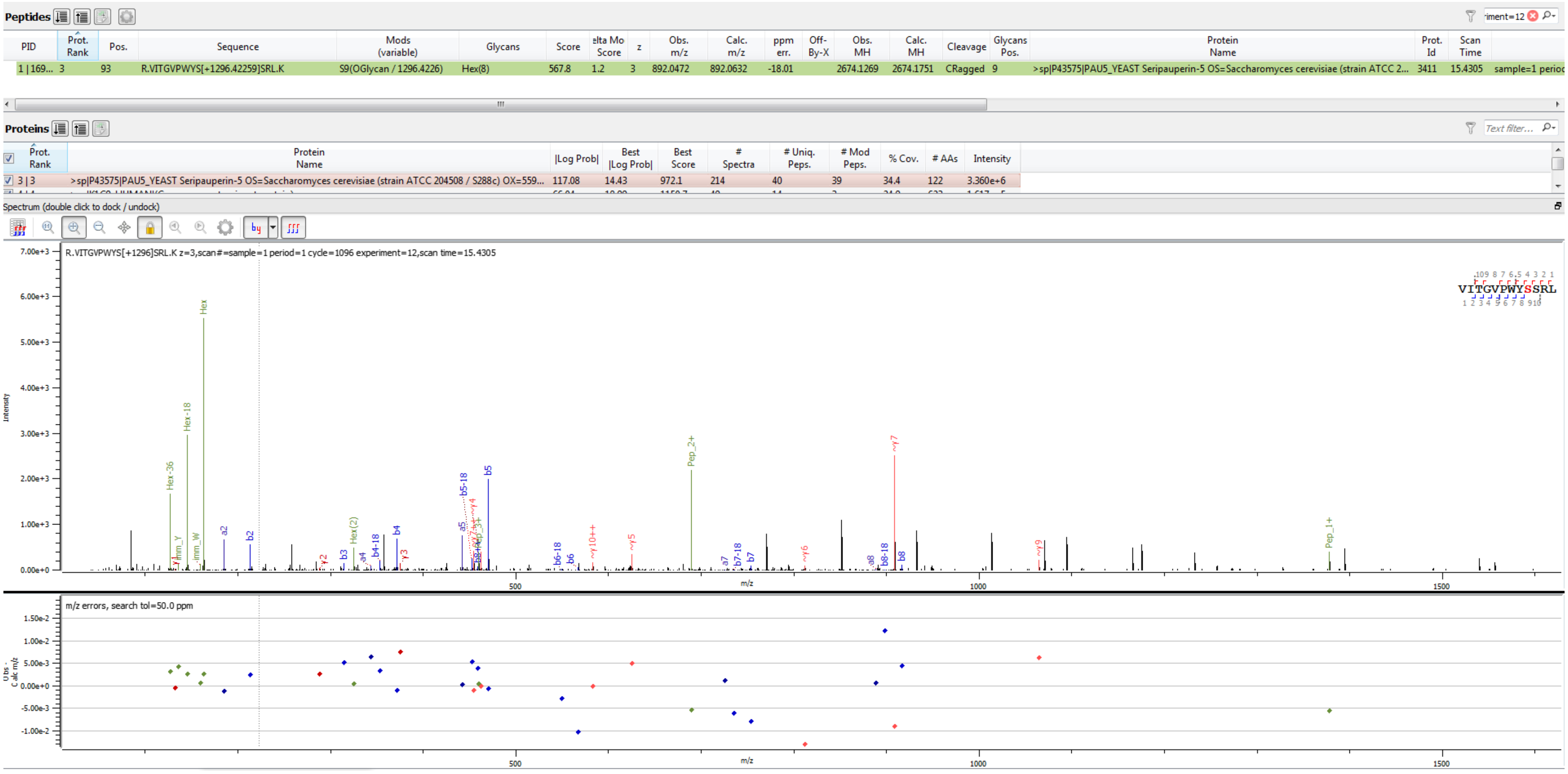


Sample S2 R.VITGVPWYS[+1296.423]SRL.K

**Peptides**

| PID | Prot. Rank | Pos. | Sequence | Mods (variable) | Glycans | Score | elta Mo Score | z | Obs. m/z | Calc. m/z | ppm err. | Off-By-X | Obs. MH | Calc. MH | Cleavage | Glycans Pos. | Protein Name | Prot. Id | Scan Time | |
|---|---|---|---|---|---|---|---|---|---|---|---|---|---|---|---|---|---|---|---|---|
| 1 \| 121... | 1 | 91 | R.MITGVPWYSS[+486.15847]R.L | S10(OGlycan / 486.1585) | Hex(3) | 686.1 | 0.0 | 2 | 891.8856 | 891.9031 | -19.62 | | 1782.7640 | 1782.7990 | Specific | 10 | >sp\|Q3E770\|PAU9_YEAST Seripauperin-9 OS=Saccharomyces cerevisiae (strain ATCC 2... | 3461 | 15.1760 | sample=1 perioc |

**Proteins**

| | Prot. Rank | Protein Name | \|Log Prob\| | Best \|Log Prob\| | Best Score | # Spectra | # Uniq. Peps. | # Mod Peps. | % Cov. | # AAs | Intensity |
|---|---|---|---|---|---|---|---|---|---|---|---|
| ☑ | 1 \| 1 | >sp\|Q3E770\|PAU9_YEAST Seripauperin-9 OS=Saccharomyces cerevisiae (strain ATCC 204508 / S288c) OX=559... | 248.32 | 15.68 | 955.8 | 342 | 58 | 55 | 67.5 | 120 | 1.797e+7 |

Spectrum (double click to dock / undock)

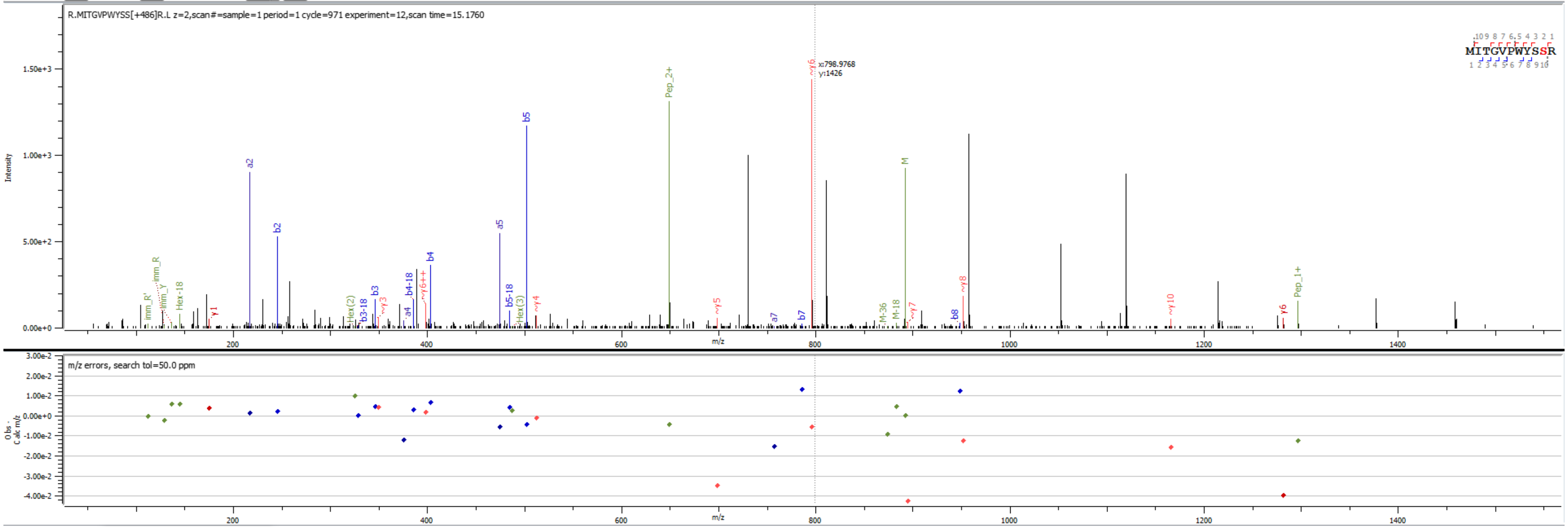


Sample S3 R.MITGVPWYSS[+486.158]R.L

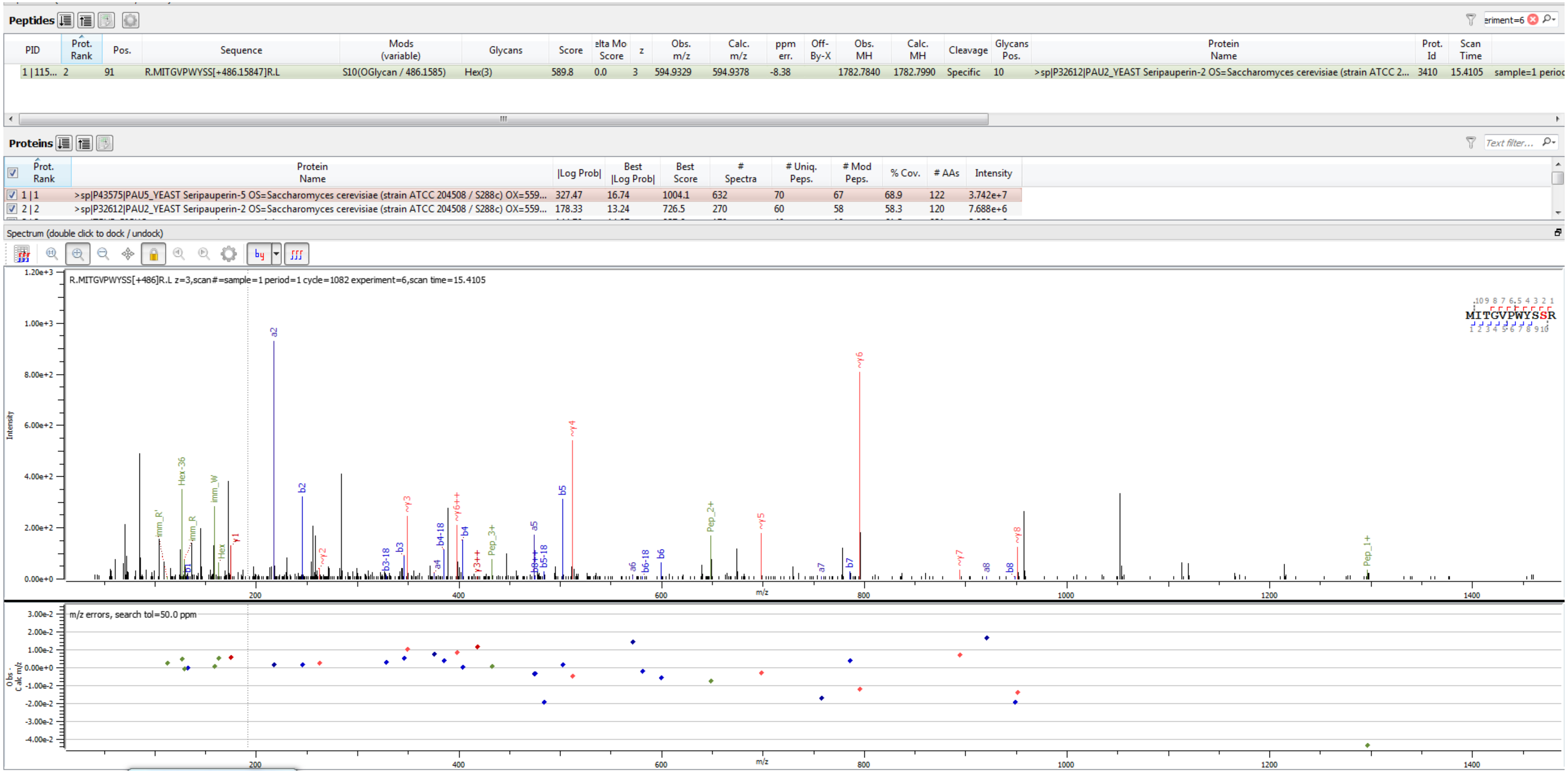


Sample S1 R.MITGVPWYSS[+486.158]R.L

**Peptides**

| PID | Prot. Rank | Pos. | Sequence | Mods (variable) | Glycans | Score | elta Mo Score | z | Obs. m/z | Calc. m/z | ppm err. | Off-By-X | Obs. MH | Calc. MH | Cleavage | Glycans Pos. | Protein Name | Prot. Id | Scan Time | |
|---|---|---|---|---|---|---|---|---|---|---|---|---|---|---|---|---|---|---|---|---|
| 1 \| 113... | 2 | 91 | R.MITGVPWYSS[+648.21129]R.L | S10(OGlycan / 648.2113) | Hex(4) | 670.7 | 0.0 | 2 | 972.9174 | 972.9295 | -12.52 | | 1944.8274 | 1944.8518 | Specific | 10 | >sp\|P32612\|PAU2_YEAST Seripauperin-2 OS=Saccharomyces cerevisiae (strain ATCC 2... | 3410 | 15.2446 | sample=1 perioc |

**Proteins**

| Prot. Rank | Protein Name | \|Log Prob\| | Best \|Log Prob\| | Best Score | # Spectra | # Uniq. Peps. | # Mod Peps. | % Cov. | # AAs | Intensity |
|---|---|---|---|---|---|---|---|---|---|---|
| 1 \| 1 | >sp\|P43575\|PAU5_YEAST Seripauperin-5 OS=Saccharomyces cerevisiae (strain ATCC 204508 / S288c) OX=559... | 327.47 | 16.74 | 1004.1 | 632 | 70 | 67 | 68.9 | 122 | 3.742e+7 |
| 2 \| 2 | >sp\|P32612\|PAU2_YEAST Seripauperin-2 OS=Saccharomyces cerevisiae (strain ATCC 204508 / S288c) OX=559... | 178.33 | 13.24 | 726.5 | 270 | 60 | 58 | 58.3 | 120 | 7.688e+6 |

Spectrum (double click to dock / undock)

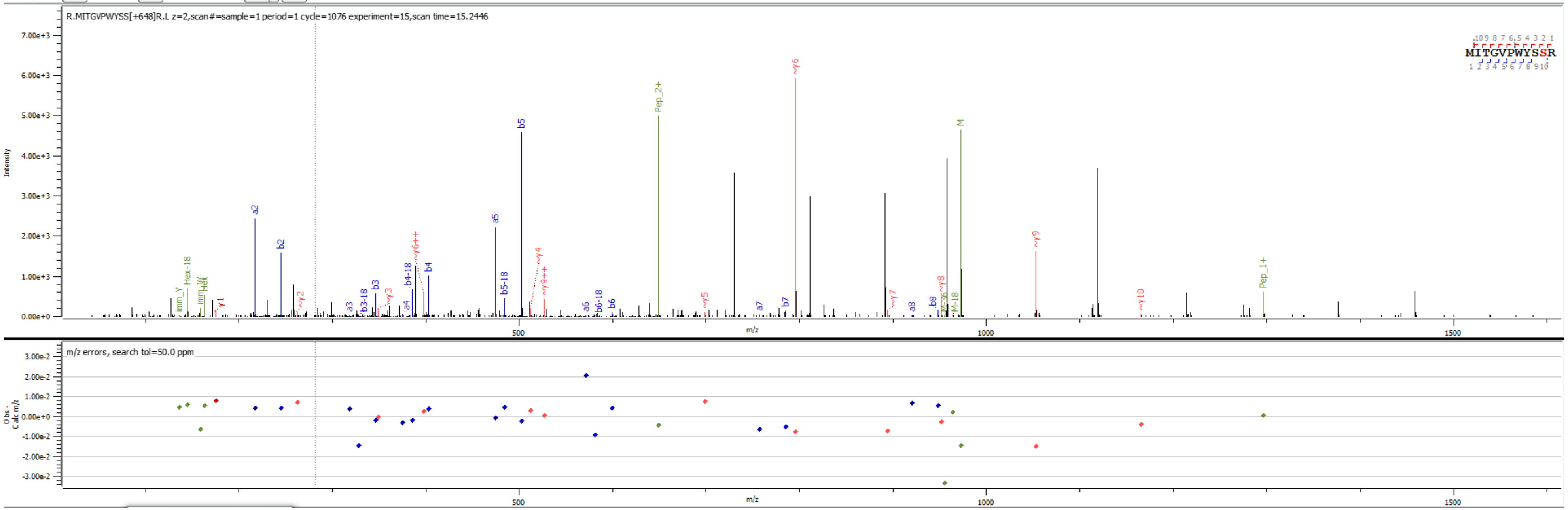


Sample S1 R.MITGVPWYSS[+648.211]R.L

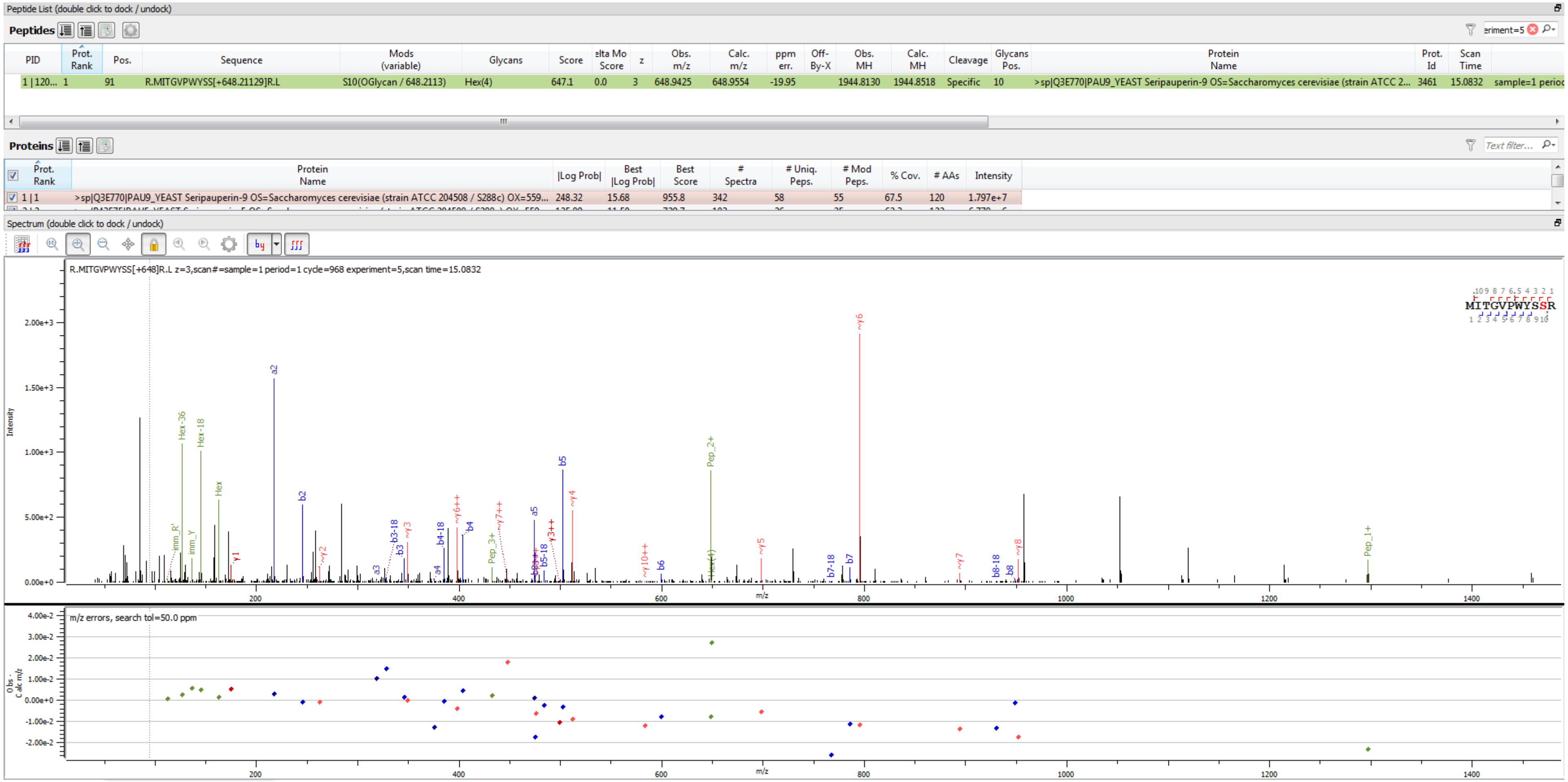


Sample S3 R.MITGVPWYSS[+648.211]R.L

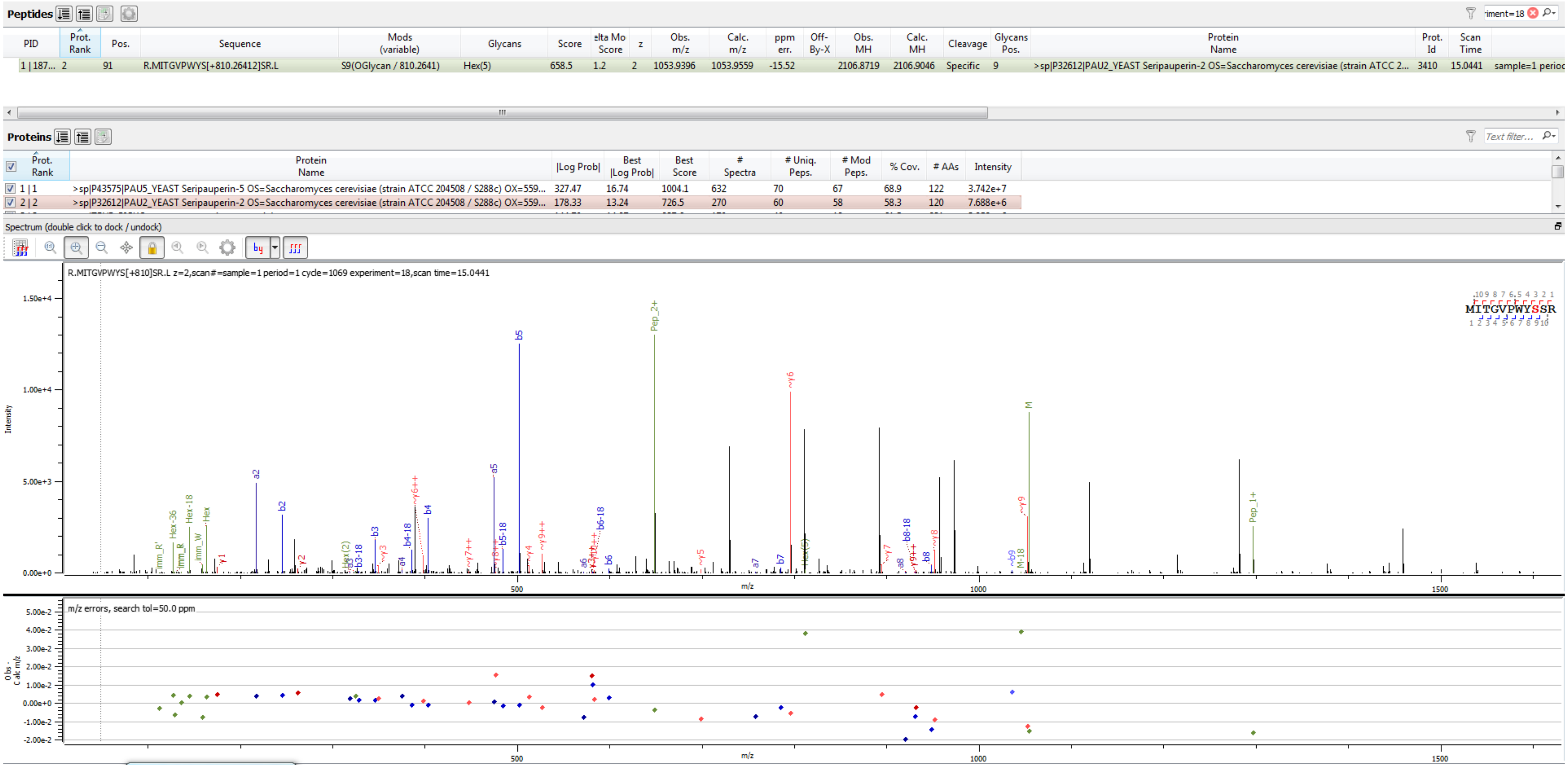


Sample S1 R.MITGVPWYS[+810.264]SR.L

**Peptides**

| PID | Prot. Rank | Pos. | Sequence | Mods (variable) | Glycans | Score | elta Mo Score | z | Obs. m/z | Calc. m/z | ppm err. | Off-By-X | Obs. MH | Calc. MH | Cleavage | Glycans Pos. | Protein Name | Prot. Id | Scan Time | |
|---|---|---|---|---|---|---|---|---|---|---|---|---|---|---|---|---|---|---|---|---|
| 1 \| 187... | 2 | 91 | R.MITGVPWYS[+810.26412]SR.L | S9(OGlycan / 810.2641) | Hex(5) | 726.5 | 1.2 | 3 | 702.9597 | 702.9731 | -19.03 | | 2106.8645 | 2106.9046 | Specific | 9 | >sp\|P32612\|PAU2_YEAST Seripauperin-2 OS=Saccharomyces cerevisiae (strain ATCC 2... | 3410 | 15.0671 | sample=1 perioc |

**Proteins**

| Prot. Rank | Protein Name | \|Log Prob\| | Best \|Log Prob\| | Best Score | # Spectra | # Uniq. Peps. | # Mod Peps. | % Cov. | # AAs | Intensity |
|---|---|---|---|---|---|---|---|---|---|---|
| 1 \| 1 | >sp\|P43575\|PAU5_YEAST Seripauperin-5 OS=Saccharomyces cerevisiae (strain ATCC 204508 / S288c) OX=559... | 327.47 | 16.74 | 1004.1 | 632 | 70 | 67 | 68.9 | 122 | 3.742e+7 |
| 2 \| 2 | >sp\|P32612\|PAU2_YEAST Seripauperin-2 OS=Saccharomyces cerevisiae (strain ATCC 204508 / S288c) OX=559... | 178.33 | 13.24 | 726.5 | 270 | 60 | 58 | 58.3 | 120 | 7.688e+6 |

Spectrum (double click to dock / undock)

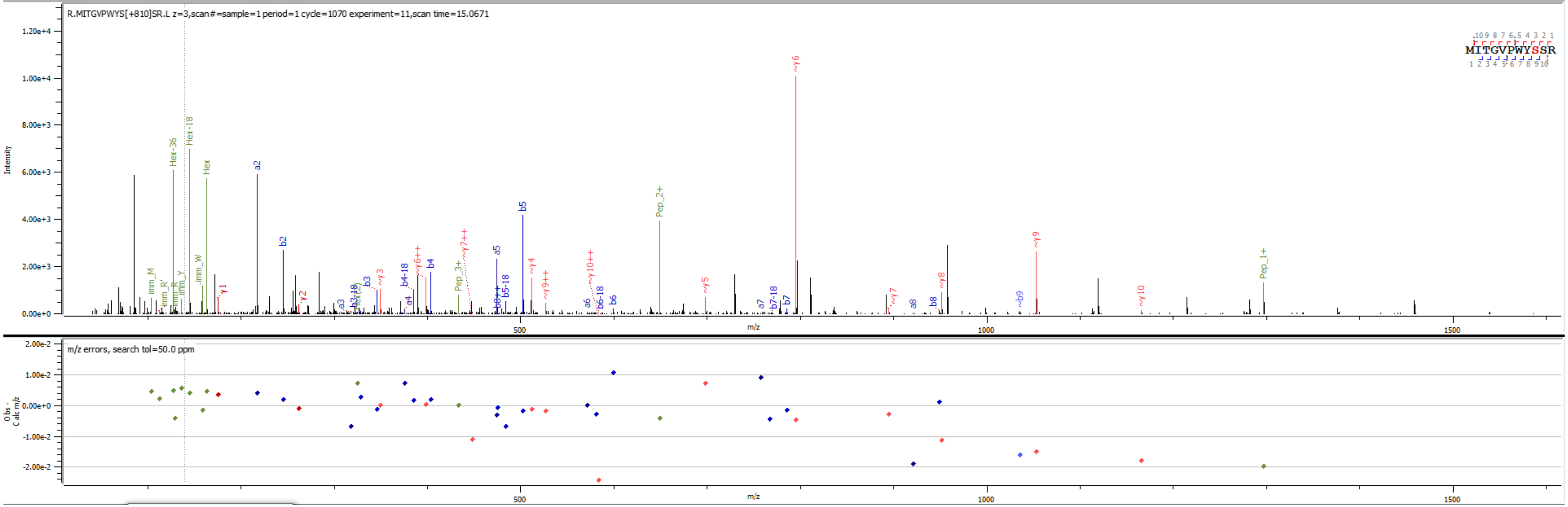


Sample S1 R.MITGVPWYS[+810.264]SR.L

**Peptides**

| PID | Prot. Rank | Pos. | Sequence | Mods (variable) | Glycans | Score | elta Mo Score | z | Obs. m/z | Calc. m/z | ppm err. | Off-By-X | Obs. MH | Calc. MH | Cleavage | Glycans Pos. | Protein Name | Prot. Id | Scan Time | |
|---|---|---|---|---|---|---|---|---|---|---|---|---|---|---|---|---|---|---|---|---|
| 1 \| 186... | 2 | 91 | R.MITGVPWYSS[+972.31694]R.L | S10(OGlycan / 972.3169) | Hex(6) | 622.3 | 0.8 | 2 | 1134.9622 | 1134.9824 | -17.74 | | 2268.9172 | 2268.9574 | Specific | 10 | >sp\|P32612\|PAU2_YEAST Seripauperin-2 OS=Saccharomyces cerevisiae (strain ATCC 2... | 3410 | 14.8720 | sample=1 perioc |

**Proteins**

| Prot. Rank | Protein Name | \|Log Prob\| | Best \|Log Prob\| | Best Score | # Spectra | # Uniq. Peps. | # Mod Peps. | % Cov. | # AAs | Intensity |
|---|---|---|---|---|---|---|---|---|---|---|
| 1 \| 1 | >sp\|P43575\|PAU5_YEAST Seripauperin-5 OS=Saccharomyces cerevisiae (strain ATCC 204508 / S288c) OX=559... | 327.47 | 16.74 | 1004.1 | 632 | 70 | 67 | 68.9 | 122 | 3.742e+7 |
| 2 \| 2 | >sp\|P32612\|PAU2_YEAST Seripauperin-2 OS=Saccharomyces cerevisiae (strain ATCC 204508 / S288c) OX=559... | 178.33 | 13.24 | 726.5 | 270 | 60 | 58 | 58.3 | 120 | 7.688e+6 |

Spectrum (double click to dock / undock)

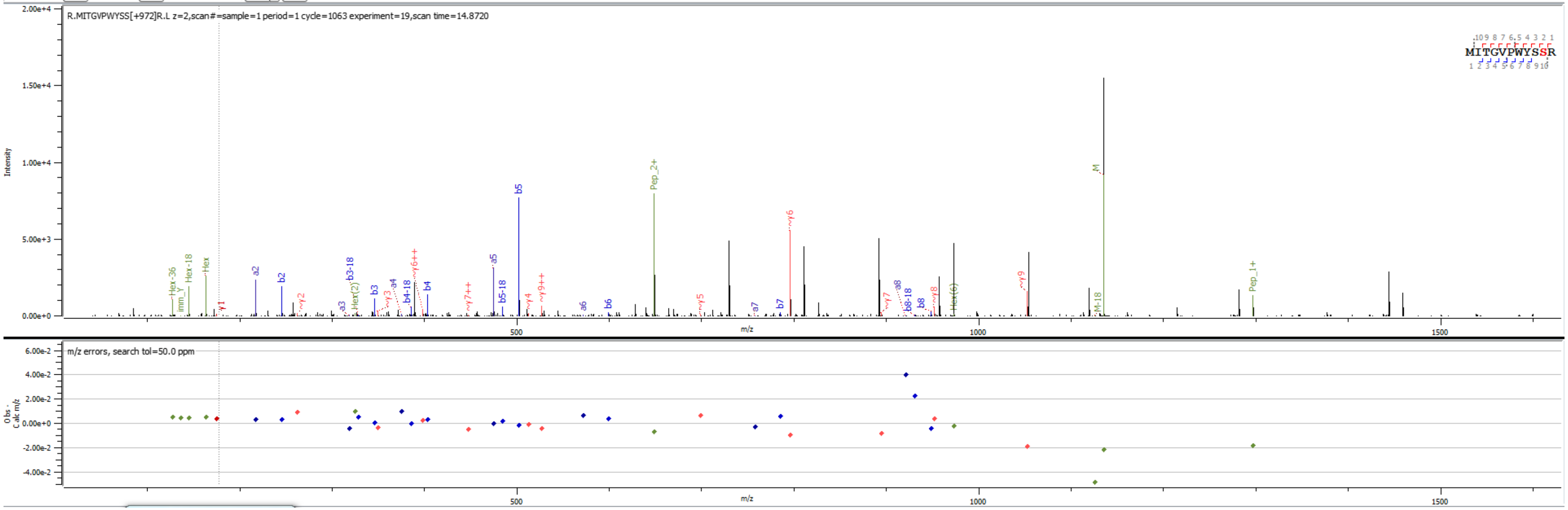


Sample S1 R.MITGVPWYS[+972.317]SR.L

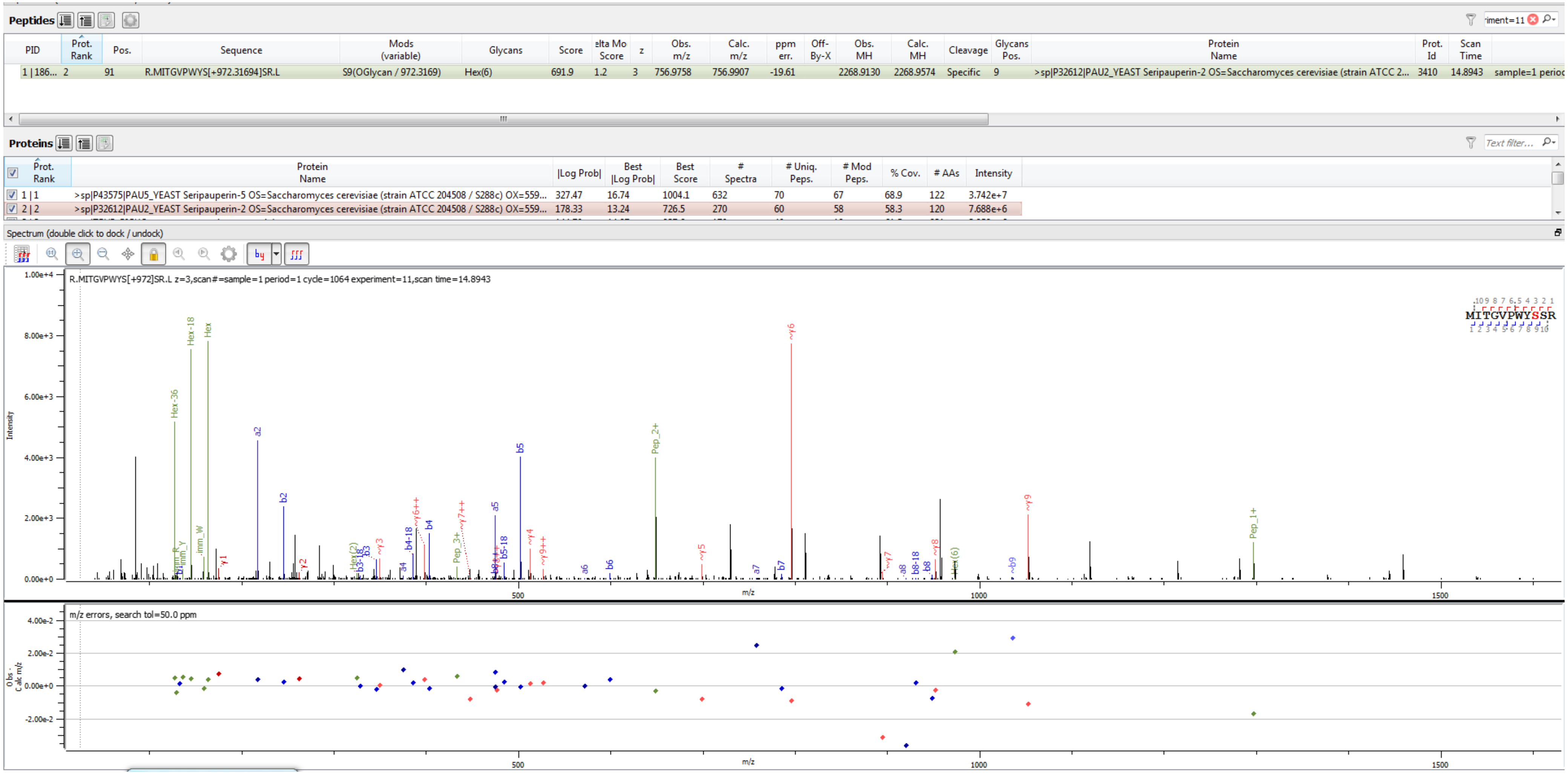


Sample S1 R.MITGVPWYS[+972.317]SR.L

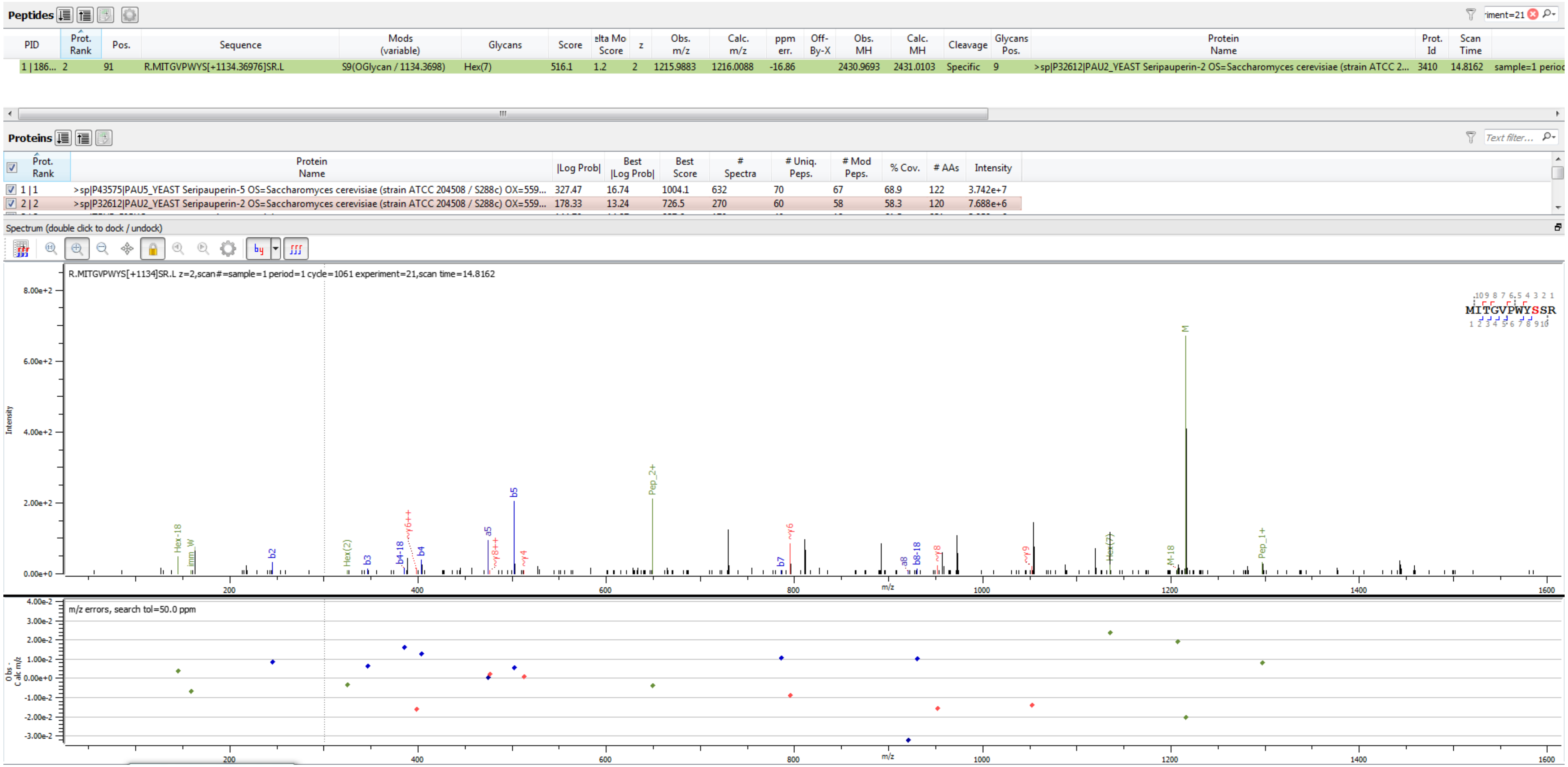


Sample S1 R.MITGVPWYS[+1134.370]SR.L

**Peptides**

| PID | Prot. Rank | Pos. | Sequence | Mods (variable) | Glycans | Score | elta Mo Score | z | Obs. m/z | Calc. m/z | ppm err. | Off-By-X | Obs. MH | Calc. MH | Cleavage | Glycans Pos. | Protein Name | Prot. Id | Scan Time | |
|---|---|---|---|---|---|---|---|---|---|---|---|---|---|---|---|---|---|---|---|---|
| 1 \| 132... | 1 | 91 | R.MITGVPWYS[+1134.36976]SR.L | S9(OGlycan / 1134.3698) | Hex(7) | 636.2 | 1.2 | 3 | 810.9955 | 811.0083 | -15.75 | | 2430.9720 | 2431.0103 | Specific | 9 | >sp\|Q3E770\|PAU9_YEAST Seripauperin-9 OS=Saccharomyces cerevisiae (strain ATCC 2... | 3461 | 14.7986 | sample=1 perioc |

**Proteins**

| Prot. Rank | Protein Name | \|Log Prob\| | Best \|Log Prob\| | Best Score | # Spectra | # Uniq. Peps. | # Mod Peps. | % Cov. | # AAs | Intensity |
|---|---|---|---|---|---|---|---|---|---|---|
| 1 \| 1 | >sp\|Q3E770\|PAU9_YEAST Seripauperin-9 OS=Saccharomyces cerevisiae (strain ATCC 204508 / S288c) OX=559... | 288.60 | 16.66 | 988.6 | 452 | 61 | 56 | 60.8 | 120 | 1.023e+7 |
| 2 \| 2 | >sp\|Q07987\|PAU23_YEAST Seripauperin-23 OS=Saccharomyces cerevisiae (strain ATCC 204508 / S288c) OX=5... | 160.38 | 13.28 | 718.6 | 124 | 39 | 36 | 43.5 | 124 | 2.323e+6 |
| 3 \| 3 | >sp\|P43575\|PAU5_YEAST Seripauperin-5 OS=Saccharomyces cerevisiae (strain ATCC 204508 / S288c) OX=559... | 142.55 | 14.15 | 737.1 | 167 | 35 | 33 | 42.6 | 122 | 6.136e+6 |
| 4 \| 4 | >sp\|K2C1_HUMAN\|(Common contaminant protein) | 101.54 | 13.43 | 953.1 | 37 | 11 | 0 | 20.4 | 643 | 1.801e+5 |

Spectrum (double click to dock / undock)

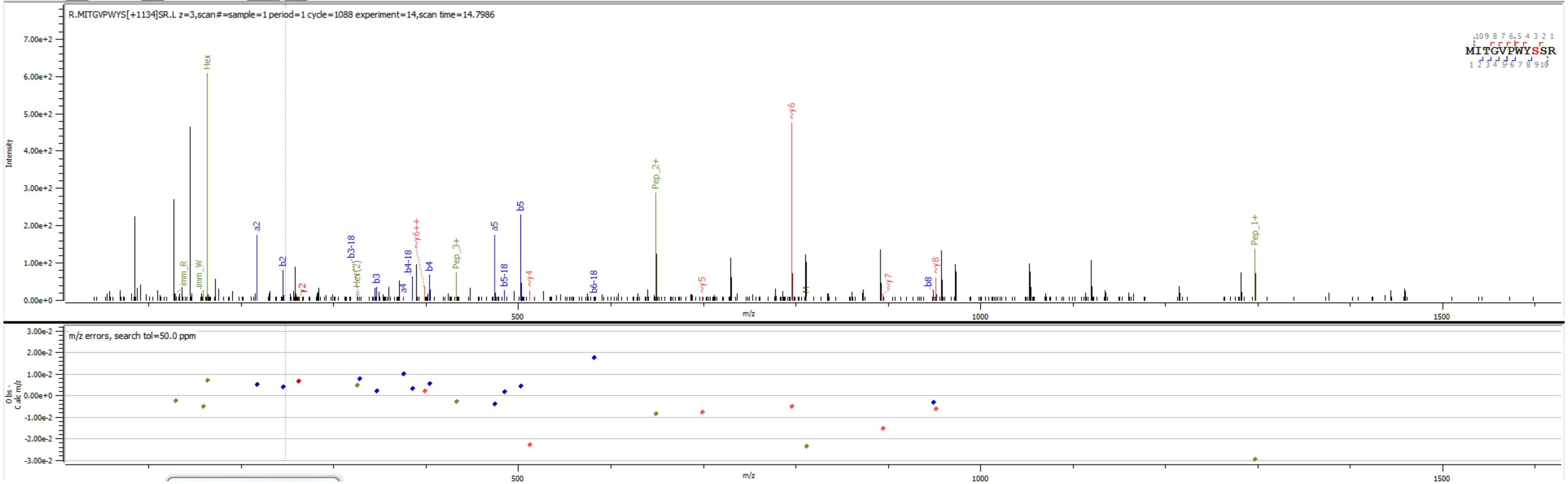


Sample S6 R.MITGVPWYS[+1134.370]SR.L

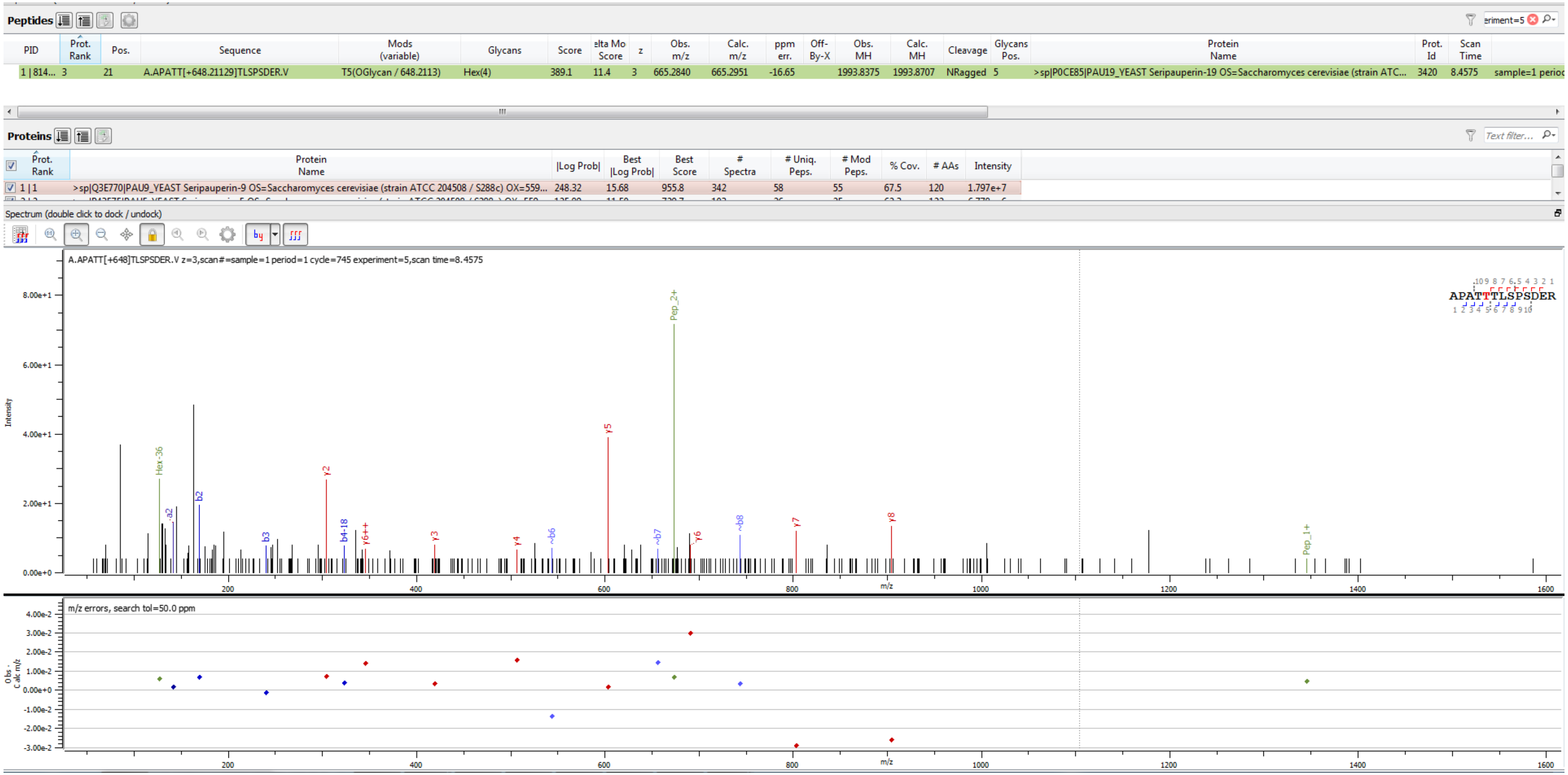


Sample S3 A.APATT[+648.211]TLSPSDER.V

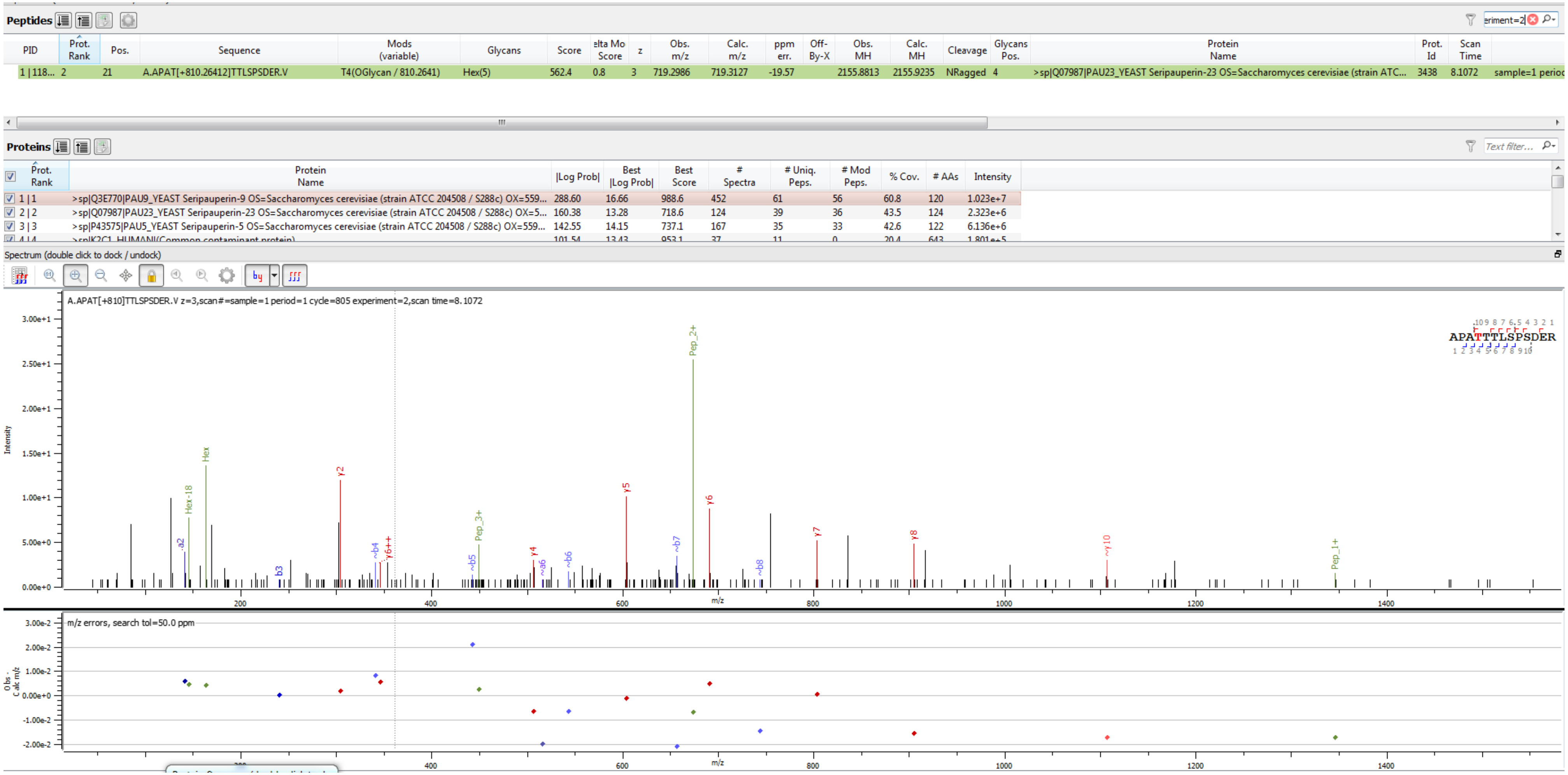


Sample S6 A.APAT[+810.264]TTLSPSDER.V

**Peptides**

| PID | Prot. Rank | Pos. | Sequence | Mods (variable) | Glycans | Score | elta Mo Score | z | Obs. m/z | Calc. m/z | ppm err. | Off-By-X | Obs. MH | Calc. MH | Cleavage | Glycans Pos. | Protein Name | Prot. Id | Scan Time | |
|---|---|---|---|---|---|---|---|---|---|---|---|---|---|---|---|---|---|---|---|---|
| 1 \| 118... | 2 | 21 | A.APATTT[+972.31694]LSPSDER.V | T6(OGlycan / 972.3169) | Hex(6) | 702.4 | 0.0 | 3 | 773.3154 | 773.3303 | -19.32 | | 2317.9315 | 2317.9763 | NRagged | 6 | >sp\|Q07987\|PAU23_YEAST Seripauperin-23 OS=Saccharomyces cerevisiae (strain ATC... | 3438 | 8.1732 | sample=1 perioc |

**Proteins**

| Prot. Rank | Protein Name | \|Log Prob\| | Best \|Log Prob\| | Best Score | # Spectra | # Uniq. Peps. | # Mod Peps. | % Cov. | # AAs | Intensity |
|---|---|---|---|---|---|---|---|---|---|---|
| 1 \| 1 | >sp\|Q3E770\|PAU9_YEAST Seripauperin-9 OS=Saccharomyces cerevisiae (strain ATCC 204508 / S288c) OX=559... | 288.60 | 16.66 | 988.6 | 452 | 61 | 56 | 60.8 | 120 | 1.023e+7 |
| 2 \| 2 | >sp\|Q07987\|PAU23_YEAST Seripauperin-23 OS=Saccharomyces cerevisiae (strain ATCC 204508 / S288c) OX=5... | 160.38 | 13.28 | 718.6 | 124 | 39 | 36 | 43.5 | 124 | 2.323e+6 |
| 3 \| 3 | >sp\|P43575\|PAU5_YEAST Seripauperin-5 OS=Saccharomyces cerevisiae (strain ATCC 204508 / S288c) OX=559... | 142.55 | 14.15 | 737.1 | 167 | 35 | 33 | 42.6 | 122 | 6.136e+6 |
| 4 \| 4 | >sp\|K2C1_HUMAN\|(Common contaminant protein) | 101.54 | 13.43 | 953.1 | 37 | 11 | 0 | 20.4 | 643 | 1.801e+5 |

Spectrum (double click to dock / undock)

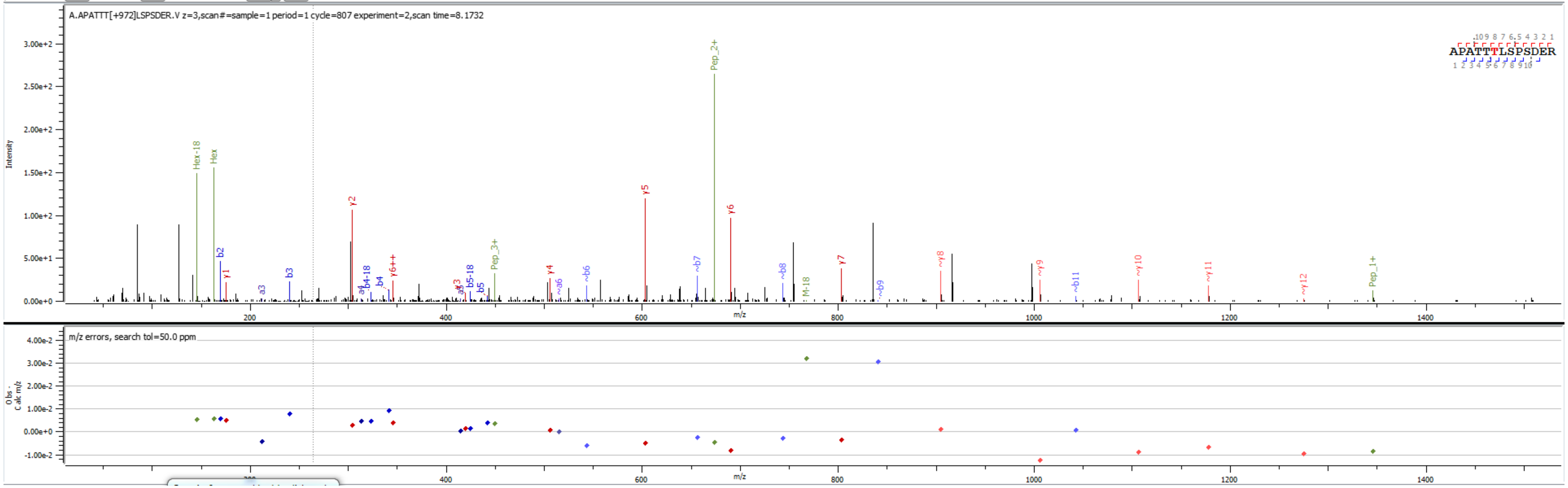


Sample S6 A.APATTT[+972.317]LSPSDER.V

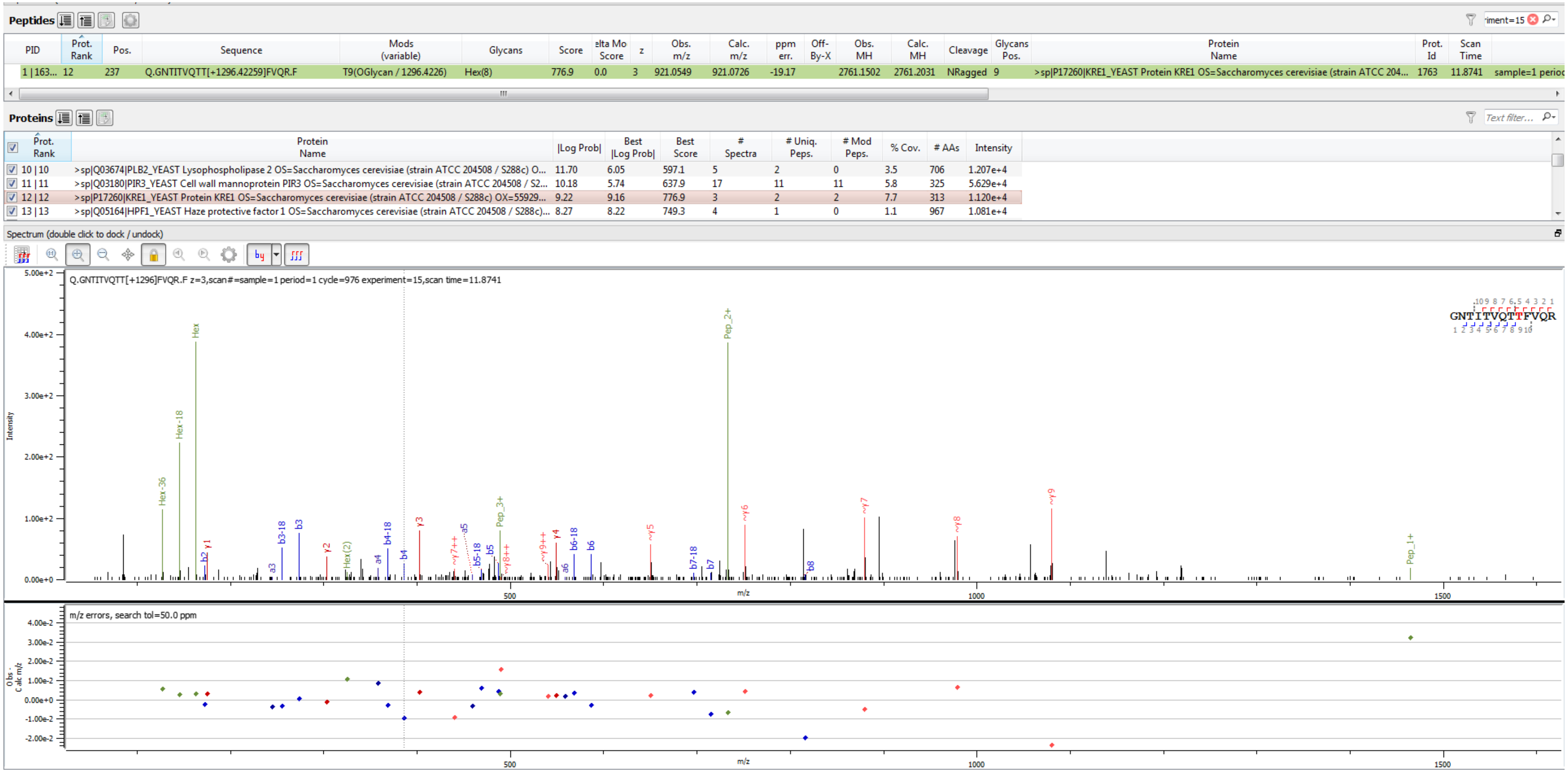


Sample S2 Q.GNTITVQTT[+1296.423]FVQR.F

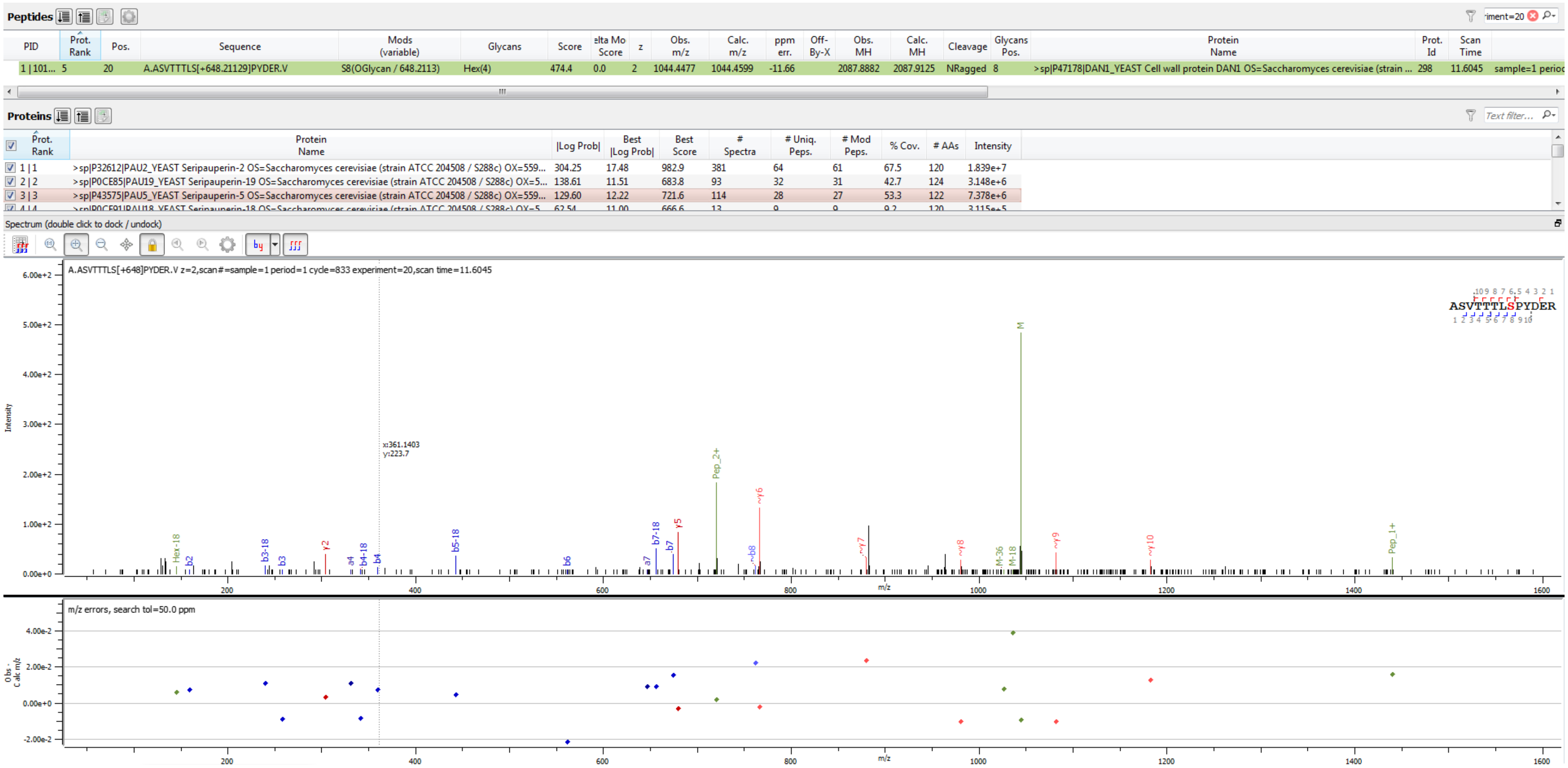


Sample S4 A.ASVTTTLS[+648.211]PYDER.V

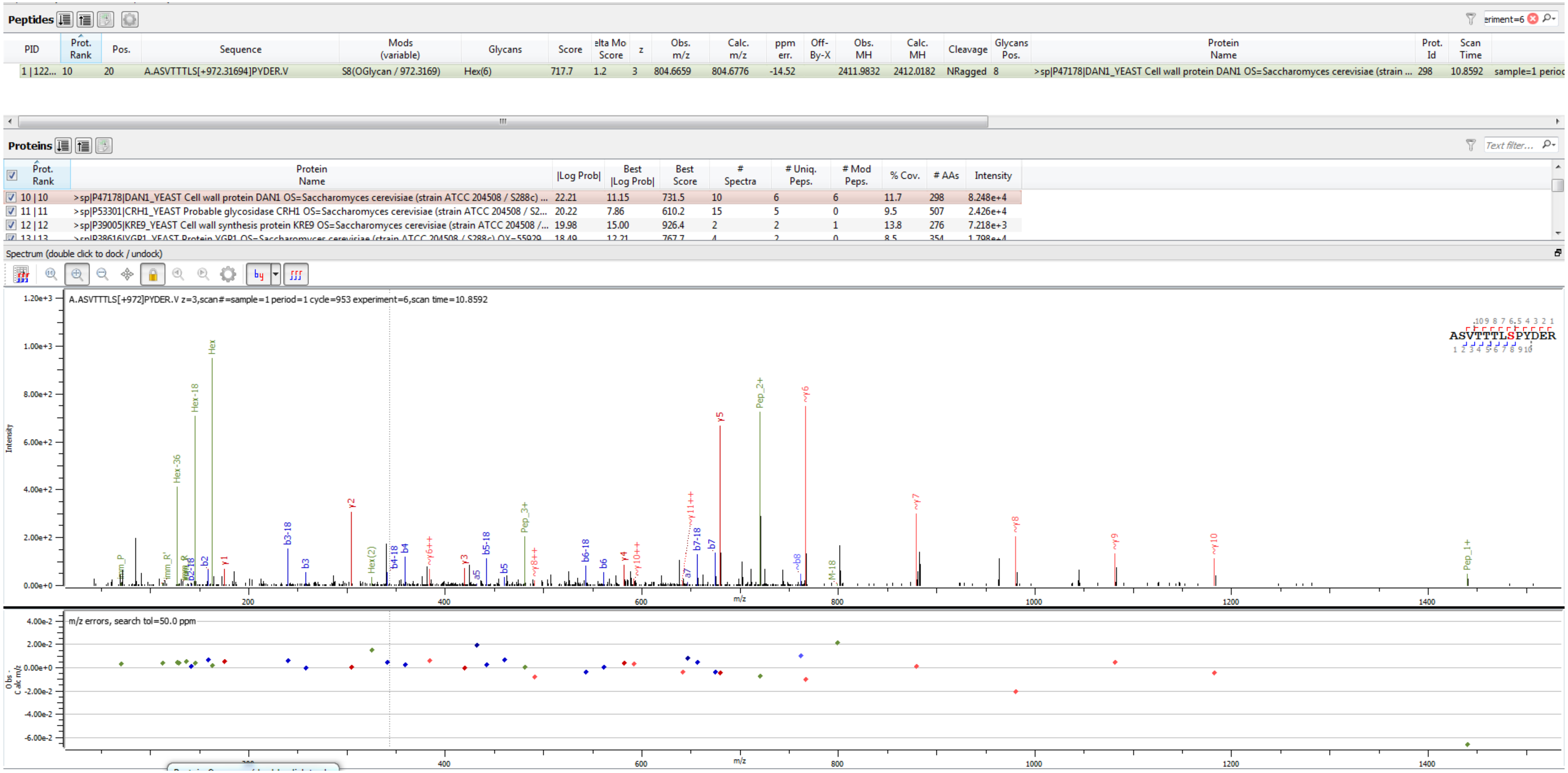


Sample S6 A.ASVTTTLS[+972.317]PYDER.V

**Peptides**

| PID | Prot. Rank | Pos. | Sequence | Mods (variable) | Glycans | Score | elta Mo Score | z | Obs. m/z | Calc. m/z | ppm err. | Off-By-X | Obs. MH | Calc. MH | Cleavage | Glycans Pos. | Protein Name | Prot. Id | Scan Time | |
|---|---|---|---|---|---|---|---|---|---|---|---|---|---|---|---|---|---|---|---|---|
| 1 \| 356... | 2 | 116 | K.DGIYT[+324.10565]AIPK.- | T5(OGlycan / 324.1056) | Hex(2) | 683.8 | 321.7 | 2 | 651.3173 | 651.3216 | -6.59 | | 1301.6273 | 1301.6358 | Specific | 5 | >sp\|P0CE85\|PAU19_YEAST Seripauperin-19 OS=Saccharomyces cerevisiae (strain ATC... | 3420 | 13.3804 | sample=1 perioc |

**Proteins**

| Prot. Rank | Protein Name | \|Log Prob\| | Best \|Log Prob\| | Best Score | # Spectra | # Uniq. Peps. | # Mod Peps. | % Cov. | # AAs | Intensity |
|---|---|---|---|---|---|---|---|---|---|---|
| 1 \| 1 | >sp\|P32612\|PAU2_YEAST Seripauperin-2 OS=Saccharomyces cerevisiae (strain ATCC 204508 / S288c) OX=559... | 304.25 | 17.48 | 982.9 | 381 | 64 | 61 | 67.5 | 120 | 1.839e+7 |
| 2 \| 2 | >sp\|P0CE85\|PAU19_YEAST Seripauperin-19 OS=Saccharomyces cerevisiae (strain ATCC 204508 / S288c) OX=5... | 138.61 | 11.51 | 683.8 | 93 | 32 | 31 | 42.7 | 124 | 3.148e+6 |
| 3 \| 3 | >sp\|P43575\|PAU5_YEAST Seripauperin-5 OS=Saccharomyces cerevisiae (strain ATCC 204508 / S288c) OX=559... | 129.60 | 12.22 | 721.6 | 114 | 28 | 27 | 53.3 | 122 | 7.378e+6 |
| 4 \| 4 | >sp\|P0CE91\|PAU18_YEAST Seripauperin-18 OS=Saccharomyces cerevisiae (strain ATCC 204508 / S288c) OX=5... | 62.54 | 11.00 | 666.6 | 13 | 9 | 9 | 9.2 | 120 | 3.115e+5 |

Spectrum (double click to dock / undock)

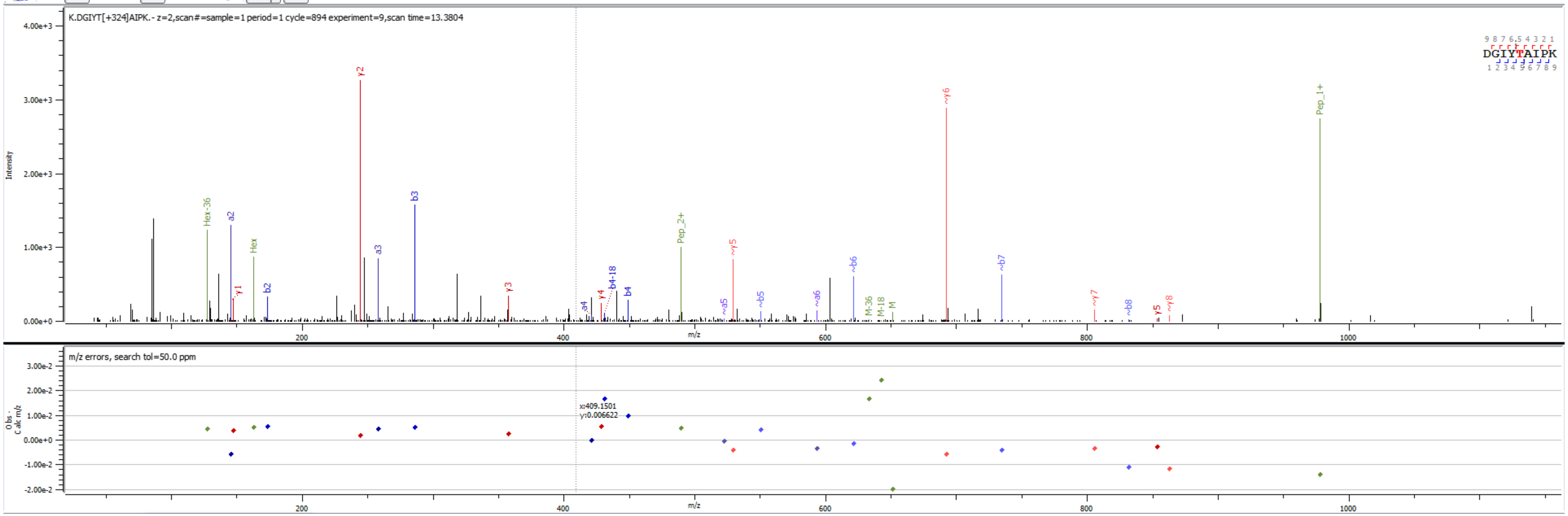


Sample S4 K.DGIYT[+324.106]AIPK.-

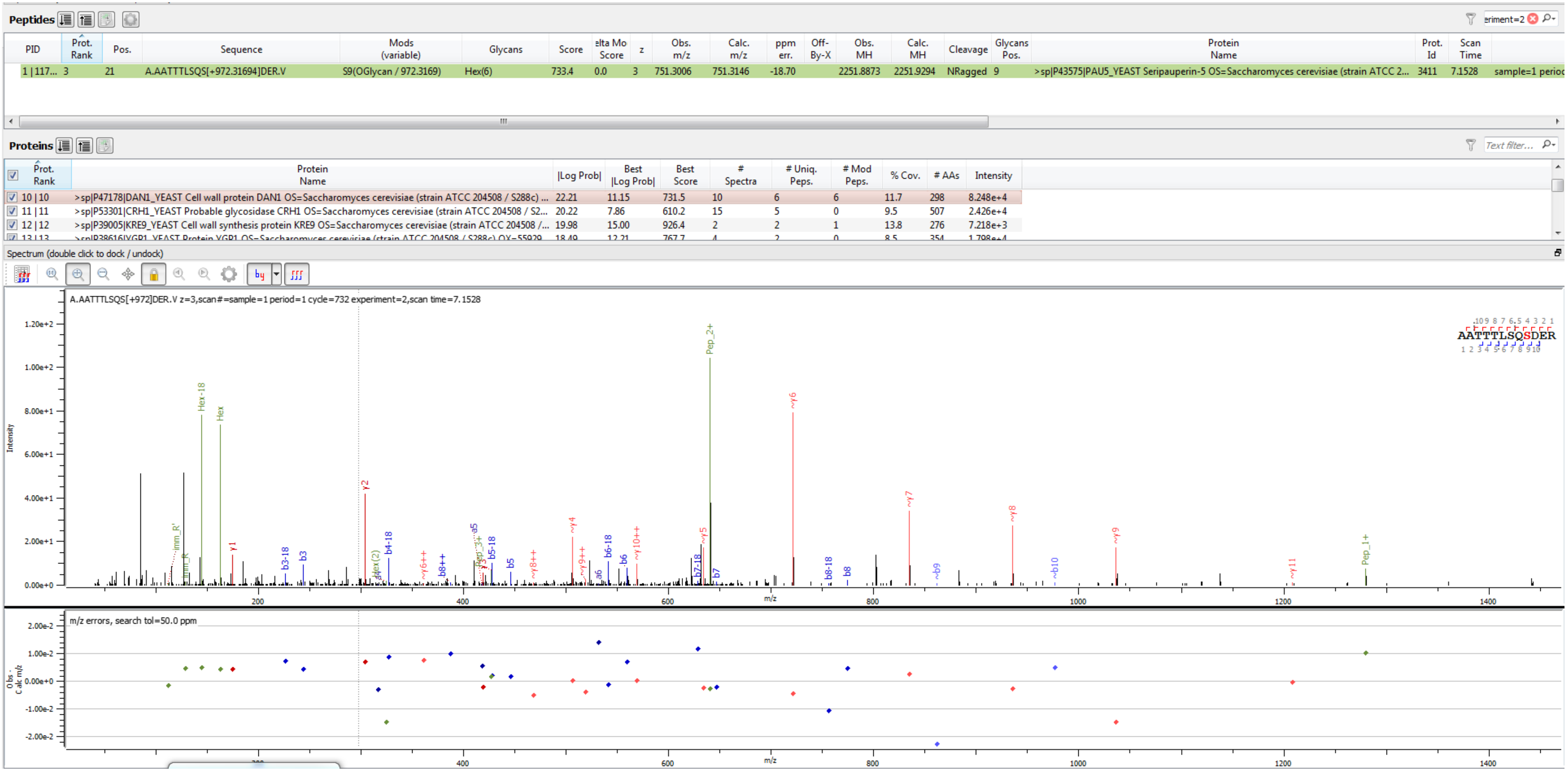


Sample S6 A.AATTTLSQS[+972.317]DER.V

**Peptides**

| PID | Prot. Rank | Pos. | Sequence | Mods (variable) | Glycans | Score | elta Mo Score | z | Obs. m/z | Calc. m/z | ppm err. | Off-By-X | Obs. MH | Calc. MH | Cleavage | Glycans Pos. | Protein Name | Prot. Id | Scan Time | |
|---|---|---|---|---|---|---|---|---|---|---|---|---|---|---|---|---|---|---|---|---|
| 1 \| 106... | 14 | 109 | K.ALSKDGIYT[+648.21129]IAN.- | T9(OGlycan / 648.2113) | Hex(4) | 519.5 | 8.6 | 2 | 957.4411 | 957.4461 | -5.16 | | 1913.8750 | 1913.8849 | Specific | 9 | >sp\|P53427\|PAU4_YEAST Seripauperin-4 OS=Saccharomyces cerevisiae (strain ATCC 2... | 3439 | 12.3100 | sample=1 perioc |

**Proteins**

| Prot. Rank | Protein Name | \|Log Prob\| | Best \|Log Prob\| | Best Score | # Spectra | # Uniq. Peps. | # Mod Peps. | % Cov. | # AAs | Intensity |
|---|---|---|---|---|---|---|---|---|---|---|
| 1 \| 1 | >sp\|P32612\|PAU2_YEAST Seripauperin-2 OS=Saccharomyces cerevisiae (strain ATCC 204508 / S288c) OX=559... | 304.25 | 17.48 | 982.9 | 381 | 64 | 61 | 67.5 | 120 | 1.839e+7 |
| 2 \| 2 | >sp\|P0CE85\|PAU19_YEAST Seripauperin-19 OS=Saccharomyces cerevisiae (strain ATCC 204508 / S288c) OX=5... | 138.61 | 11.51 | 683.8 | 93 | 32 | 31 | 42.7 | 124 | 3.148e+6 |
| 3 \| 3 | >sp\|P43575\|PAU5_YEAST Seripauperin-5 OS=Saccharomyces cerevisiae (strain ATCC 204508 / S288c) OX=559... | 129.60 | 12.22 | 721.6 | 114 | 28 | 27 | 53.3 | 122 | 7.378e+6 |
| 4 \| 4 | >sp\|P0CE91\|PAU18_YEAST Seripauperin-18 OS=Saccharomyces cerevisiae (strain ATCC 204508 / S288c) OX=5... | 62.54 | 11.00 | 666.6 | 13 | 9 | 9 | 9.2 | 120 | 3.115e+5 |

Spectrum (double click to dock / undock)

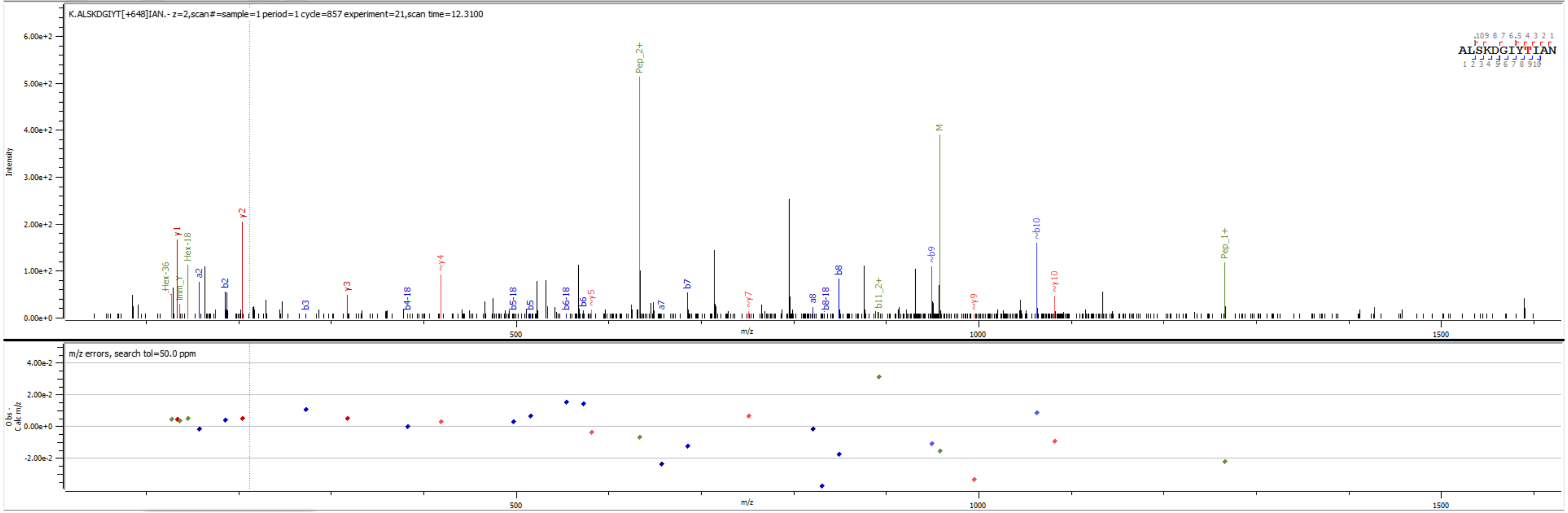


Sample S4 K.ALSKDGIYT[+648.211]IAN.-

**Peptides**

| PID | Prot. Rank | Pos. | Sequence | Mods (variable) | Glycans | Score | elta Mo Score | z | Obs. m/z | Calc. m/z | ppm err. | Off-By-X | Obs. MH | Calc. MH | Cleavage | Glycans Pos. | Protein Name | Prot. Id | Scan Time | |
|---|---|---|---|---|---|---|---|---|---|---|---|---|---|---|---|---|---|---|---|---|
| 1 \| 381... | 2 | 113 | K.DGIYT[+324.10565]IAN.- | T5(OGlycan / 324.1056) | Hex(2) | 479.4 | 146.7 | 2 | 595.7590 | 595.7692 | -17.06 | | 1190.5108 | 1190.5311 | Specific | 5 | >sp\|P32612\|PAU2_YEAST Seripauperin-2 OS=Saccharomyces cerevisiae (strain ATCC 2... | 3410 | 13.2611 | sample=1 perioc |

**Proteins**

| Prot. Rank | Protein Name | \|Log Prob\| | Best \|Log Prob\| | Best Score | # Spectra | # Uniq. Peps. | # Mod Peps. | % Cov. | # AAs | Intensity |
|---|---|---|---|---|---|---|---|---|---|---|
| 1 \| 1 | >sp\|P43575\|PAU5_YEAST Seripauperin-5 OS=Saccharomyces cerevisiae (strain ATCC 204508 / S288c) OX=559... | 327.47 | 16.74 | 1004.1 | 632 | 70 | 67 | 68.9 | 122 | 3.742e+7 |
| 2 \| 2 | >sp\|P32612\|PAU2_YEAST Seripauperin-2 OS=Saccharomyces cerevisiae (strain ATCC 204508 / S288c) OX=559... | 178.33 | 13.24 | 726.5 | 270 | 60 | 58 | 58.3 | 120 | 7.688e+6 |

Spectrum (double click to dock / undock)

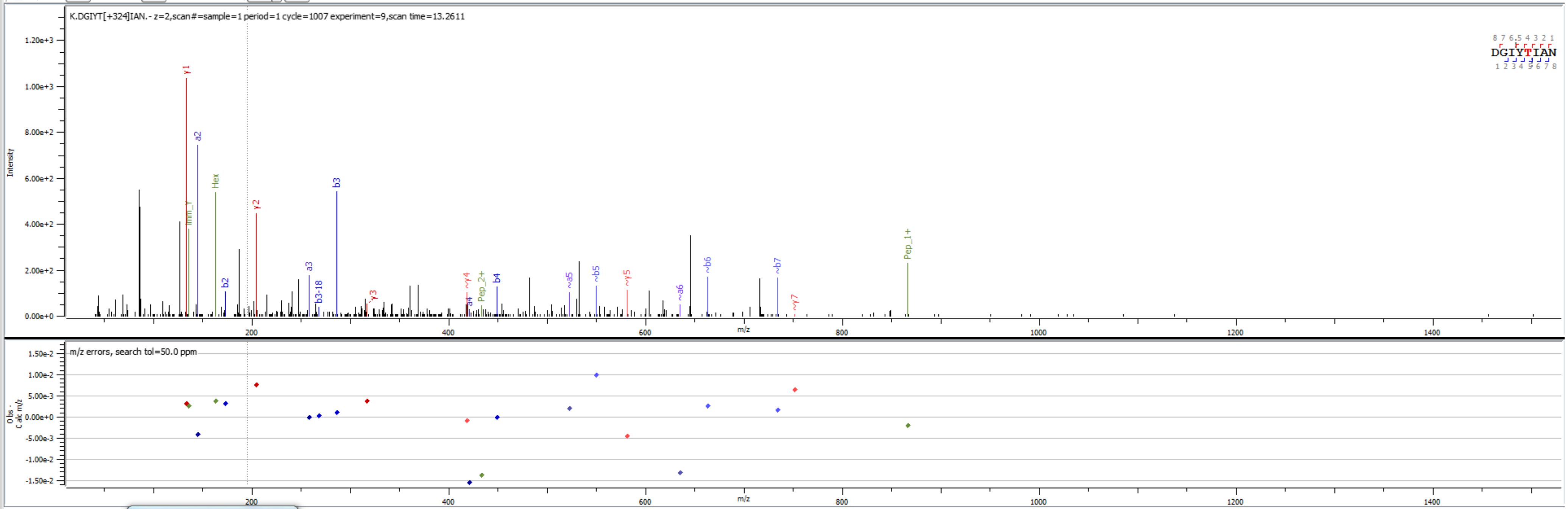


Sample S1 K.DGIYT[+324.106]IAN.-

**Peptides**

| PID | Prot. Rank | Pos. | Sequence | Mods (variable) | Glycans | Score | elta Mo Score | z | Obs. m/z | Calc. m/z | ppm err. | Off-By-X | Obs. MH | Calc. MH | Cleavage | Glycans Pos. | Protein Name | Prot. Id | Scan Time | |
|---|---|---|---|---|---|---|---|---|---|---|---|---|---|---|---|---|---|---|---|---|
| 1 \| 368... | 11 | 24 | K.DPWST[+648.21129]LTPS.A | T5(OGlycan / 648.2113) | Hex(4) | 395.5 | 0.0 | 2 | 826.3449 | 826.3458 | -1.17 | | 1651.6824 | 1651.6844 | CRagged | 5 | >sp\|Q03180\|PIR3_YEAST Cell wall mannoprotein PIR3 OS=Saccharomyces cerevisiae (s... | 3365 | 13.6993 | sample=1 perioc |

**Proteins**

| Prot. Rank | Protein Name | \|Log Prob\| | Best \|Log Prob\| | Best Score | # Spectra | # Uniq. Peps. | # Mod Peps. | % Cov. | # AAs | Intensity |
|---|---|---|---|---|---|---|---|---|---|---|
| 10 \| 10 | >sp\|Q03674\|PLB2_YEAST Lysophospholipase 2 OS=Saccharomyces cerevisiae (strain ATCC 204508 / S288c) O... | 11.70 | 6.05 | 597.1 | 5 | 2 | 0 | 3.5 | 706 | 1.207e+4 |
| 11 \| 11 | >sp\|Q03180\|PIR3_YEAST Cell wall mannoprotein PIR3 OS=Saccharomyces cerevisiae (strain ATCC 204508 / S2... | 10.18 | 5.74 | 637.9 | 17 | 11 | 11 | 5.8 | 325 | 5.629e+4 |
| 12 \| 12 | >sp\|P17260\|KRE1_YEAST Protein KRE1 OS=Saccharomyces cerevisiae (strain ATCC 204508 / S288c) OX=55929... | 9.22 | 9.16 | 776.9 | 3 | 2 | 2 | 7.7 | 313 | 1.120e+4 |
| 13 \| 13 | >sp\|Q05164\|HPF1_YEAST Haze protective factor 1 OS=Saccharomyces cerevisiae (strain ATCC 204508 / S288c)... | 8.27 | 8.22 | 749.3 | 4 | 1 | 0 | 1.1 | 967 | 1.081e+4 |

Spectrum (double click to dock / undock)

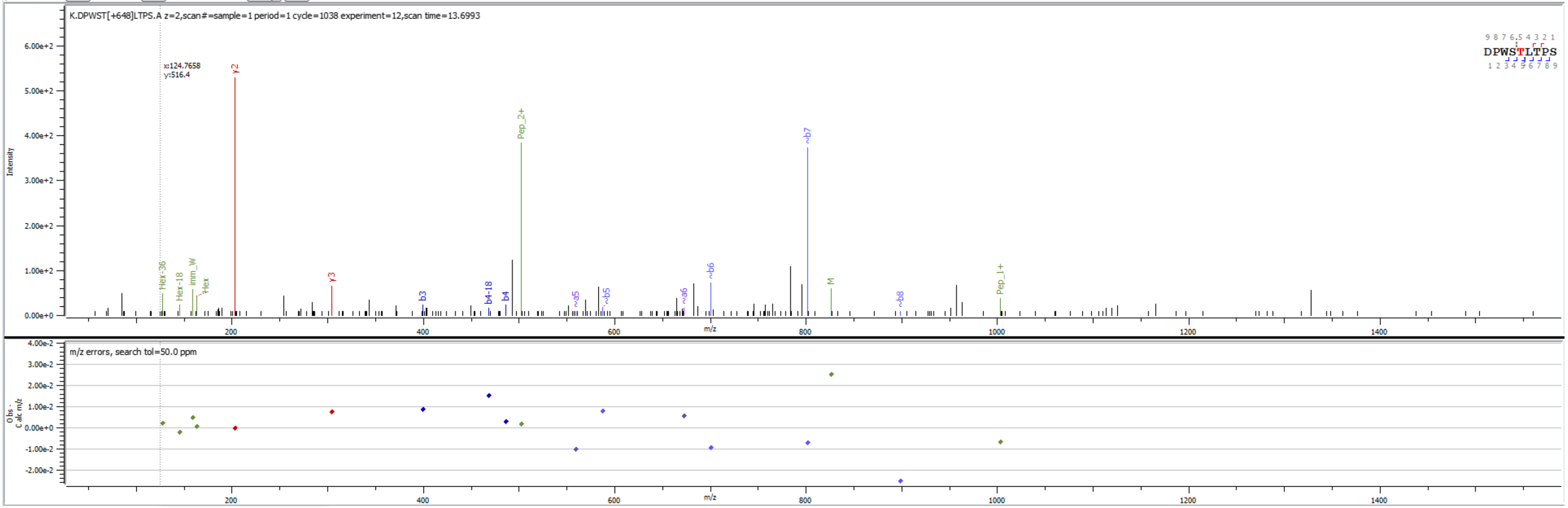


Sample S2 K.DPWST[+648.211]LTPS.A

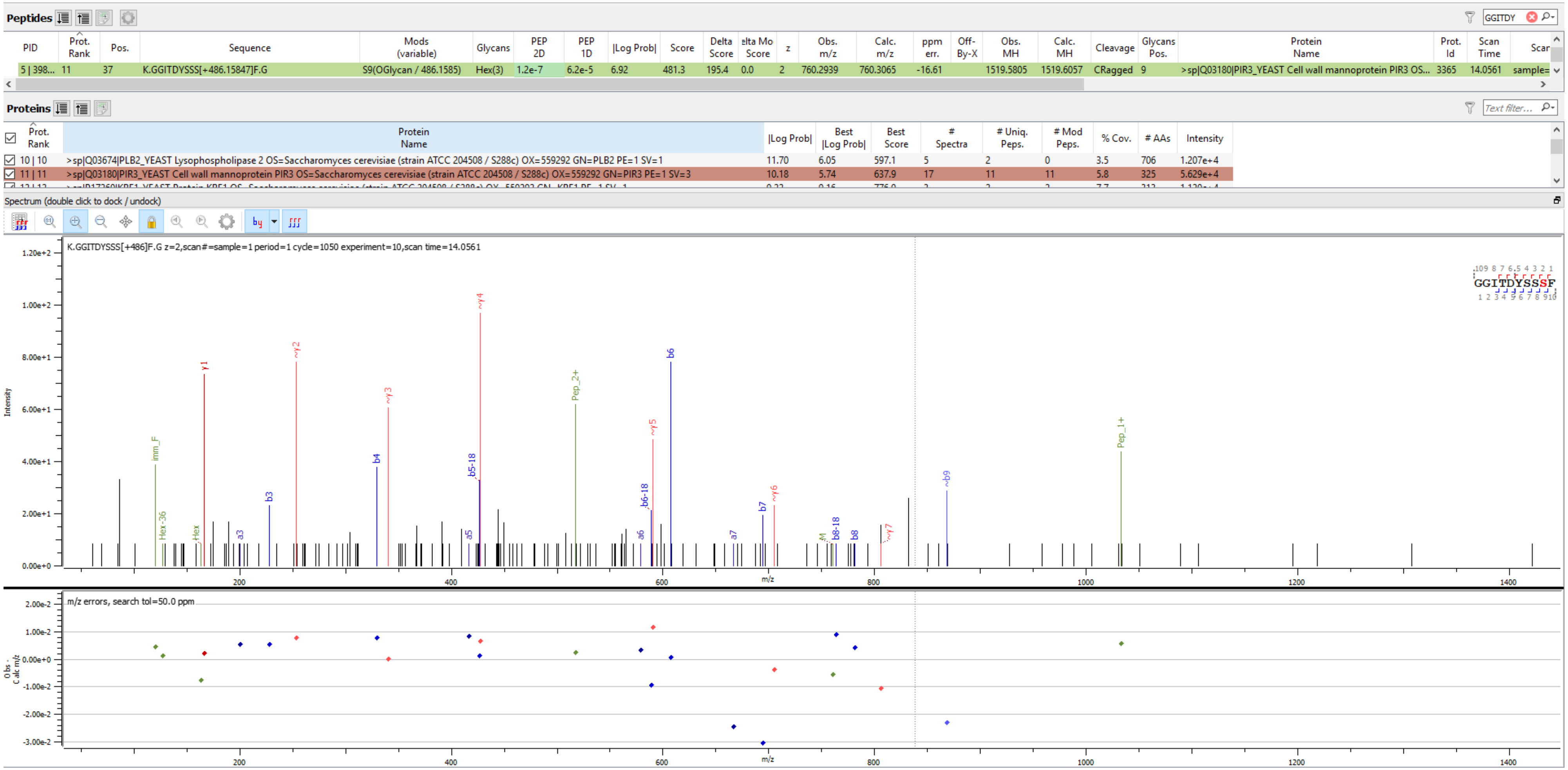


Sample S2 K.GGITDYSSS[+486.158]F.G

**Peptides**

| PID | Prot. Rank | Pos. | Sequence | Mods (variable) | Glycans | Score | elta Mo Score | z | Obs. m/z | Calc. m/z | ppm err. | Off-By-X | Obs. MH | Calc. MH | Cleavage | Glycans Pos. | Protein Name | Prot. Id | Scan Time | |
|---|---|---|---|---|---|---|---|---|---|---|---|---|---|---|---|---|---|---|---|---|
| 1 \| 342... | 11 | 37 | K.GGITDYS[+648.21129]SSF.G | S7(OGlycan / 648.2113) | Hex(4) | 637.9 | 0.0 | 2 | 841.3211 | 841.3329 | -14.04 | | 1681.6350 | 1681.6586 | CRagged | 7 | >sp\|Q03180\|PIR3_YEAST Cell wall mannoprotein PIR3 OS=Saccharomyces cerevisiae (s... | 3365 | 13.4938 | sample=1 perioc |

**Proteins**

| | Prot. Rank | Protein Name | \|Log Prob\| | Best \|Log Prob\| | Best Score | # Spectra | # Uniq. Peps. | # Mod Peps. | % Cov. | # AAs | Intensity |
|---|---|---|---|---|---|---|---|---|---|---|---|
| ☑ | 10 \| 10 | >sp\|Q03674\|PLB2_YEAST Lysophospholipase 2 OS=Saccharomyces cerevisiae (strain ATCC 204508 / S288c) O... | 11.70 | 6.05 | 597.1 | 5 | 2 | 0 | 3.5 | 706 | 1.207e+4 |
| ☑ | 11 \| 11 | >sp\|Q03180\|PIR3_YEAST Cell wall mannoprotein PIR3 OS=Saccharomyces cerevisiae (strain ATCC 204508 / S2... | 10.18 | 5.74 | 637.9 | 17 | 11 | 11 | 5.8 | 325 | 5.629e+4 |
| ☑ | 12 \| 12 | >sp\|P17260\|KRE1_YEAST Protein KRE1 OS=Saccharomyces cerevisiae (strain ATCC 204508 / S288c) OX=55929... | 9.22 | 9.16 | 776.9 | 3 | 2 | 2 | 7.7 | 313 | 1.120e+4 |
| ☑ | 13 \| 13 | >sp\|Q05164\|HPF1_YEAST Haze protective factor 1 OS=Saccharomyces cerevisiae (strain ATCC 204508 / S288c)... | 8.27 | 8.22 | 749.3 | 4 | 1 | 0 | 1.1 | 967 | 1.081e+4 |

Spectrum (double click to dock / undock)

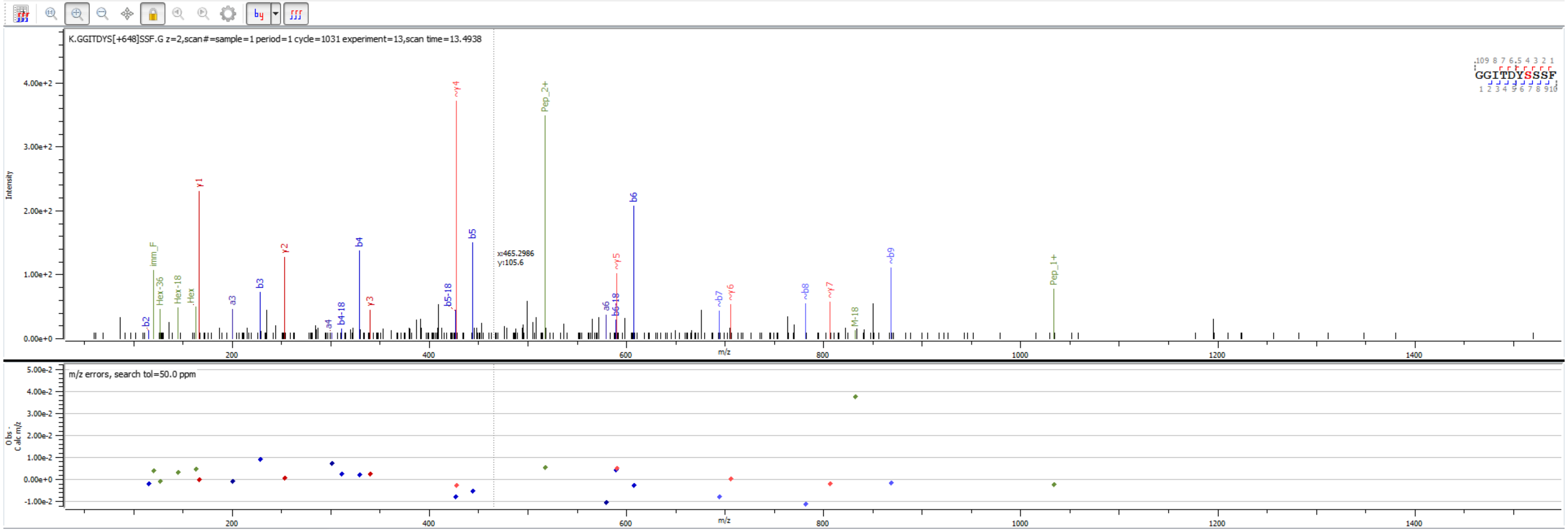


Sample S2 K.GGITDYS[+648.211]SSF.G

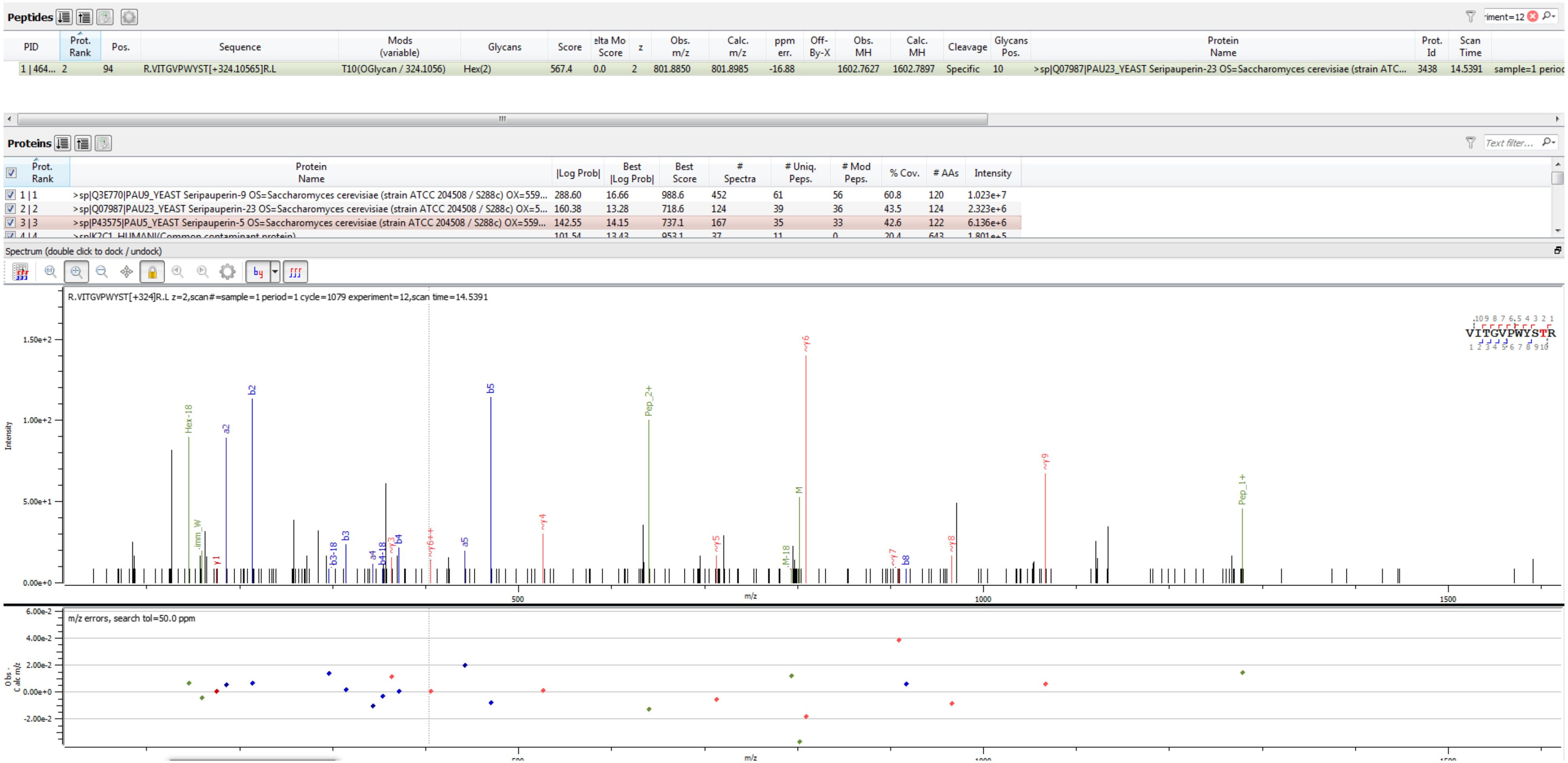


Sample S6 R.VITGVPWYST[+324.106]R.L

**Peptides**

| PID | Prot. Rank | Pos. | Sequence | Mods (variable) | Glycans | Score | elta Mo Score | z | Obs. m/z | Calc. m/z | ppm err. | Off-By-X | Obs. MH | Calc. MH | Cleavage | Glycans Pos. | Protein Name | Prot. Id | Scan Time | |
|---|---|---|---|---|---|---|---|---|---|---|---|---|---|---|---|---|---|---|---|---|
| 1 \| 121... | 2 | 94 | R.VITGVPWYS[+486.15847]TR.L | S9(OGlycan / 486.1585) | Hex(3) | 485.5 | 0.0 | 2 | 882.9132 | 882.9249 | -13.24 | | 1764.8192 | 1764.8426 | Specific | 9 | >sp\|P0CE85\|PAU19_YEAST Seripauperin-19 OS=Saccharomyces cerevisiae (strain ATC... | 3420 | 14.9705 | sample=1 perioc |

**Proteins**

| Prot. Rank | Protein Name | \|Log Prob\| | Best \|Log Prob\| | Best Score | # Spectra | # Uniq. Peps. | # Mod Peps. | % Cov. | # AAs | Intensity |
|---|---|---|---|---|---|---|---|---|---|---|
| 1 \| 1 | >sp\|P32612\|PAU2_YEAST Seripauperin-2 OS=Saccharomyces cerevisiae (strain ATCC 204508 / S288c) OX=559... | 304.25 | 17.48 | 982.9 | 381 | 64 | 61 | 67.5 | 120 | 1.839e+7 |
| 2 \| 2 | >sp\|P0CE85\|PAU19_YEAST Seripauperin-19 OS=Saccharomyces cerevisiae (strain ATCC 204508 / S288c) OX=5... | 138.61 | 11.51 | 683.8 | 93 | 32 | 31 | 42.7 | 124 | 3.148e+6 |
| 3 \| 3 | >sp\|P43575\|PAU5_YEAST Seripauperin-5 OS=Saccharomyces cerevisiae (strain ATCC 204508 / S288c) OX=559... | 129.60 | 12.22 | 721.6 | 114 | 28 | 27 | 53.3 | 122 | 7.378e+6 |
| 4 \| 4 | >sp\|P0CE91\|PAU18_YEAST Seripauperin-18 OS=Saccharomyces cerevisiae (strain ATCC 204508 / S288c) OX=5... | 62.54 | 11.00 | 666.6 | 13 | 9 | 9 | 9.2 | 120 | 3.115e+5 |

Spectrum (double click to dock / undock)

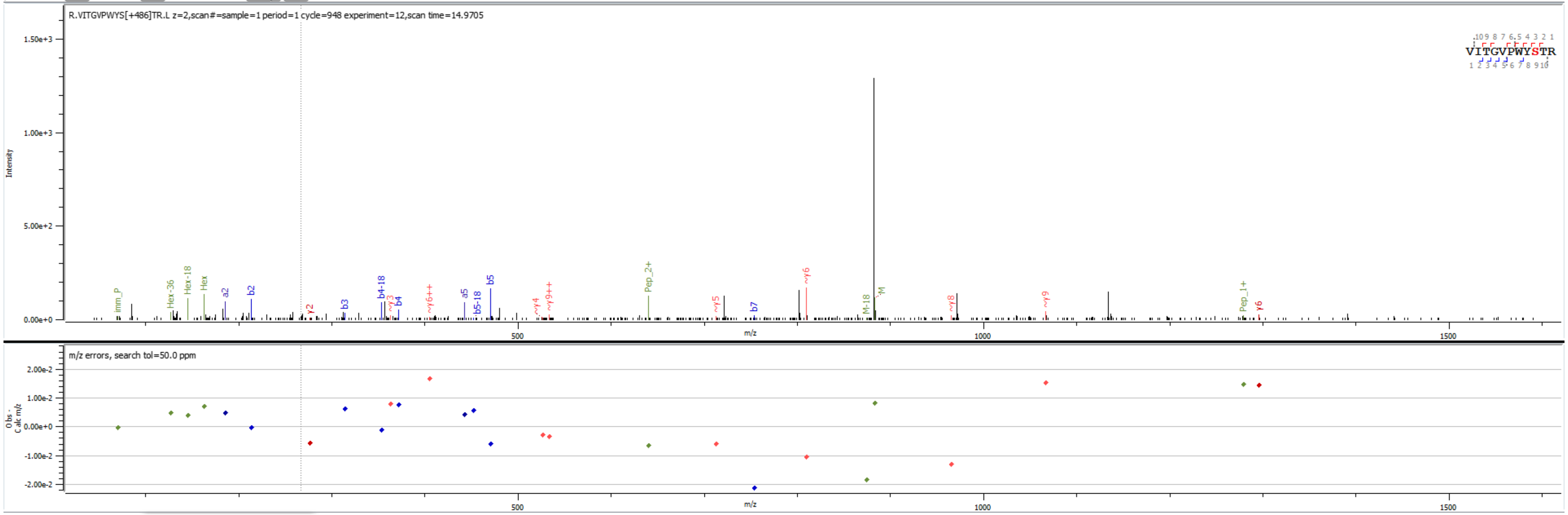


Sample S4 R.VITGVPWYS[+486.158]TR.L

**Peptides**

| PID | Prot. Rank | Pos. | Sequence | Mods (variable) | Glycans | Score | elta Mo Score | z | Obs. m/z | Calc. m/z | ppm err. | Off-By-X | Obs. MH | Calc. MH | Cleavage | Glycans Pos. | Protein Name | Prot. Id | Scan Time | |
|---|---|---|---|---|---|---|---|---|---|---|---|---|---|---|---|---|---|---|---|---|
| 1 \| 119... | 2 | 94 | R.VITGVPWYS[+486.15847]TR.L | S9(OGlycan / 486.1585) | Hex(3) | 468.2 | 0.0 | 3 | 588.9442 | 588.9524 | -13.92 | | 1764.8180 | 1764.8426 | Specific | 9 | >sp\|P0CE85\|PAU19_YEAST Seripauperin-19 OS=Saccharomyces cerevisiae (strain ATC... | 3420 | 14.6791 | sample=1 perioc |

**Proteins**

| Prot. Rank | Protein Name | \|Log Prob\| | Best \|Log Prob\| | Best Score | # Spectra | # Uniq. Peps. | # Mod Peps. | % Cov. | # AAs | Intensity |
|---|---|---|---|---|---|---|---|---|---|---|
| 1 \| 1 | >sp\|P32612\|PAU2_YEAST Seripauperin-2 OS=Saccharomyces cerevisiae (strain ATCC 204508 / S288c) OX=559... | 304.25 | 17.48 | 982.9 | 381 | 64 | 61 | 67.5 | 120 | 1.839e+7 |
| 2 \| 2 | >sp\|P0CE85\|PAU19_YEAST Seripauperin-19 OS=Saccharomyces cerevisiae (strain ATCC 204508 / S288c) OX=5... | 138.61 | 11.51 | 683.8 | 93 | 32 | 31 | 42.7 | 124 | 3.148e+6 |
| 3 \| 3 | >sp\|P43575\|PAU5_YEAST Seripauperin-5 OS=Saccharomyces cerevisiae (strain ATCC 204508 / S288c) OX=559... | 129.60 | 12.22 | 721.6 | 114 | 28 | 27 | 53.3 | 122 | 7.378e+6 |
| 4 \| 4 | >sp\|P0CE91\|PAU18_YEAST Seripauperin-18 OS=Saccharomyces cerevisiae (strain ATCC 204508 / S288c) OX=5... | 62.54 | 11.00 | 666.6 | 13 | 9 | 9 | 9.2 | 120 | 3.115e+5 |

Spectrum (double click to dock / undock)

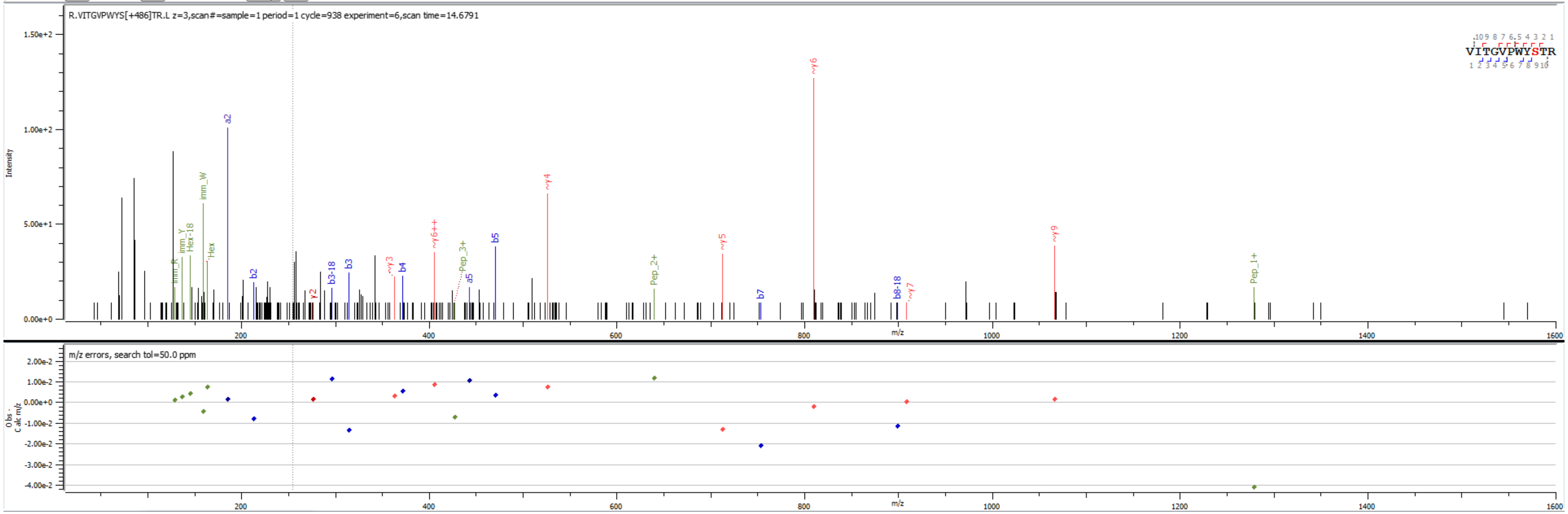


Sample S4 R.VITGVPWYS[+486.158]TR.L

**Peptides**

| PID | Prot. Rank | Pos. | Sequence | Mods (variable) | Glycans | Score | elta Mo Score | z | Obs. m/z | Calc. m/z | ppm err. | Off-By-X | Obs. MH | Calc. MH | Cleavage | Glycans Pos. | Protein Name | Prot. Id | Scan Time | |
|---|---|---|---|---|---|---|---|---|---|---|---|---|---|---|---|---|---|---|---|---|
| 1 \| 936... | 2 | 94 | R.VITGVPWYST[+648.21129]R.L | T10(OGlycan / 648.2113) | Hex(4) | 659.4 | 0.0 | 2 | 963.9353 | 963.9513 | -16.62 | | 1926.8633 | 1926.8954 | Specific | 10 | >sp\|Q07987\|PAU23_YEAST Seripauperin-23 OS=Saccharomyces cerevisiae (strain ATC... | 3438 | 14.9432 | sample=1 perioc |

**Proteins**

| Prot. Rank | Protein Name | \|Log Prob\| | Best \|Log Prob\| | Best Score | # Spectra | # Uniq. Peps. | # Mod Peps. | % Cov. | # AAs | Intensity |
|---|---|---|---|---|---|---|---|---|---|---|
| 1 \| 1 | >sp\|Q3E770\|PAU9_YEAST Seripauperin-9 OS=Saccharomyces cerevisiae (strain ATCC 204508 / S288c) OX=559... | 288.60 | 16.66 | 988.6 | 452 | 61 | 56 | 60.8 | 120 | 1.023e+7 |
| 2 \| 2 | >sp\|Q07987\|PAU23_YEAST Seripauperin-23 OS=Saccharomyces cerevisiae (strain ATCC 204508 / S288c) OX=5... | 160.38 | 13.28 | 718.6 | 124 | 39 | 36 | 43.5 | 124 | 2.323e+6 |
| 3 \| 3 | >sp\|P43575\|PAU5_YEAST Seripauperin-5 OS=Saccharomyces cerevisiae (strain ATCC 204508 / S288c) OX=559... | 142.55 | 14.15 | 737.1 | 167 | 35 | 33 | 42.6 | 122 | 6.136e+6 |
| 4 \| 4 | >sp\|K2C1_HUMAN\|(Common contaminant protein) | 101.54 | 13.43 | 953.1 | 37 | 11 | 0 | 20.4 | 643 | 1.801e+5 |

Spectrum (double click to dock / undock)

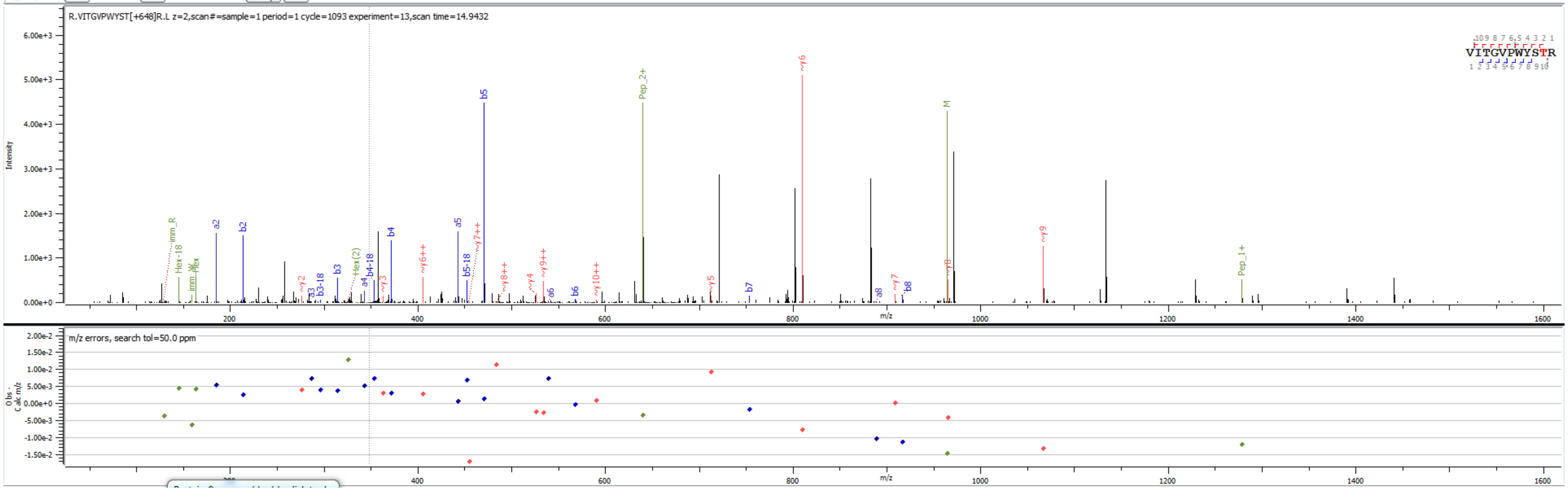


Sample S6 R.VITGVPWYST[+648.211]R.L

**Peptides**

| PID | Prot. Rank | Pos. | Sequence | Mods (variable) | Glycans | Score | elta Mo Score | z | Obs. m/z | Calc. m/z | ppm err. | Off-By-X | Obs. MH | Calc. MH | Cleavage | Glycans Pos. | Protein Name | Prot. Id | Scan Time | |
|---|---|---|---|---|---|---|---|---|---|---|---|---|---|---|---|---|---|---|---|---|
| 1 \| 115... | 3 | 94 | R.VITGVPWYST[+648.21129]R.L | T10(OGlycan / 648.2113) | Hex(4) | 594.7 | 0.0 | 3 | 642.9572 | 642.9700 | -19.83 | | 1926.8572 | 1926.8954 | Specific | 10 | >sp\|P0CE85\|PAU19_YEAST Seripauperin-19 OS=Saccharomyces cerevisiae (strain ATC... | 3420 | 14.4496 | sample=1 perioc |

**Proteins**

| Prot. Rank | Protein Name | \|Log Prob\| | Best \|Log Prob\| | Best Score | # Spectra | # Uniq. Peps. | # Mod Peps. | % Cov. | # AAs | Intensity |
|---|---|---|---|---|---|---|---|---|---|---|
| 1 \| 1 | >sp\|Q3E770\|PAU9_YEAST Seripauperin-9 OS=Saccharomyces cerevisiae (strain ATCC 204508 / S288c) OX=559... | 248.32 | 15.68 | 955.8 | 342 | 58 | 55 | 67.5 | 120 | 1.797e+7 |

Spectrum (double click to dock / undock)

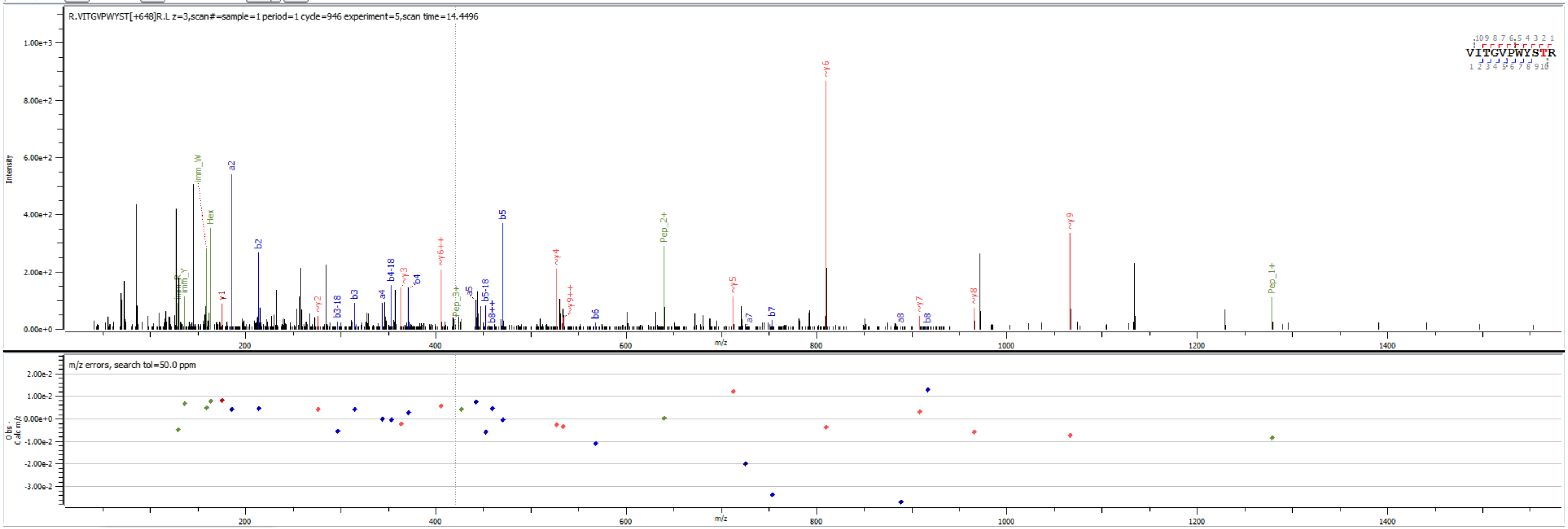


Sample S3 R.VITGVPWYST[+648.211]R.L

**Peptides**

| PID | Prot. Rank | Pos. | Sequence | Mods (variable) | Glycans | Score | elta Mo Score | z | Obs. m/z | Calc. m/z | ppm err. | Off-By-X | Obs. MH | Calc. MH | Cleavage | Glycans Pos. | Protein Name | Prot. Id | Scan Time | |
|---|---|---|---|---|---|---|---|---|---|---|---|---|---|---|---|---|---|---|---|---|
| 1 \| 108... | 4 | 94 | R.VITGVPWYS[+810.26412]TR.L | S9(OGlycan / 810.2641) | Hex(5) | 647.2 | 1.2 | 2 | 1044.9594 | 1044.9777 | -17.60 | | 2088.9114 | 2088.9482 | Specific | 9 | >sp\|Q07987\|PAU23_YEAST Seripauperin-23 OS=Saccharomyces cerevisiae (strain ATC... | 3438 | 14.6709 | sample=1 perioc |

**Proteins**

| Prot. Rank | Protein Name | \|Log Prob\| | Best \|Log Prob\| | Best Score | # Spectra | # Uniq. Peps. | # Mod Peps. | % Cov. | # AAs | Intensity |
|---|---|---|---|---|---|---|---|---|---|---|
| 1 \| 1 | >sp\|P43575\|PAU5_YEAST Seripauperin-5 OS=Saccharomyces cerevisiae (strain ATCC 204508 / S288c) OX=559... | 327.47 | 16.74 | 1004.1 | 632 | 70 | 67 | 68.9 | 122 | 3.742e+7 |
| 2 \| 2 | >sp\|P32612\|PAU2_YEAST Seripauperin-2 OS=Saccharomyces cerevisiae (strain ATCC 204508 / S288c) OX=559... | 178.33 | 13.24 | 726.5 | 270 | 60 | 58 | 58.3 | 120 | 7.688e+6 |

Spectrum (double click to dock / undock)

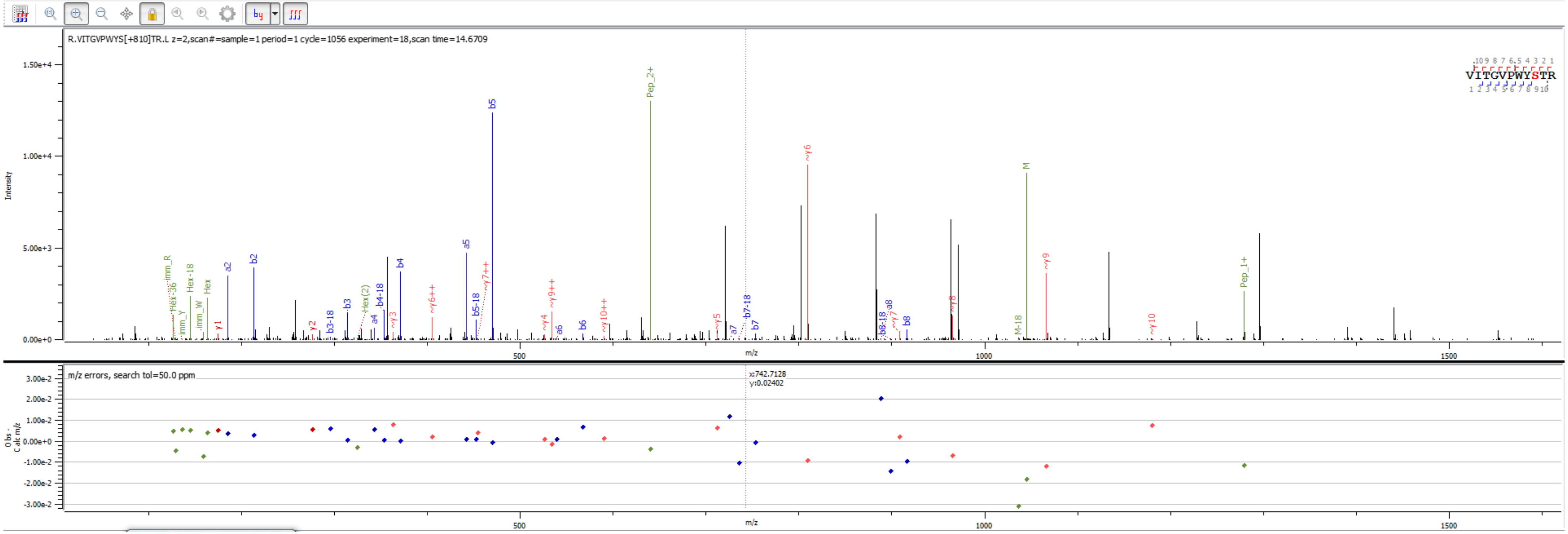


Sample S1 R.VITGVPWYS[+810.264]TR.L

**Peptides**

| PID | Prot. Rank | Pos. | Sequence | Mods (variable) | Glycans | Score | elta Mo Score | z | Obs. m/z | Calc. m/z | ppm err. | Off-By-X | Obs. MH | Calc. MH | Cleavage | Glycans Pos. | Protein Name | Prot. Id | Scan Time | |
|---|---|---|---|---|---|---|---|---|---|---|---|---|---|---|---|---|---|---|---|---|
| 1 \| 927... | 2 | 94 | R.VITGVPWYS[+810.26412]TR.L | S9(OGlycan / 810.2641) | Hex(5) | 711.5 | 1.2 | 3 | 696.9746 | 696.9876 | -18.69 | | 2088.9092 | 2088.9482 | Specific | 9 | >sp\|Q07987\|PAU23_YEAST Seripauperin-23 OS=Saccharomyces cerevisiae (strain ATC... | 3438 | 14.8259 | sample=1 perioc |

**Proteins**

| Prot. Rank | Protein Name | \|Log Prob\| | Best \|Log Prob\| | Best Score | # Spectra | # Uniq. Peps. | # Mod Peps. | % Cov. | # AAs | Intensity |
|---|---|---|---|---|---|---|---|---|---|---|
| 1 \| 1 | >sp\|Q3E770\|PAU9_YEAST Seripauperin-9 OS=Saccharomyces cerevisiae (strain ATCC 204508 / S288c) OX=559... | 288.60 | 16.66 | 988.6 | 452 | 61 | 56 | 60.8 | 120 | 1.023e+7 |
| 2 \| 2 | >sp\|Q07987\|PAU23_YEAST Seripauperin-23 OS=Saccharomyces cerevisiae (strain ATCC 204508 / S288c) OX=5... | 160.38 | 13.28 | 718.6 | 124 | 39 | 36 | 43.5 | 124 | 2.323e+6 |
| 3 \| 3 | >sp\|P43575\|PAU5_YEAST Seripauperin-5 OS=Saccharomyces cerevisiae (strain ATCC 204508 / S288c) OX=559... | 142.55 | 14.15 | 737.1 | 167 | 35 | 33 | 42.6 | 122 | 6.136e+6 |
| 4 \| 4 | >sp\|K2C1_HUMAN\|(Common contaminant protein) | 101.54 | 13.43 | 953.1 | 37 | 11 | 0 | 20.4 | 643 | 1.801e+5 |

Spectrum (double click to dock / undock)

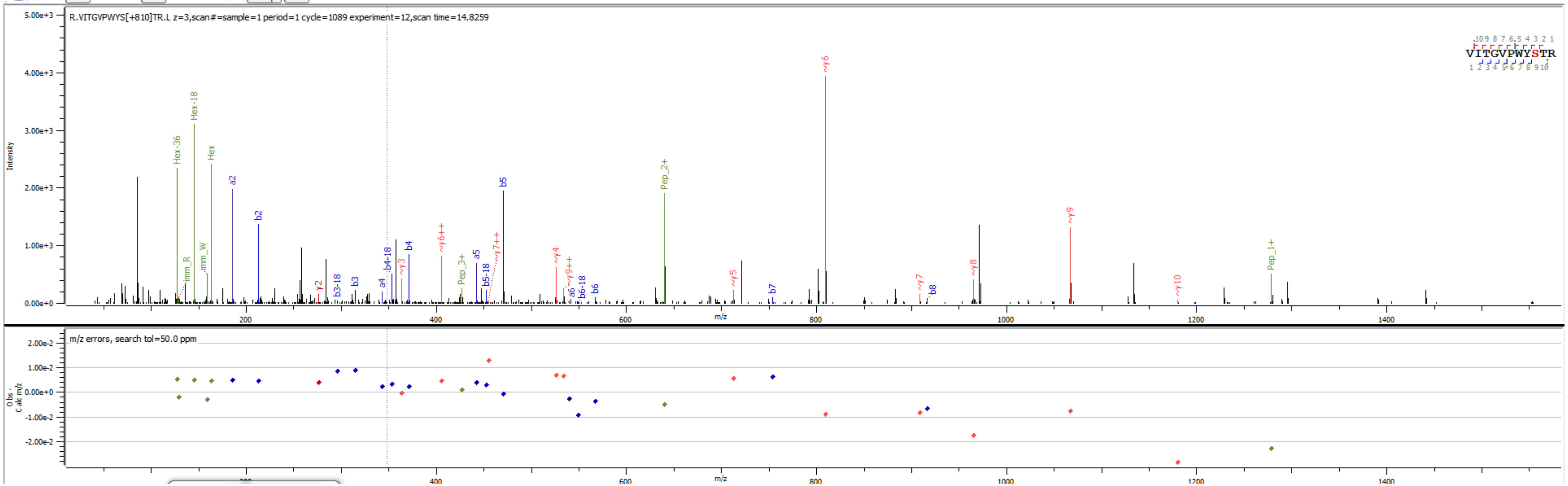


Sample S6 R.VITGVPWYS[+810.264]TR.L

**Peptides**

| PID | Prot. Rank | Pos. | Sequence | Mods (variable) | Glycans | Score | elta Mo Score | z | Obs. m/z | Calc. m/z | ppm err. | Off-By-X | Obs. MH | Calc. MH | Cleavage | Glycans Pos. | Protein Name | Prot. Id | Scan Time | |
|---|---|---|---|---|---|---|---|---|---|---|---|---|---|---|---|---|---|---|---|---|
| 1 \| 187... | 2 | 94 | R.VITGVPWYS[+972.31694]TR.L | S9(OGlycan / 972.3169) | Hex(6) | 622.4 | 1.2 | 2 | 1125.9857 | 1126.0041 | -16.40 | | 2250.9641 | 2251.0010 | Specific | 9 | >sp\|P0CE85\|PAU19_YEAST Seripauperin-19 OS=Saccharomyces cerevisiae (strain ATC... | 3420 | 14.3431 | sample=1 perioc |

**Proteins**

| Prot. Rank | Protein Name | \|Log Prob\| | Best \|Log Prob\| | Best Score | # Spectra | # Uniq. Peps. | # Mod Peps. | % Cov. | # AAs | Intensity |
|---|---|---|---|---|---|---|---|---|---|---|
| 1 \| 1 | >sp\|P32612\|PAU2_YEAST Seripauperin-2 OS=Saccharomyces cerevisiae (strain ATCC 204508 / S288c) OX=559... | 304.25 | 17.48 | 982.9 | 381 | 64 | 61 | 67.5 | 120 | 1.839e+7 |
| 2 \| 2 | >sp\|P0CE85\|PAU19_YEAST Seripauperin-19 OS=Saccharomyces cerevisiae (strain ATCC 204508 / S288c) OX=5... | 138.61 | 11.51 | 683.8 | 93 | 32 | 31 | 42.7 | 124 | 3.148e+6 |
| 3 \| 3 | >sp\|P43575\|PAU5_YEAST Seripauperin-5 OS=Saccharomyces cerevisiae (strain ATCC 204508 / S288c) OX=559... | 129.60 | 12.22 | 721.6 | 114 | 28 | 27 | 53.3 | 122 | 7.378e+6 |
| 4 \| 4 | >sp\|P0CE91\|PAU18_YEAST Seripauperin-18 OS=Saccharomyces cerevisiae (strain ATCC 204508 / S288c) OX=5... | 62.54 | 11.00 | 666.6 | 13 | 9 | 9 | 9.2 | 120 | 3.115e+5 |

Spectrum (double click to dock / undock)

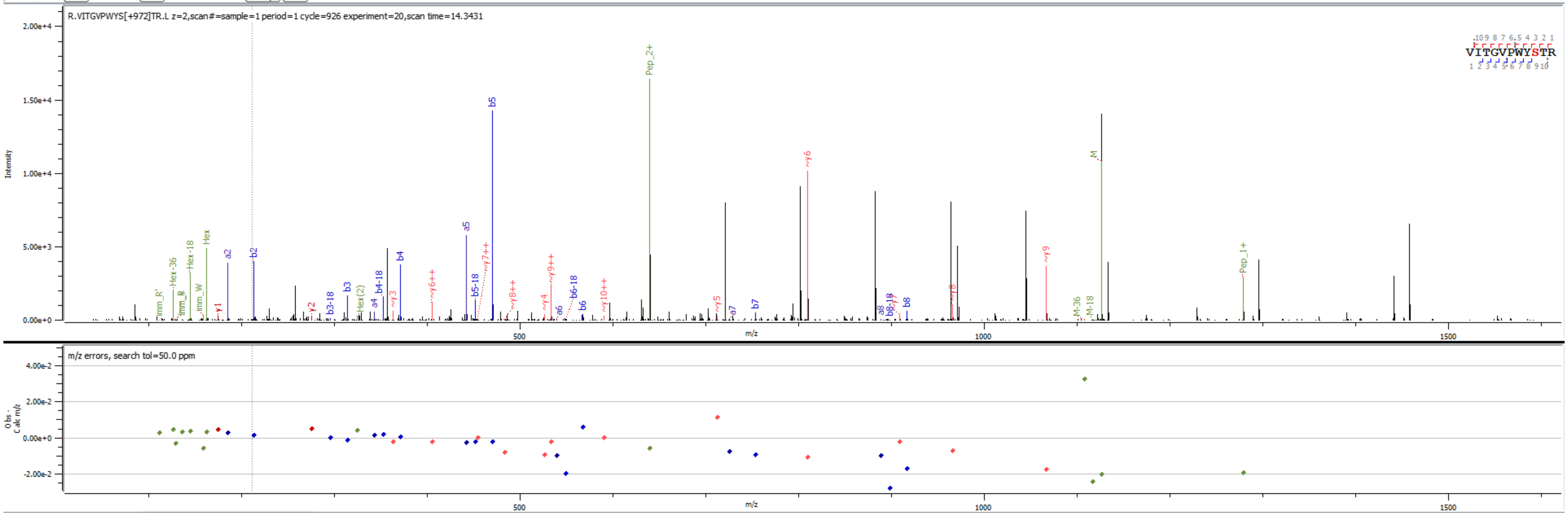


Sample S4 R.VITGVPWYS[+972.317]TR.L

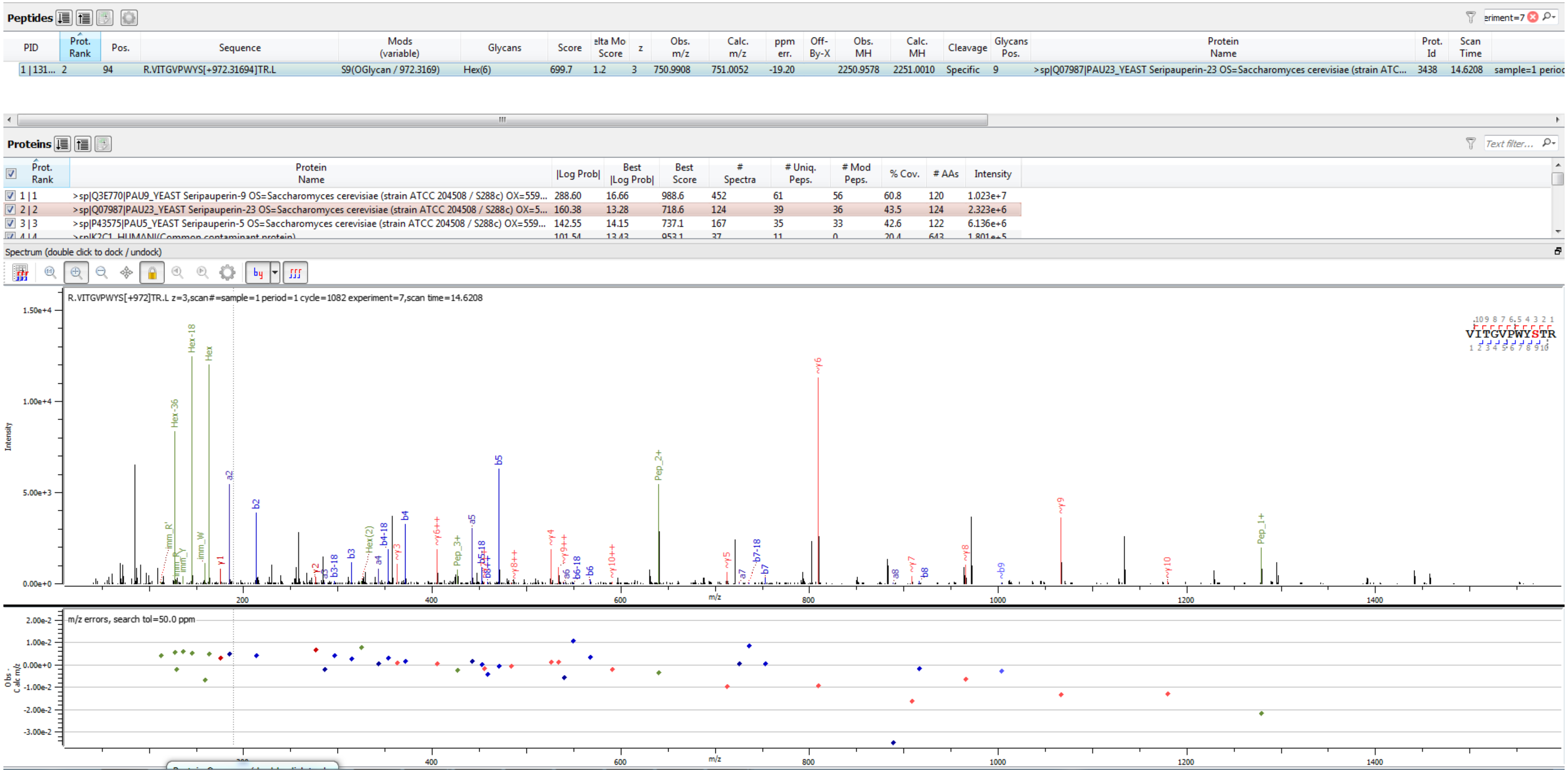


Sample S6 R.VITGVPWYS[+972.317]TR.L

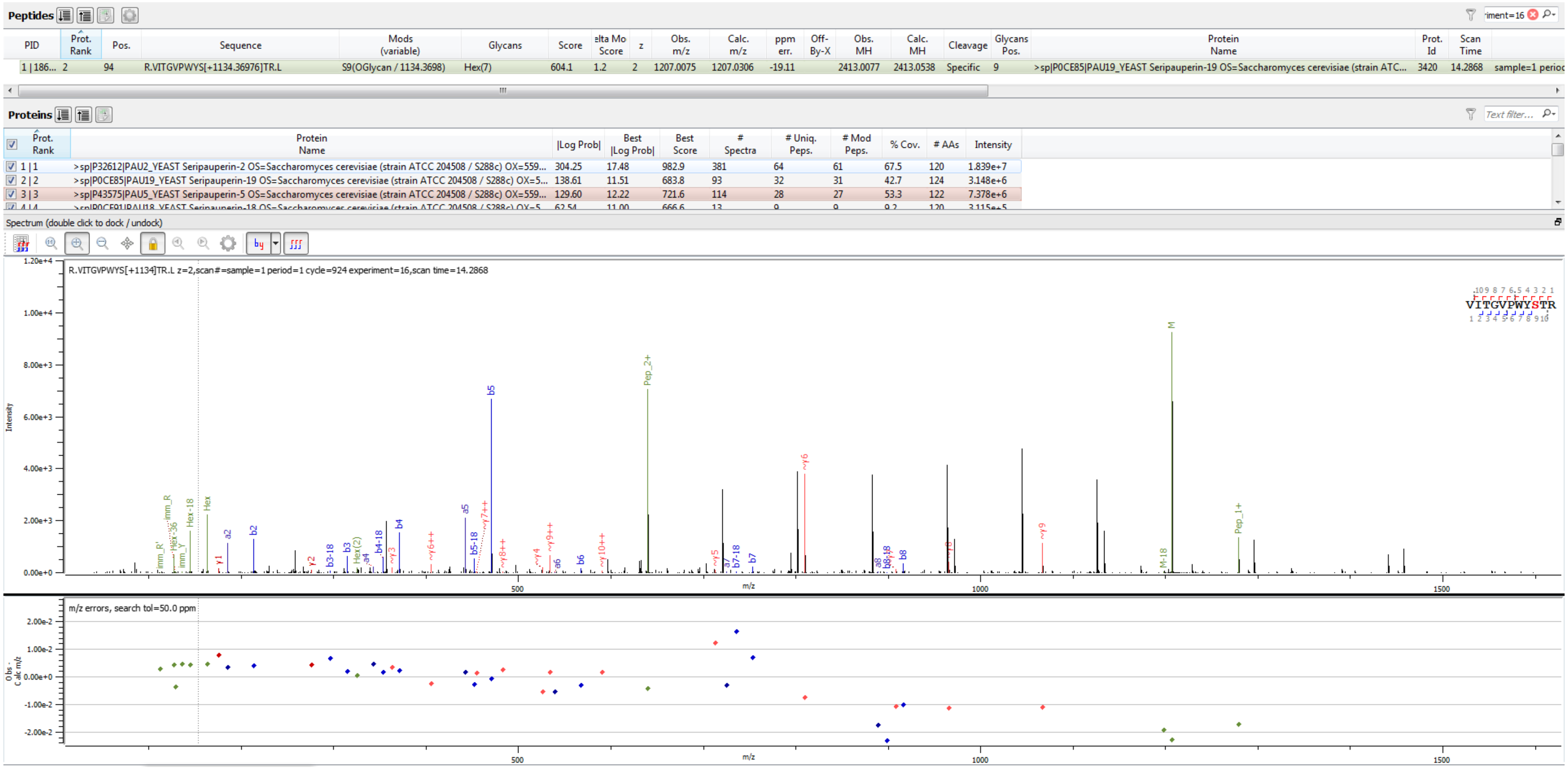


Sample S4 R.VITGVPWYS[+1134.370]TR.L

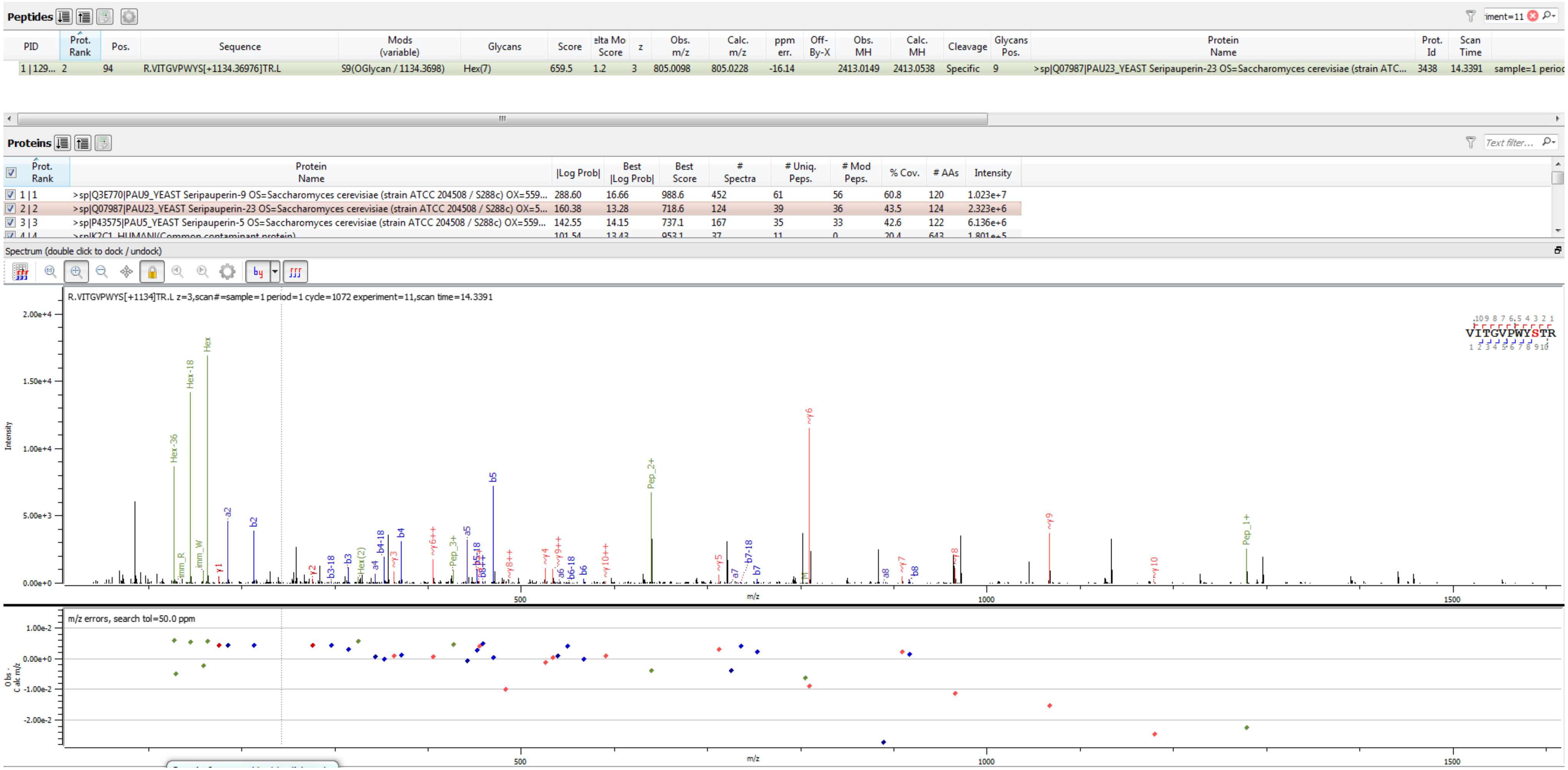


Sample S6 R.VITGVPWYS[+1134.370]TR.L

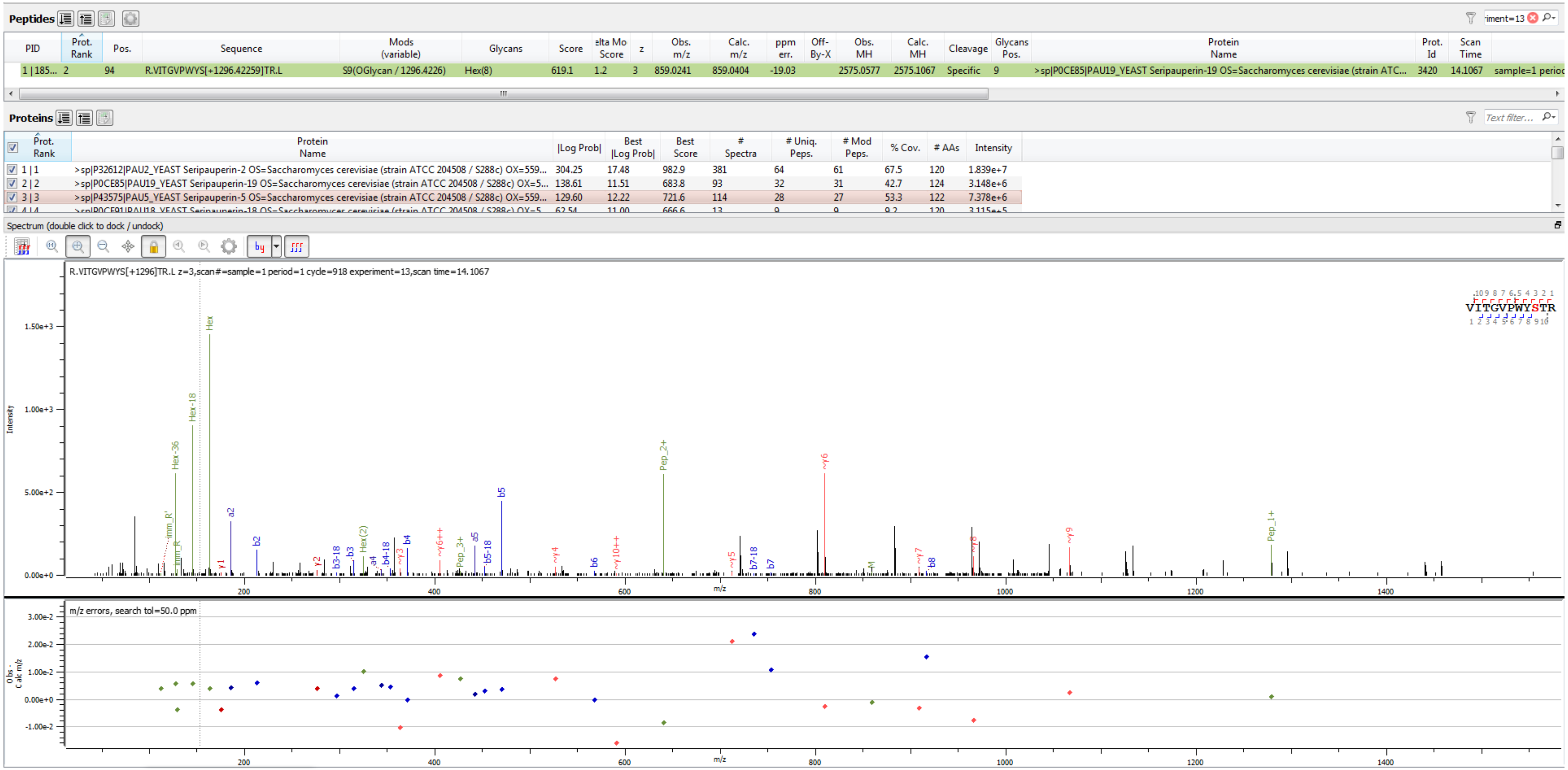


Sample S4 R.VITGVPWYS[+1296.423]TR.L

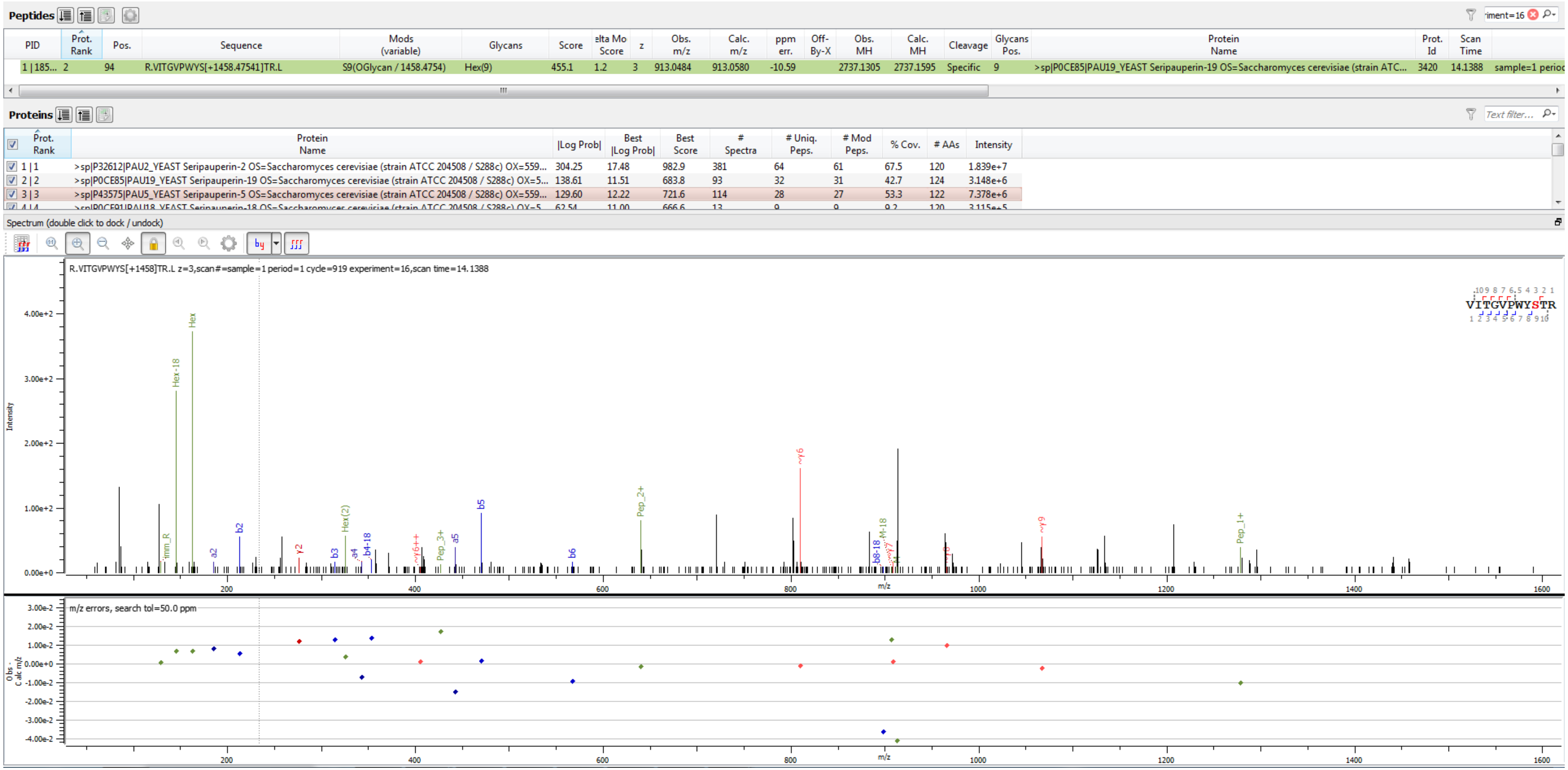


Sample S4 R.VITGVPWYS[+1458.475]TR.L

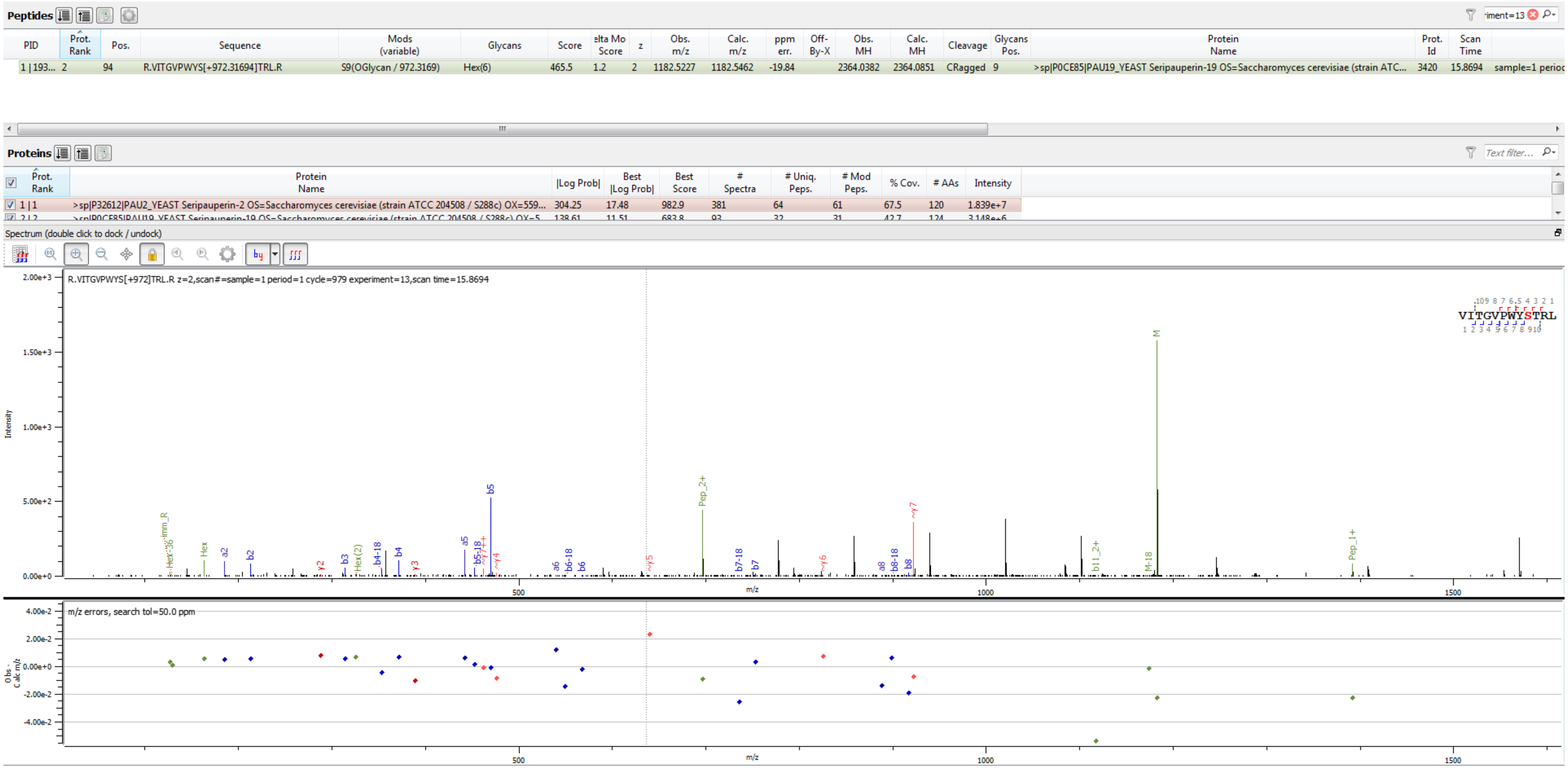


Sample S    R.VITGVPWYS[+972.317]TRL.R

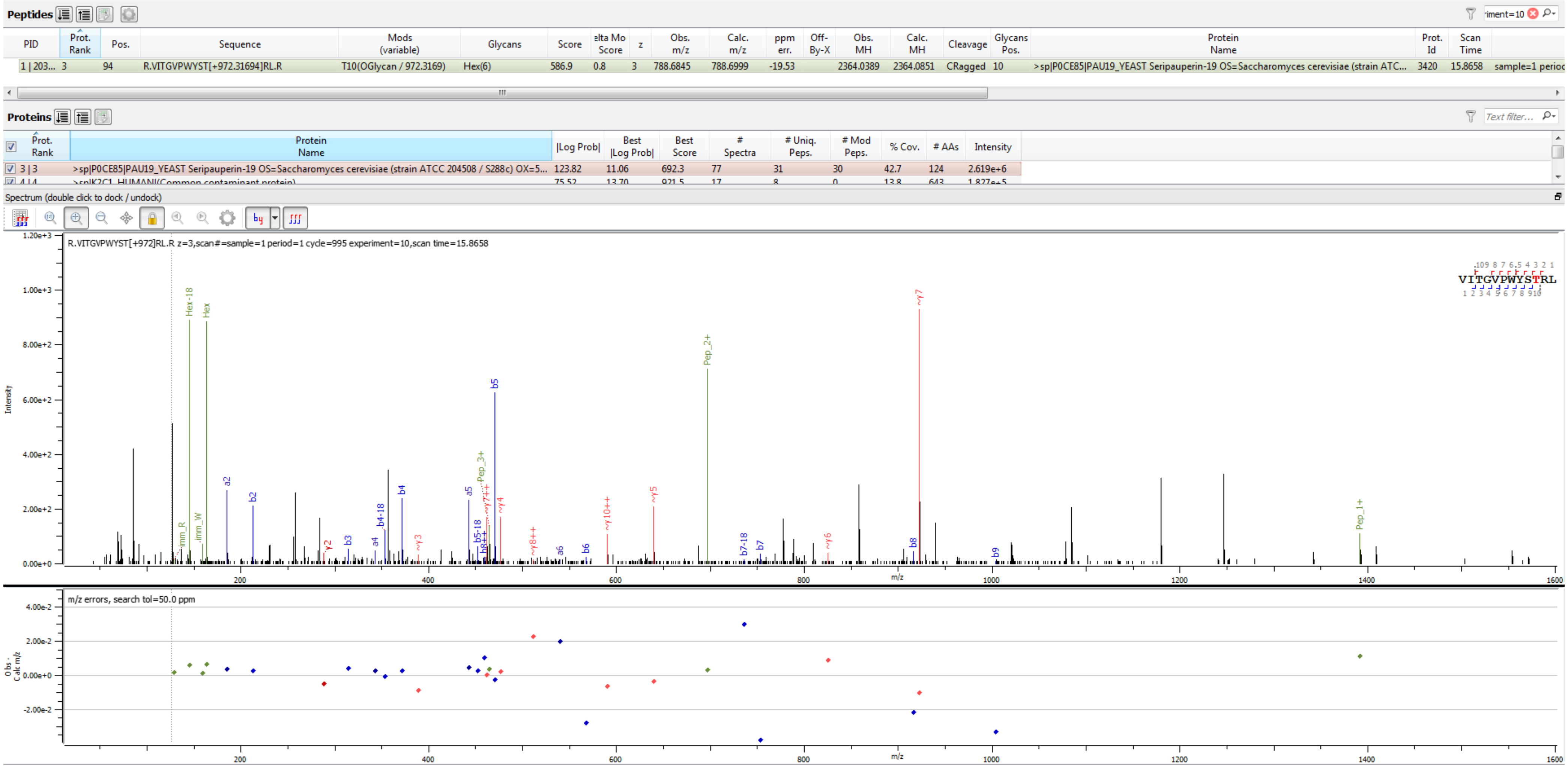


Sample S3 R.VITGVPWYST[+972.317]RL.R

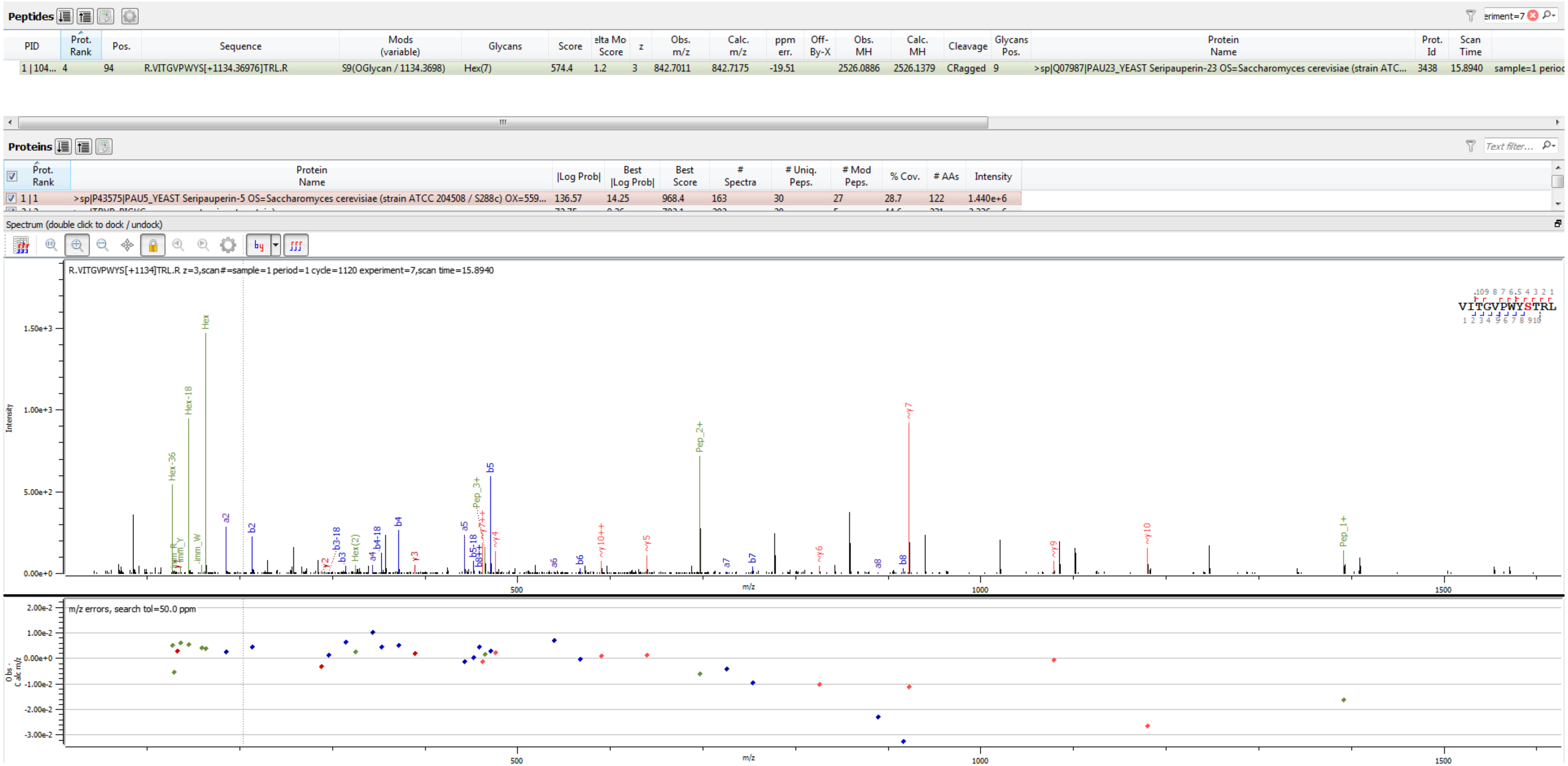


Sample S5 R.VITGVPWYS[+1134.370]TRL.R

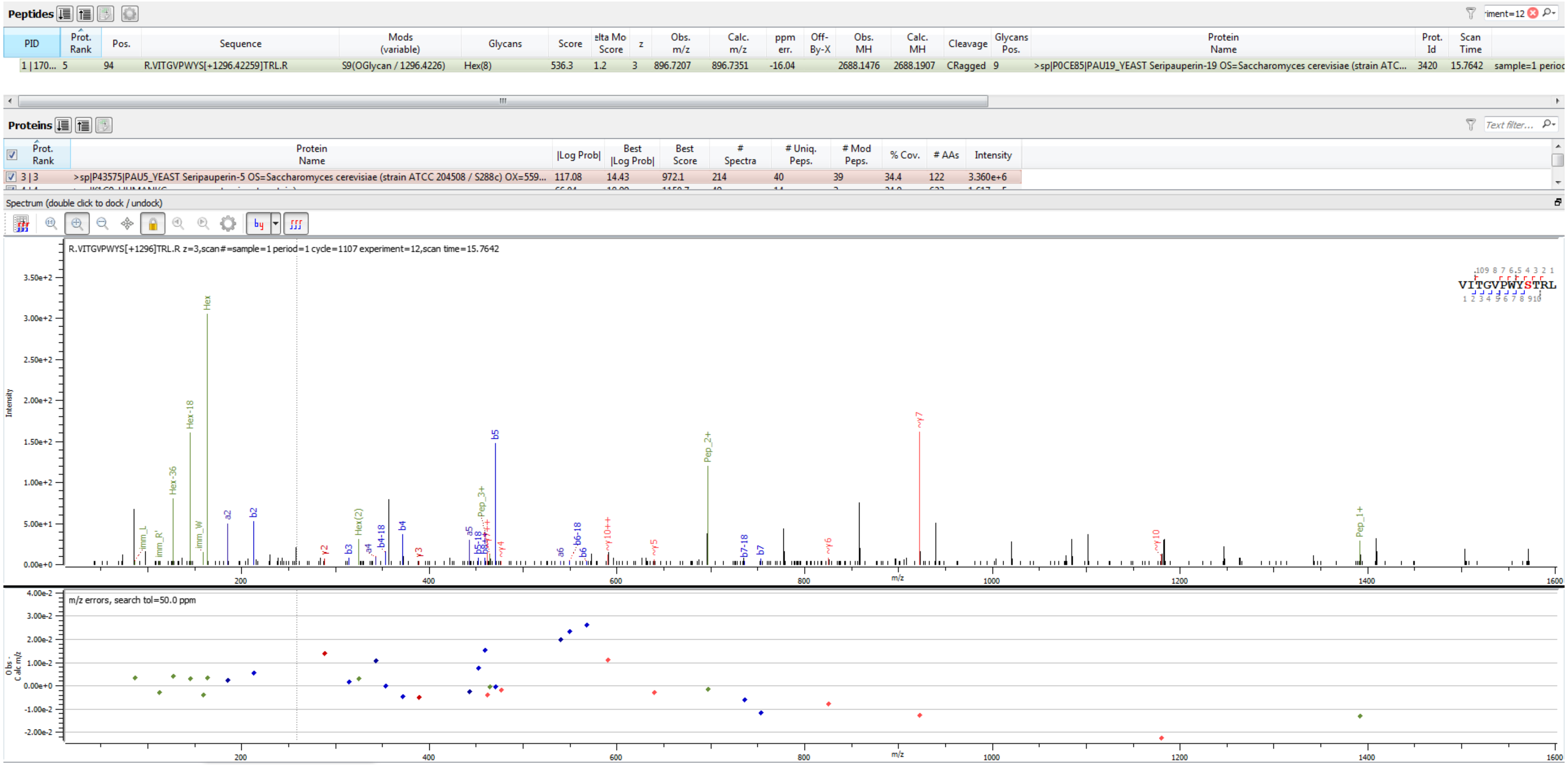


Sample S2 R.VITGVPWYS[+1296.423]TRL.R

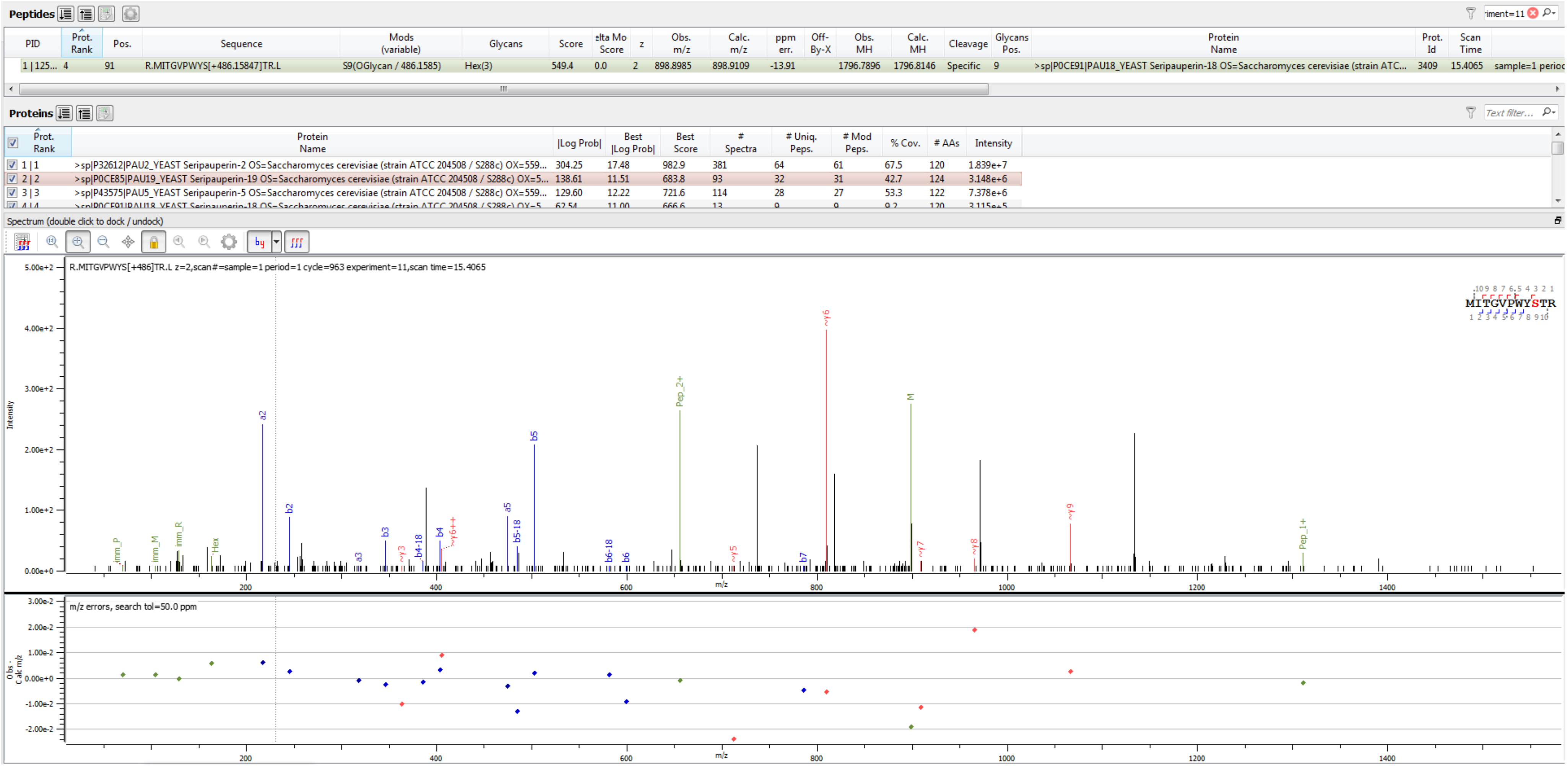


Sample S4 R.MITGVPWYS[+486.158]TR.L

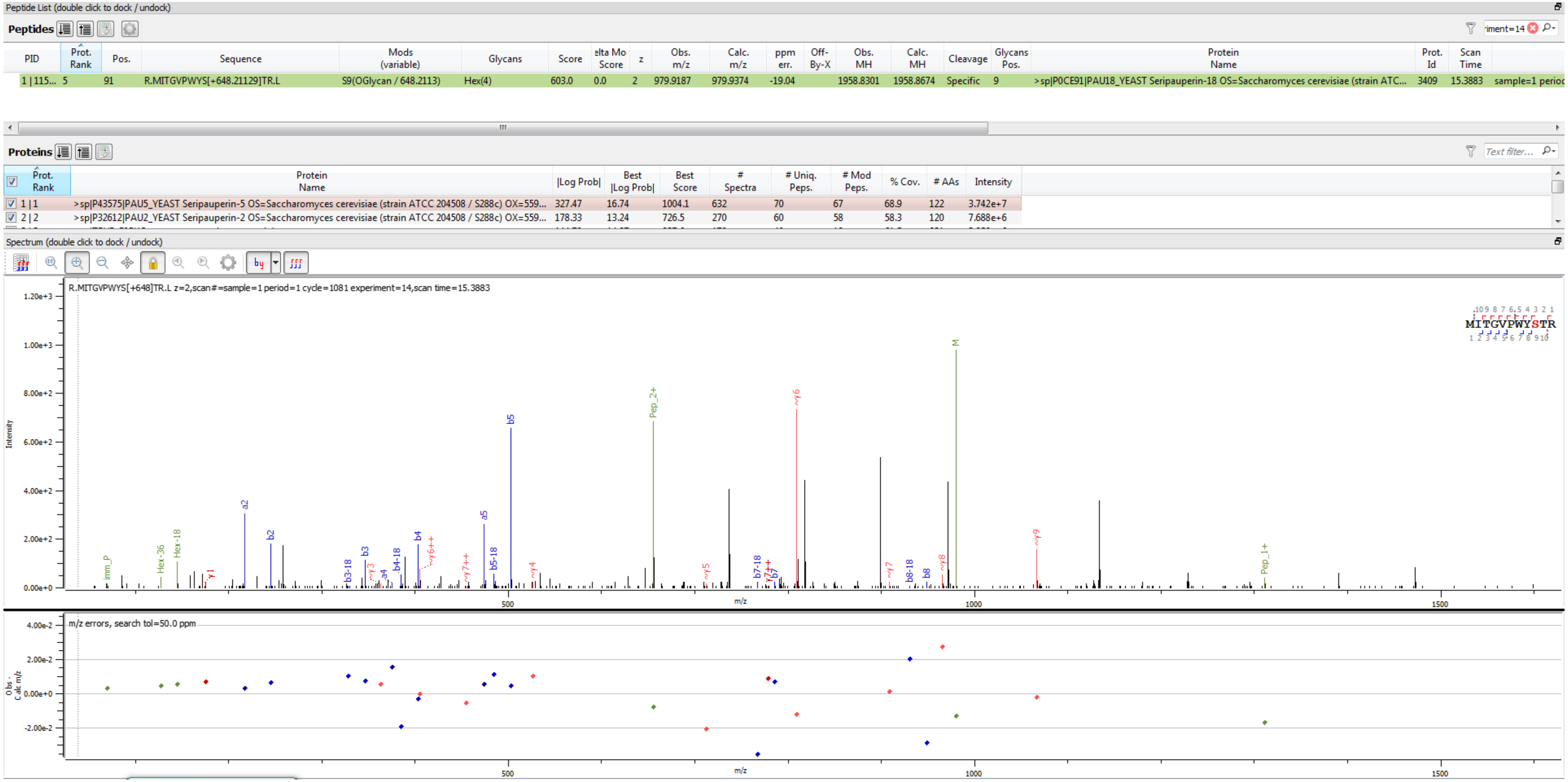

Sample S1 R.MITGVPWYS[+648.211]TR.L

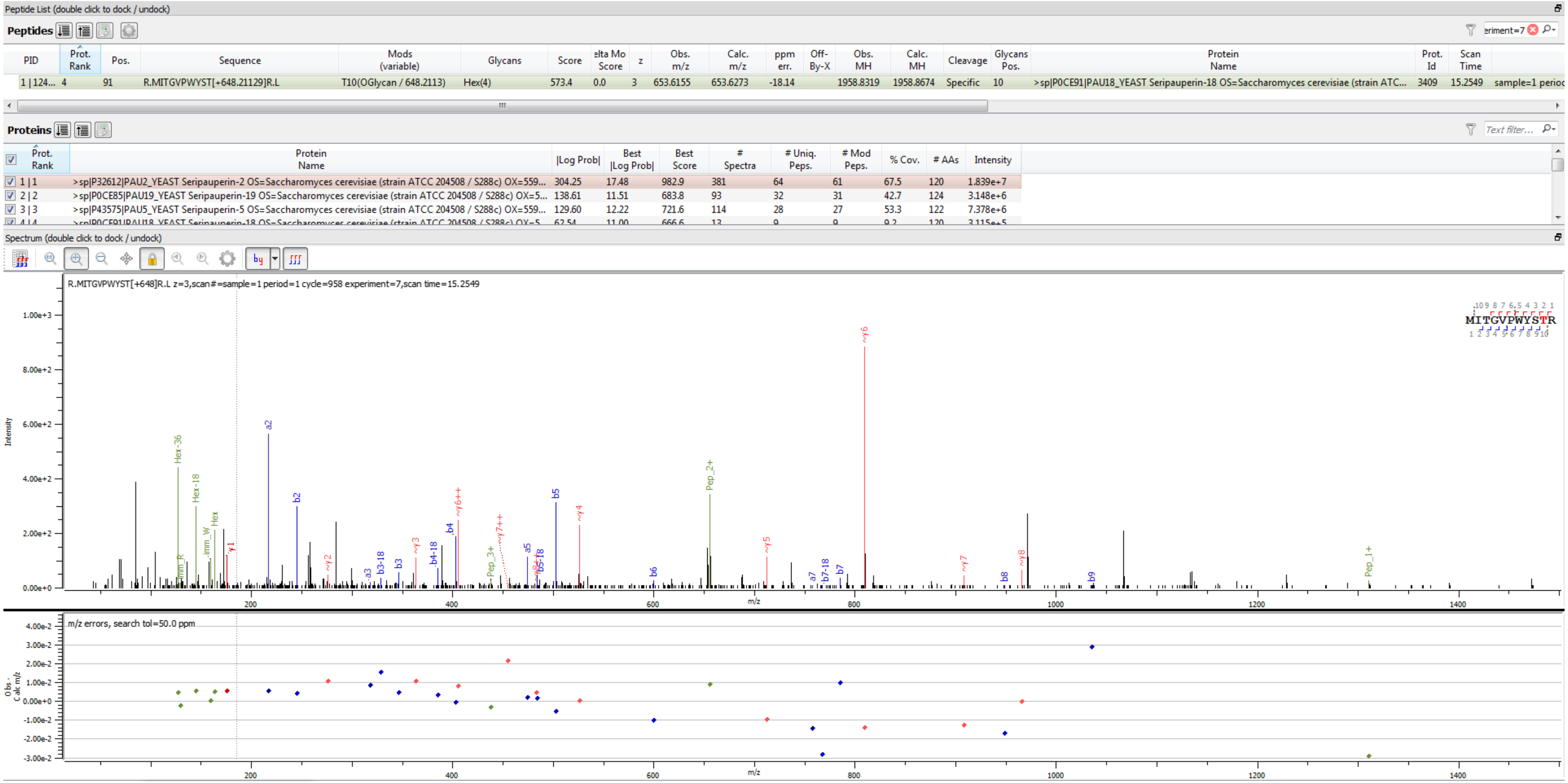


Sample S4 R.MITGVPWYS[+648.211]TR.L

**Peptides**

| PID | Prot. Rank | Pos. | Sequence | Mods (variable) | Glycans | Score | elta Mo Score | z | Obs. m/z | Calc. m/z | ppm err. | Off-By-X | Obs. MH | Calc. MH | Cleavage | Glycans Pos. | Protein Name | Prot. Id | Scan Time | |
|---|---|---|---|---|---|---|---|---|---|---|---|---|---|---|---|---|---|---|---|---|
| 1 \| 122... | 4 | 91 | R.MITGVPWYS[+810.26412]TR.L | S9(OGlycan / 810.2641) | Hex(5) | 644.0 | 1.2 | 2 | 1060.9436 | 1060.9638 | -19.05 | | 2120.8799 | 2120.9203 | Specific | 9 | >sp\|P0CE91\|PAU18_YEAST Seripauperin-18 OS=Saccharomyces cerevisiae (strain ATC... | 3409 | 15.0311 | sample=1 perioc |

**Proteins**

| Prot. Rank | Protein Name | \|Log Prob\| | Best \|Log Prob\| | Best Score | # Spectra | # Uniq. Peps. | # Mod Peps. | % Cov. | # AAs | Intensity |
|---|---|---|---|---|---|---|---|---|---|---|
| 1 \| 1 | >sp\|P32612\|PAU2_YEAST Seripauperin-2 OS=Saccharomyces cerevisiae (strain ATCC 204508 / S288c) OX=559... | 304.25 | 17.48 | 982.9 | 381 | 64 | 61 | 67.5 | 120 | 1.839e+7 |
| 2 \| 2 | >sp\|P0CE85\|PAU19_YEAST Seripauperin-19 OS=Saccharomyces cerevisiae (strain ATCC 204508 / S288c) OX=5... | 138.61 | 11.51 | 683.8 | 93 | 32 | 31 | 42.7 | 124 | 3.148e+6 |
| 3 \| 3 | >sp\|P43575\|PAU5_YEAST Seripauperin-5 OS=Saccharomyces cerevisiae (strain ATCC 204508 / S288c) OX=559... | 129.60 | 12.22 | 721.6 | 114 | 28 | 27 | 53.3 | 122 | 7.378e+6 |
| 4 \| 4 | >sp\|P0CE91\|PAU18_YEAST Seripauperin-18 OS=Saccharomyces cerevisiae (strain ATCC 204508 / S288c) OX=5... | 62.54 | 11.00 | 666.6 | 13 | 9 | 9 | 9.2 | 120 | 3.115e+5 |

Spectrum (double click to dock / undock)

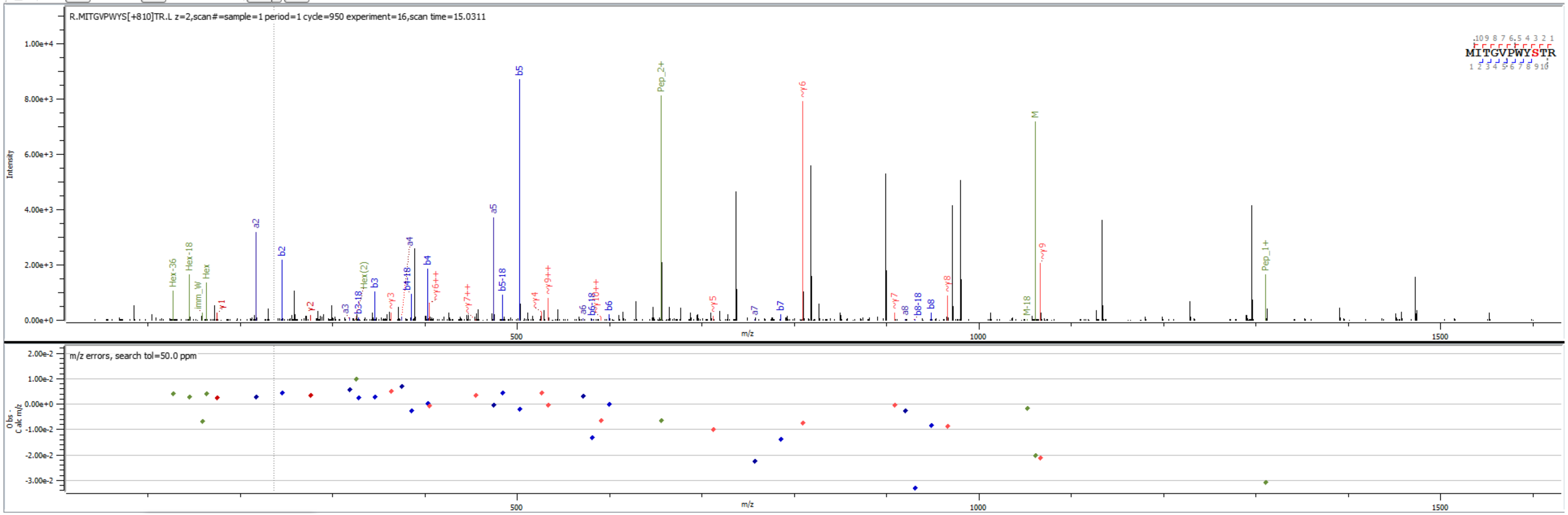


Sample S4 R.MITGVPWYS[+810.264]TR.L

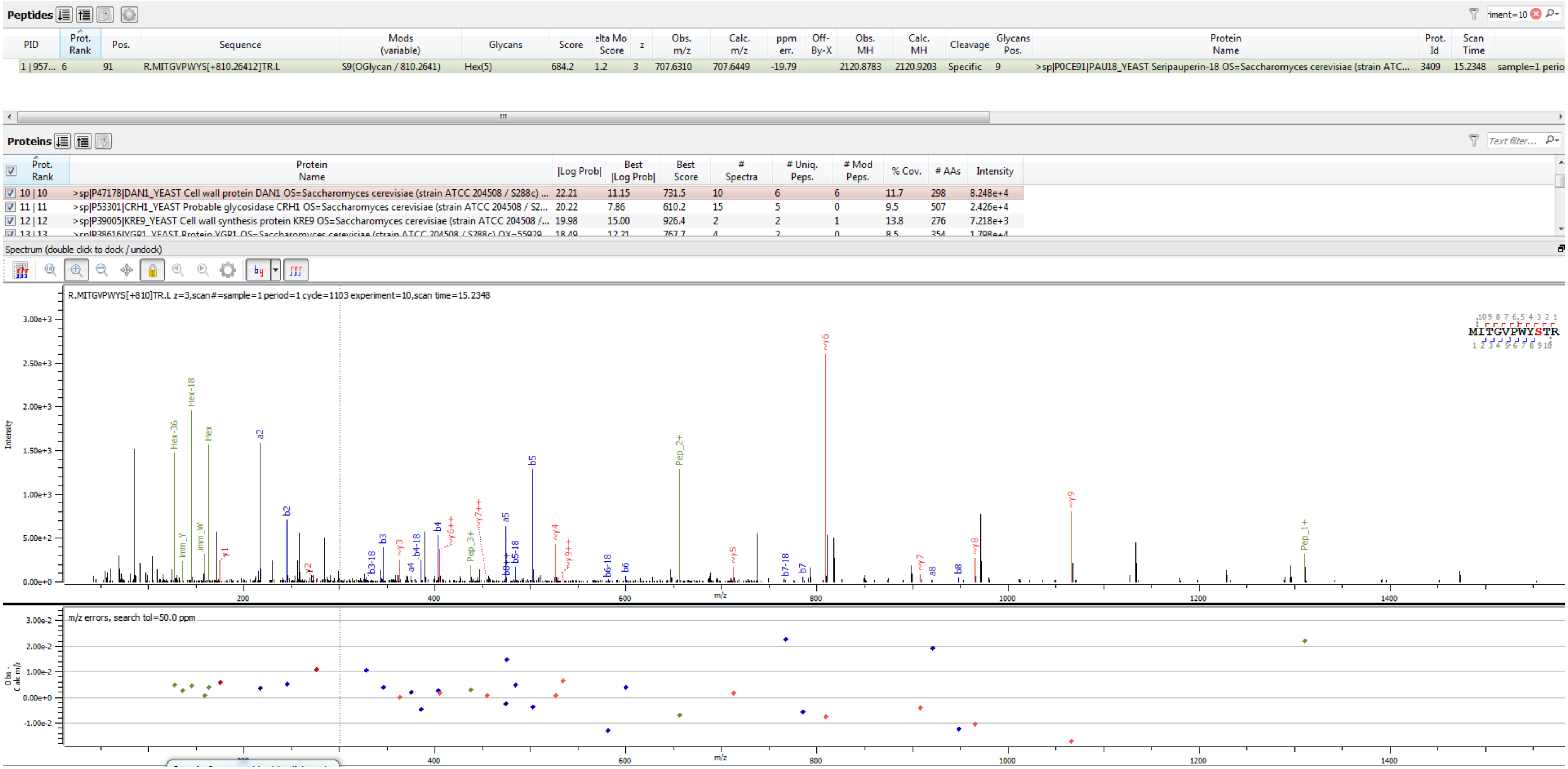


Sample S6 R.MITGVPWYS[+810.264]TR.L

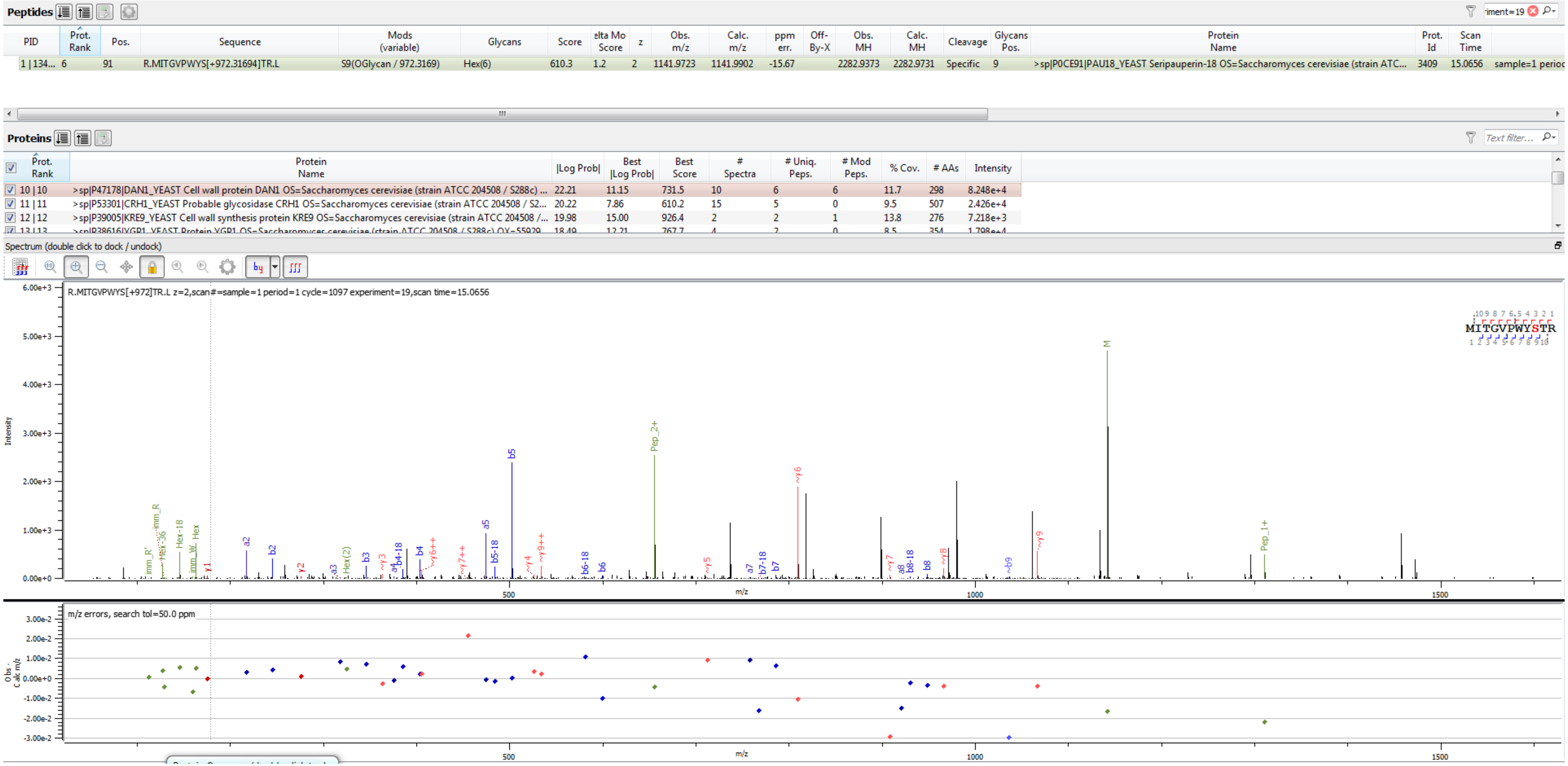


Sample S6 R.MITGVPWYS[+972.317]TR.L

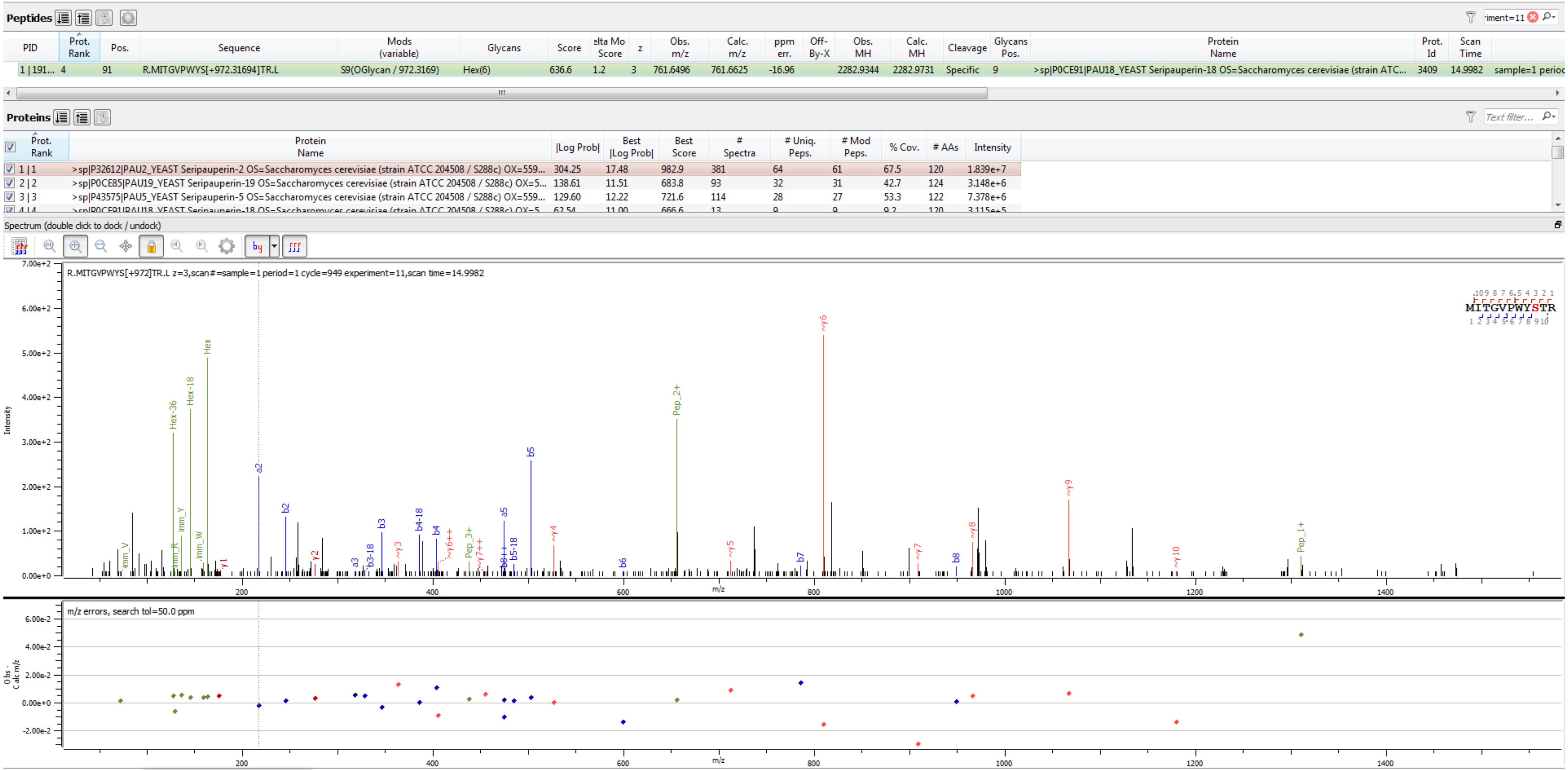


Sample S4 R.MITGVPWYS[+972.317]TR.L

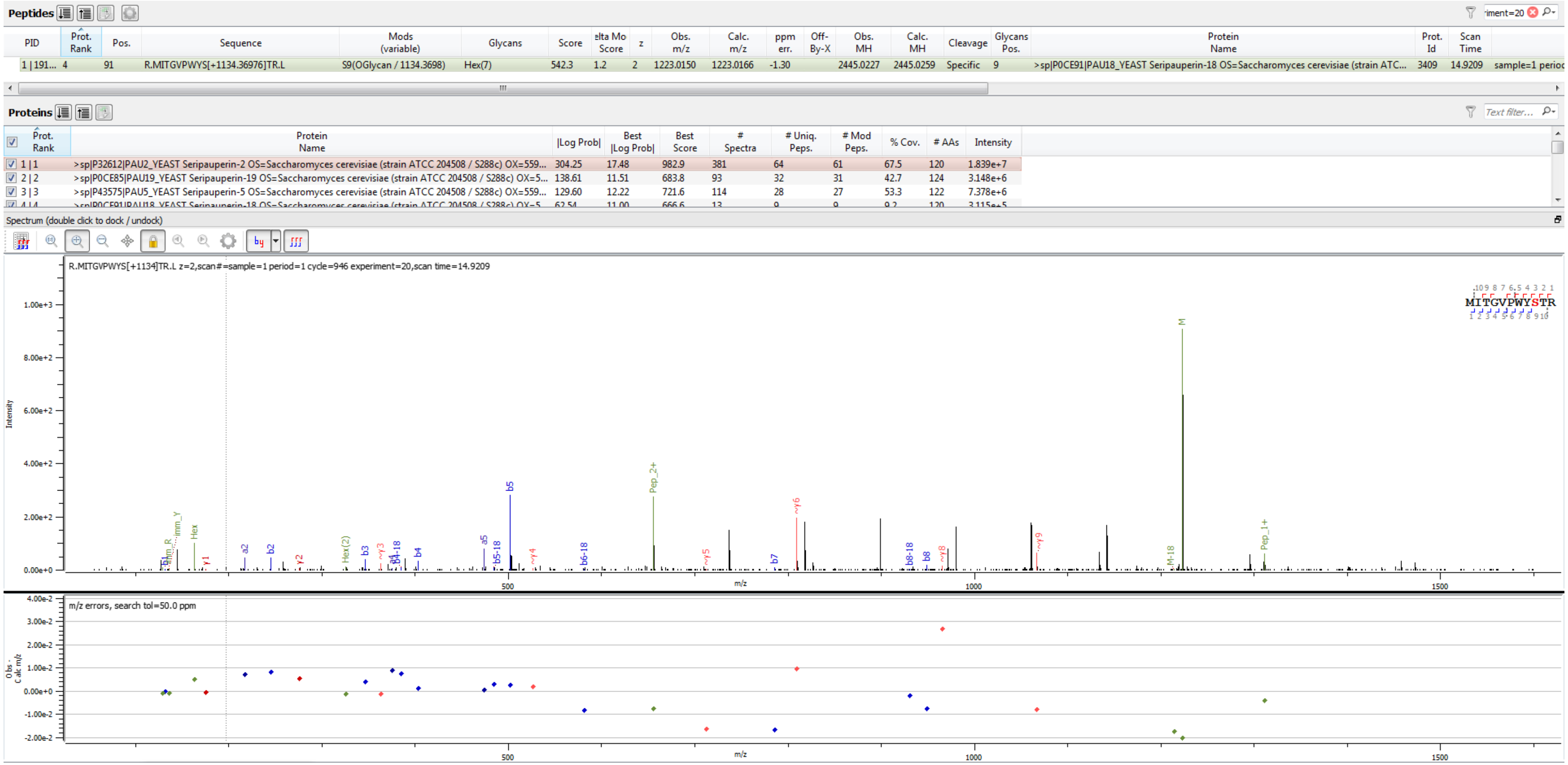

Peptides
PID | Prot. Rank | Pos. | Sequence | Mods (variable) | Glycans | Score | elta Mo Score | z | Obs. m/z | Calc. m/z | ppm err. | Off-By-X | Obs. MH | Calc. MH | Cleavage | Glycans Pos. | Protein Name | Prot. Id | Scan Time
1 | 191... 4 | 91 | R.MITGVPWYS[+1134.36976]TR.L | S9(OGlycan / 1134.3698) | Hex(7) | 542.3 | 1.2 | 2 | 1223.0150 | 1223.0166 | -1.30 | | 2445.0227 | 2445.0259 | Specific | 9 | >sp|P0CE91|PAU18_YEAST Seripauperin-18 OS=Saccharomyces cerevisiae (strain ATC... | 3409 | 14.9209 | sample=1 perioc
Proteins
Prot. Rank | Protein Name | |Log Prob| | Best |Log Prob| | Best Score | # Spectra | # Uniq. Peps. | # Mod Peps. | % Cov. | # AAs | Intensity
1 | 1 | >sp|P32612|PAU2_YEAST Seripauperin-2 OS=Saccharomyces cerevisiae (strain ATCC 204508 / S288c) OX=559... | 304.25 | 17.48 | 982.9 | 381 | 64 | 61 | 67.5 | 120 | 1.839e+7
2 | 2 | >sp|P0CE85|PAU19_YEAST Seripauperin-19 OS=Saccharomyces cerevisiae (strain ATCC 204508 / S288c) OX=5... | 138.61 | 11.51 | 683.8 | 93 | 32 | 31 | 42.7 | 124 | 3.148e+6
3 | 3 | >sp|P43575|PAU5_YEAST Seripauperin-5 OS=Saccharomyces cerevisiae (strain ATCC 204508 / S288c) OX=559... | 129.60 | 12.22 | 721.6 | 114 | 28 | 27 | 53.3 | 122 | 7.378e+6
Spectrum (double click to dock / undock)
R.MITGVPWYS[+1134]TR.L z=2,scan#=sample=1 period=1 cycle=946 experiment=20,scan time=14.9209
MITGVPWYSTR
Intensity
m/z
m/z errors, search tol=50.0 ppm
Obs - Calc m/z


Sample S4

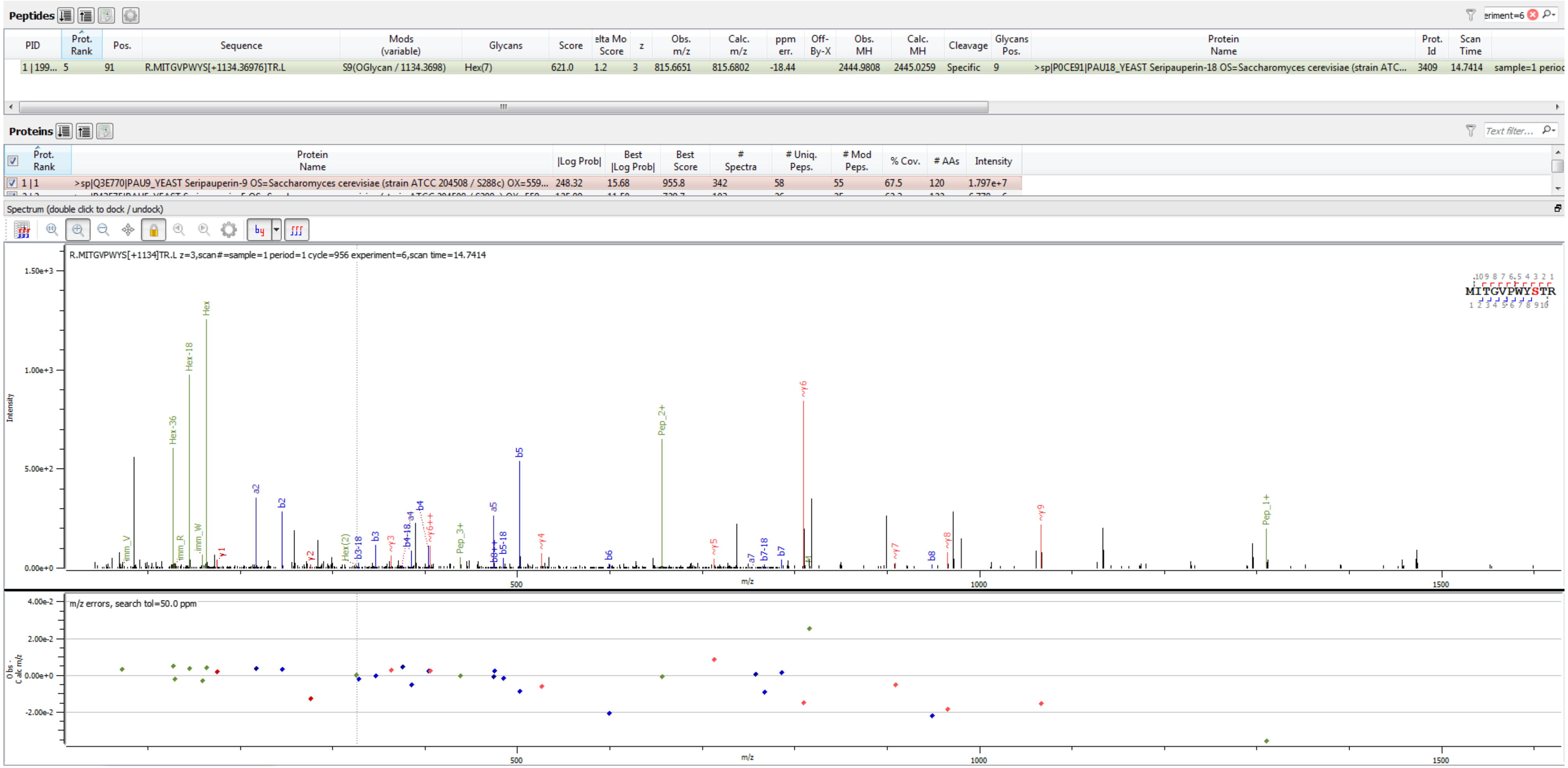


Sample S3 R.MITGVPWYS[+1134.370]TR.L

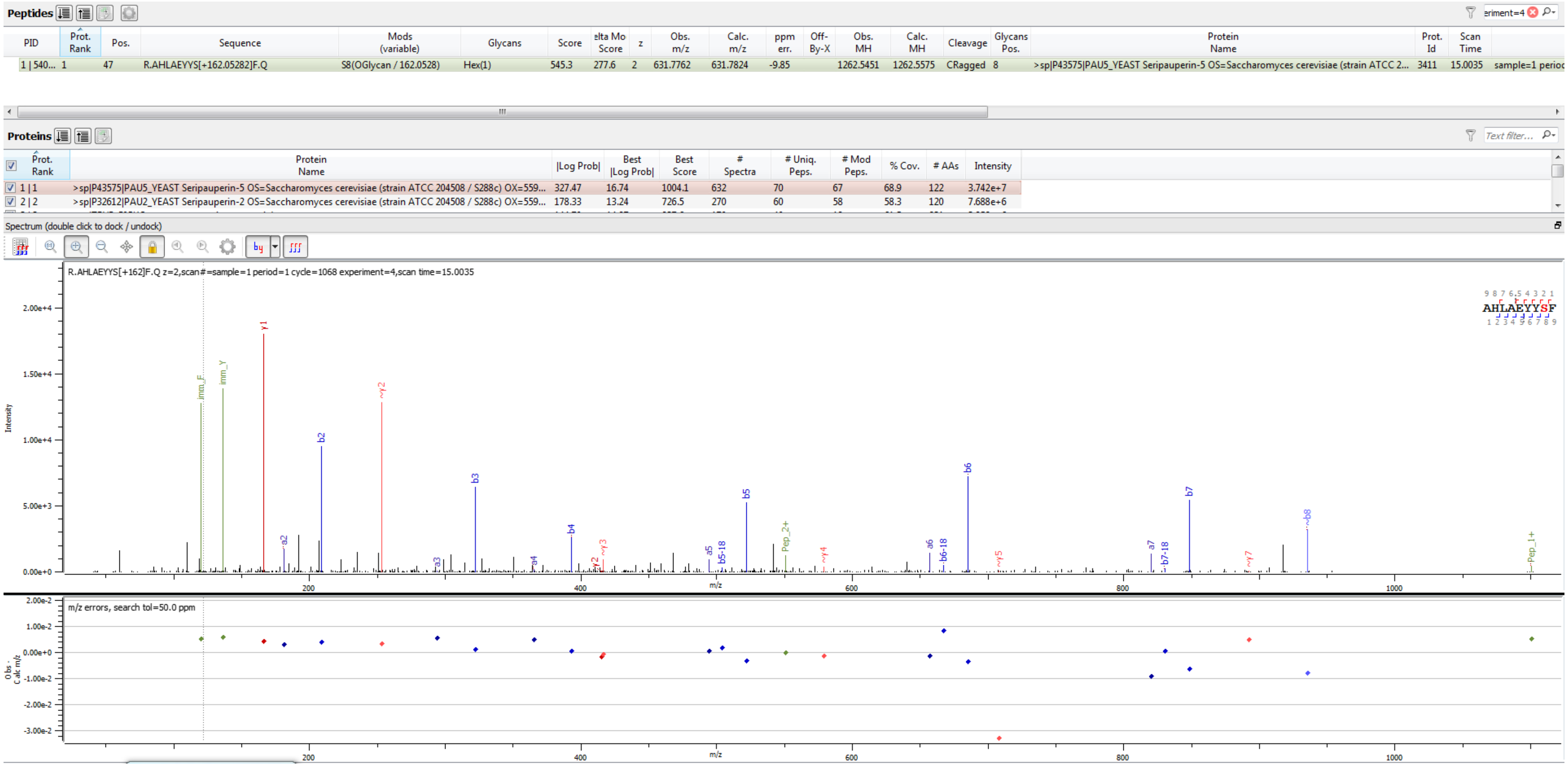


Sample S1 R.AHLAEYYS[+162.053]F.Q

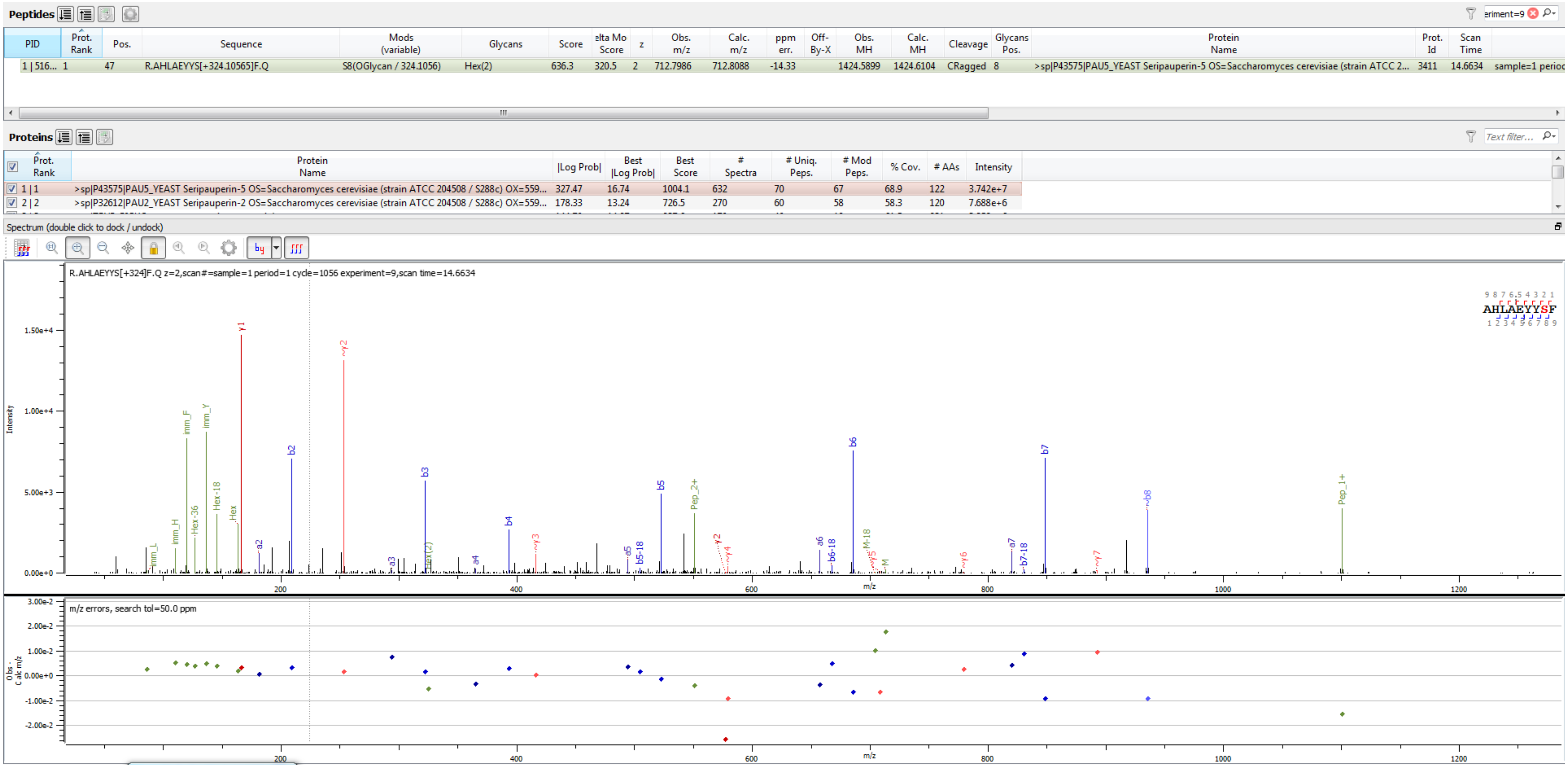


Sample S1 R.AHLAEYYS[+324.106]F.Q

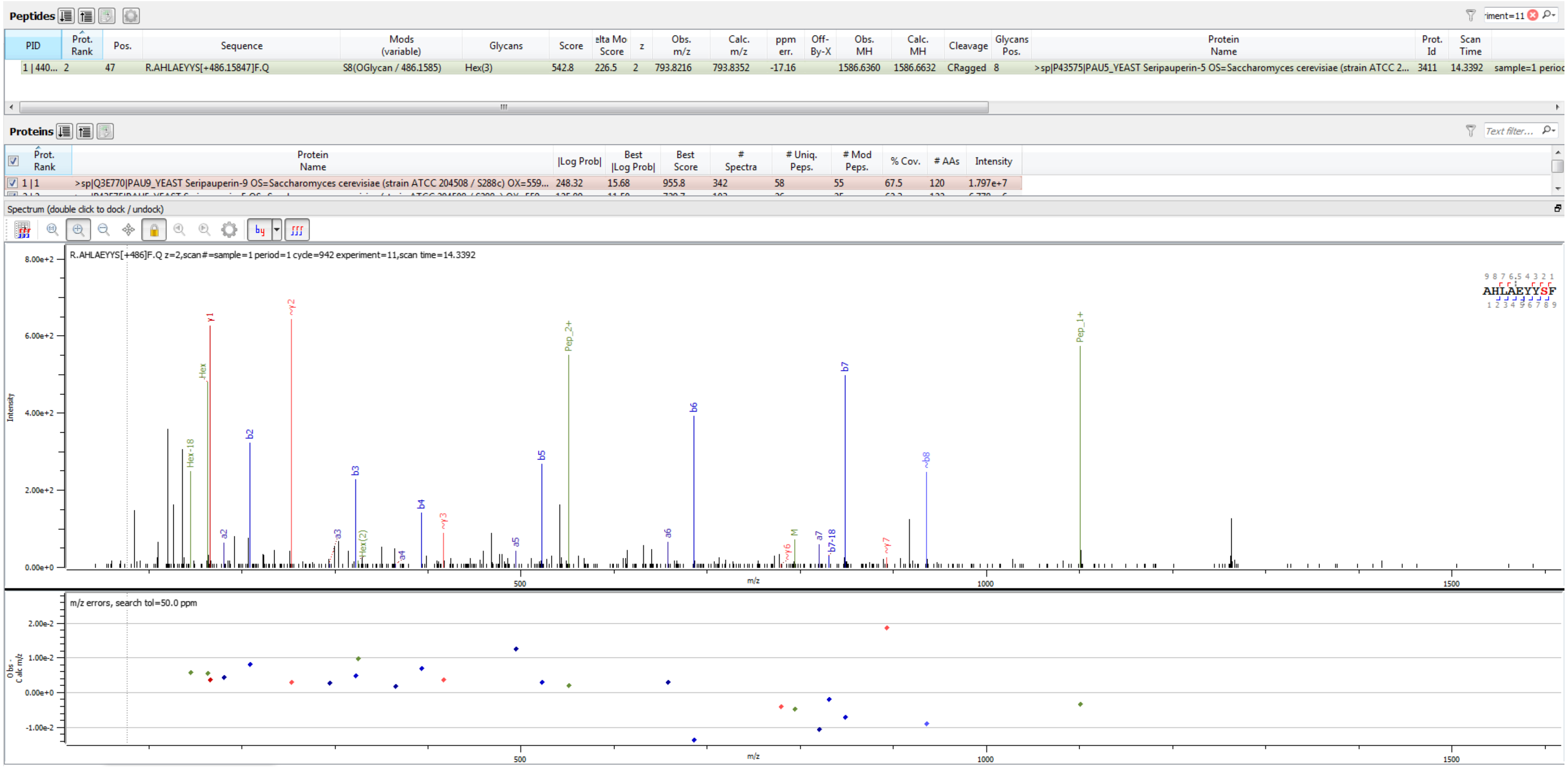


Sample S3 R.AHLAEYYS[+486.158]F.Q

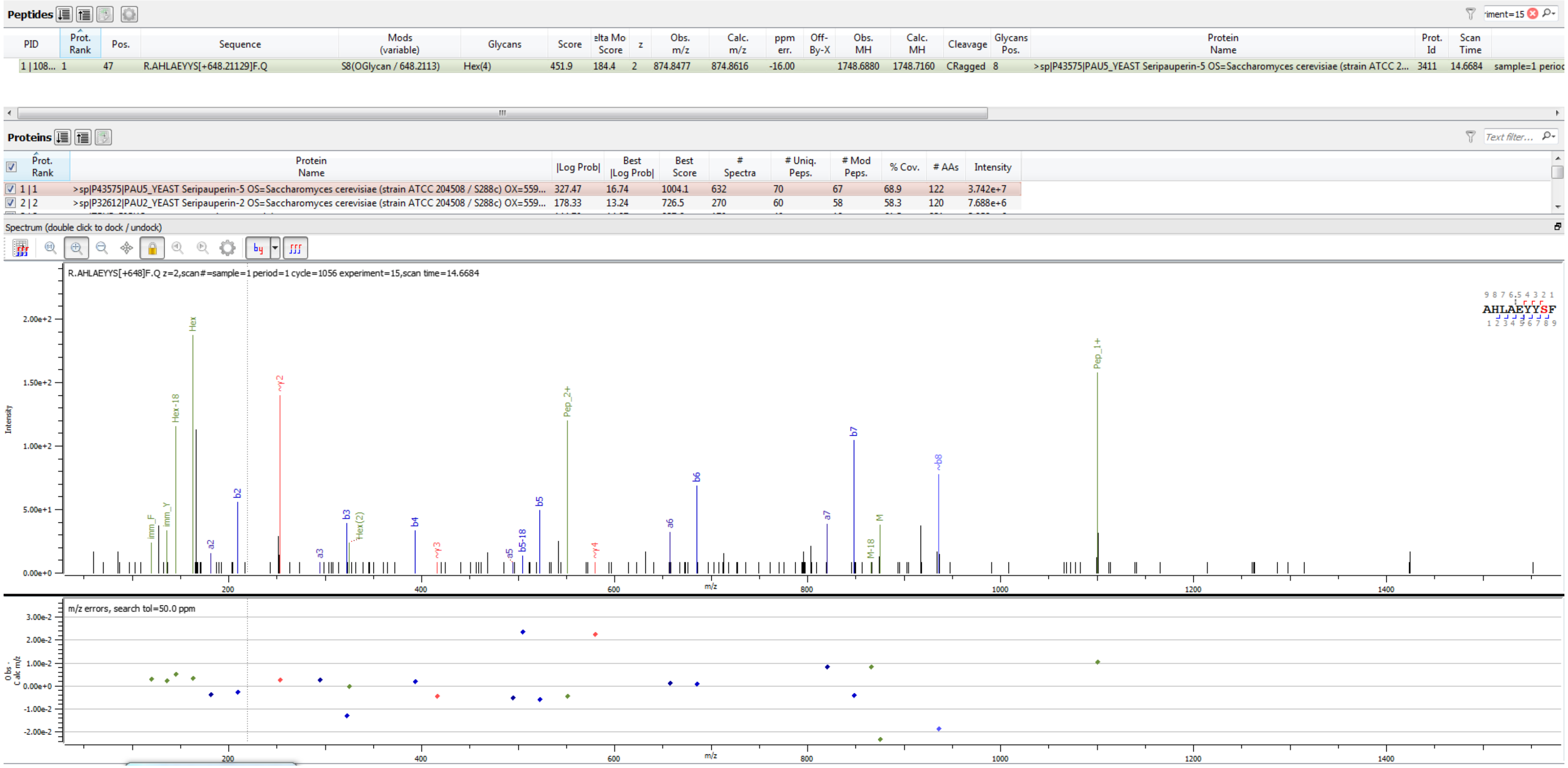


Sample S1 R.AHLAEYYS[+648.211]F.Q

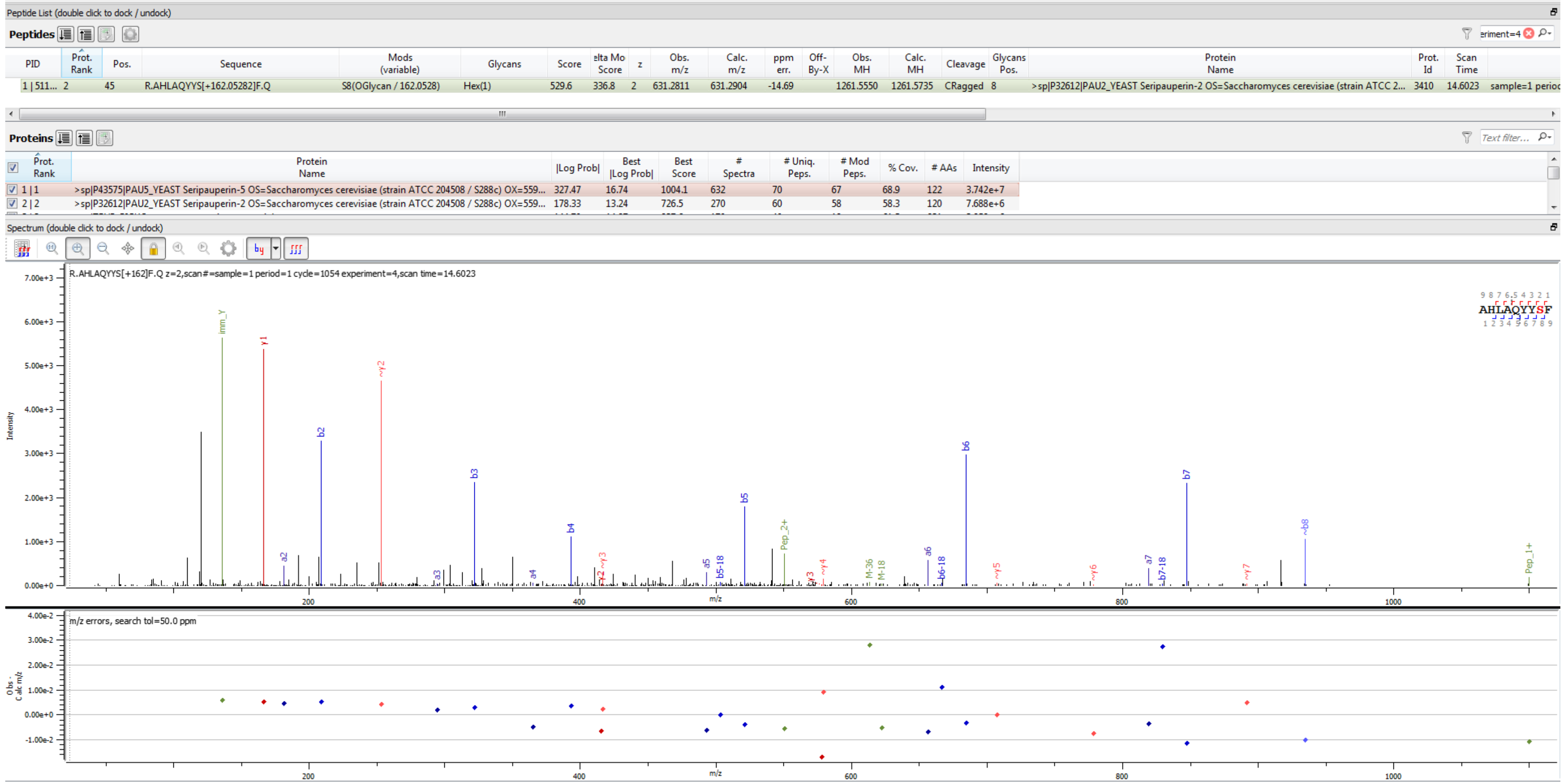


Sample S1 R.AHLAQYYS[+162.053]F.Q

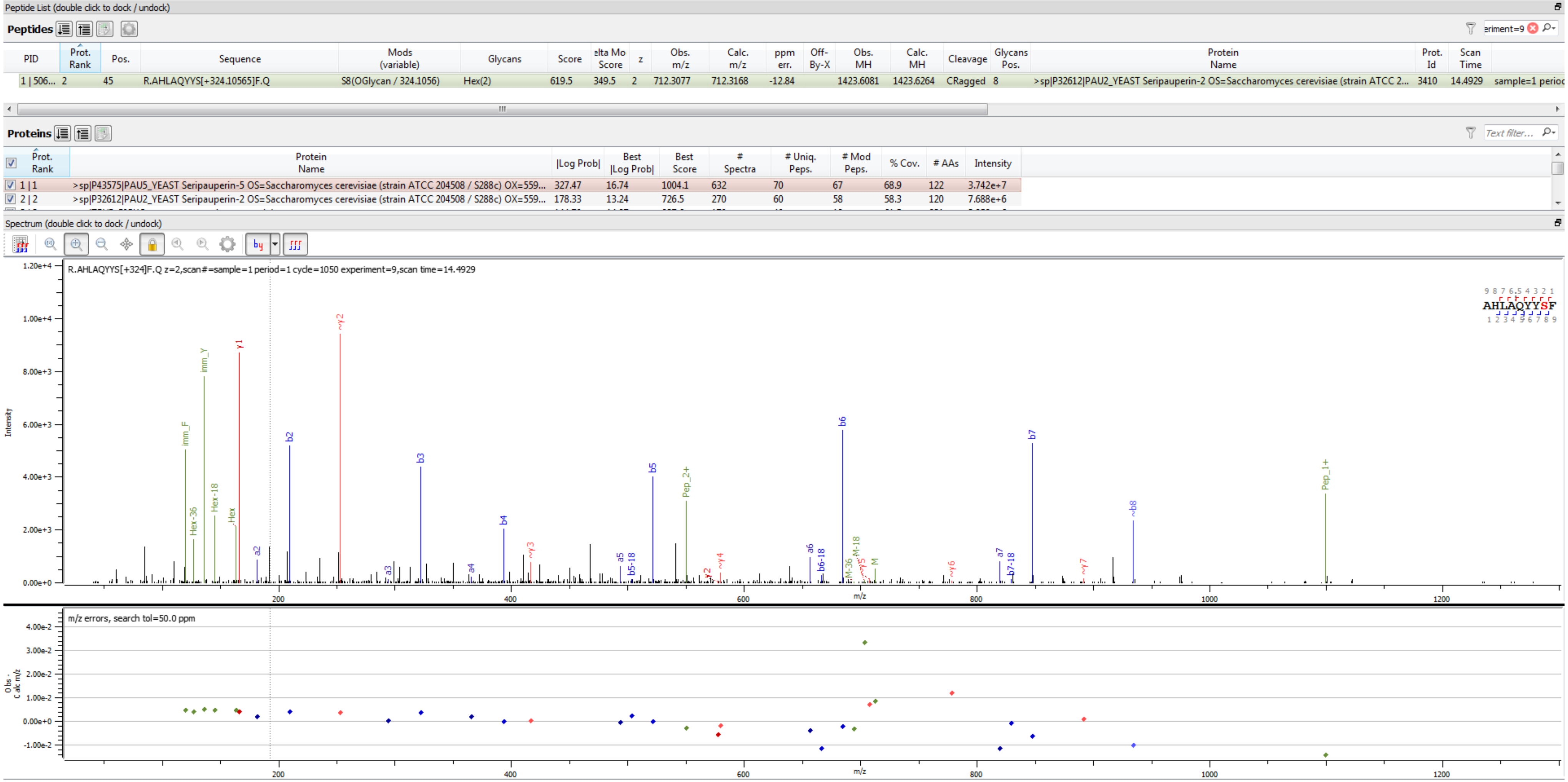


Sample S1 R.AHLAQYYS[+324.106]F.Q

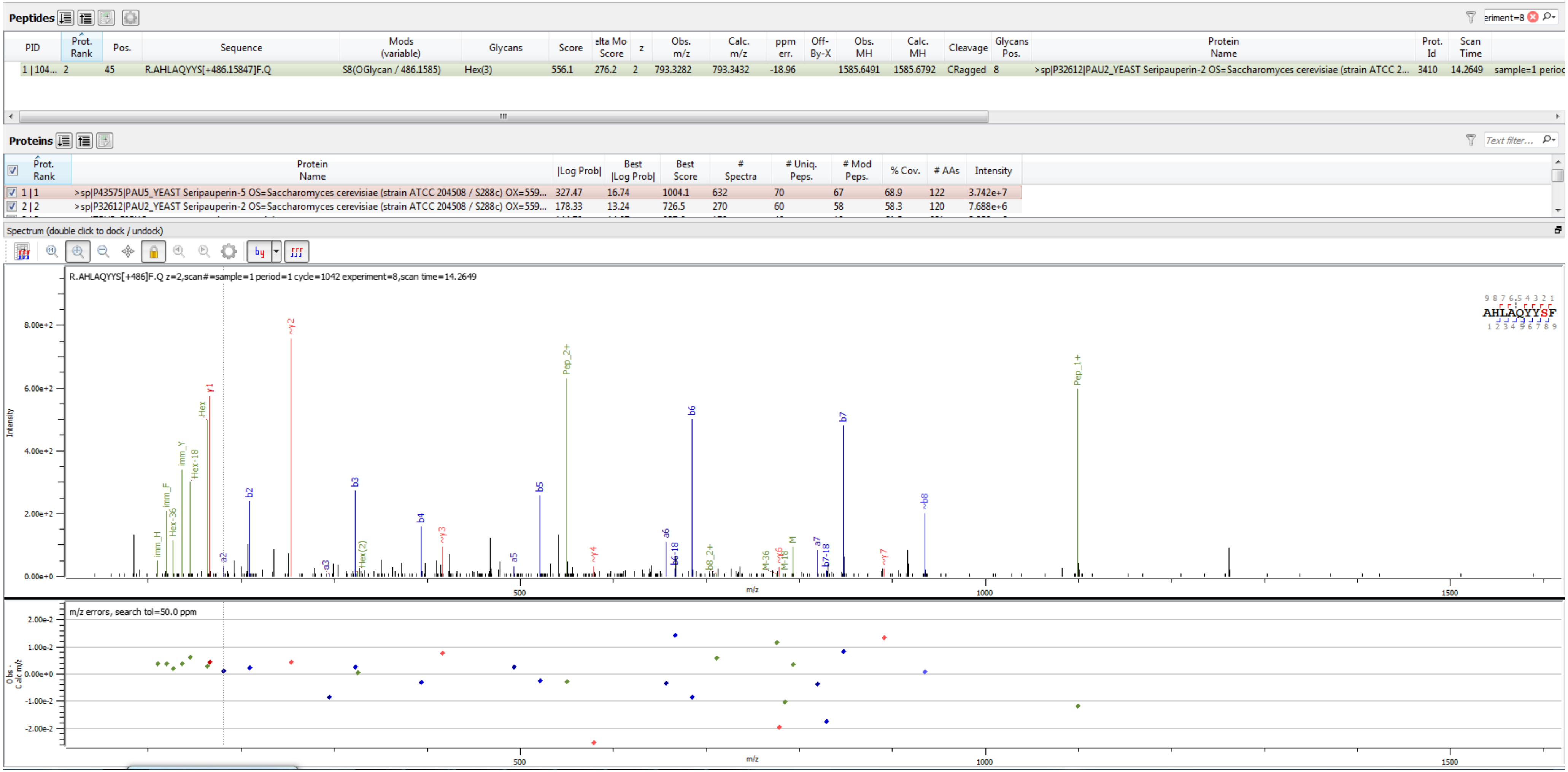


Sample S1 R.AHLAQYYS[+486.158]F.Q

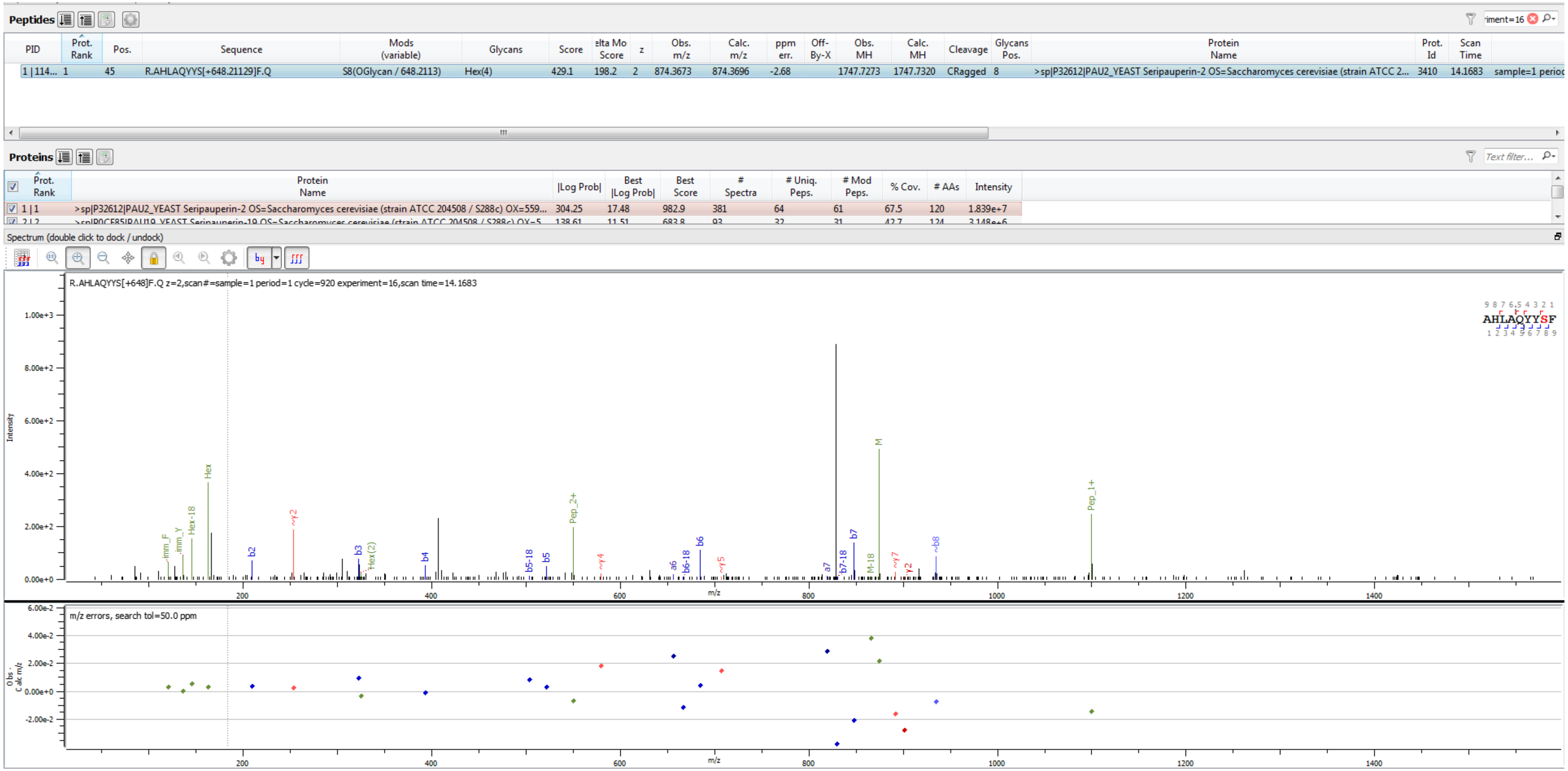

Sample S4 R.AHLAQYYS[+648.211]F.Q

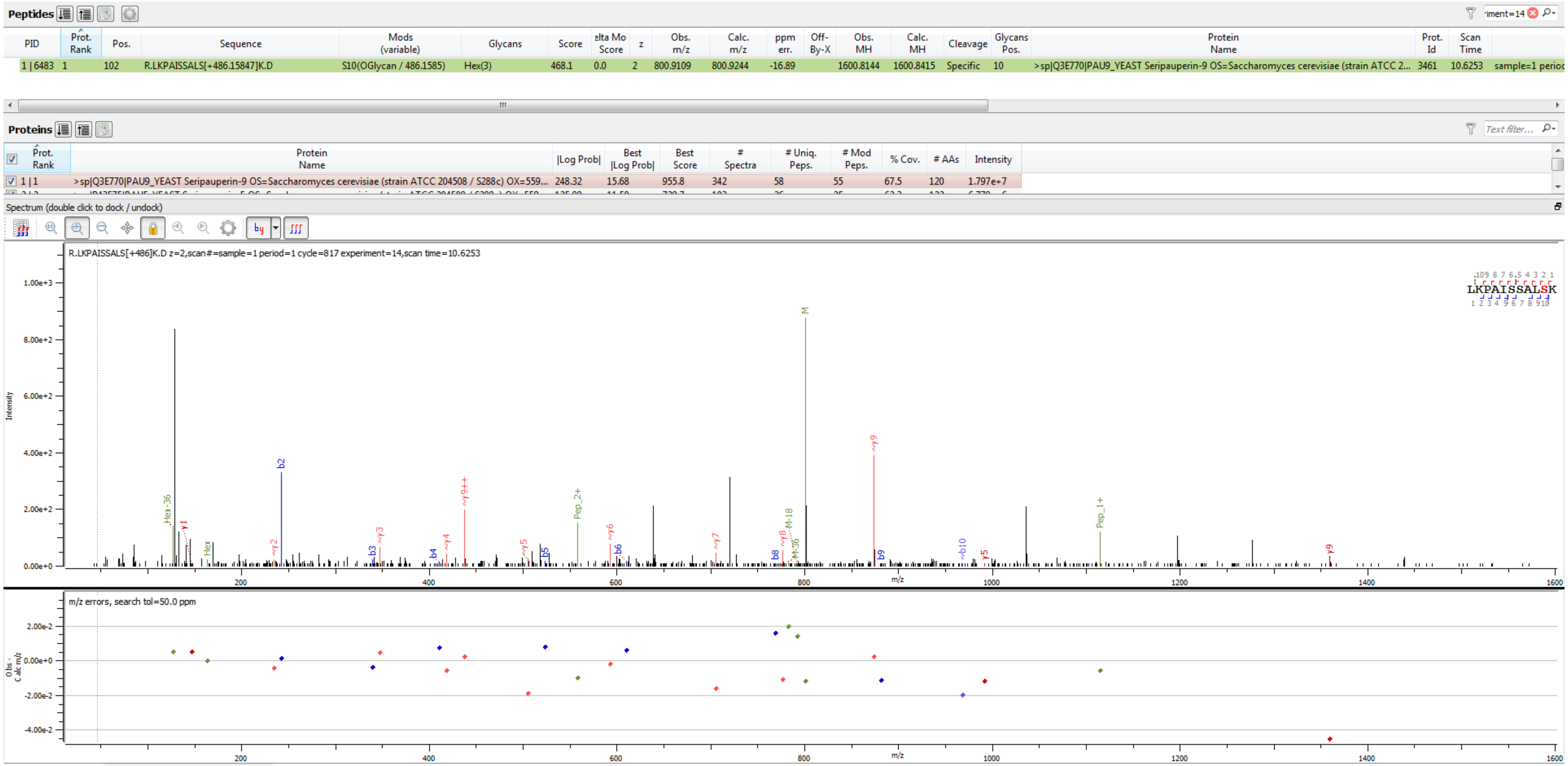


Sample S3 R.LKPAISSALS[+486.158]K.D

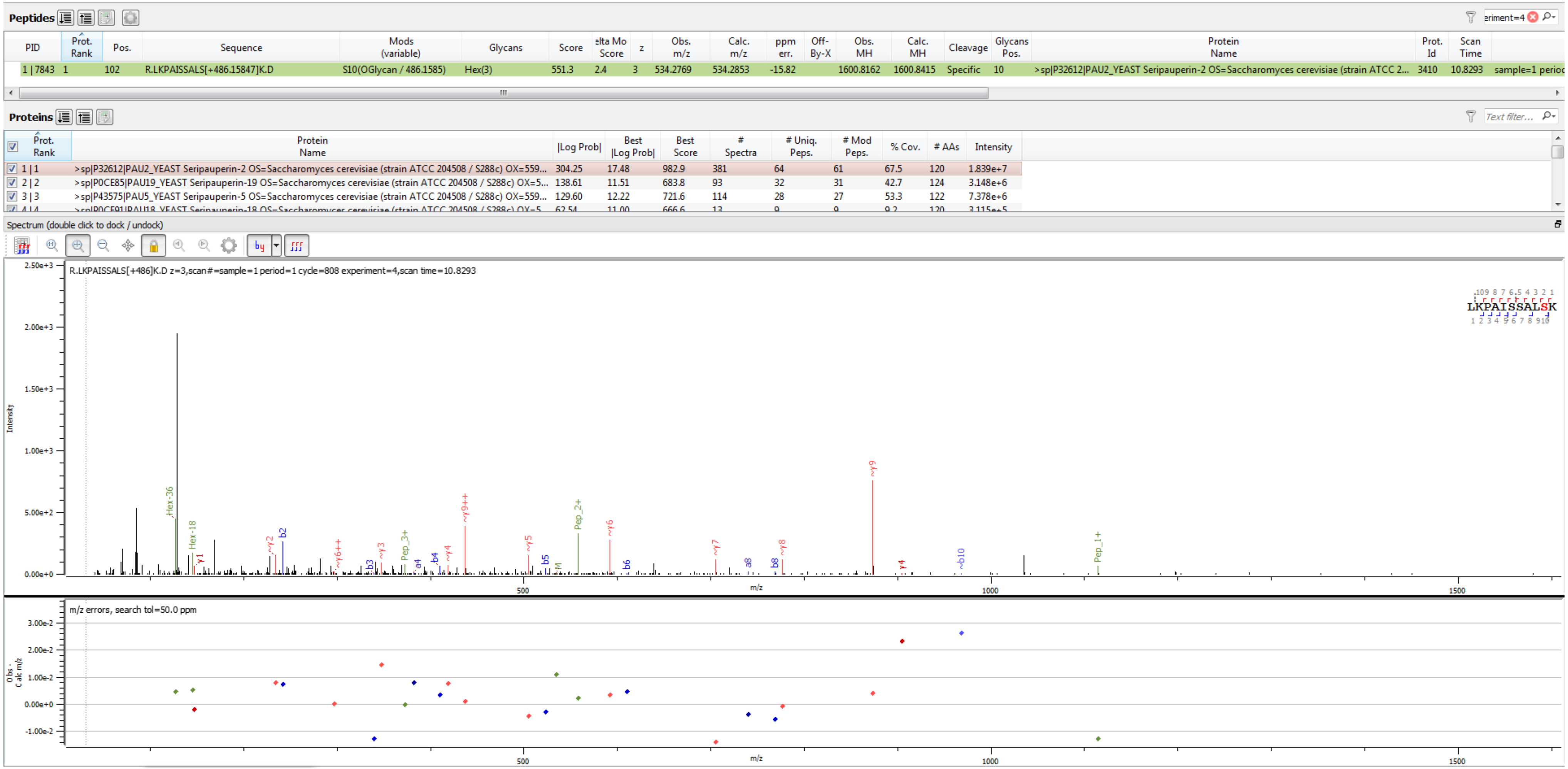


Sample S4 R.LKPAISSALS[+486.158]K.D

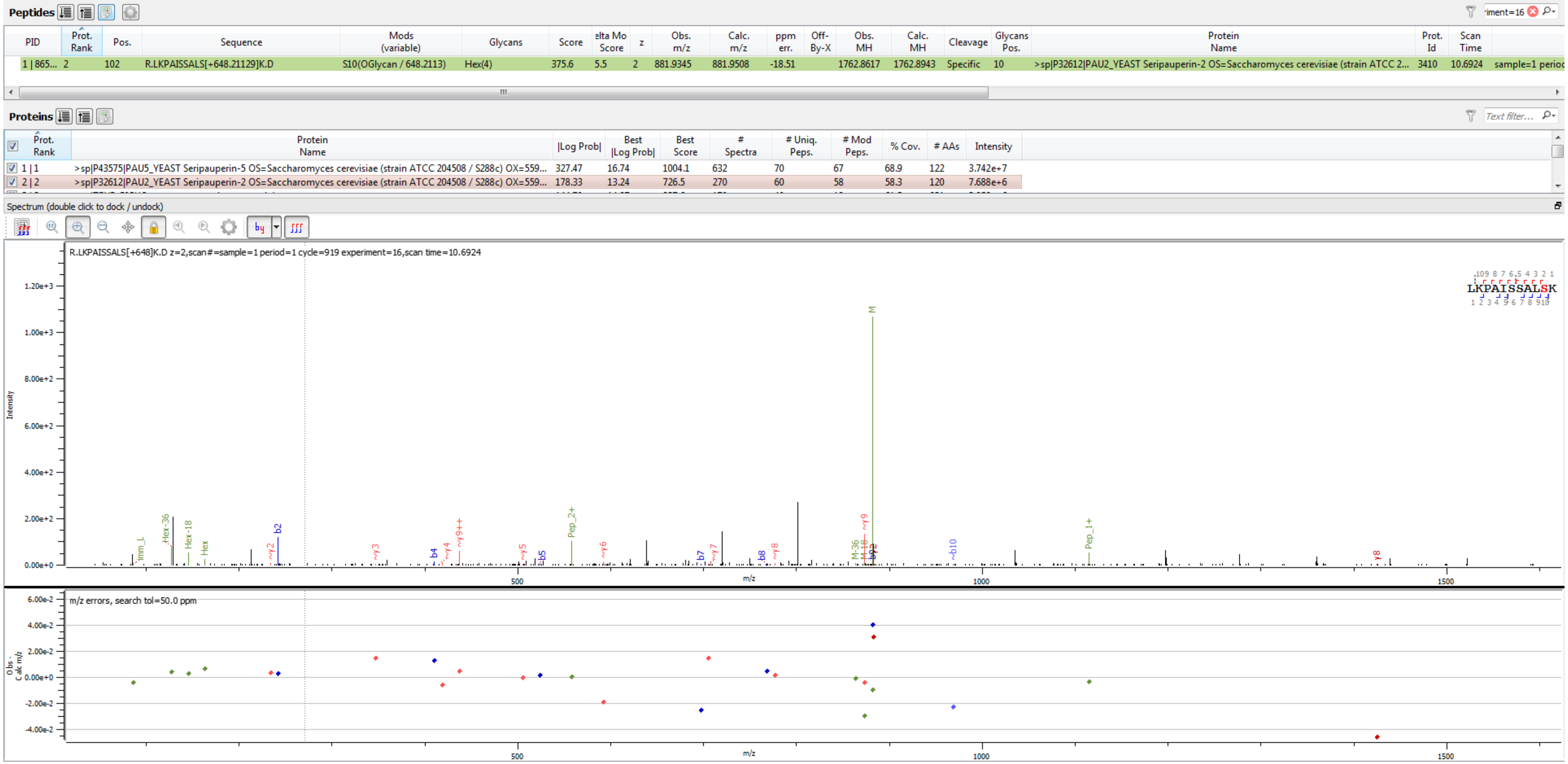


Sample S1 R.LKPAISSALS[+648.211]K.D

**Peptides**

| PID | Prot. Rank | Pos. | Sequence | Mods (variable) | Glycans | Score | elta Mo Score | z | Obs. m/z | Calc. m/z | ppm err. | Off-By-X | Obs. MH | Calc. MH | Cleavage | Glycans Pos. | Protein Name | Prot. Id | Scan Time | |
|---|---|---|---|---|---|---|---|---|---|---|---|---|---|---|---|---|---|---|---|---|
| 1 \| 2290 | 1 | 102 | R.LKPAISS[+648.21129]ALSK.D | S7(OGlycan / 648.2113) | Hex(4) | 485.6 | 0.0 | 3 | 588.2979 | 588.3030 | -8.60 | | 1762.8791 | 1762.8943 | Specific | 7 | >sp\|Q3E770\|PAU9_YEAST Seripauperin-9 OS=Saccharomyces cerevisiae (strain ATCC 2... | 3461 | 10.5024 | sample=1 perioc |

**Proteins**

| Prot. Rank | Protein Name | \|Log Prob\| | Best \|Log Prob\| | Best Score | # Spectra | # Uniq. Peps. | # Mod Peps. | % Cov. | # AAs | Intensity |
|---|---|---|---|---|---|---|---|---|---|---|
| 1 \| 1 | >sp\|Q3E770\|PAU9_YEAST Seripauperin-9 OS=Saccharomyces cerevisiae (strain ATCC 204508 / S288c) OX=559... | 288.60 | 16.66 | 988.6 | 452 | 61 | 56 | 60.8 | 120 | 1.023e+7 |
| 2 \| 2 | >sp\|Q07987\|PAU23_YEAST Seripauperin-23 OS=Saccharomyces cerevisiae (strain ATCC 204508 / S288c) OX=5... | 160.38 | 13.28 | 718.6 | 124 | 39 | 36 | 43.5 | 124 | 2.323e+6 |
| 3 \| 3 | >sp\|P43575\|PAU5_YEAST Seripauperin-5 OS=Saccharomyces cerevisiae (strain ATCC 204508 / S288c) OX=559... | 142.55 | 14.15 | 737.1 | 167 | 35 | 33 | 42.6 | 122 | 6.136e+6 |

Spectrum (double click to dock / undock)

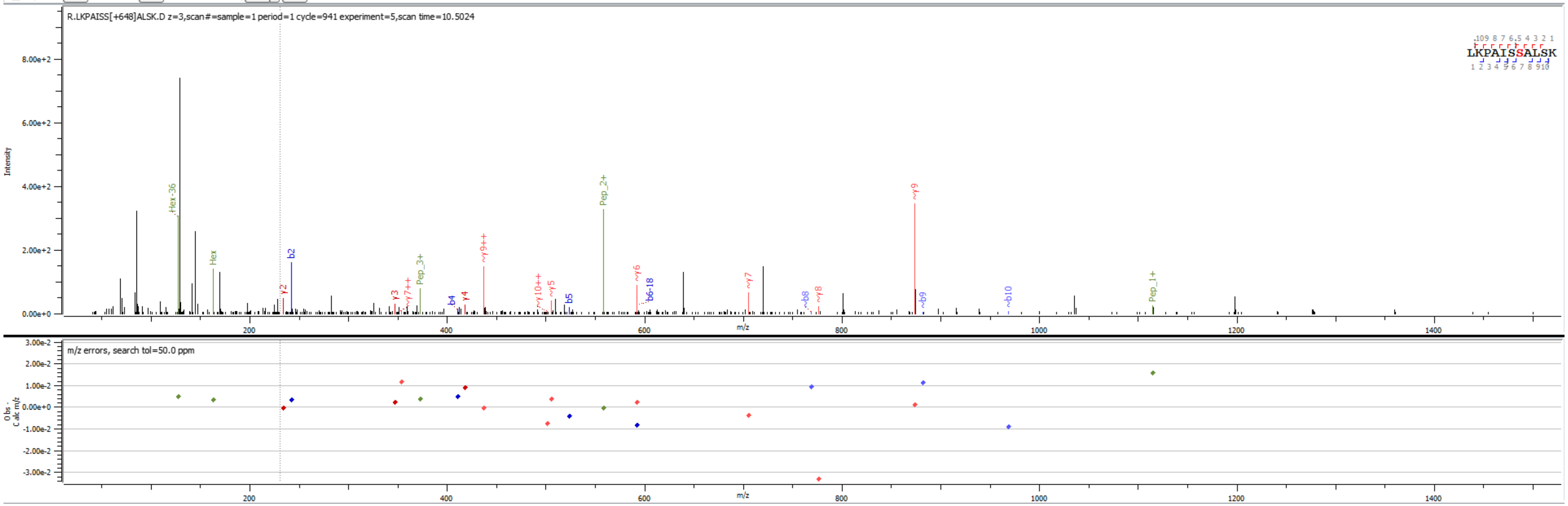


Sample S6 R.LKPAISS[+648.211]ALSK.D

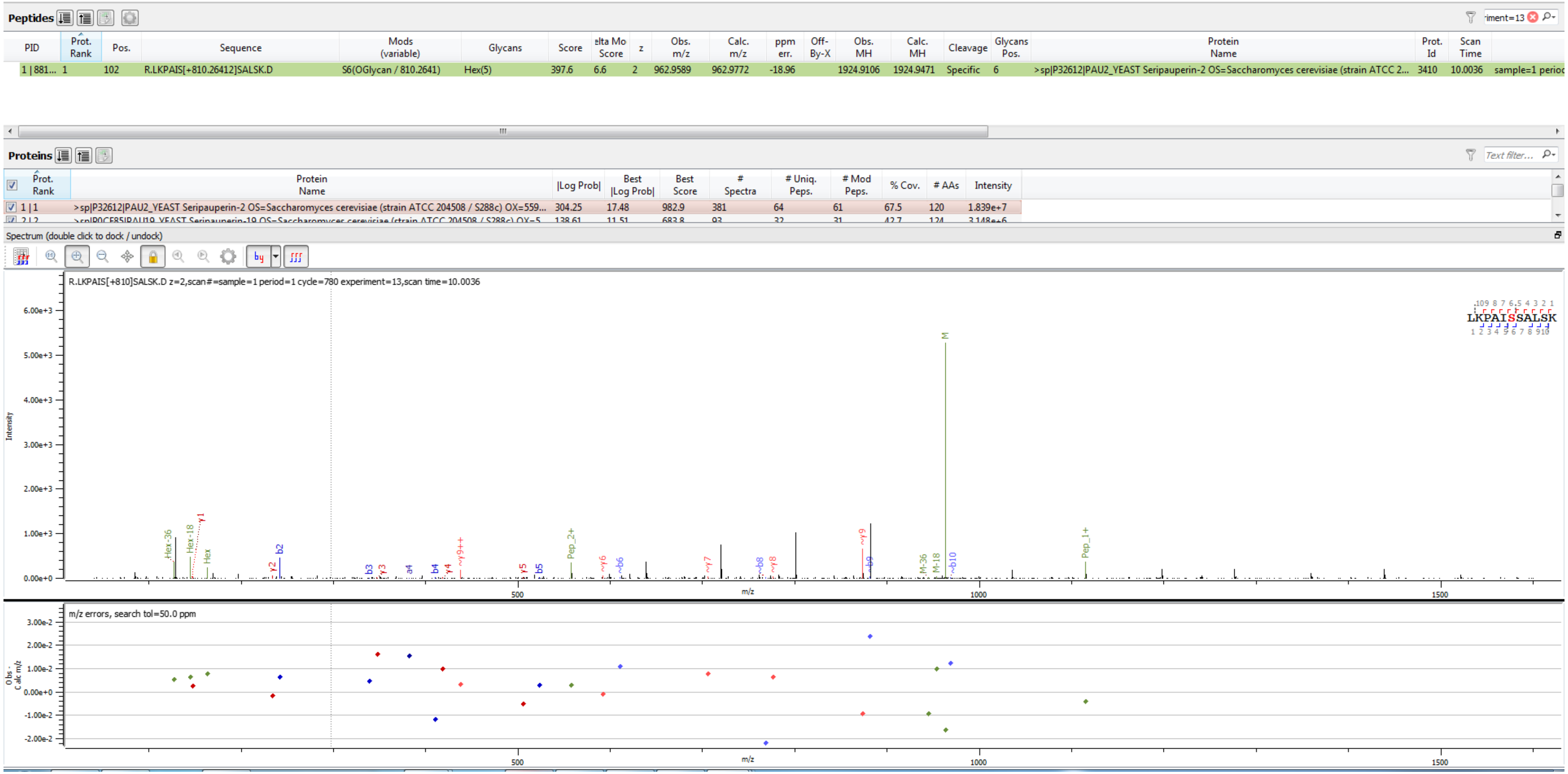


Sample S1 R.LKPAIS[+810.264]SALSK.D

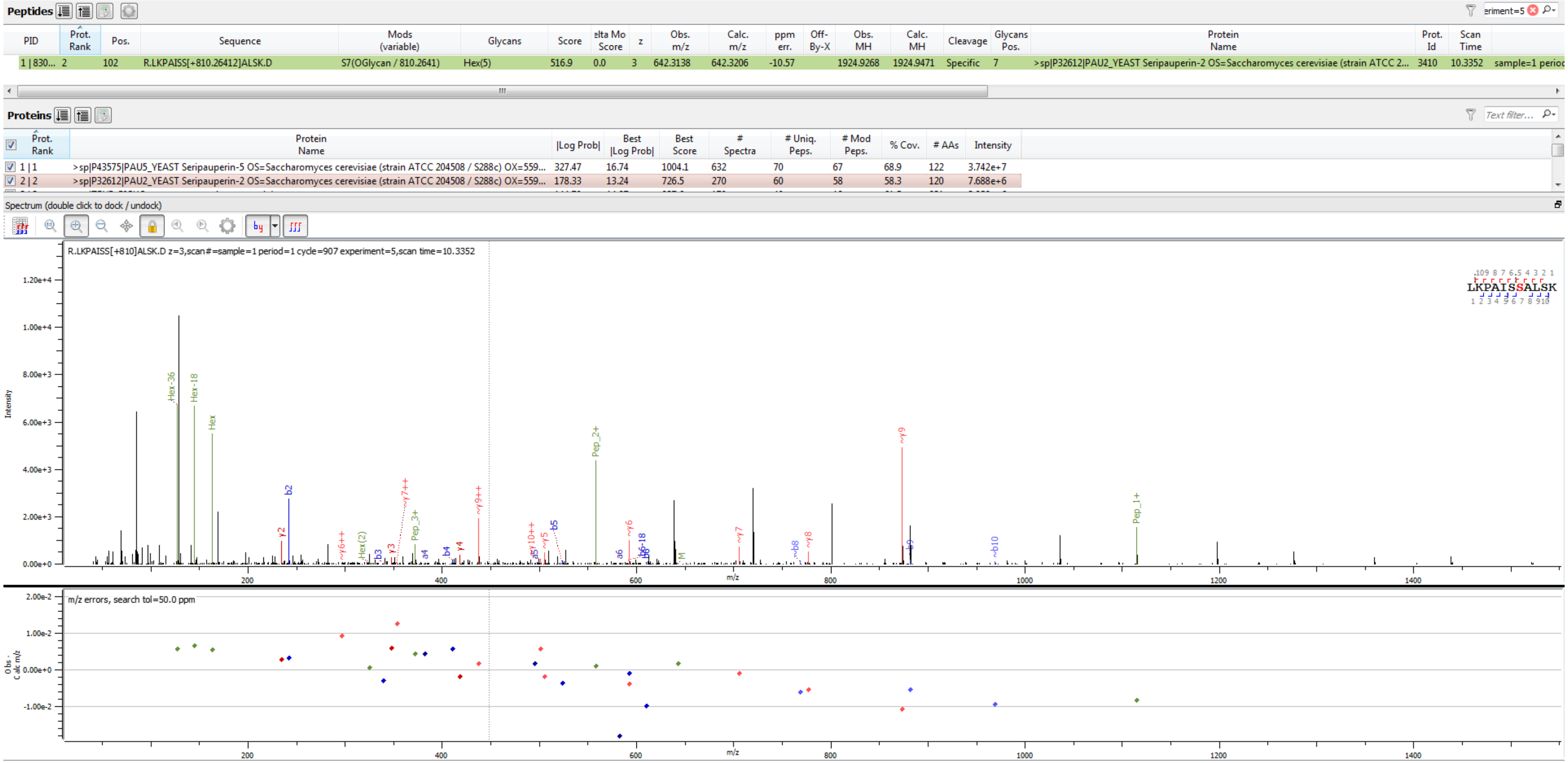


Sample S1 R.LKPAISS[+810.264]ALSK.D

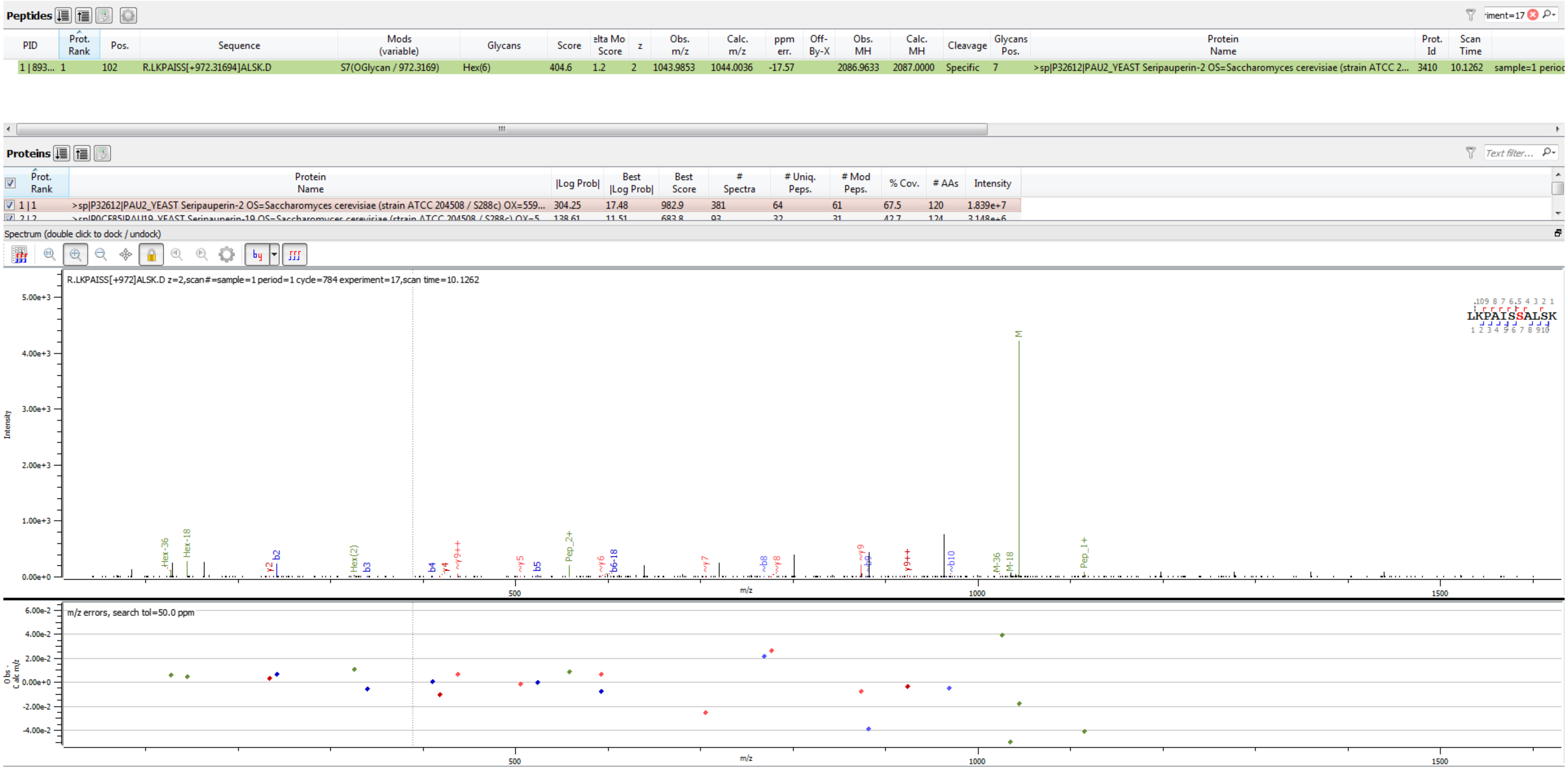


Sample S4 R.LKPAISS[+972.317]ALSK.D

**Peptides**

| PID | Prot. Rank | Pos. | Sequence | Mods (variable) | Glycans | Score | elta Mo Score | z | Obs. m/z | Calc. m/z | ppm err. | Off-By-X | Obs. MH | Calc. MH | Cleavage | Glycans Pos. | Protein Name | Prot. Id | Scan Time | |
|---|---|---|---|---|---|---|---|---|---|---|---|---|---|---|---|---|---|---|---|---|
| 1 \| 875... | 1 | 102 | R.LKPAISS[+972.31694]ALSK.D | S7(OGlycan / 972.3169) | Hex(6) | 524.9 | 1.2 | 3 | 696.3297 | 696.3382 | -12.13 | | 2086.9746 | 2087.0000 | Specific | 7 | >sp\|P32612\|PAU2_YEAST Seripauperin-2 OS=Saccharomyces cerevisiae (strain ATCC 2... | 3410 | 9.9411 | sample=1 perioc |

**Proteins**

| Prot. Rank | Protein Name | \|Log Prob\| | Best \|Log Prob\| | Best Score | # Spectra | # Uniq. Peps. | # Mod Peps. | % Cov. | # AAs | Intensity |
|---|---|---|---|---|---|---|---|---|---|---|
| 1 \| 1 | >sp\|P32612\|PAU2_YEAST Seripauperin-2 OS=Saccharomyces cerevisiae (strain ATCC 204508 / S288c) OX=559... | 304.25 | 17.48 | 982.9 | 381 | 64 | 61 | 67.5 | 120 | 1.839e+7 |
| 2 \| 2 | >sp\|P0CE85\|PAU19_YEAST Seripauperin-19 OS=Saccharomyces cerevisiae (strain ATCC 204508 / S288c) OX=5... | 138.61 | 11.51 | 683.8 | 93 | 32 | 31 | 42.7 | 124 | 3.148e+6 |

Spectrum (double click to dock / undock)

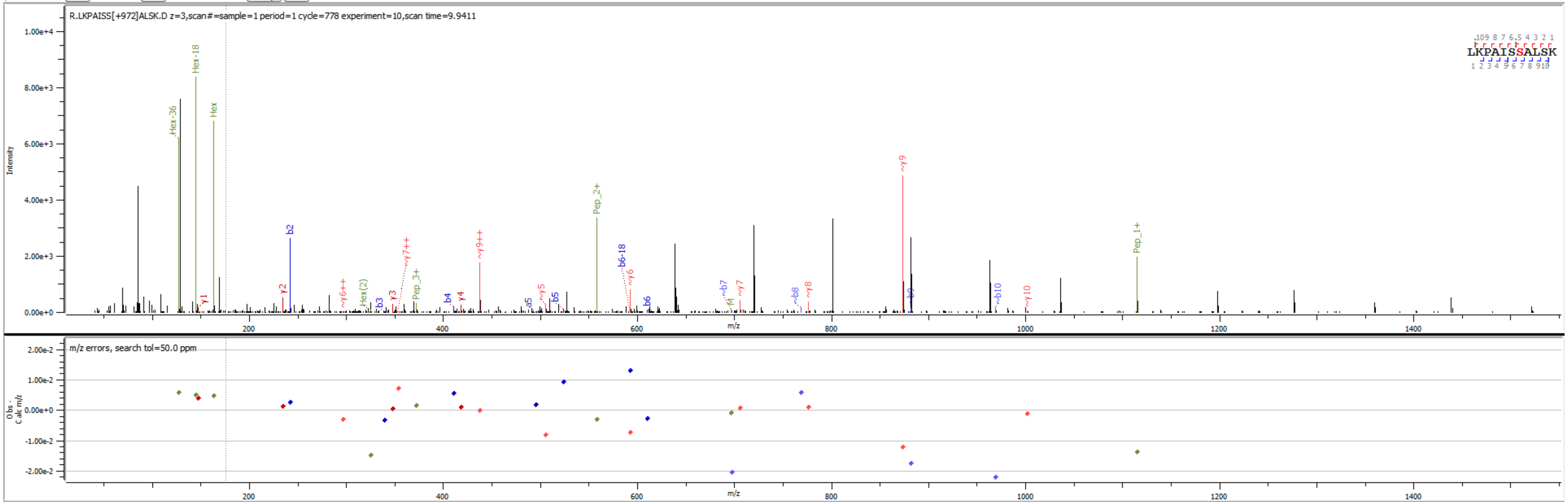


Sample S4 R.LKPAISS[+972.317]ALSK.D

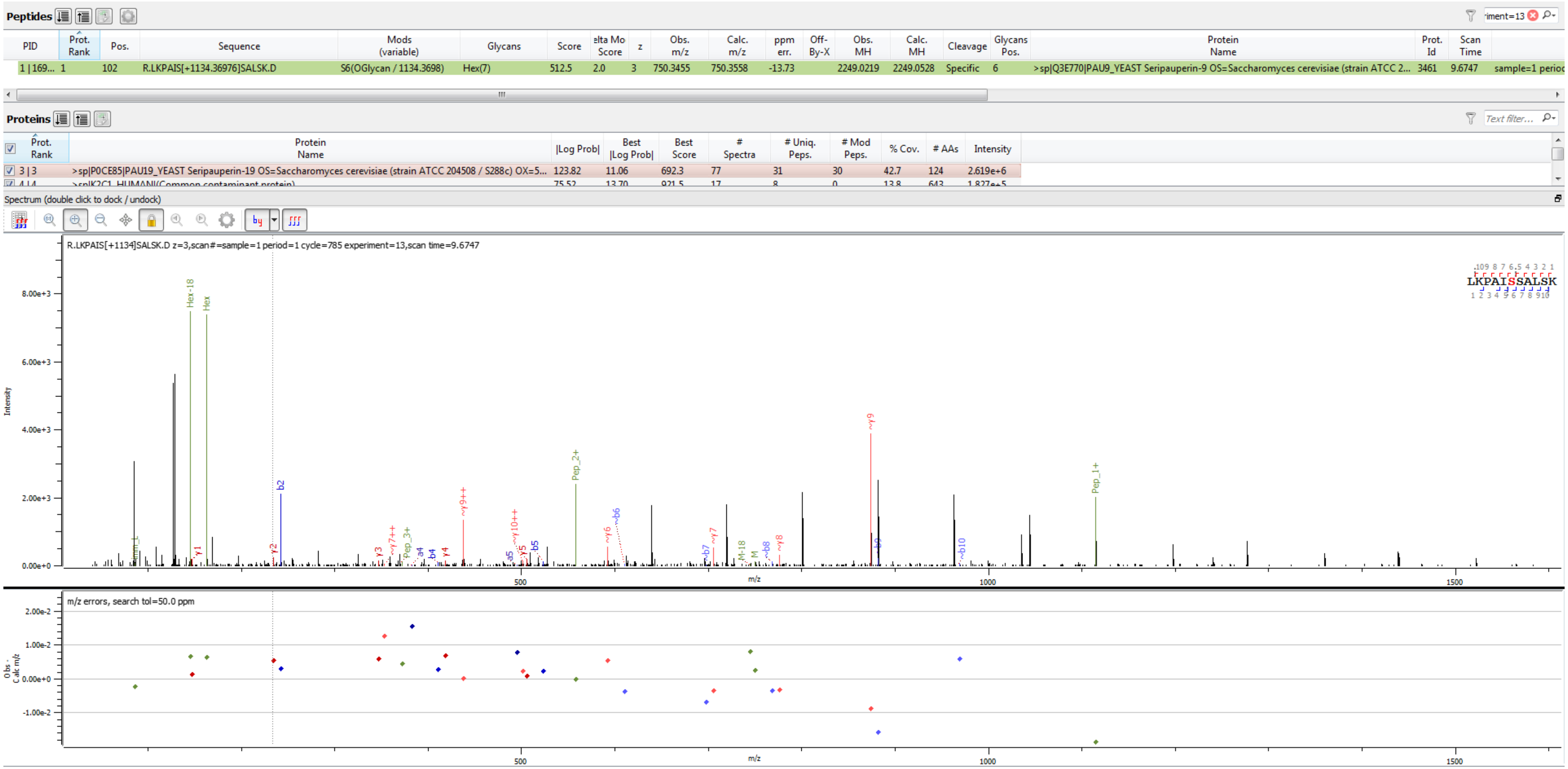


Sample S3 R.LKPAIS[+1134.370]SALSK.D

Peptide List (double click to dock / undock)

**Peptides**

| PID | Prot. Rank | Pos. | Sequence | Mods (variable) | Glycans | Score | elta Mo Score | z | Obs. m/z | Calc. m/z | ppm err. | Off-By-X | Obs. MH | Calc. MH | Cleavage | Glycans Pos. | Protein Name | Prot. Id | Scan Time | |
|---|---|---|---|---|---|---|---|---|---|---|---|---|---|---|---|---|---|---|---|---|
| 1 \| 161... | 2 | 102 | R.LKPAIS[+1296.42259]SALSK.D | S6(OGlycan / 1296.4226) | Hex(8) | 461.3 | 2.0 | 3 | 804.3581 | 804.3734 | -19.06 | | 2411.0596 | 2411.1056 | Specific | 6 | >sp\|P32612\|PAU2_YEAST Seripauperin-2 OS=Saccharomyces cerevisiae (strain ATCC 2... | 3410 | 9.7348 | sample=1 perioc |

**Proteins**

| Prot. Rank | Protein Name | \|Log Prob\| | Best \|Log Prob\| | Best Score | # Spectra | # Uniq. Peps. | # Mod Peps. | % Cov. | # AAs | Intensity |
|---|---|---|---|---|---|---|---|---|---|---|
| 1 \| 1 | >sp\|P43575\|PAU5_YEAST Seripauperin-5 OS=Saccharomyces cerevisiae (strain ATCC 204508 / S288c) OX=559... | 327.47 | 16.74 | 1004.1 | 632 | 70 | 67 | 68.9 | 122 | 3.742e+7 |
| 2 \| 2 | >sp\|P32612\|PAU2_YEAST Seripauperin-2 OS=Saccharomyces cerevisiae (strain ATCC 204508 / S288c) OX=559... | 178.33 | 13.24 | 726.5 | 270 | 60 | 58 | 58.3 | 120 | 7.688e+6 |

Spectrum (double click to dock / undock)

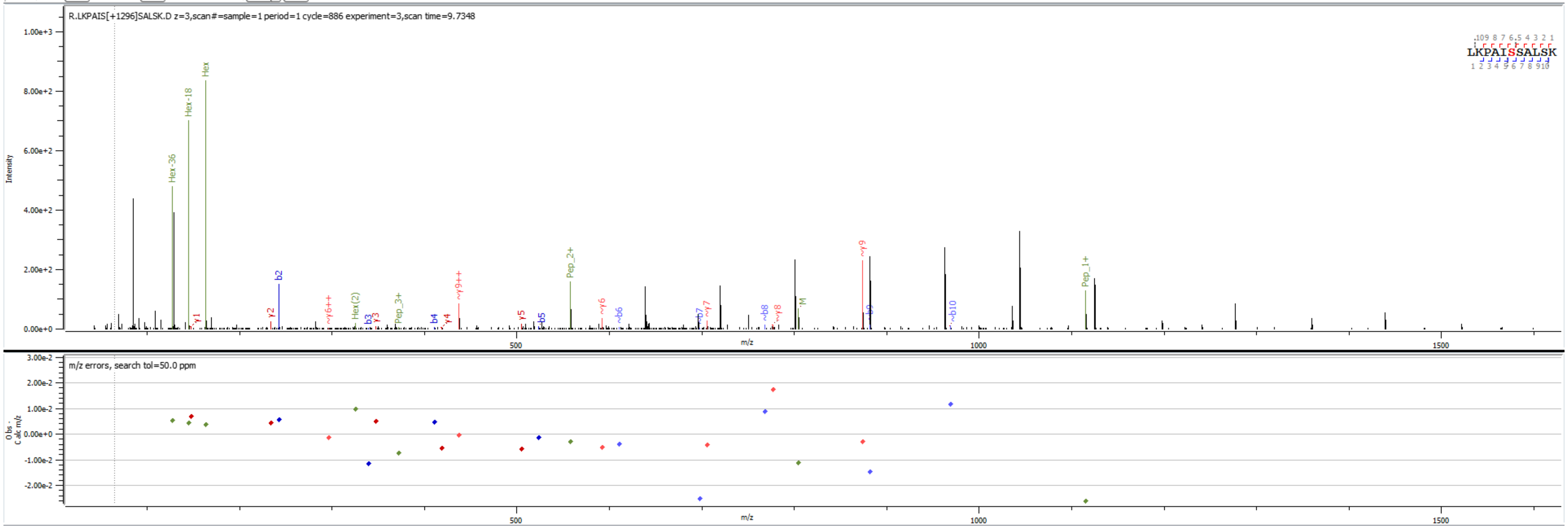


Sample S1 R.LKPAIS[+1296.423]SALSK.D

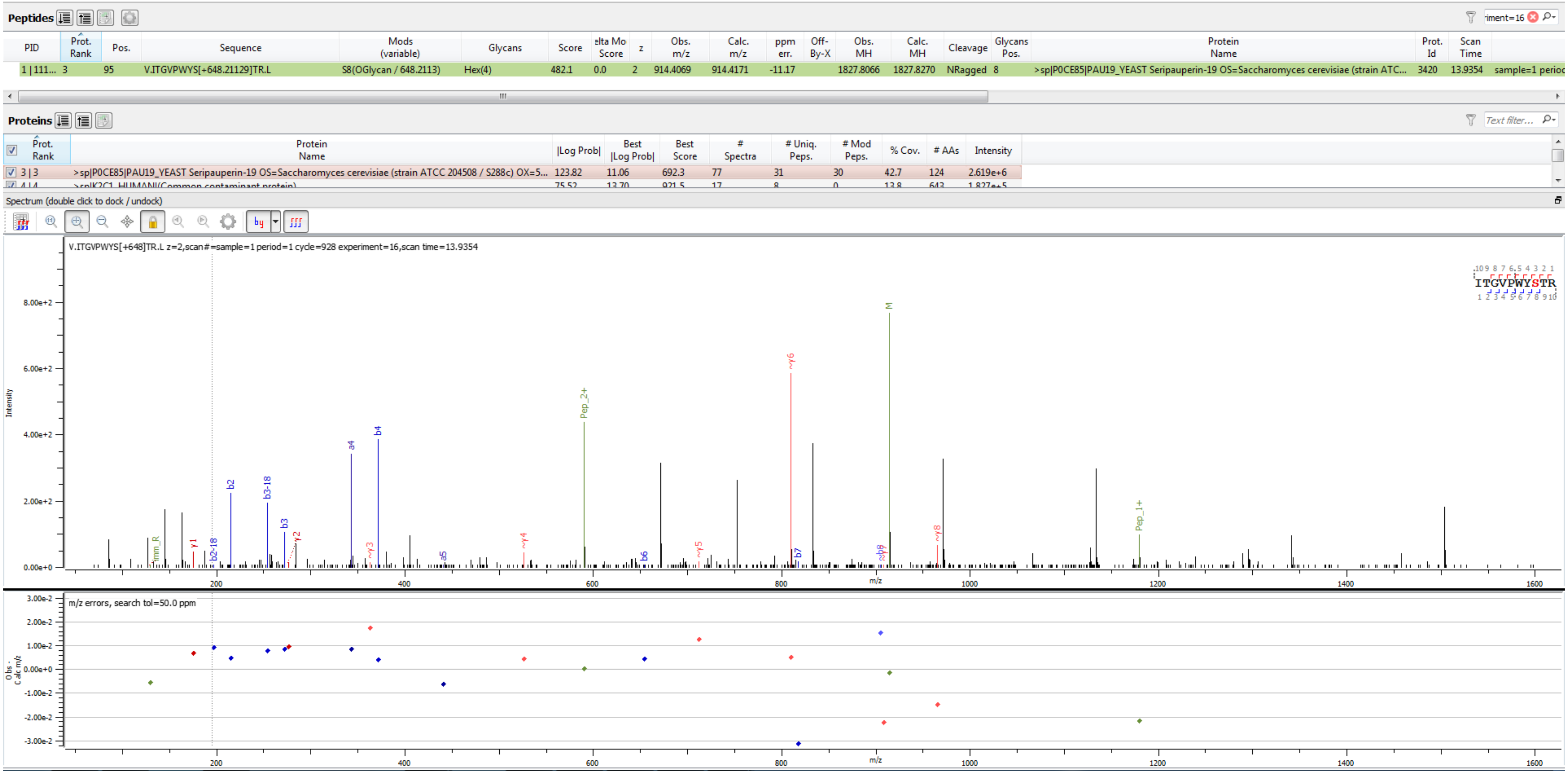


Sample S3 V.ITGVPWYS[+648.211]TR.L

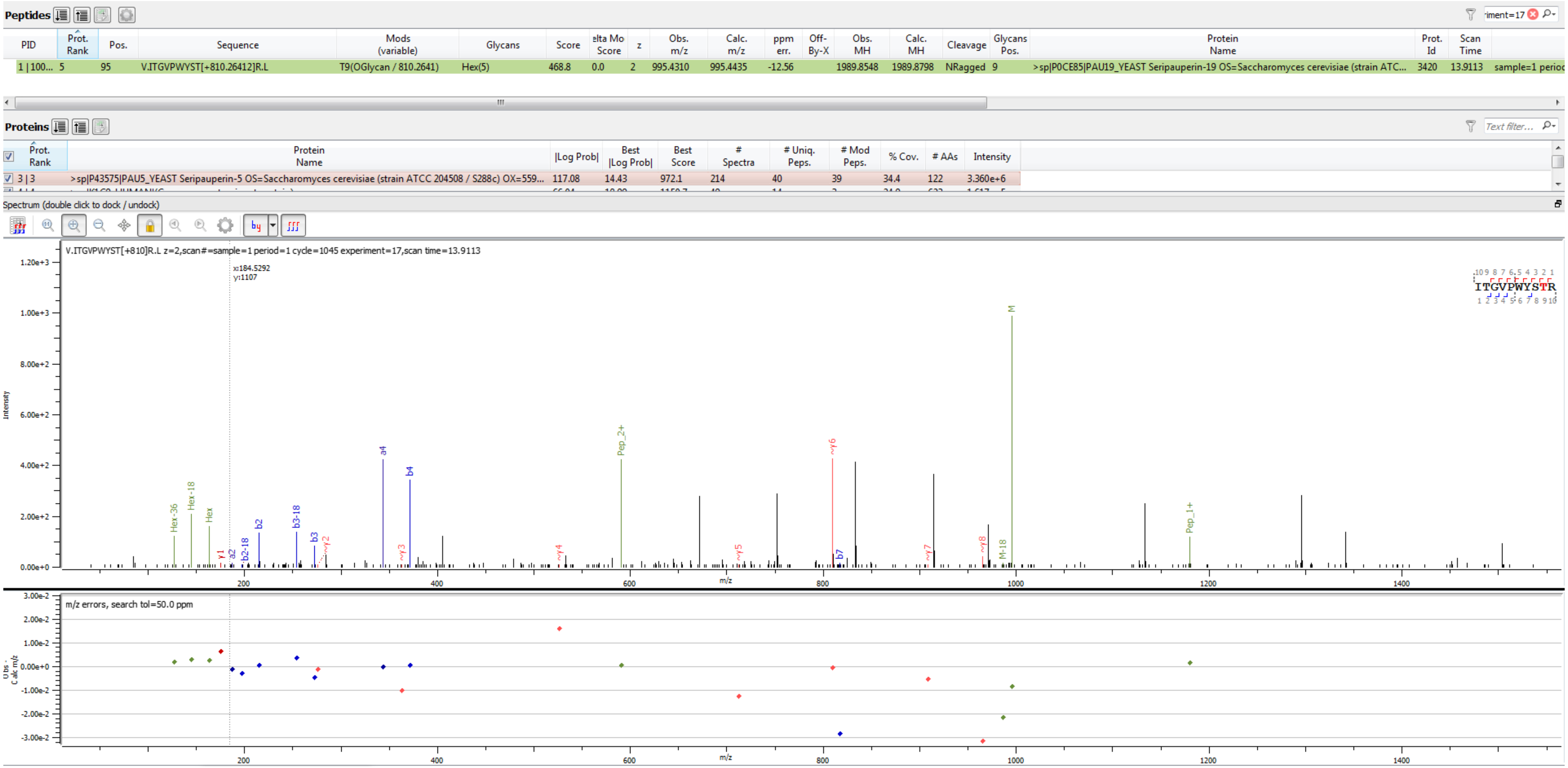


Sample S2 V.ITGVPWYST[+810.264]R.L

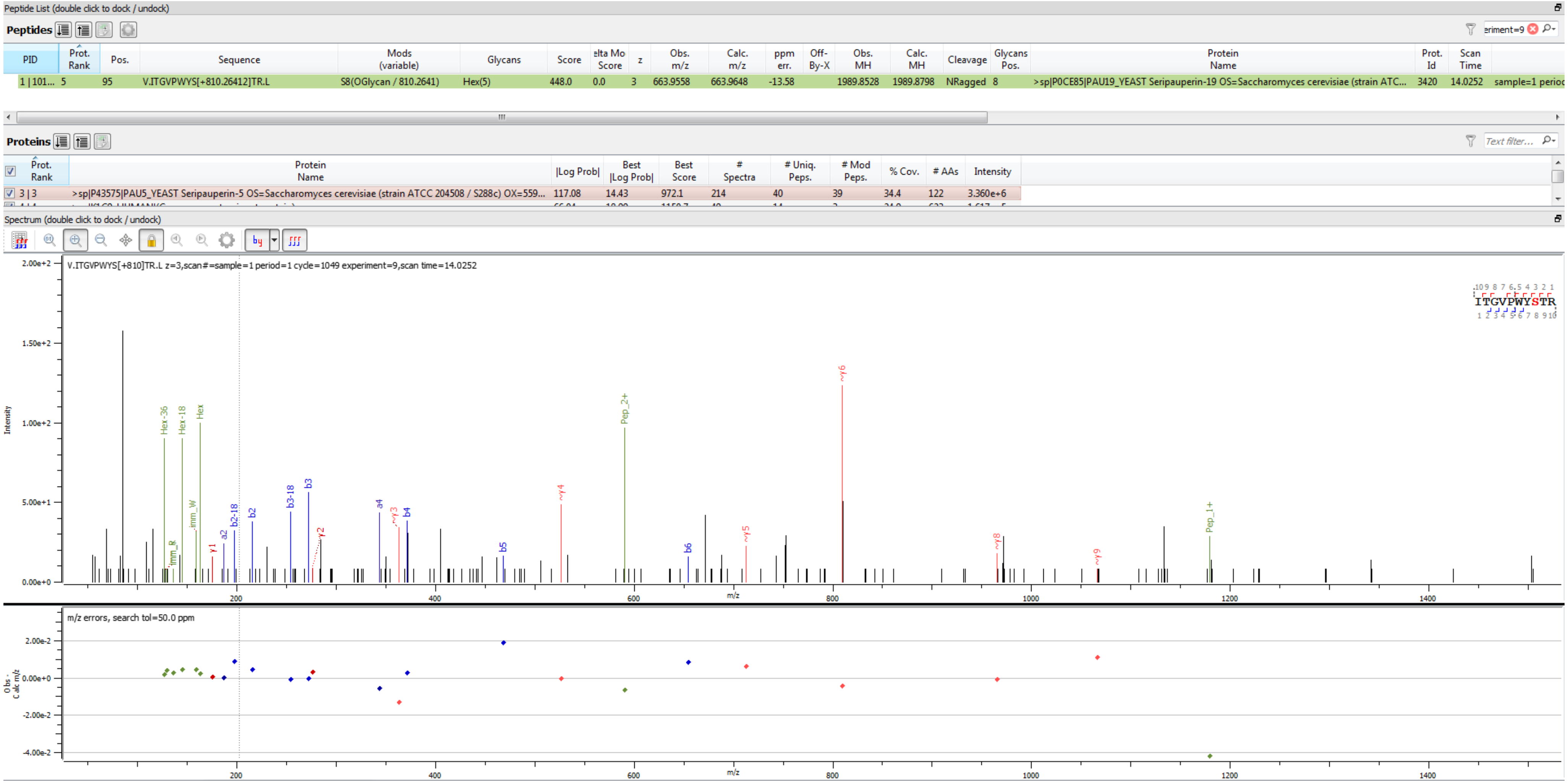


Sample S2 V.ITGVPWYS[+810.264]TR.L

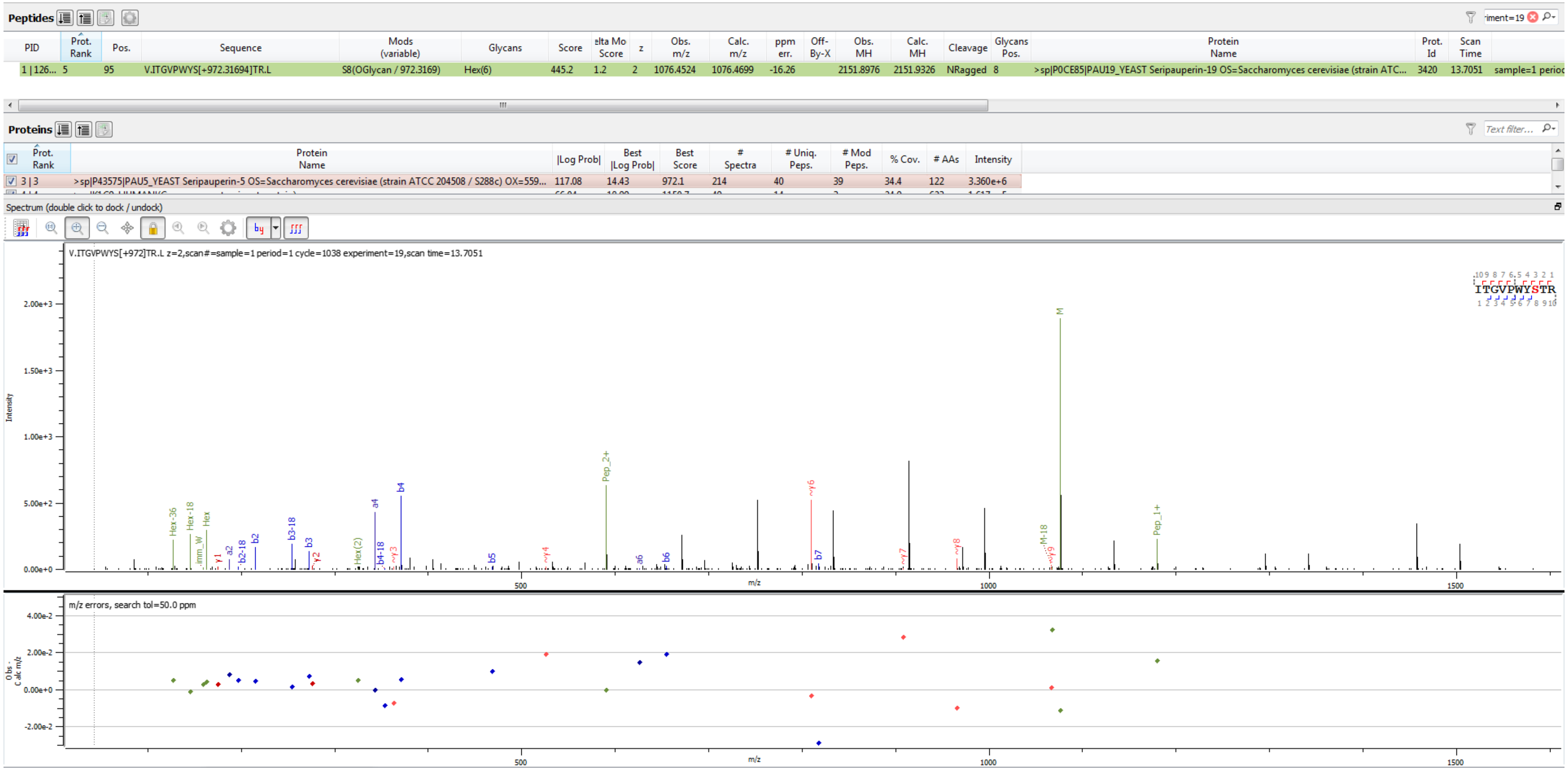


Sample S2 V.ITGVPWYS[+972.317]TR.L

**Peptides**

| PID | Prot. Rank | Pos. | Sequence | Mods (variable) | Glycans | Score | elta Mo Score | z | Obs. m/z | Calc. m/z | ppm err. | Off-By-X | Obs. MH | Calc. MH | Cleavage | Glycans Pos. | Protein Name | Prot. Id | Scan Time | |
|---|---|---|---|---|---|---|---|---|---|---|---|---|---|---|---|---|---|---|---|---|
| 1 \| 126... | 5 | 95 | V.ITGVPWYS[+972.31694]TR.L | S8(OGlycan / 972.3169) | Hex(6) | 538.7 | 1.2 | 3 | 717.9694 | 717.9824 | -18.08 | | 2151.8937 | 2151.9326 | NRagged | 8 | >sp\|P0CE85\|PAU19_YEAST Seripauperin-19 OS=Saccharomyces cerevisiae (strain ATC... | 3420 | 13.7270 | sample=1 perioc |

**Proteins**

| | Prot. Rank | Protein Name | \|Log Prob\| | Best \|Log Prob\| | Best Score | # Spectra | # Uniq. Peps. | # Mod Peps. | % Cov. | # AAs | Intensity |
|---|---|---|---|---|---|---|---|---|---|---|---|
| ☑ | 3 \| 3 | >sp\|P43575\|PAU5_YEAST Seripauperin-5 OS=Saccharomyces cerevisiae (strain ATCC 204508 / S288c) OX=559... | 117.08 | 14.43 | 972.1 | 214 | 40 | 39 | 34.4 | 122 | 3.360e+6 |

Spectrum (double click to dock / undock)

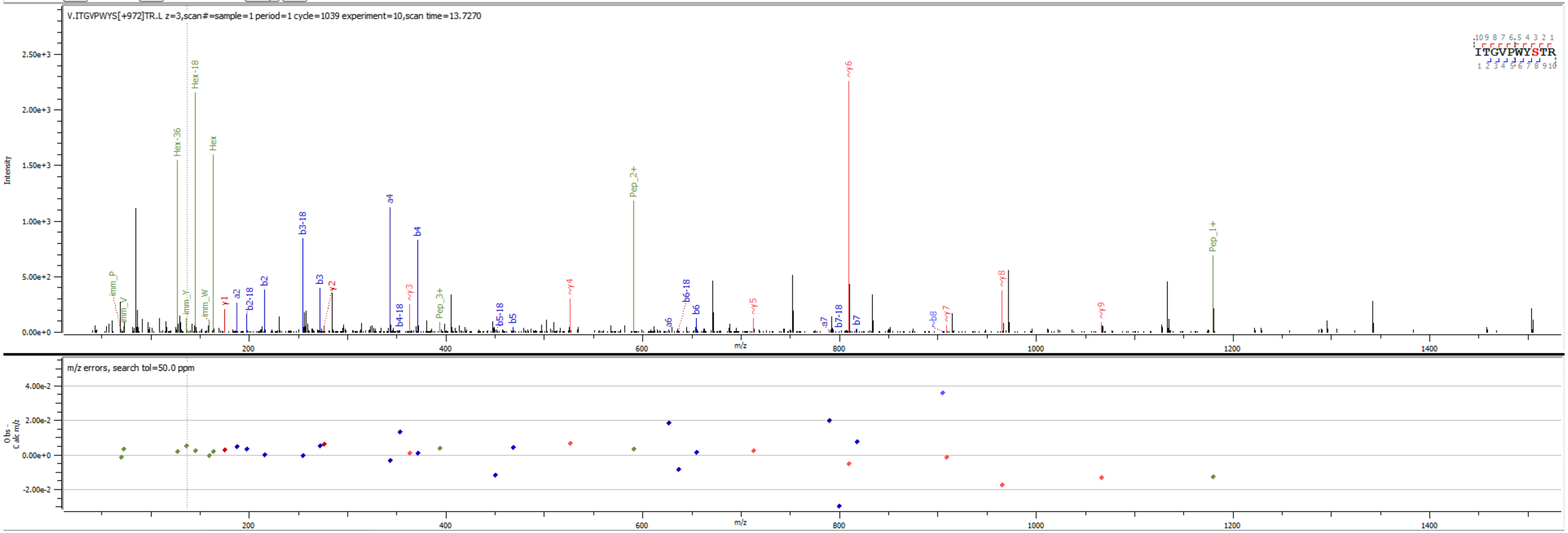


Sample S2 V.ITGVPWYS[+972.317]TR.L

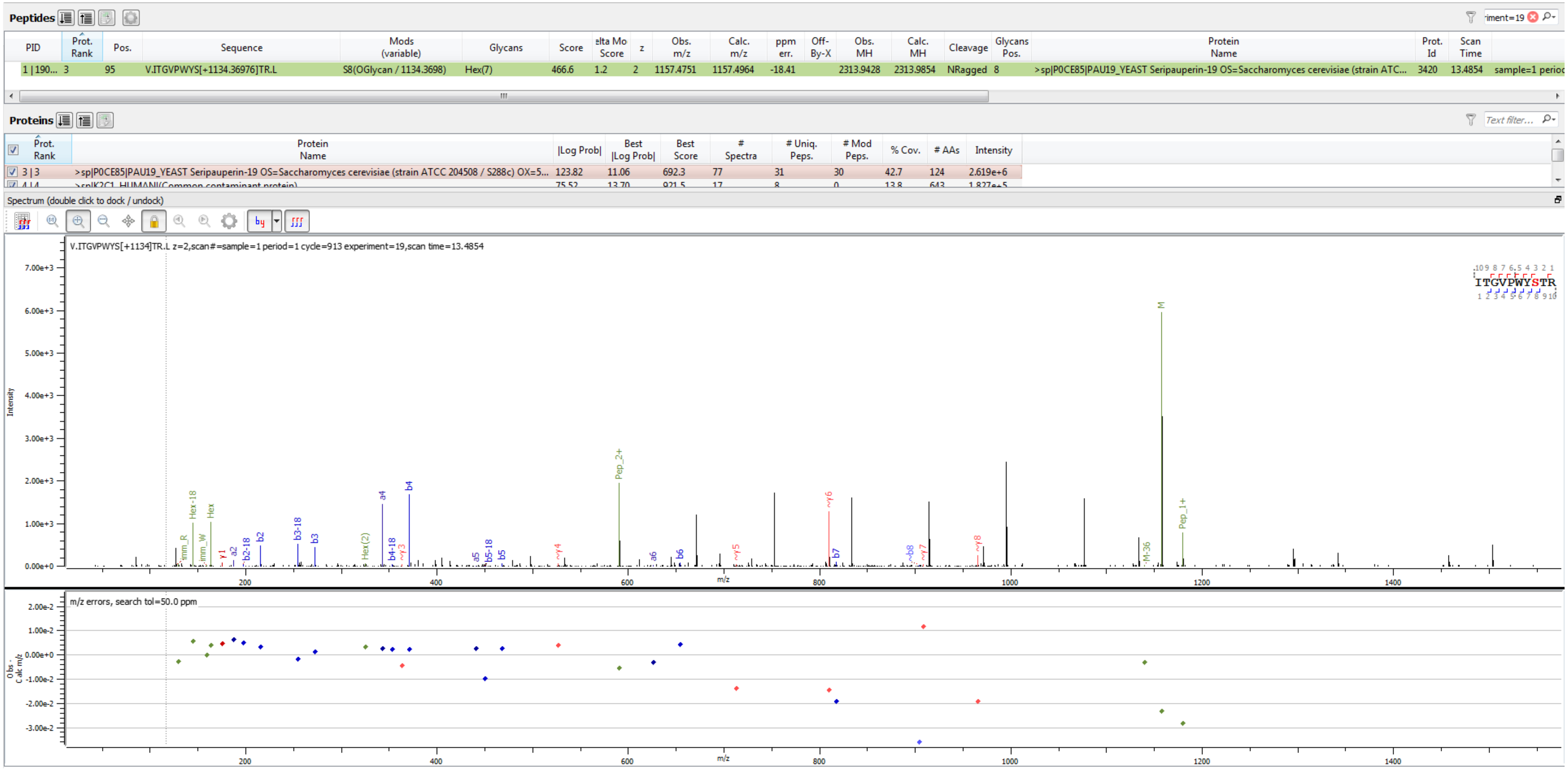


Sample S3 V.ITGVPWYS[+1134.370]TR.L

**Peptides**

| PID | Prot. Rank | Pos. | Sequence | Mods (variable) | Glycans | Score | elta Mo Score | z | Obs. m/z | Calc. m/z | ppm err. | Off-By-X | Obs. MH | Calc. MH | Cleavage | Glycans Pos. | Protein Name | Prot. Id | Scan Time | |
|---|---|---|---|---|---|---|---|---|---|---|---|---|---|---|---|---|---|---|---|---|
| 1 \| 150... | 5 | 95 | V.ITGVPWYS[+1134.36976]TR.L | S8(OGlycan / 1134.3698) | Hex(7) | 492.5 | 1.2 | 3 | 771.9868 | 772.0000 | -17.08 | | 2313.9459 | 2313.9854 | NRagged | 8 | >sp\|P0CE85\|PAU19_YEAST Seripauperin-19 OS=Saccharomyces cerevisiae (strain ATC... | 3420 | 13.5215 | sample=1 perioc |

**Proteins**

| Prot. Rank | Protein Name | \|Log Prob\| | Best \|Log Prob\| | Best Score | # Spectra | # Uniq. Peps. | # Mod Peps. | % Cov. | # AAs | Intensity |
|---|---|---|---|---|---|---|---|---|---|---|
| 3 \| 3 | >sp\|P43575\|PAU5_YEAST Seripauperin-5 OS=Saccharomyces cerevisiae (strain ATCC 204508 / S288c) OX=559... | 117.08 | 14.43 | 972.1 | 214 | 40 | 39 | 34.4 | 122 | 3.360e+6 |

Spectrum (double click to dock / undock)

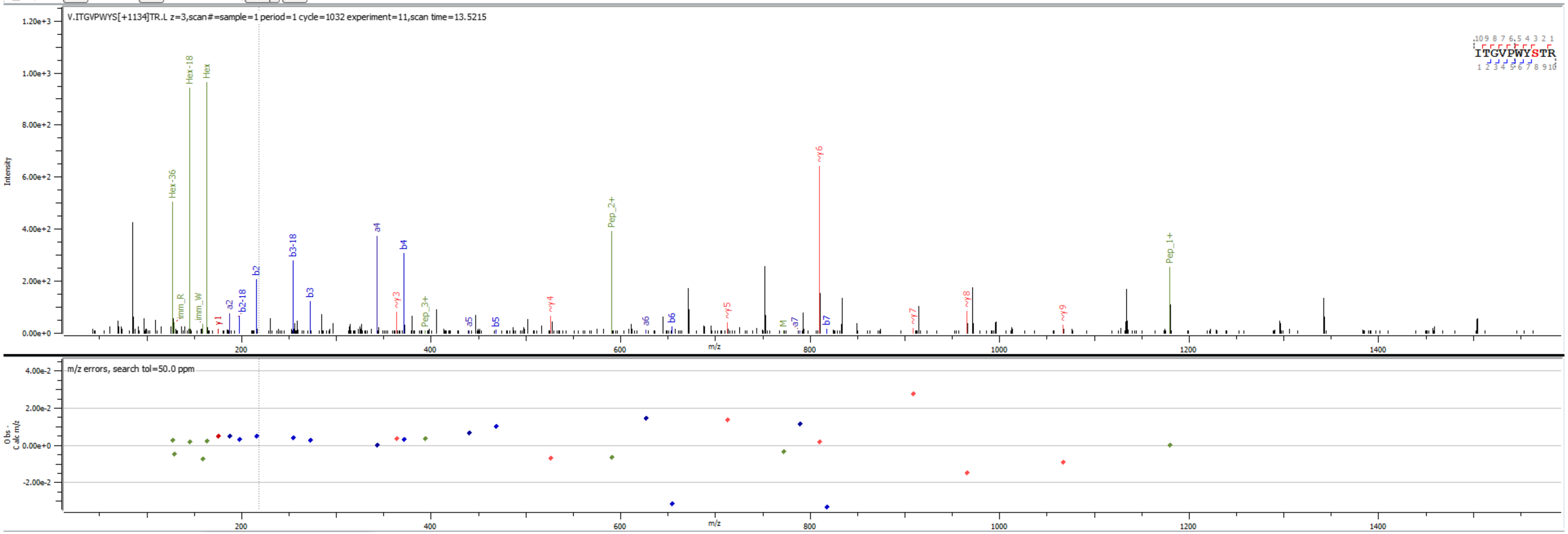


Sample S2 V.ITGVPWYS[+1134.370]TR.L

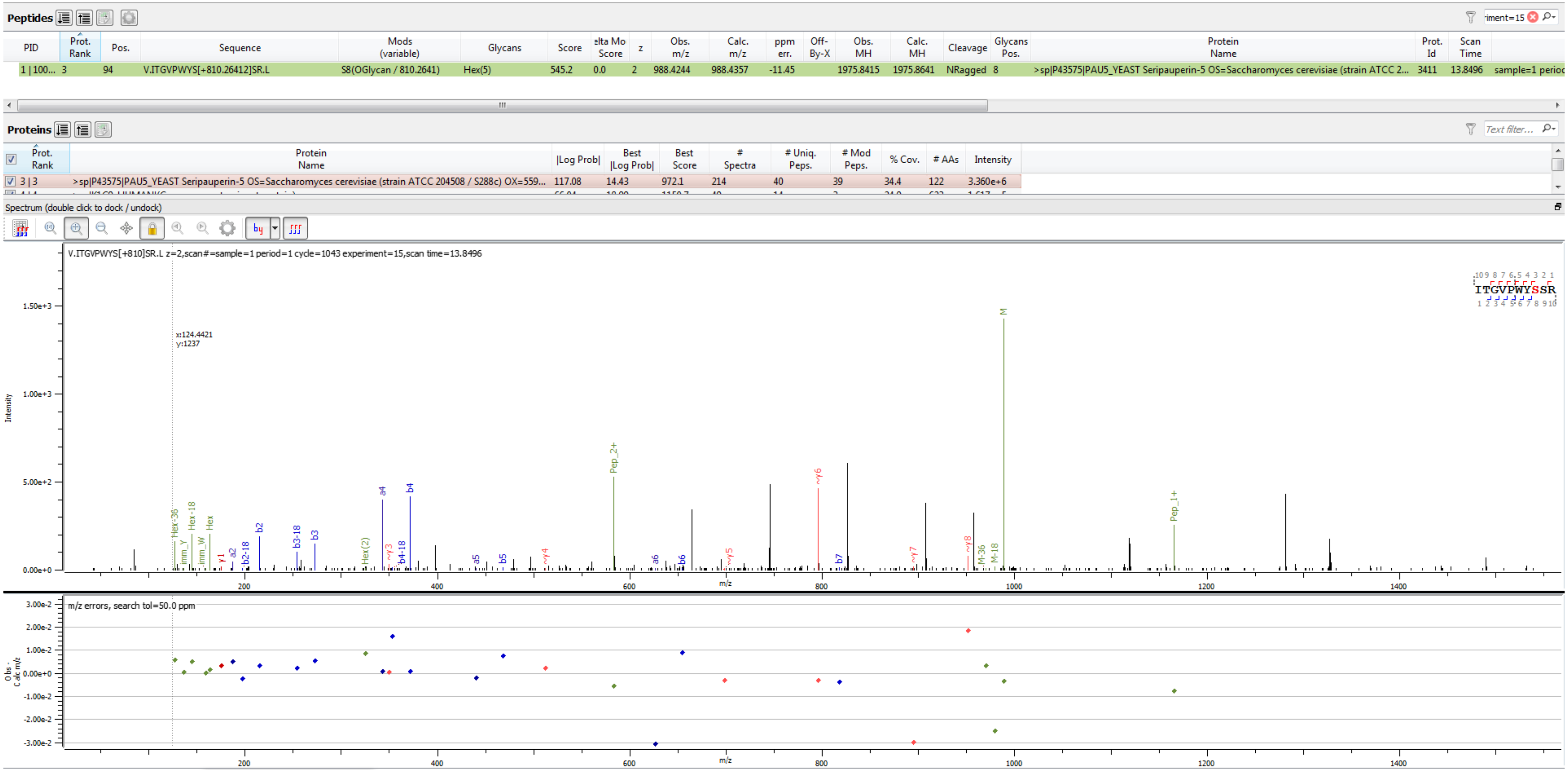


Sample S2 V.ITGVPWYS[+810.264]SR.L

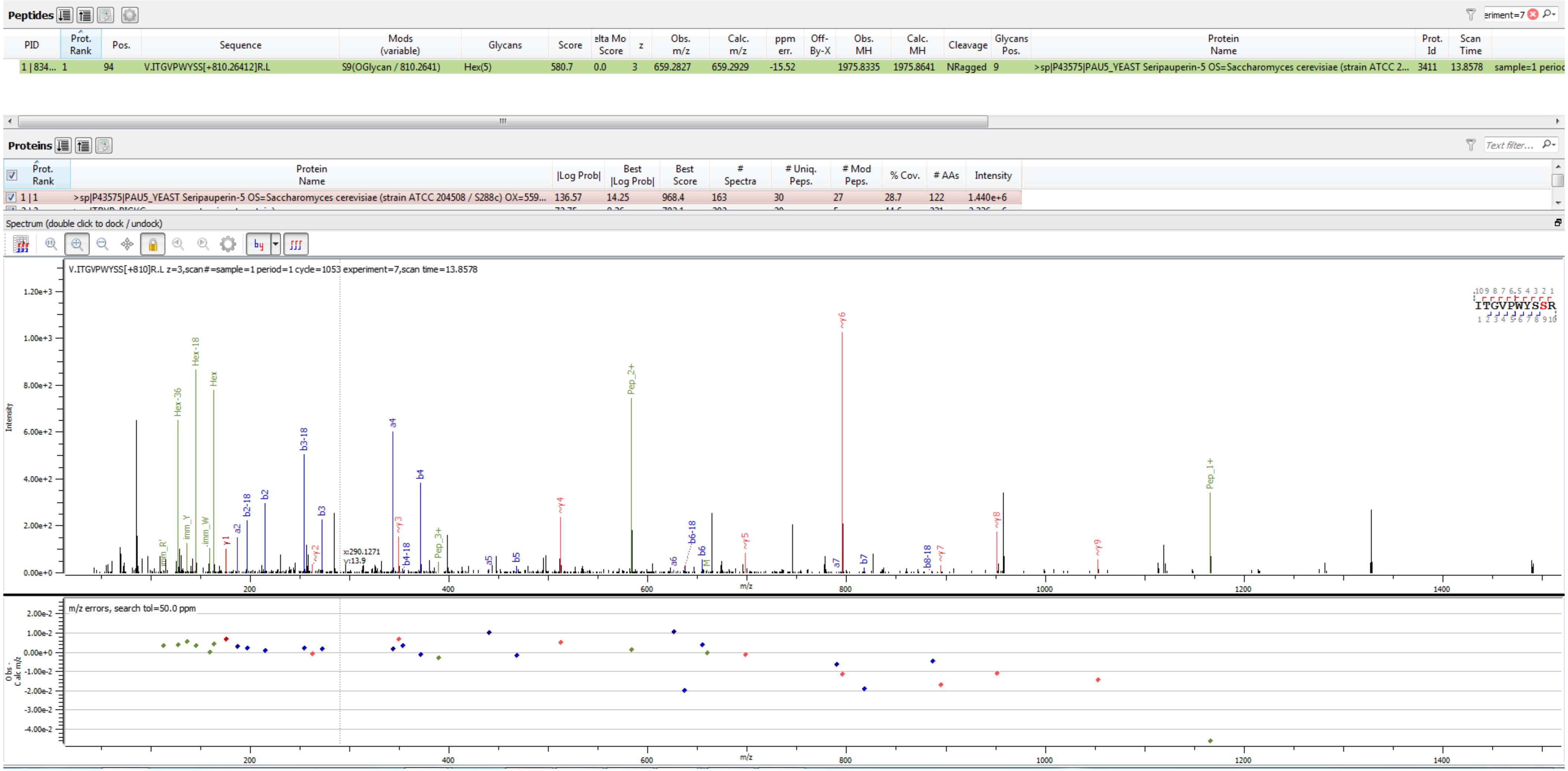


Sample S5 V.ITGVPWYSS[+810.264]R.L

**Peptides**

<table>
<tr><th>PID</th><th>Prot. Rank</th><th>Pos.</th><th>Sequence</th><th>Mods (variable)</th><th>Glycans</th><th>Score</th><th>elta Mo Score</th><th>z</th><th>Obs. m/z</th><th>Calc. m/z</th><th>ppm err.</th><th>Off-By-X</th><th>Obs. MH</th><th>Calc. MH</th><th>Cleavage</th><th>Glycans Pos.</th><th>Protein Name</th><th>Prot. Id</th><th>Scan Time</th><th></th></tr>
<tr><td>1 | 126...</td><td>3</td><td>94</td><td>V.ITGVPWYS[+972.31694]SR.L</td><td>S8(OGlycan / 972.3169)</td><td>Hex(6)</td><td>481.7</td><td>1.2</td><td>2</td><td>1069.4505</td><td>1069.4621</td><td>-10.91</td><td></td><td>2137.8936</td><td>2137.9170</td><td>NRagged</td><td>8</td><td>>sp|P43575|PAU5_YEAST Seripauperin-5 OS=Saccharomyces cerevisiae (strain ATCC 2...</td><td>3411</td><td>13.5588</td><td>sample=1 perioc</td></tr>
</table>

**Proteins**

<table>
<tr><th></th><th>Prot. Rank</th><th>Protein Name</th><th>|Log Prob|</th><th>Best |Log Prob|</th><th>Best Score</th><th># Spectra</th><th># Uniq. Peps.</th><th># Mod Peps.</th><th>% Cov.</th><th># AAs</th><th>Intensity</th></tr>
<tr><td>☑</td><td>3 | 3</td><td>>sp|P43575|PAU5_YEAST Seripauperin-5 OS=Saccharomyces cerevisiae (strain ATCC 204508 / S288c) OX=559...</td><td>117.08</td><td>14.43</td><td>972.1</td><td>214</td><td>40</td><td>39</td><td>34.4</td><td>122</td><td>3.360e+6</td></tr>
</table>

Spectrum (double click to dock / undock)

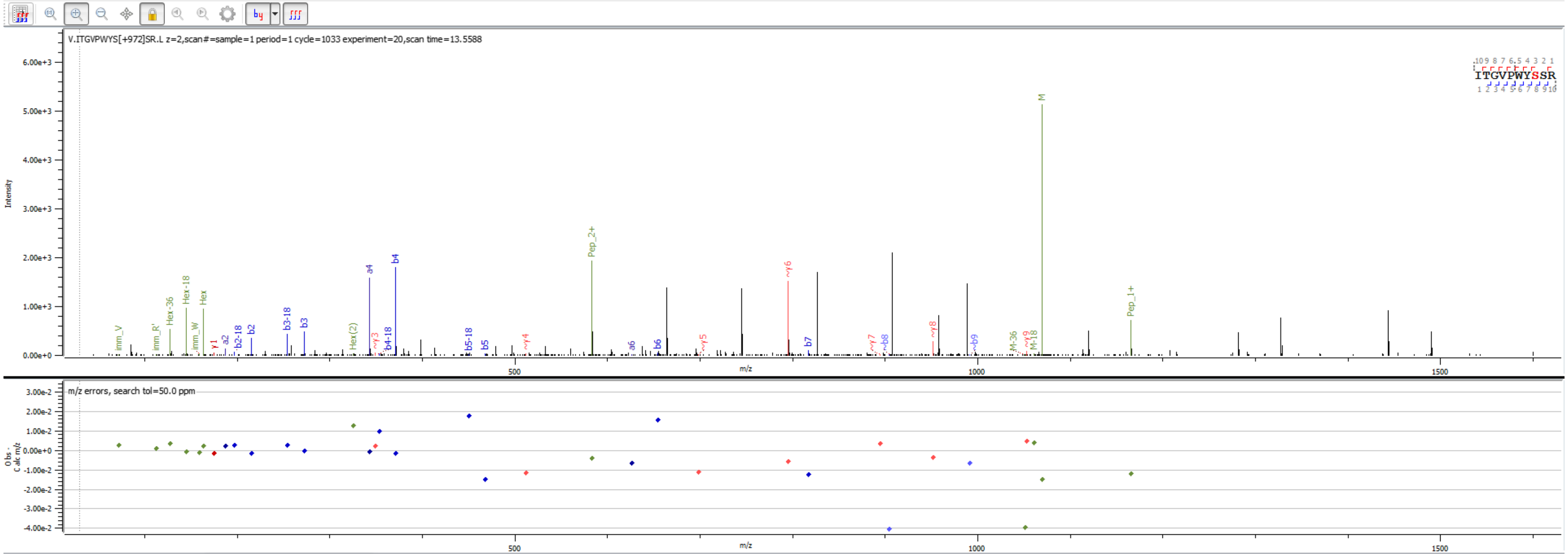


Sample S2 V.ITGVPWYS[+972.317]SR.L

**Peptides**

| PID | Prot. Rank | Pos. | Sequence | Mods (variable) | Glycans | Score | elta Mo Score | z | Obs. m/z | Calc. m/z | ppm err. | Off-By-X | Obs. MH | Calc. MH | Cleavage | Glycans Pos. | Protein Name | Prot. Id | Scan Time | |
|---|---|---|---|---|---|---|---|---|---|---|---|---|---|---|---|---|---|---|---|---|
| 1 \| 125... | 3 | 94 | V.ITGVPWYS[+972.31694]SR.L | S8(OGlycan / 972.3169) | Hex(6) | 532.3 | 1.2 | 3 | 713.3003 | 713.3105 | -14.33 | | 2137.8863 | 2137.9170 | NRagged | 8 | >sp\|P43575\|PAU5_YEAST Seripauperin-5 OS=Saccharomyces cerevisiae (strain ATCC 2... | 3411 | 13.5199 | sample=1 perioc |

**Proteins**

| Prot. Rank | Protein Name | \|Log Prob\| | Best \|Log Prob\| | Best Score | # Spectra | # Uniq. Peps. | # Mod Peps. | % Cov. | # AAs | Intensity |
|---|---|---|---|---|---|---|---|---|---|---|
| 3 \| 3 | >sp\|P43575\|PAU5_YEAST Seripauperin-5 OS=Saccharomyces cerevisiae (strain ATCC 204508 / S288c) OX=559... | 117.08 | 14.43 | 972.1 | 214 | 40 | 39 | 34.4 | 122 | 3.360e+6 |

Spectrum (double click to dock / undock)

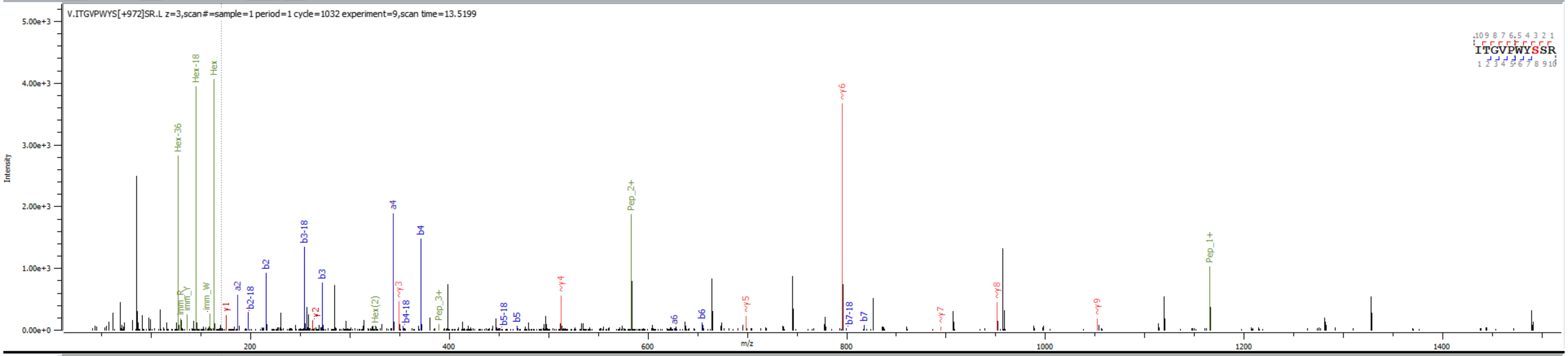


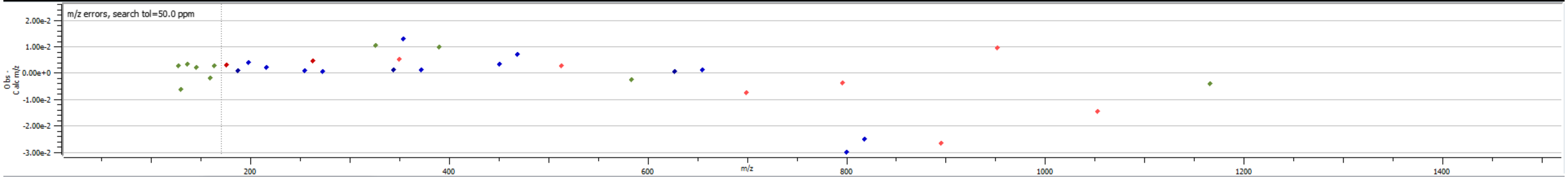


Sample S2 V.ITGVPWYS[+972.317]SR.L

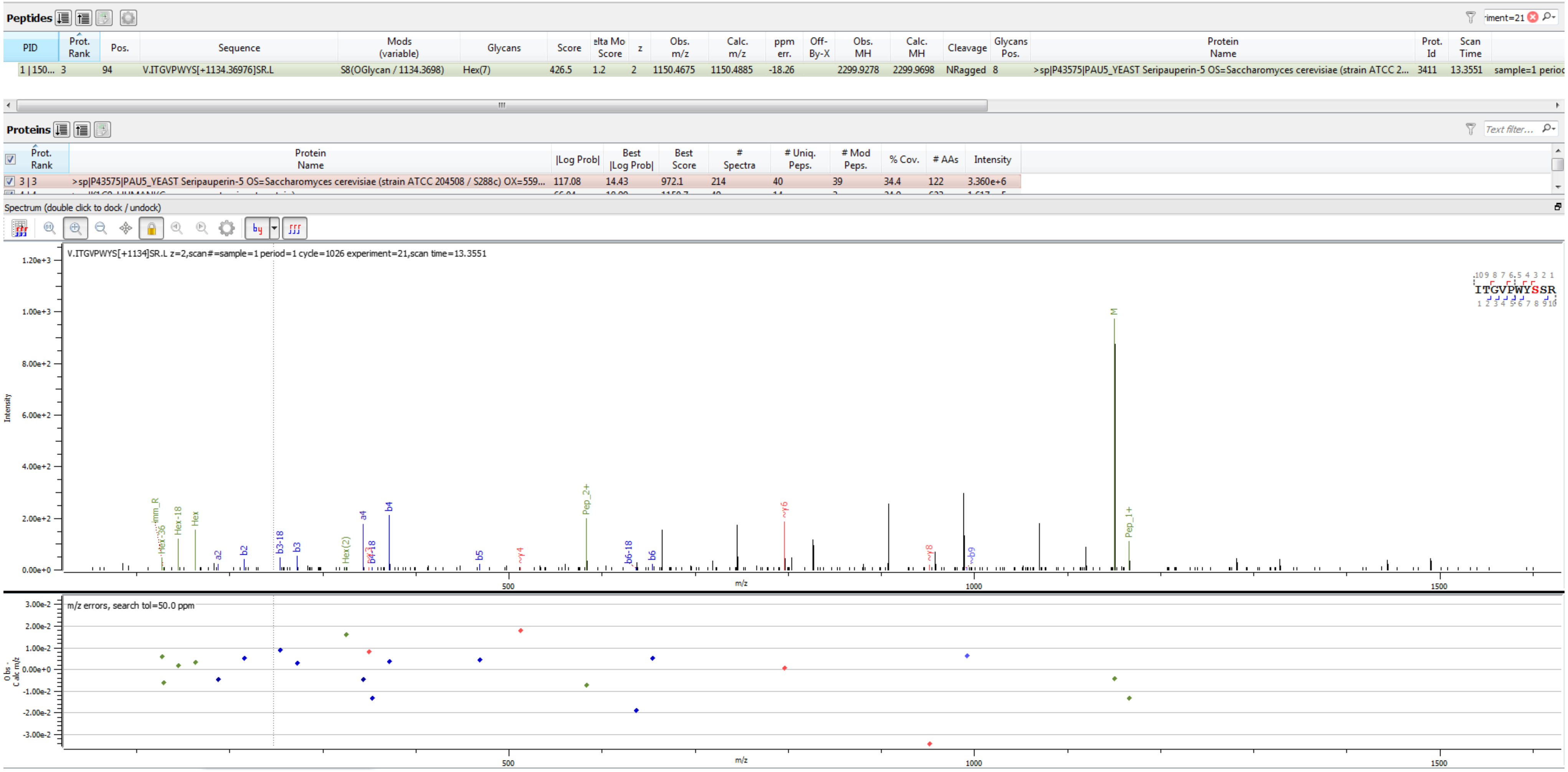


Sample S2 V.ITGVPWYS[+1134.370]SR.L

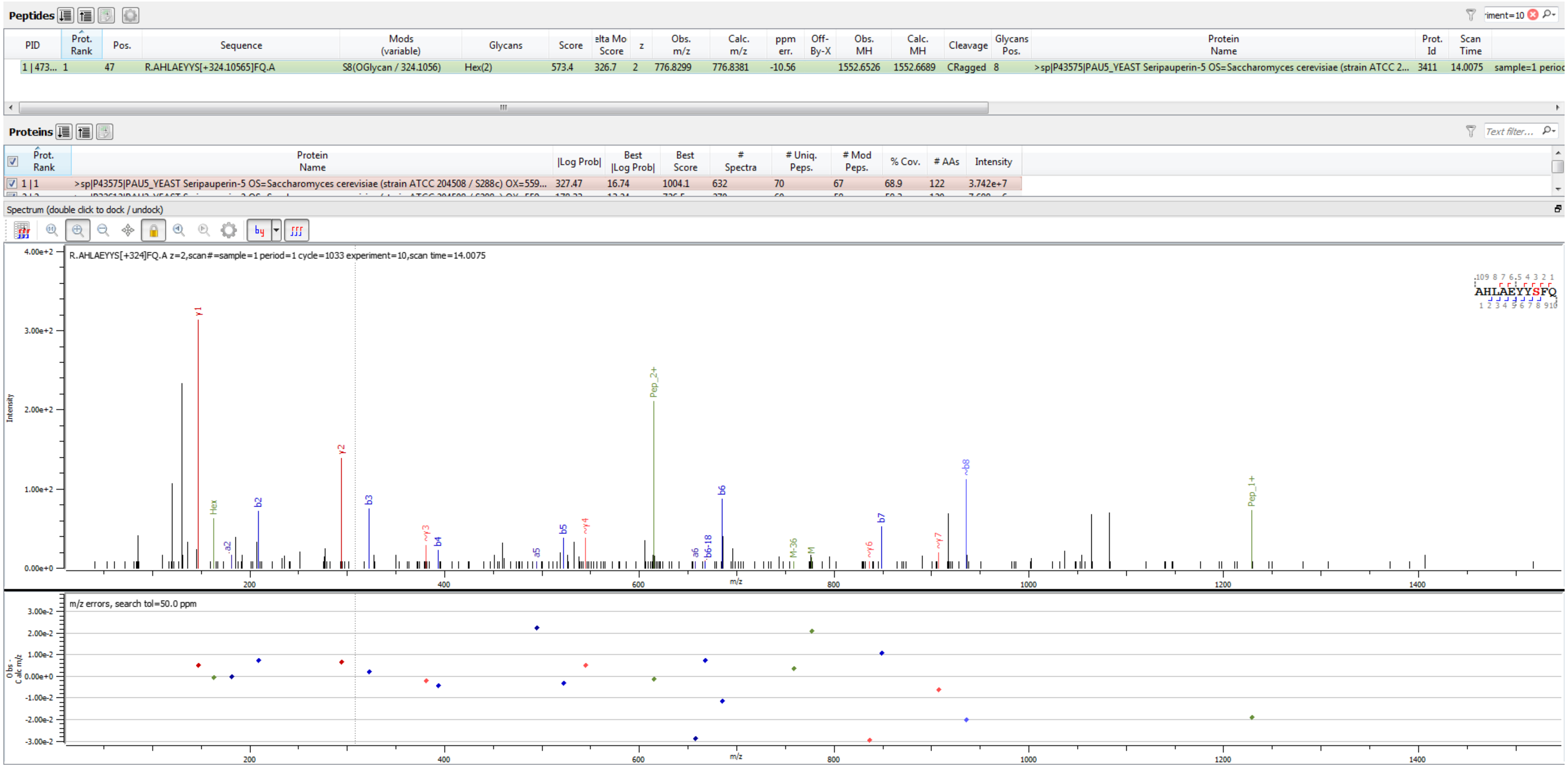


Sample S1 R.AHLAEYYS[+324.106]FQ.A

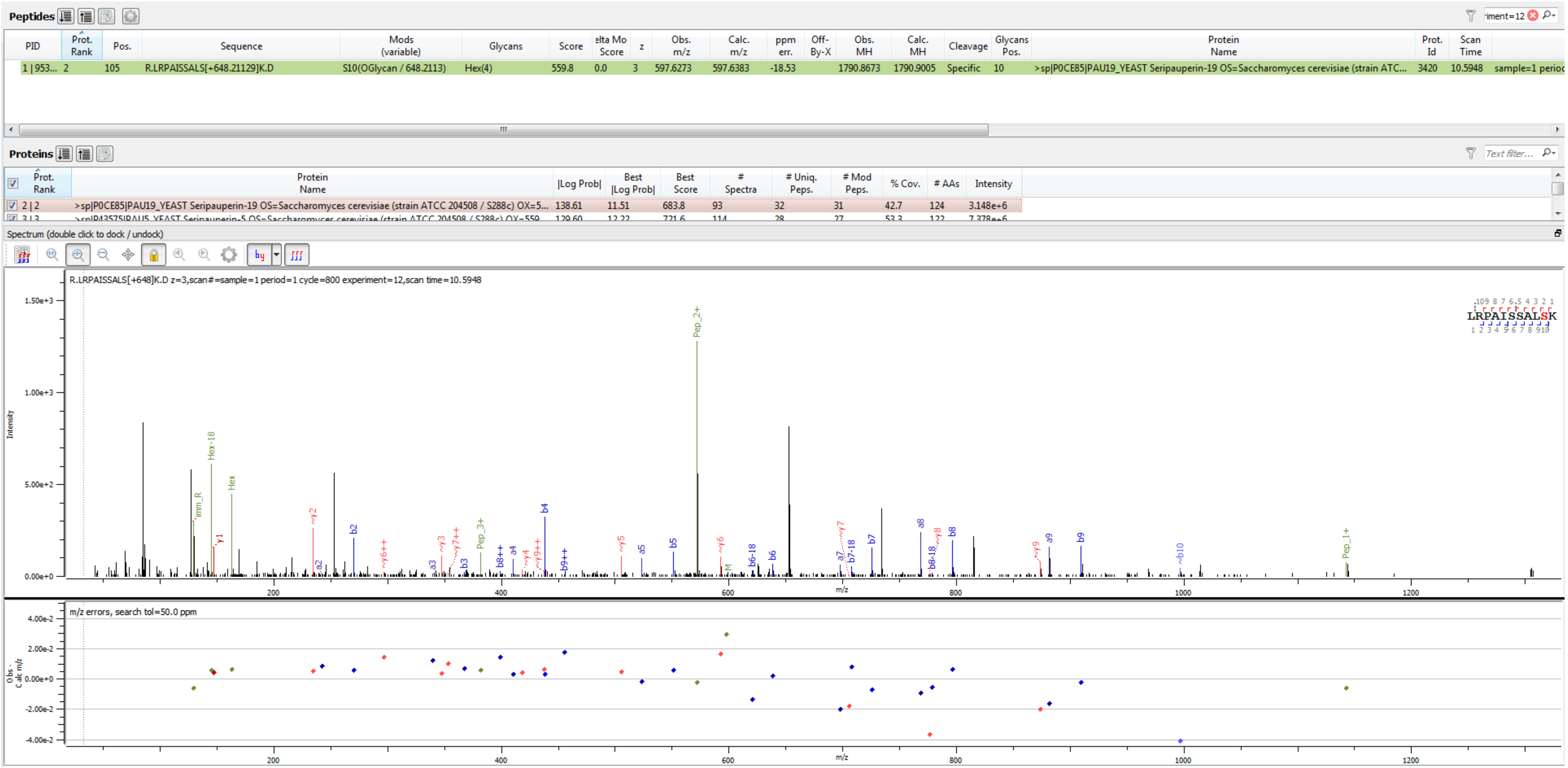


Sample S4 R.LRPAISSALS[+648.211]K.D

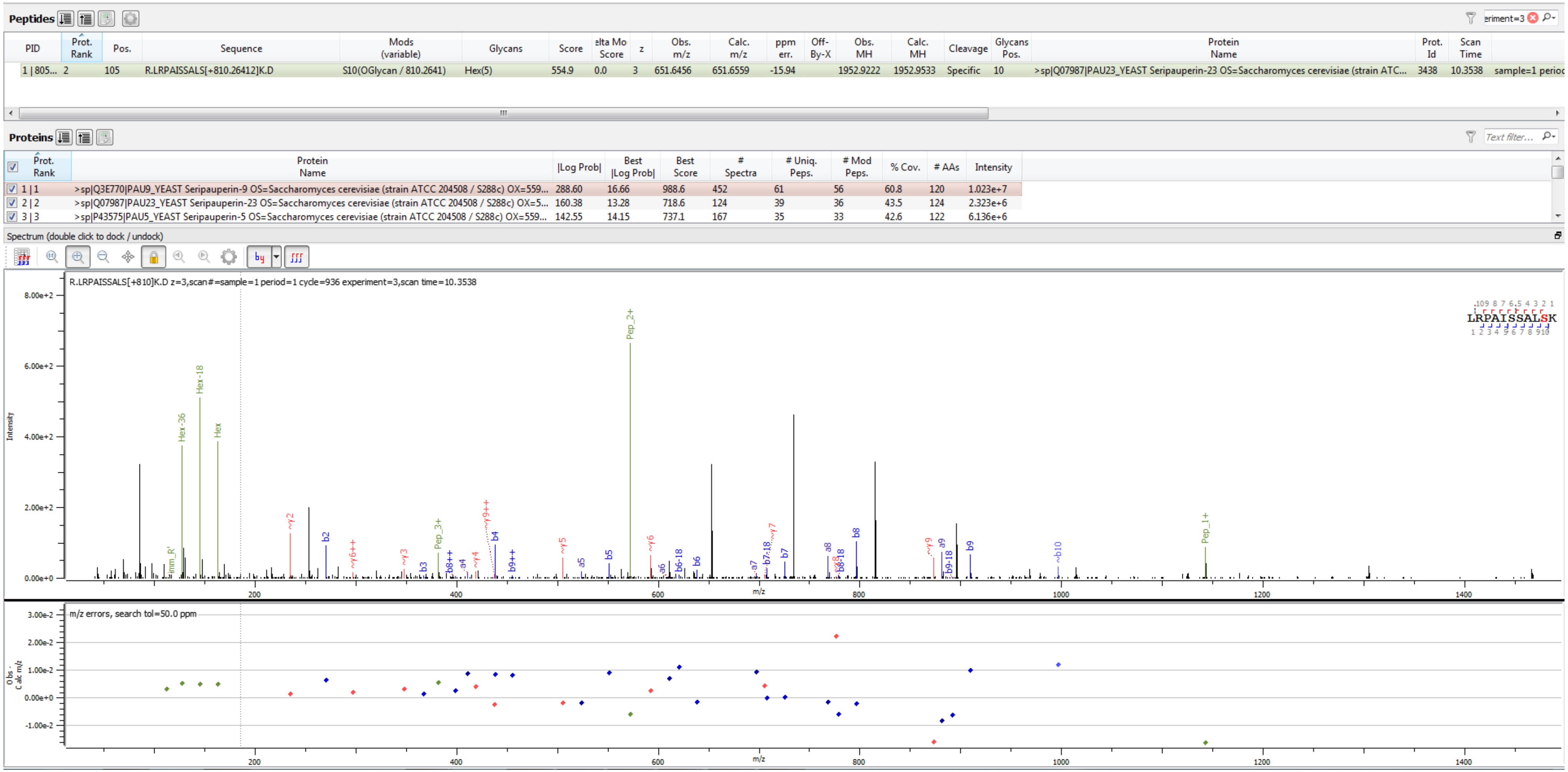


Sample S6 R.LRPAISSALS[+810.264]K.D

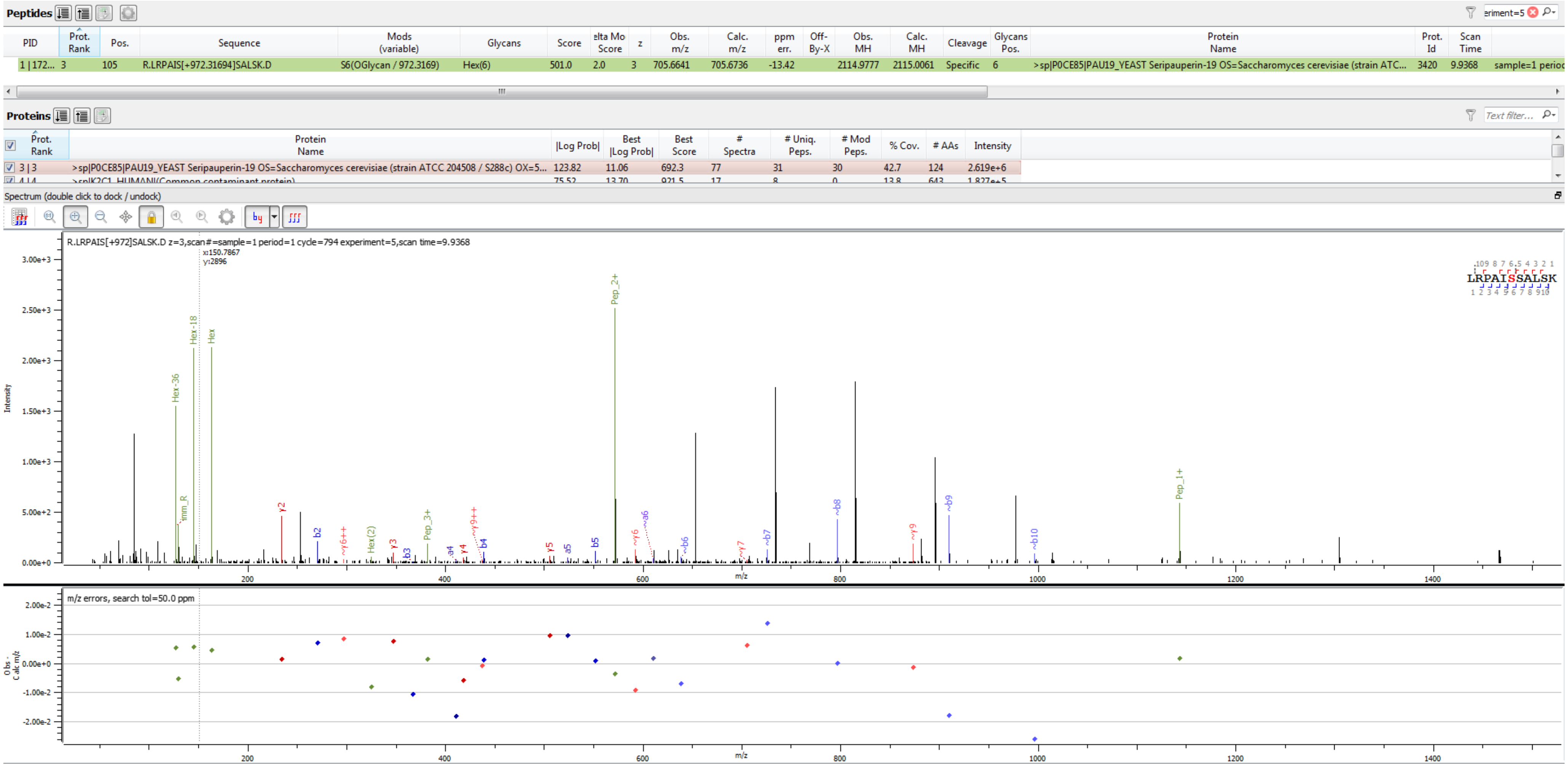


Sample S3 R.LRPAIS[+972.317]SALSK.D

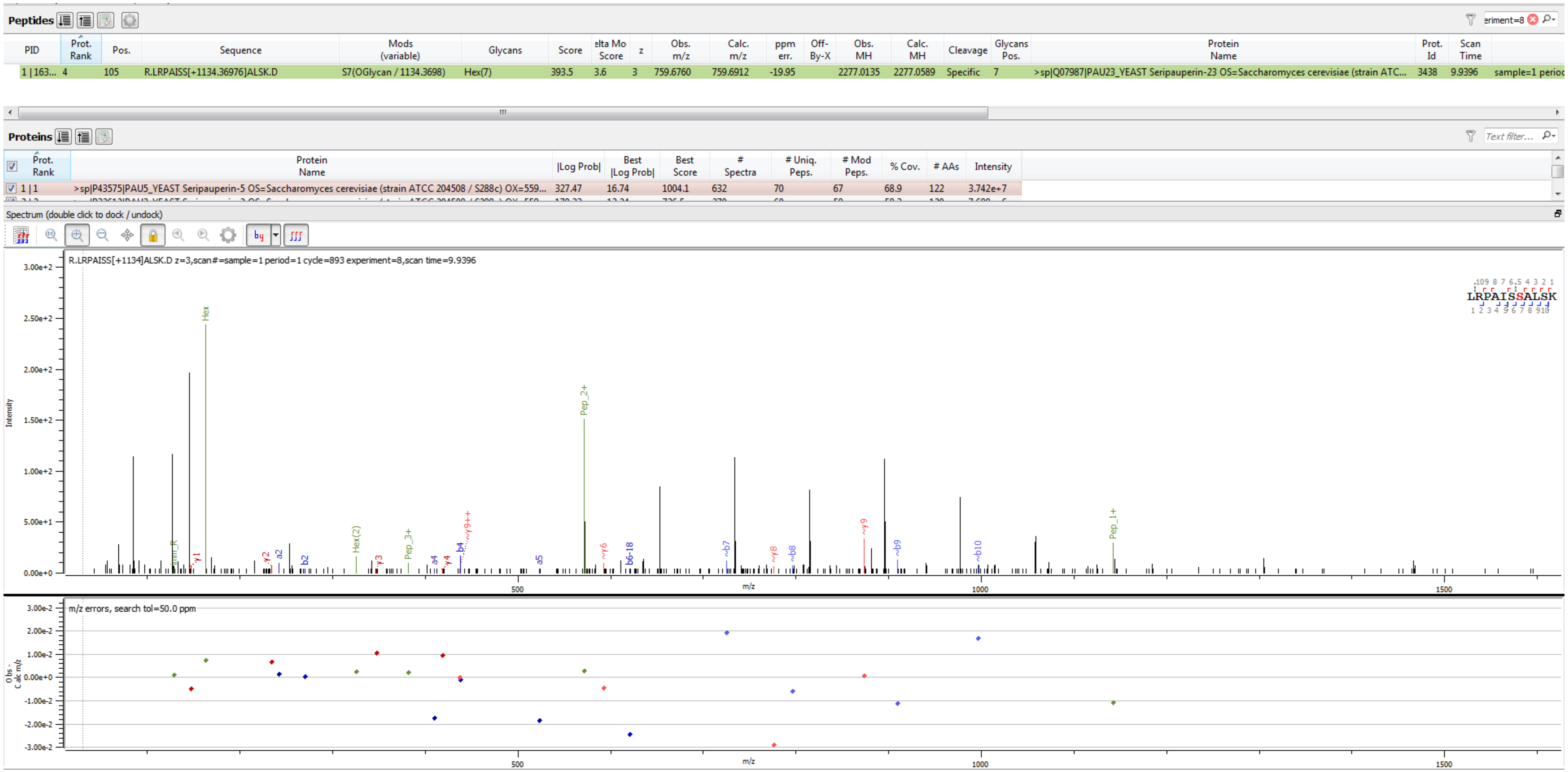


Sample S1 R.LRPAISS[+1134.370]ALSK.D

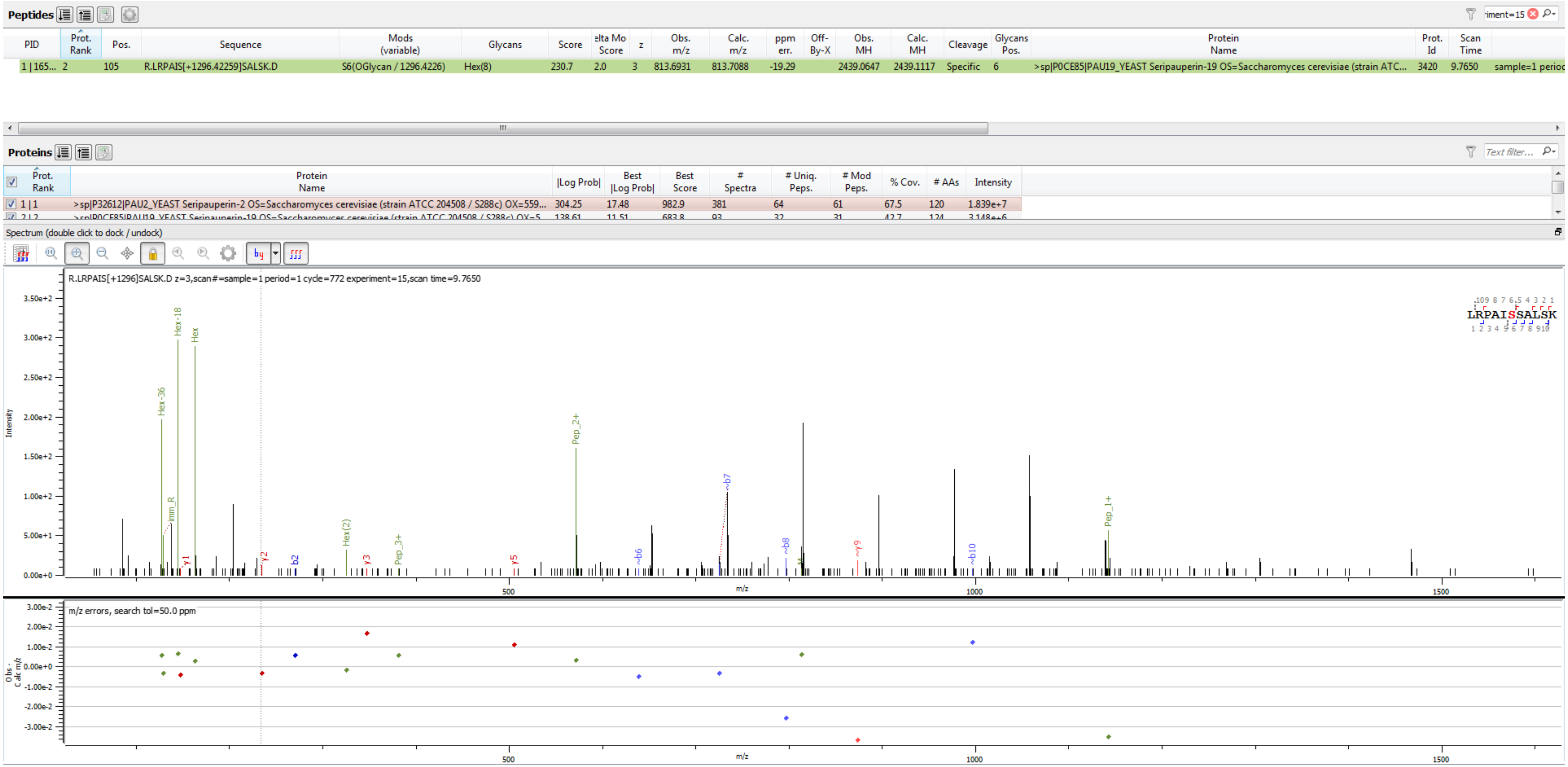


Sample S4 R.LRPAIS[+1296.423]SALSK.D

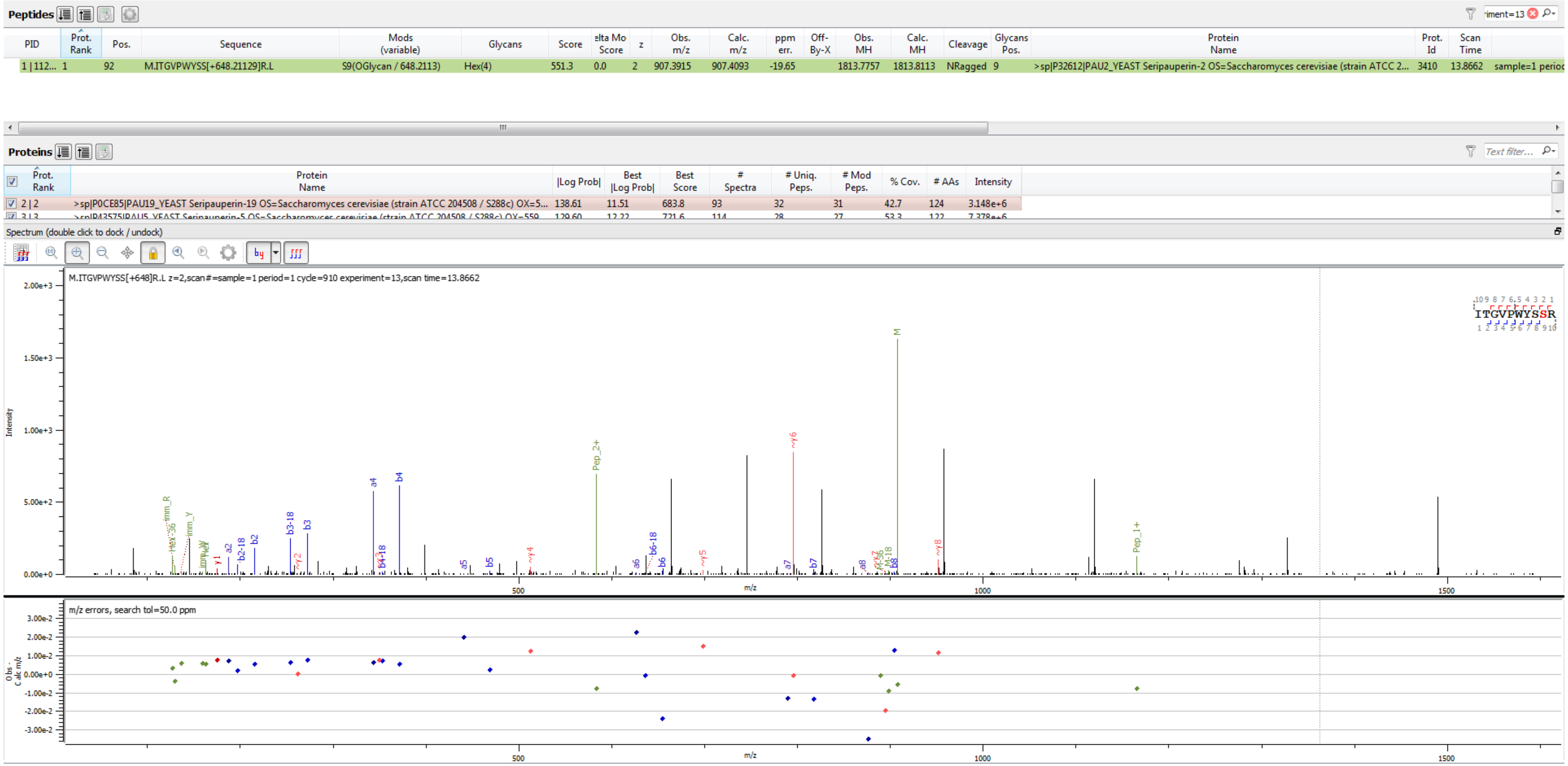


Sample S4 M.ITGVPWYSS[+648.211]R.L

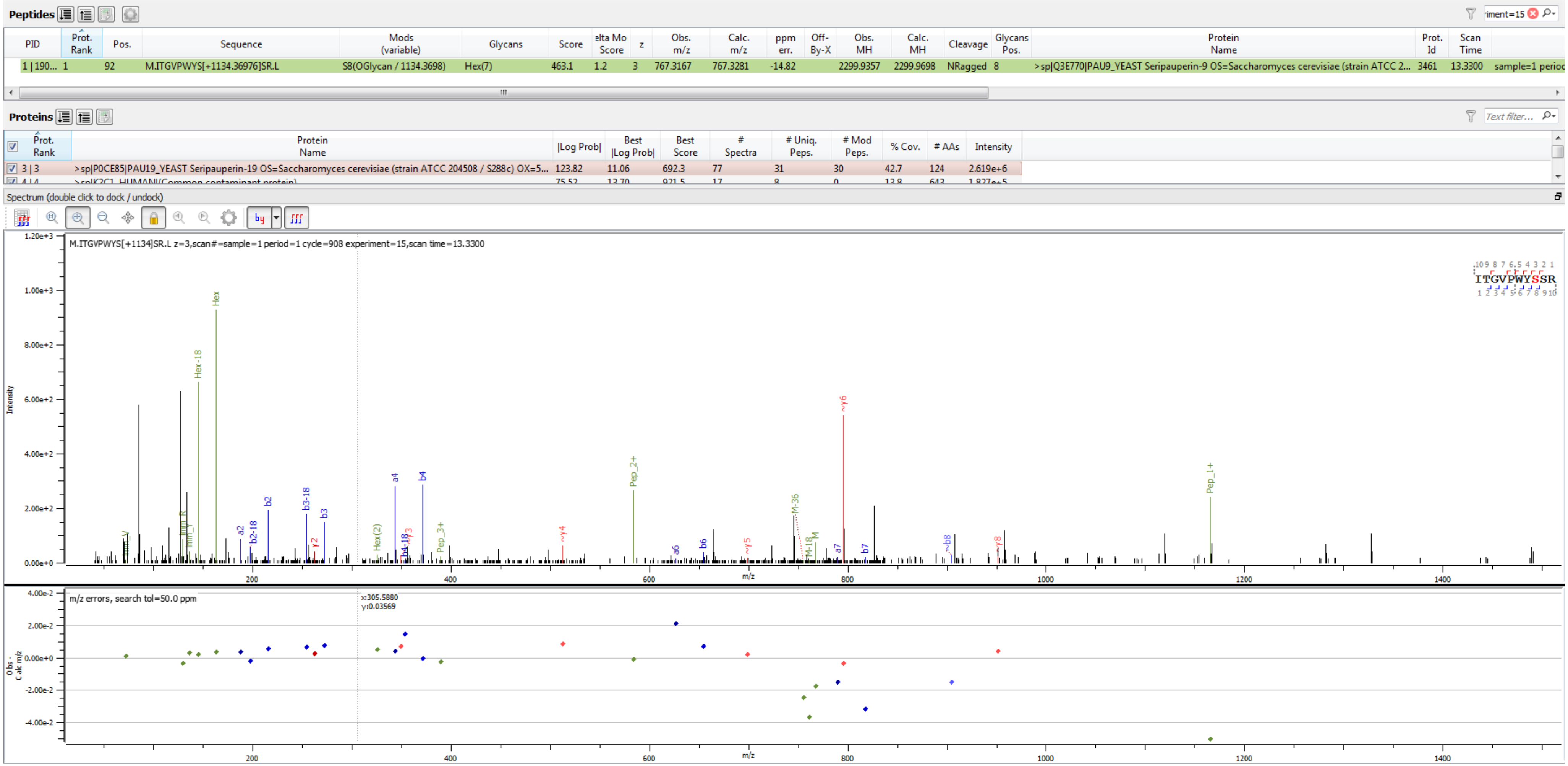


Sample S3 M.ITGVPWYS[+1134.370]SR.L

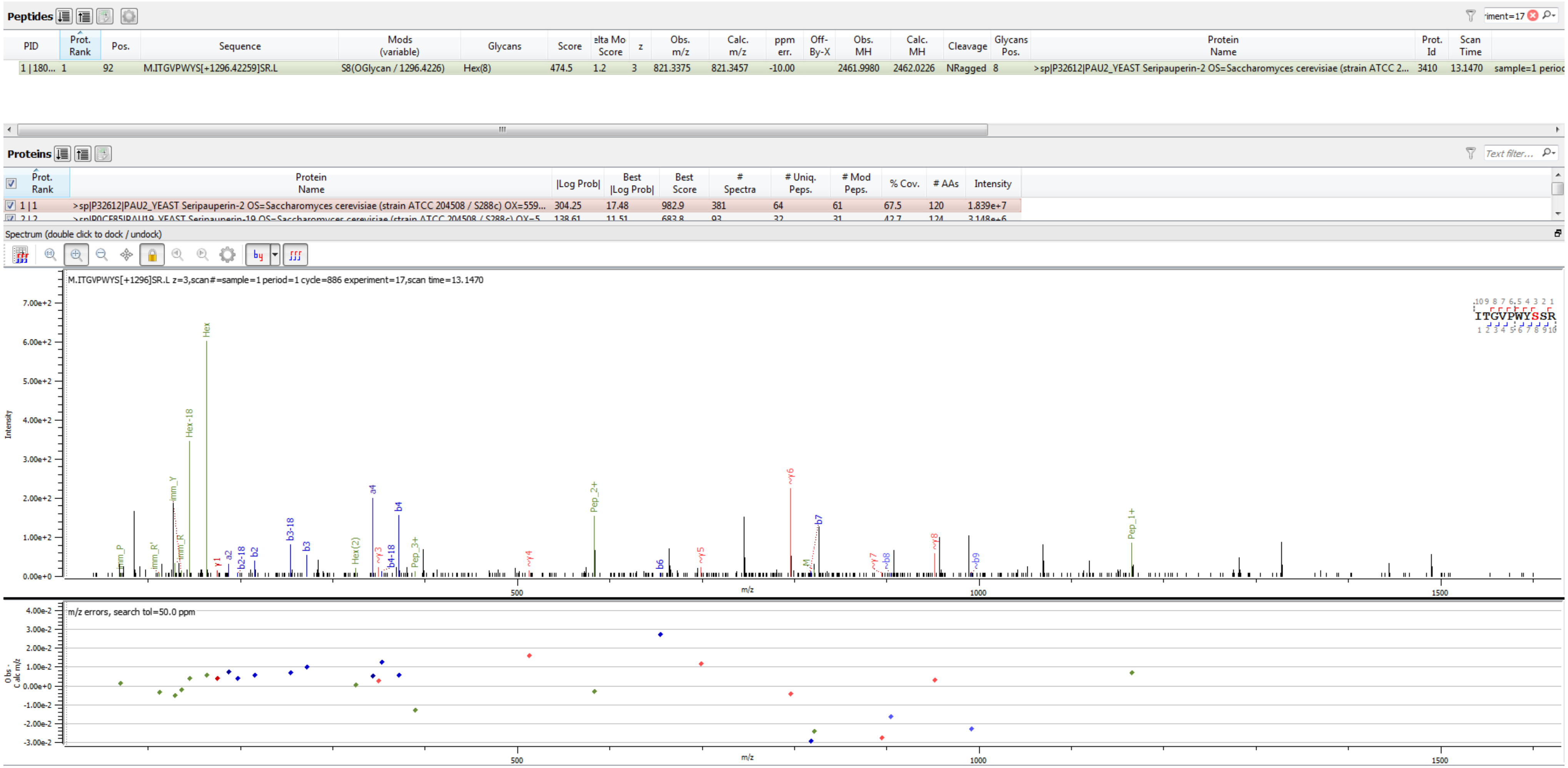


Sample S4 M.ITGVPWYS[+1296.423]SR.L

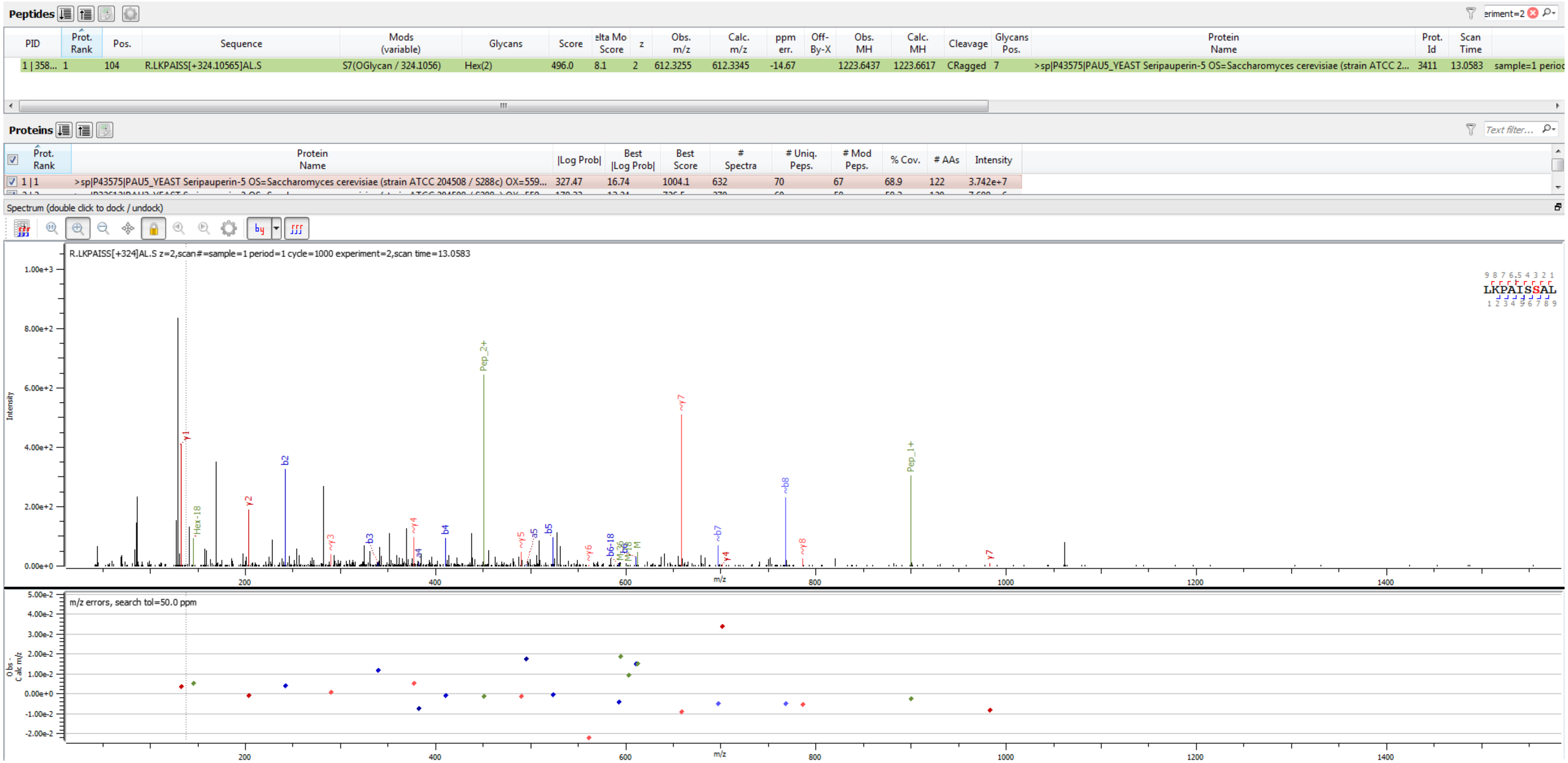


Sample S1 R.LKPAISS[+324.106]AL.S

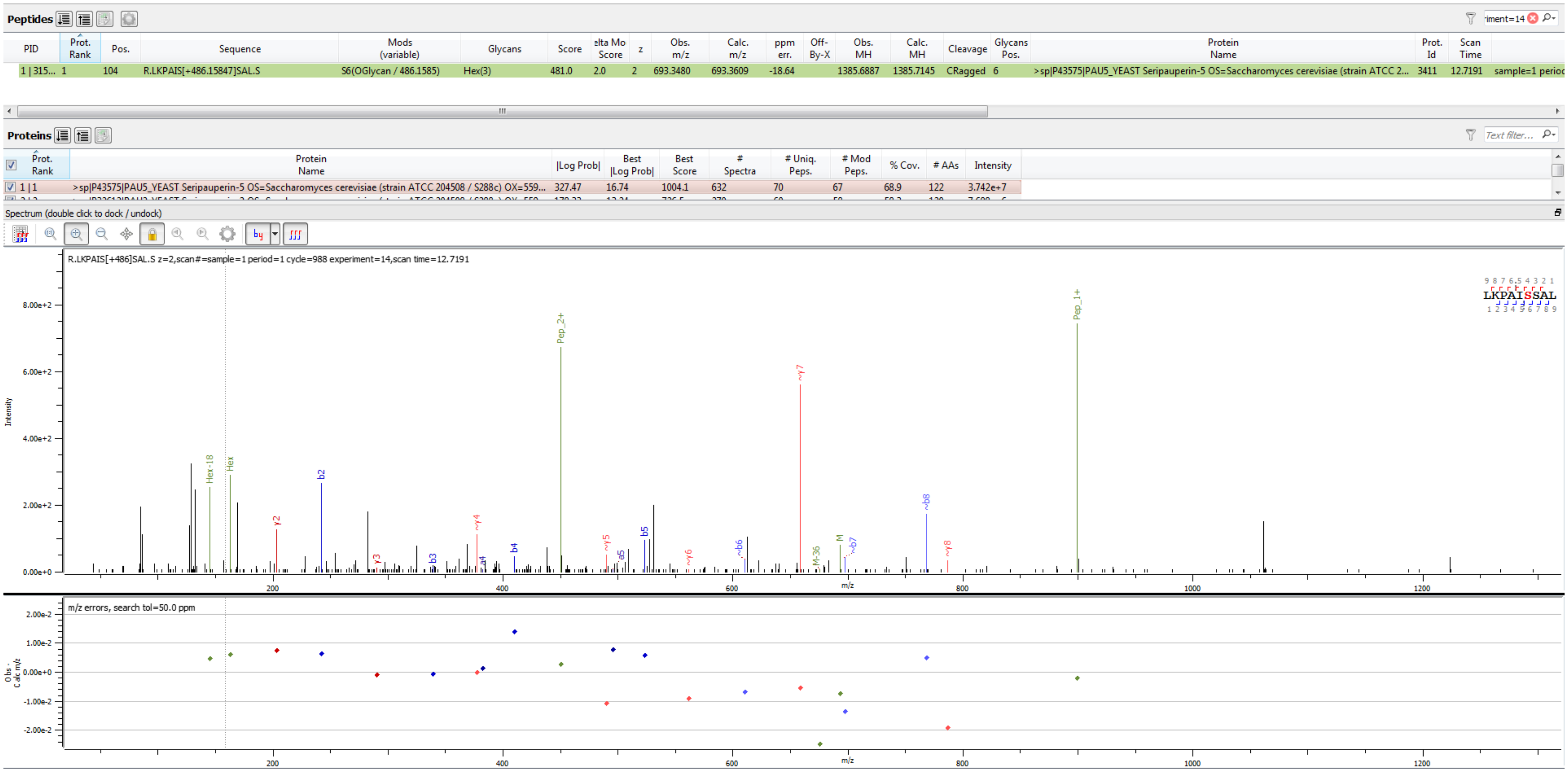


Sample S1 R.LKPAIS[+486.158]SAL.S

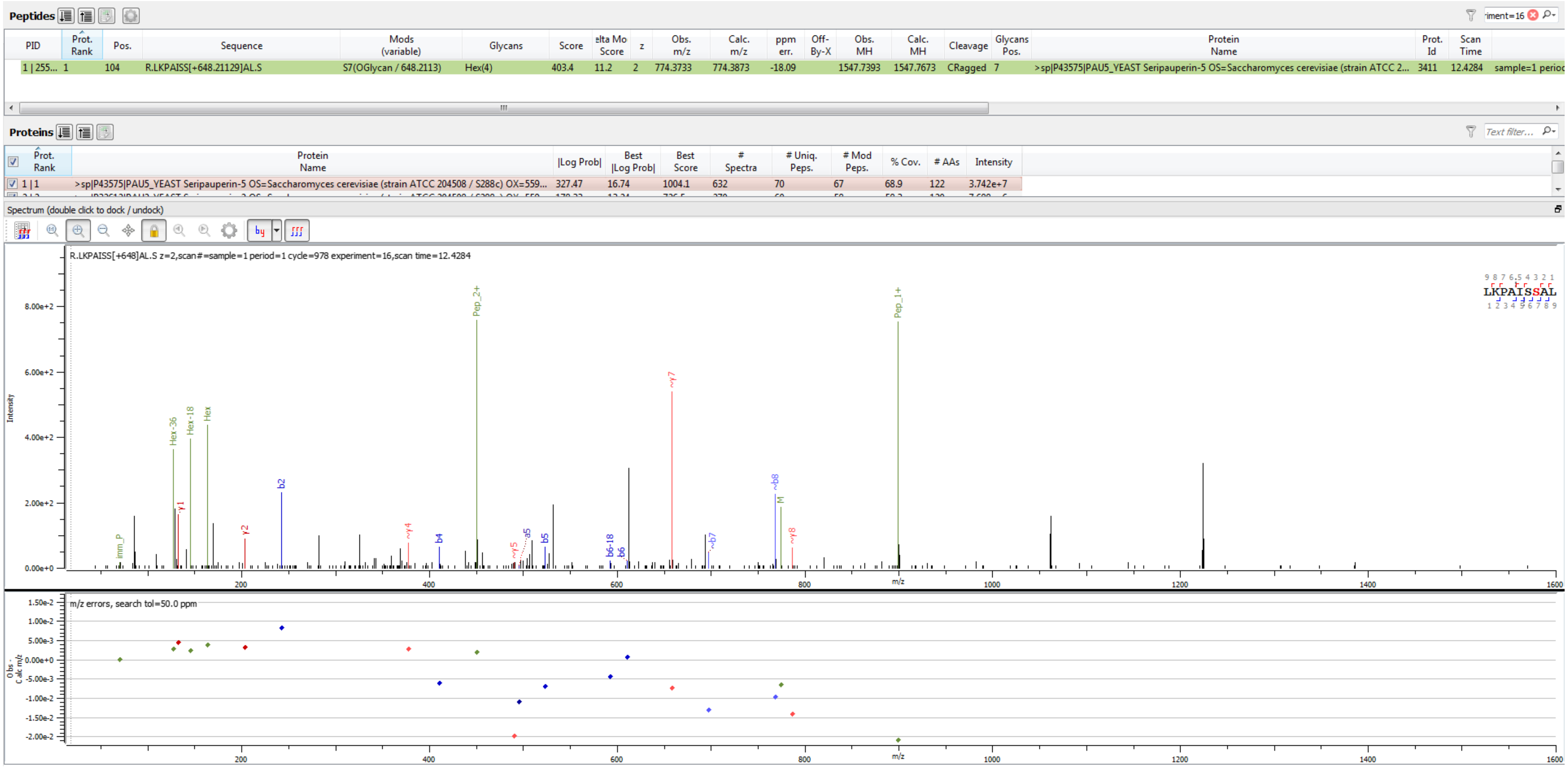


Sample S1 R.LKPAISS[+648.211]AL.S

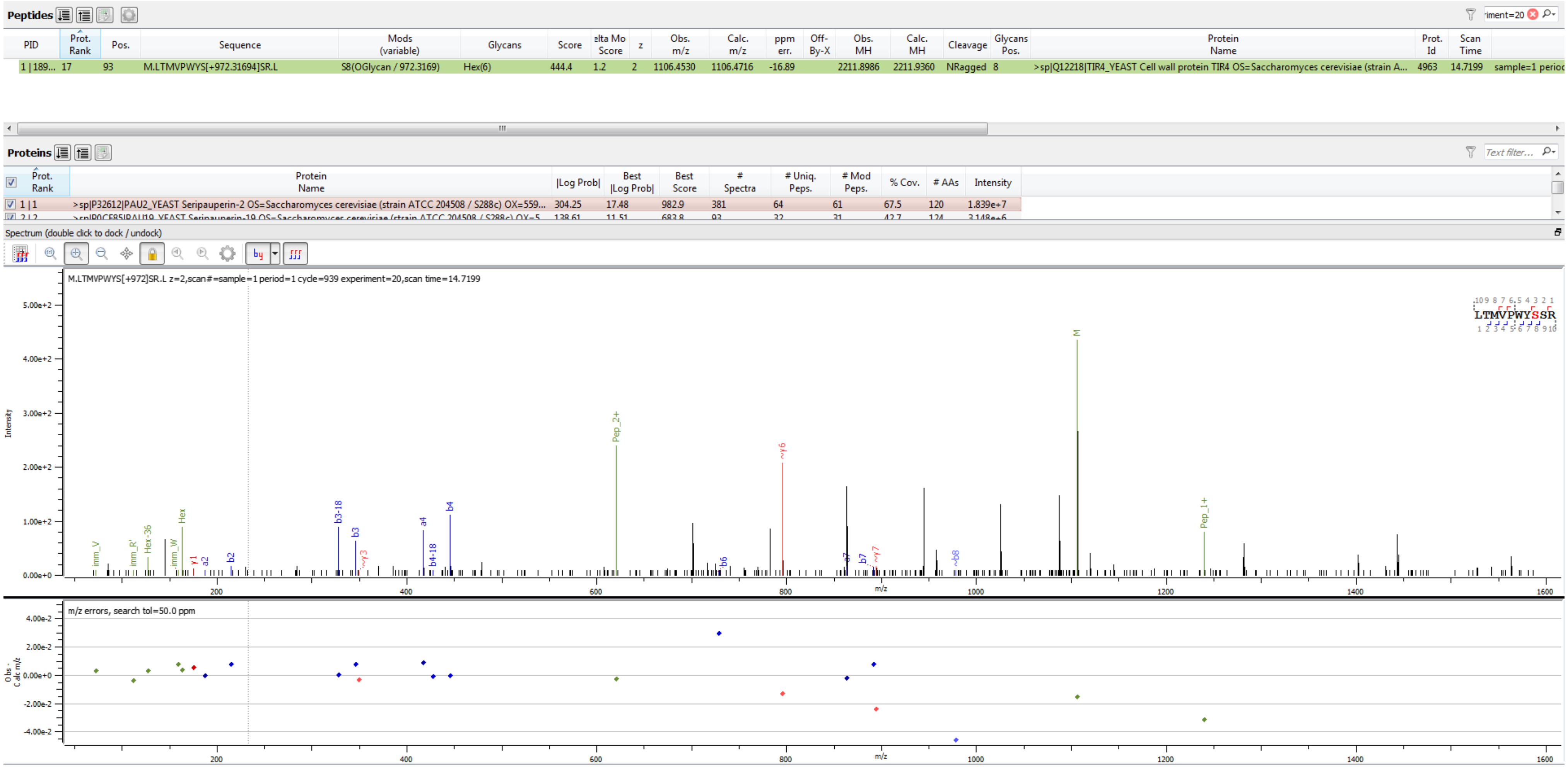


Sample S4 M.LTMVPWYS[+972.317]SR.L

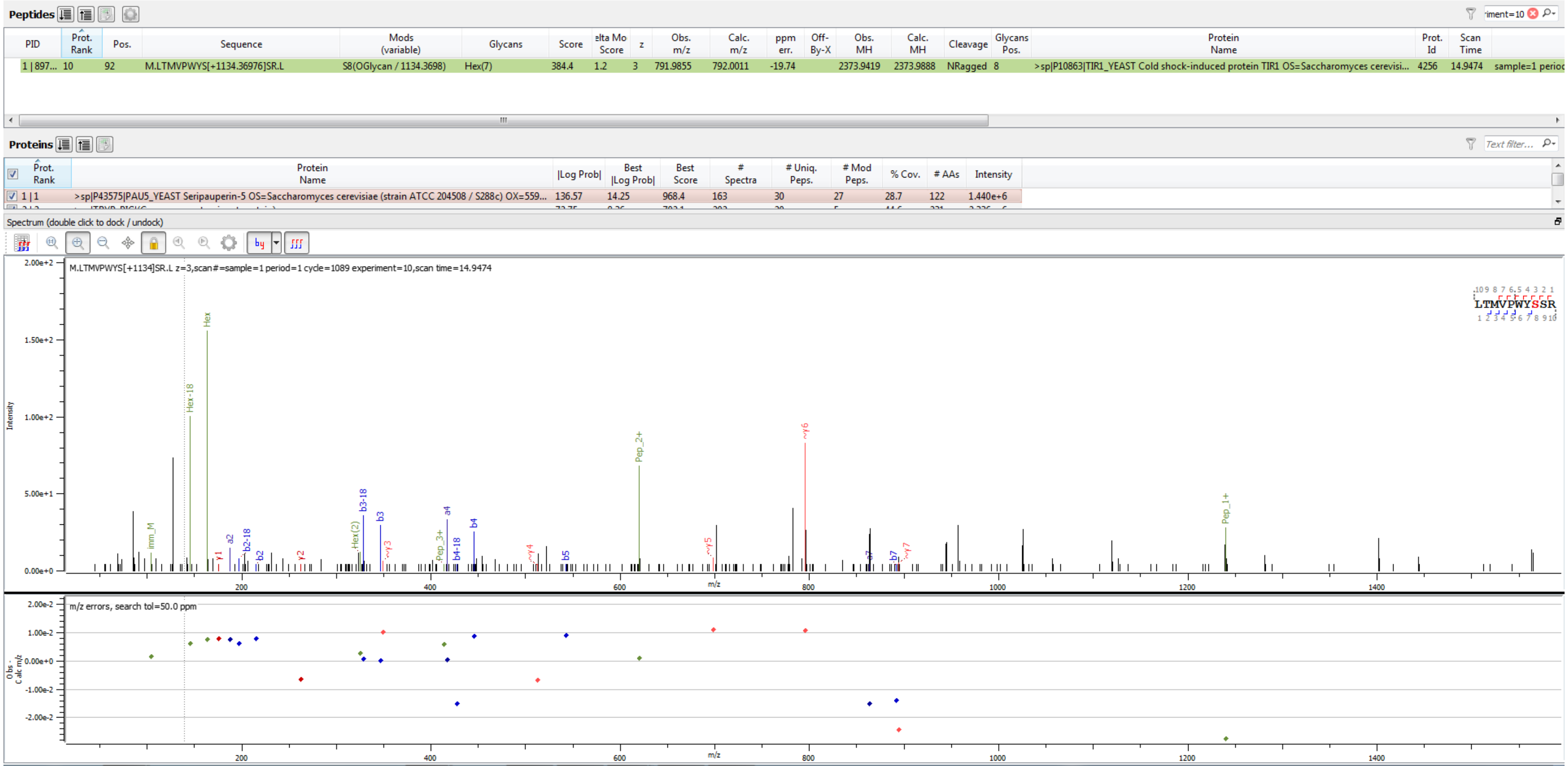


Sample S4 M.LTMVPWYS[+1134.370]SR.L

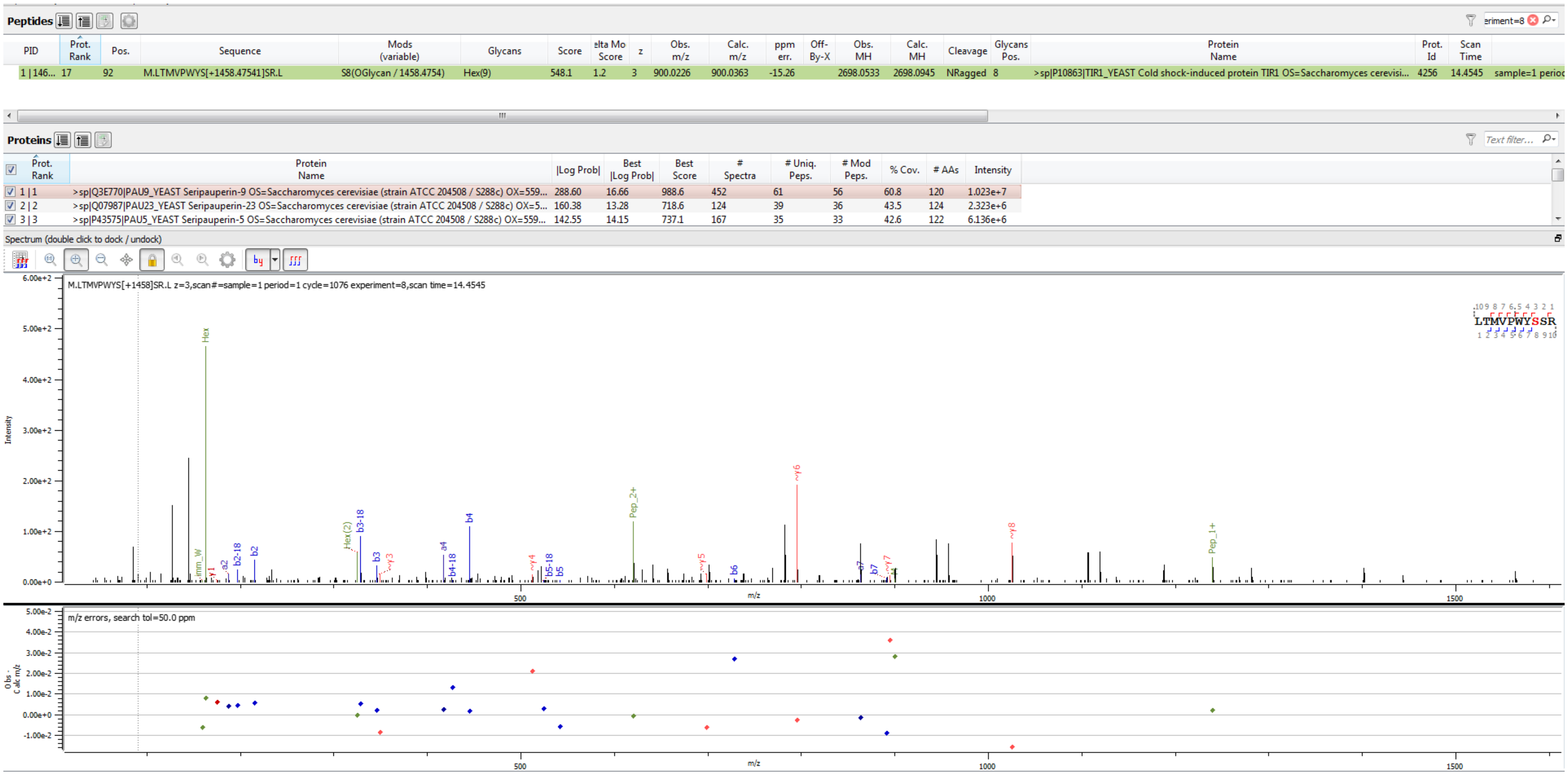

**Peptides**

| PID | Prot. Rank | Pos. | Sequence | Mods (variable) | Glycans | Score | elta Mo Score | z | Obs. m/z | Calc. m/z | ppm err. | Off-By-X | Obs. MH | Calc. MH | Cleavage | Glycans Pos. | Protein Name | Prot. Id | Scan Time | |
|---|---|---|---|---|---|---|---|---|---|---|---|---|---|---|---|---|---|---|---|---|
| 1 \| 146... | 17 | 92 | M.LTMVPWYS[+1458.47541]SR.L | S8(OGlycan / 1458.4754) | Hex(9) | 548.1 | 1.2 | 3 | 900.0226 | 900.0363 | -15.26 | | 2698.0533 | 2698.0945 | NRagged | 8 | >sp\|P10863\|TIR1_YEAST Cold shock-induced protein TIR1 OS=Saccharomyces cerevisi... | 4256 | 14.4545 | sample=1 perioc |

**Proteins**

| | Prot. Rank | Protein Name | \|Log Prob\| | Best \|Log Prob\| | Best Score | # Spectra | # Uniq. Peps. | # Mod Peps. | % Cov. | # AAs | Intensity |
|---|---|---|---|---|---|---|---|---|---|---|---|
| ☑ | 1 \| 1 | >sp\|Q3E770\|PAU9_YEAST Seripauperin-9 OS=Saccharomyces cerevisiae (strain ATCC 204508 / S288c) OX=559... | 288.60 | 16.66 | 988.6 | 452 | 61 | 56 | 60.8 | 120 | 1.023e+7 |
| ☑ | 2 \| 2 | >sp\|Q07987\|PAU23_YEAST Seripauperin-23 OS=Saccharomyces cerevisiae (strain ATCC 204508 / S288c) OX=5... | 160.38 | 13.28 | 718.6 | 124 | 39 | 36 | 43.5 | 124 | 2.323e+6 |
| ☑ | 3 \| 3 | >sp\|P43575\|PAU5_YEAST Seripauperin-5 OS=Saccharomyces cerevisiae (strain ATCC 204508 / S288c) OX=559... | 142.55 | 14.15 | 737.1 | 167 | 35 | 33 | 42.6 | 122 | 6.136e+6 |



Sample S6 M.LTMVPWYS[+1458.475]SR.L

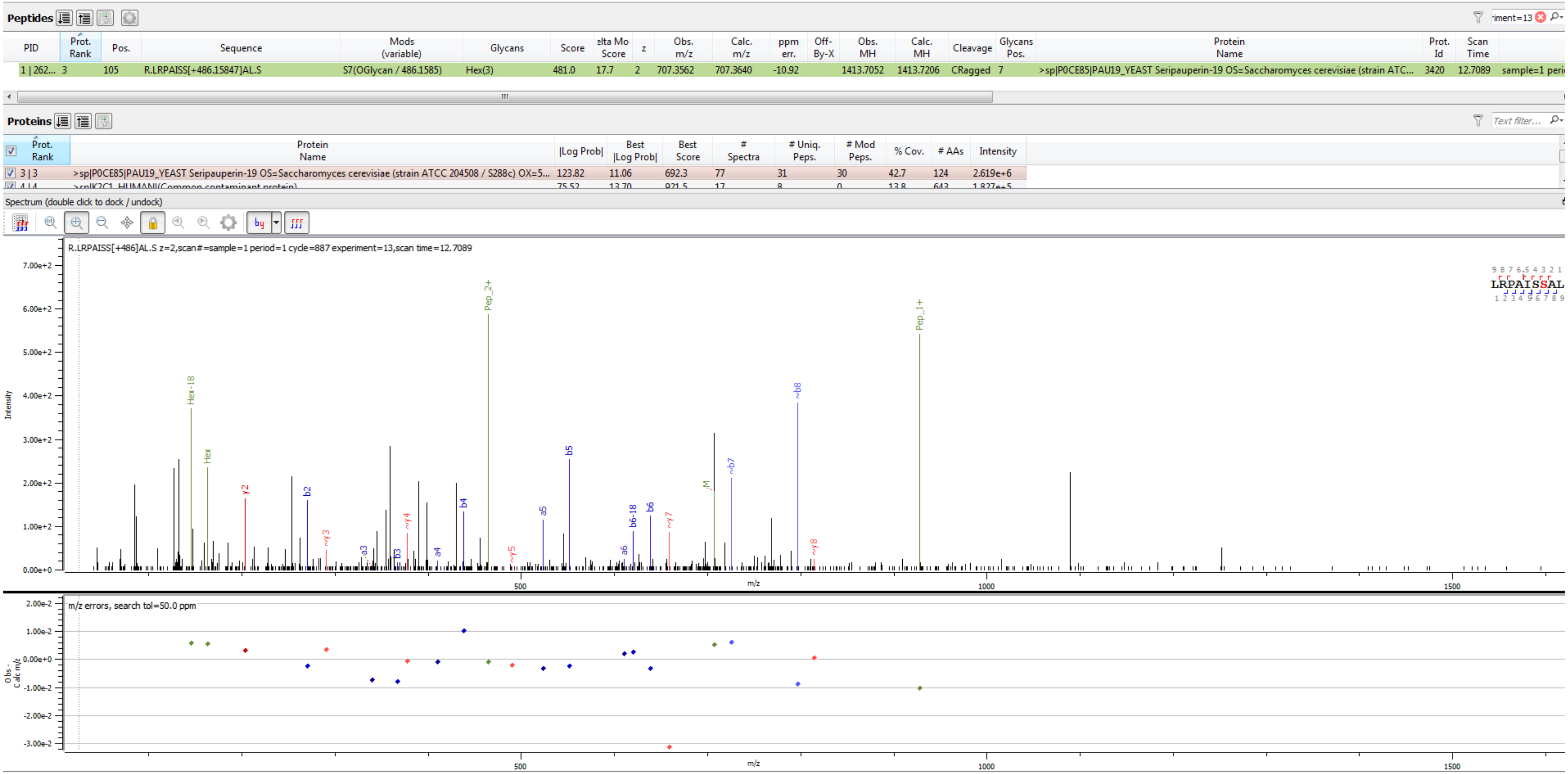


Sample S3 R.LRPAISS[+486.158]AL.S

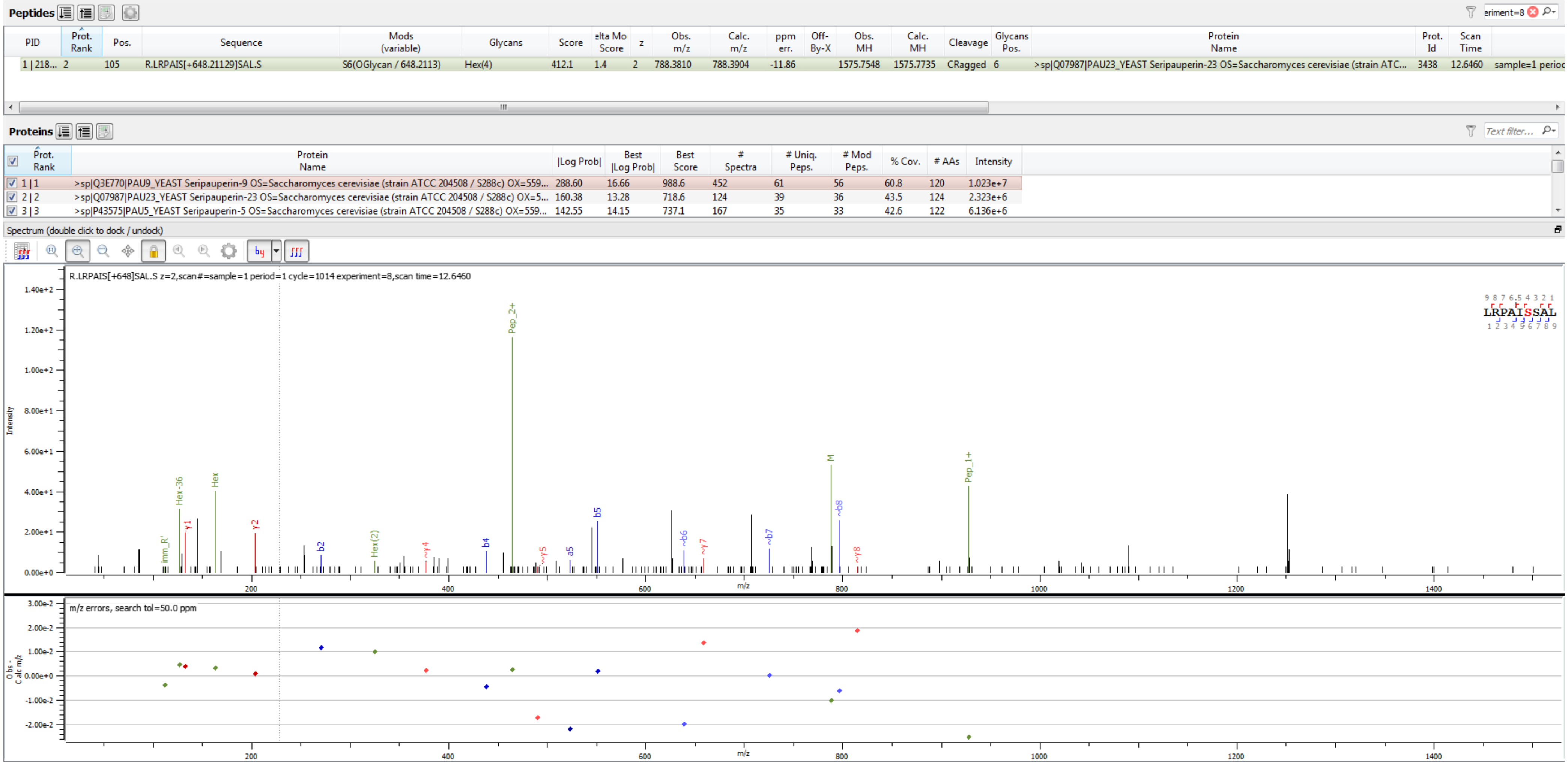


Sample S6 R.LRPAIS[+648.211]SAL.S

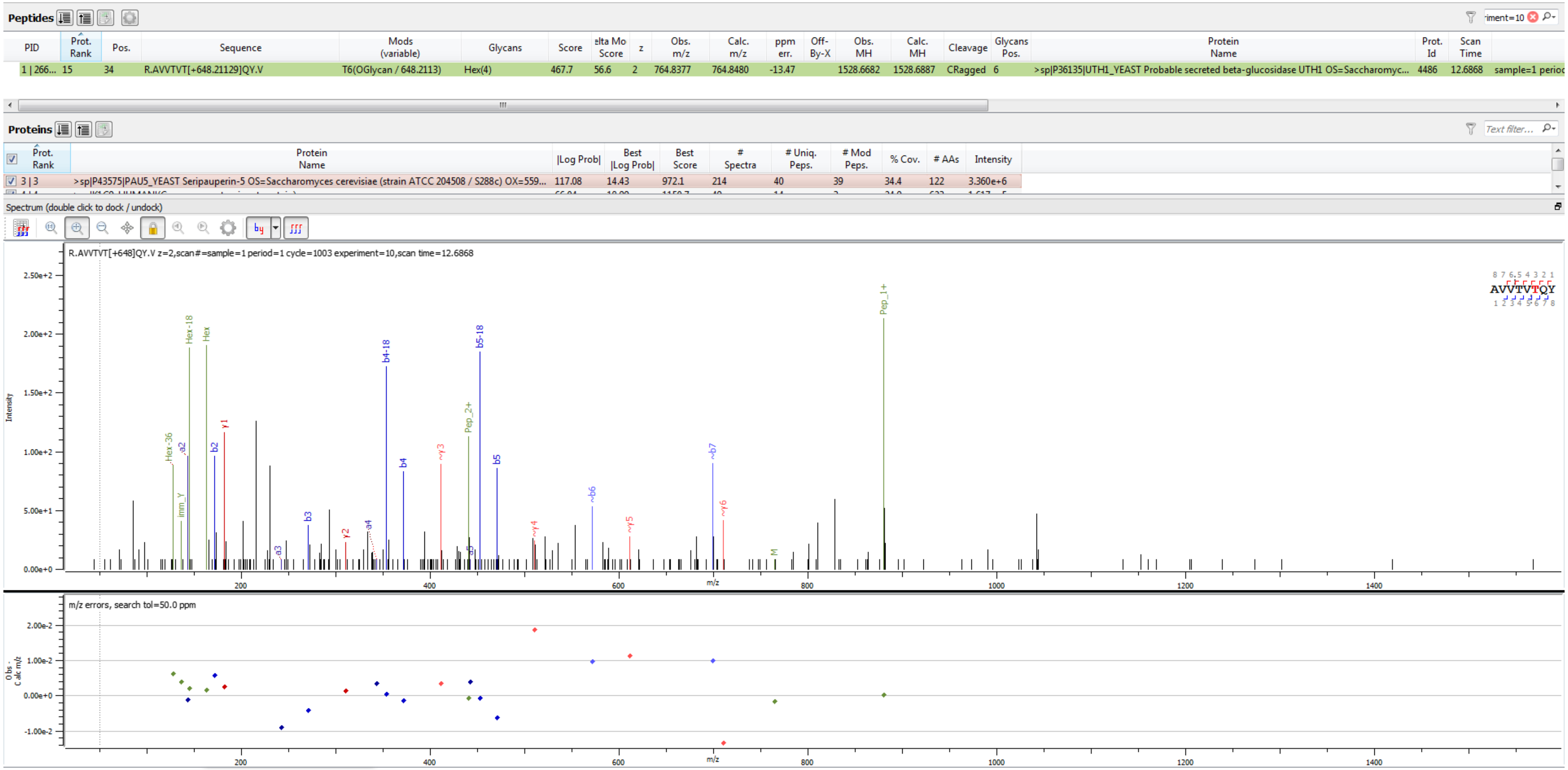

Sample S2 R.AVVTVT[+648.211]QY.V

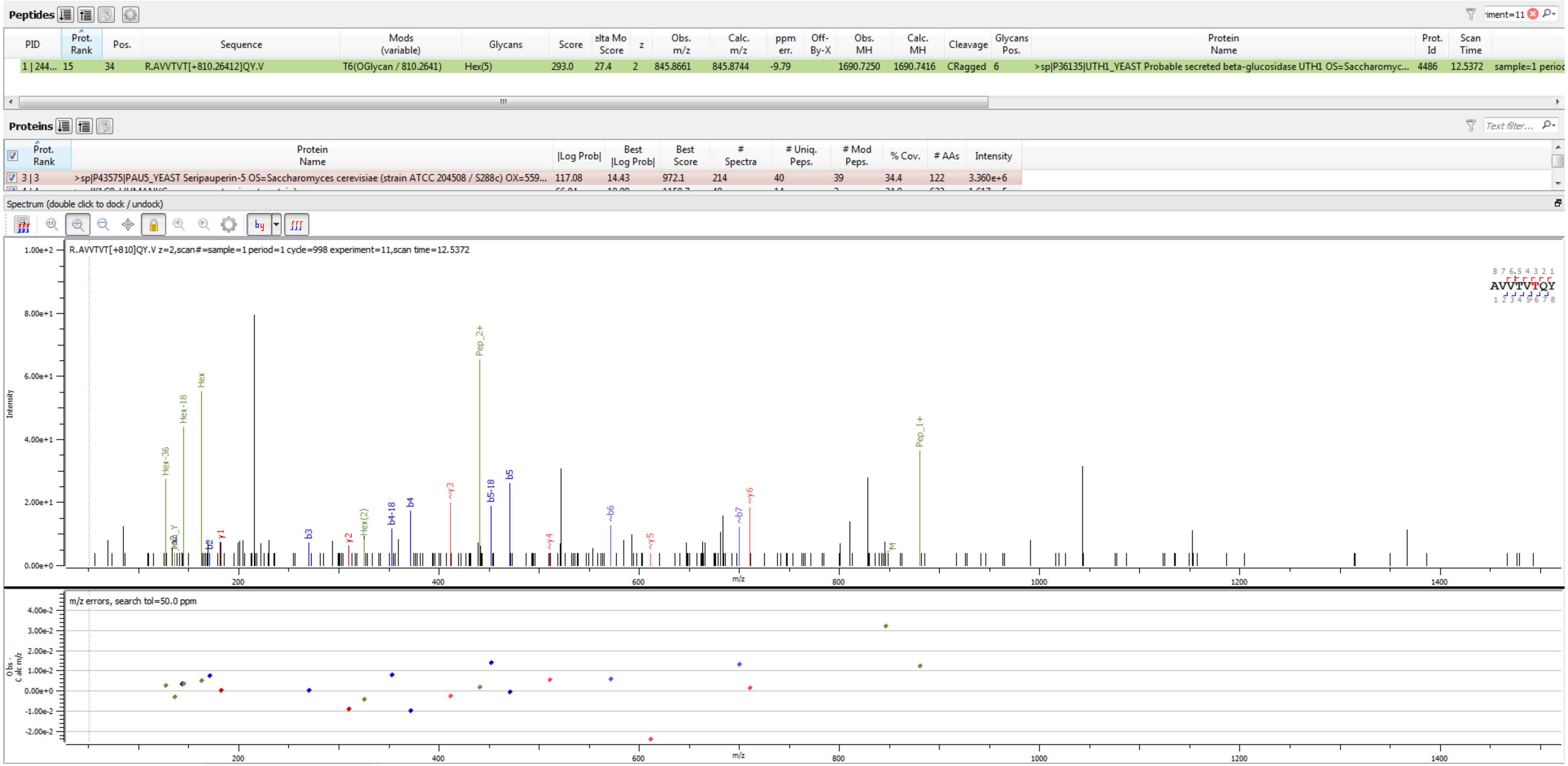


Sample S2 R.AVVTVT[+810.264]QY.V

**Peptides**

<table>
<tr><th>PID</th><th>Prot. Rank</th><th>Pos.</th><th>Sequence</th><th>Mods (variable)</th><th>Glycans</th><th>Score</th><th>elta Mo Score</th><th>z</th><th>Obs. m/z</th><th>Calc. m/z</th><th>ppm err.</th><th>Off-By-X</th><th>Obs. MH</th><th>Calc. MH</th><th>Cleavage</th><th>Glycans Pos.</th><th>Protein Name</th><th>Prot. Id</th><th>Scan Time</th><th></th></tr>
<tr><td>1 | 425...</td><td>19</td><td>93</td><td>K.LPWYT[+486.15847]TR.L</td><td>T5(OGlycan / 486.1585)</td><td>Hex(3)</td><td>460.4</td><td>24.3</td><td>2</td><td>711.8204</td><td>711.8298</td><td>-13.10</td><td></td><td>1422.6336</td><td>1422.6522</td><td>Specific</td><td>5</td><td>>sp|P27654|TIP1_YEAST Temperature shock-inducible protein 1 OS=Saccharomyces c...</td><td>4237</td><td>13.5003</td><td>sample=1 perioc</td></tr>
</table>

**Proteins**

<table>
<tr><th>Prot. Rank</th><th>Protein Name</th><th>|Log Prob|</th><th>Best |Log Prob|</th><th>Best Score</th><th># Spectra</th><th># Uniq. Peps.</th><th># Mod Peps.</th><th>% Cov.</th><th># AAs</th><th>Intensity</th></tr>
<tr><td>1 | 1</td><td>>sp|P43575|PAU5_YEAST Seripauperin-5 OS=Saccharomyces cerevisiae (strain ATCC 204508 / S288c) OX=559...</td><td>327.47</td><td>16.74</td><td>1004.1</td><td>632</td><td>70</td><td>67</td><td>68.9</td><td>122</td><td>3.742e+7</td></tr>
</table>

Spectrum (double click to dock / undock)

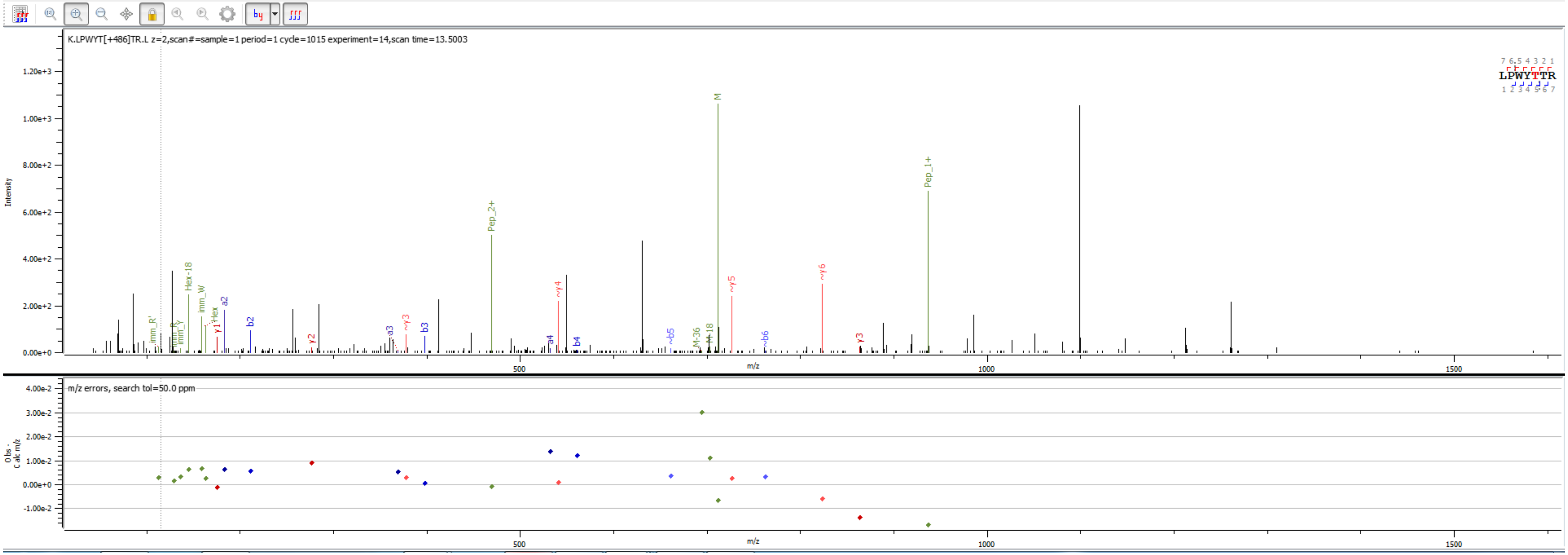


Sample S1 K.LPWYT[+486.158]TR.L

**Peptides**

<table>
<tr><th>PID</th><th>Prot. Rank</th><th>Pos.</th><th>Sequence</th><th>Mods (variable)</th><th>Glycans</th><th>Score</th><th>elta Mo Score</th><th>z</th><th>Obs. m/z</th><th>Calc. m/z</th><th>ppm err.</th><th>Off-By-X</th><th>Obs. MH</th><th>Calc. MH</th><th>Cleavage</th><th>Glycans Pos.</th><th>Protein Name</th><th>Prot. Id</th><th>Scan Time</th><th></th></tr>
<tr><td>1 | 316...</td><td>22</td><td>93</td><td>K.LPWYT[+648.21129]TR.L</td><td>T5(OGlycan / 648.2113)</td><td>Hex(4)</td><td>408.8</td><td>1.0</td><td>2</td><td>792.8440</td><td>792.8562</td><td>-15.32</td><td></td><td>1584.6808</td><td>1584.7051</td><td>Specific</td><td>5</td><td>>sp|P27654|TIP1_YEAST Temperature shock-inducible protein 1 OS=Saccharomyces c...</td><td>4237</td><td>13.4180</td><td>sample=1 perioc</td></tr>
</table>

**Proteins**

<table>
<tr><th>Prot. Rank</th><th>Protein Name</th><th>|Log Prob|</th><th>Best |Log Prob|</th><th>Best Score</th><th># Spectra</th><th># Uniq. Peps.</th><th># Mod Peps.</th><th>% Cov.</th><th># AAs</th><th>Intensity</th></tr>
<tr><td>1 | 1</td><td>>sp|Q3E770|PAU9_YEAST Seripauperin-9 OS=Saccharomyces cerevisiae (strain ATCC 204508 / S288c) OX=559...</td><td>288.60</td><td>16.66</td><td>988.6</td><td>452</td><td>61</td><td>56</td><td>60.8</td><td>120</td><td>1.023e+7</td></tr>
<tr><td>2 | 2</td><td>>sp|Q07987|PAU23_YEAST Seripauperin-23 OS=Saccharomyces cerevisiae (strain ATCC 204508 / S288c) OX=5...</td><td>160.38</td><td>13.28</td><td>718.6</td><td>124</td><td>39</td><td>36</td><td>43.5</td><td>124</td><td>2.323e+6</td></tr>
<tr><td>3 | 3</td><td>>sp|P43575|PAU5_YEAST Seripauperin-5 OS=Saccharomyces cerevisiae (strain ATCC 204508 / S288c) OX=559...</td><td>142.55</td><td>14.15</td><td>737.1</td><td>167</td><td>35</td><td>33</td><td>42.6</td><td>122</td><td>6.136e+6</td></tr>
</table>

Spectrum (double click to dock / undock)

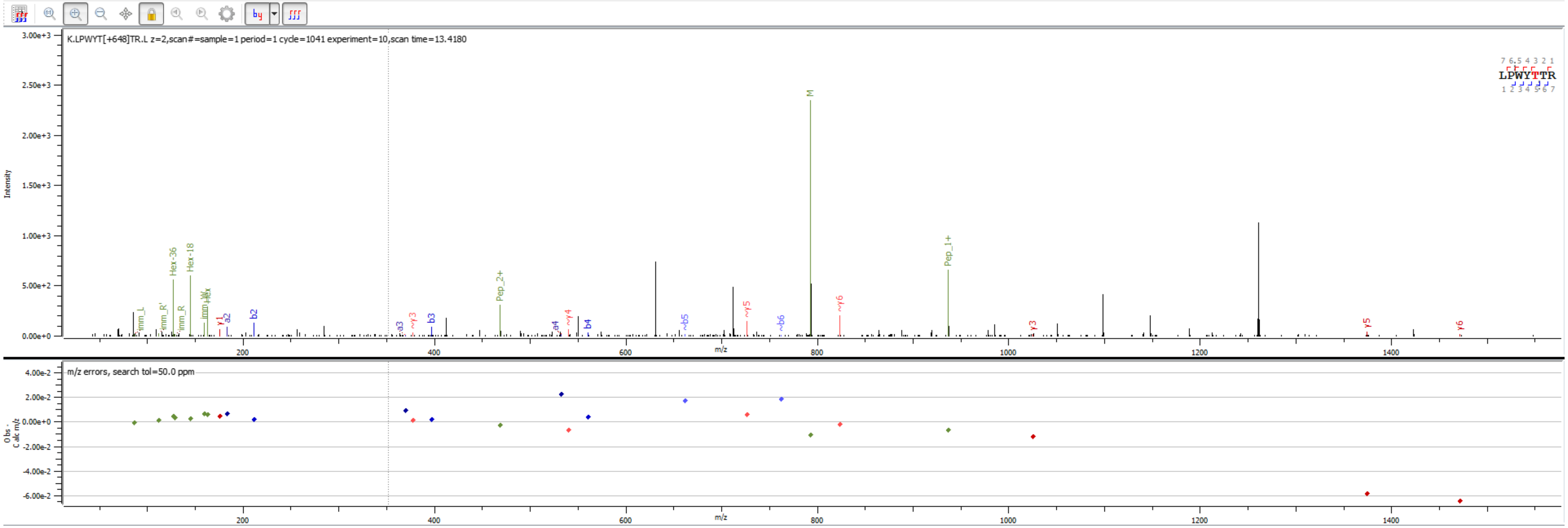


Sample S6 K.LPWYT[+648.211]TR.L

**Peptides**

| PID | Prot. Rank | Pos. | Sequence | Mods (variable) | Glycans | Score | elta Mo Score | z | Obs. m/z | Calc. m/z | ppm err. | Off-By-X | Obs. MH | Calc. MH | Cleavage | Glycans Pos. | Protein Name | Prot. Id | Scan Time | |
|---|---|---|---|---|---|---|---|---|---|---|---|---|---|---|---|---|---|---|---|---|
| 1 \| 109... | 21 | 33 | R.VNLIELAVYVS[+324.10565]DIR.A | S11(OGlycan / 324.1056) | Hex(2) | 552.7 | 316.1 | 2 | 964.4951 | 964.5091 | -14.52 | | 1927.9830 | 1928.0110 | Specific | 11 | >sp\|P47179\|DAN4_YEAST Cell wall protein DAN4 OS=Saccharomyces cerevisiae (strain ... | 273 | 19.6318 | sample=1 perioc |

**Proteins**

| Prot. Rank | Protein Name | \|Log Prob\| | Best \|Log Prob\| | Best Score | # Spectra | # Uniq. Peps. | # Mod Peps. | % Cov. | # AAs | Intensity |
|---|---|---|---|---|---|---|---|---|---|---|
| 1 \| 1 | >sp\|Q3E770\|PAU9_YEAST Seripauperin-9 OS=Saccharomyces cerevisiae (strain ATCC 204508 / S288c) OX=559... | 288.60 | 16.66 | 988.6 | 452 | 61 | 56 | 60.8 | 120 | 1.023e+7 |
| 2 \| 2 | >sp\|Q07987\|PAU23_YEAST Seripauperin-23 OS=Saccharomyces cerevisiae (strain ATCC 204508 / S288c) OX=5... | 160.38 | 13.28 | 718.6 | 124 | 39 | 36 | 43.5 | 124 | 2.323e+6 |
| 3 \| 3 | >sp\|P43575\|PAU5_YEAST Seripauperin-5 OS=Saccharomyces cerevisiae (strain ATCC 204508 / S288c) OX=559... | 142.55 | 14.15 | 737.1 | 167 | 35 | 33 | 42.6 | 122 | 6.136e+6 |

Spectrum (double click to dock / undock)

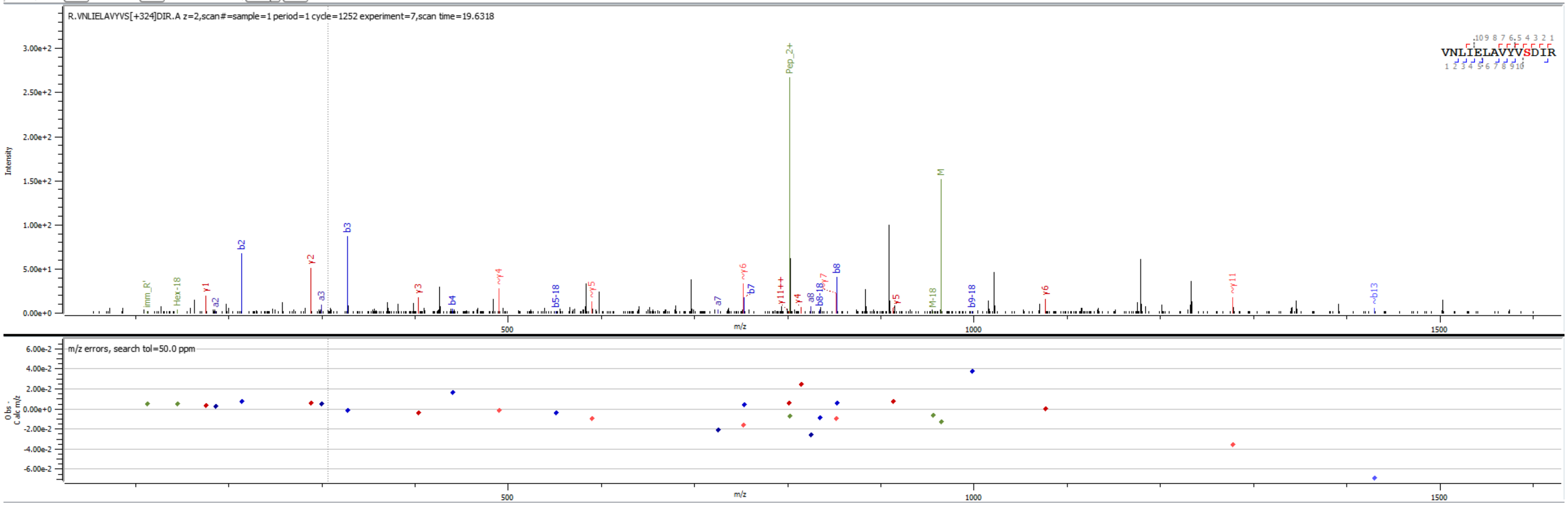


Sample S6 R.VNLIELAVYVS[+324.106]DIR.A

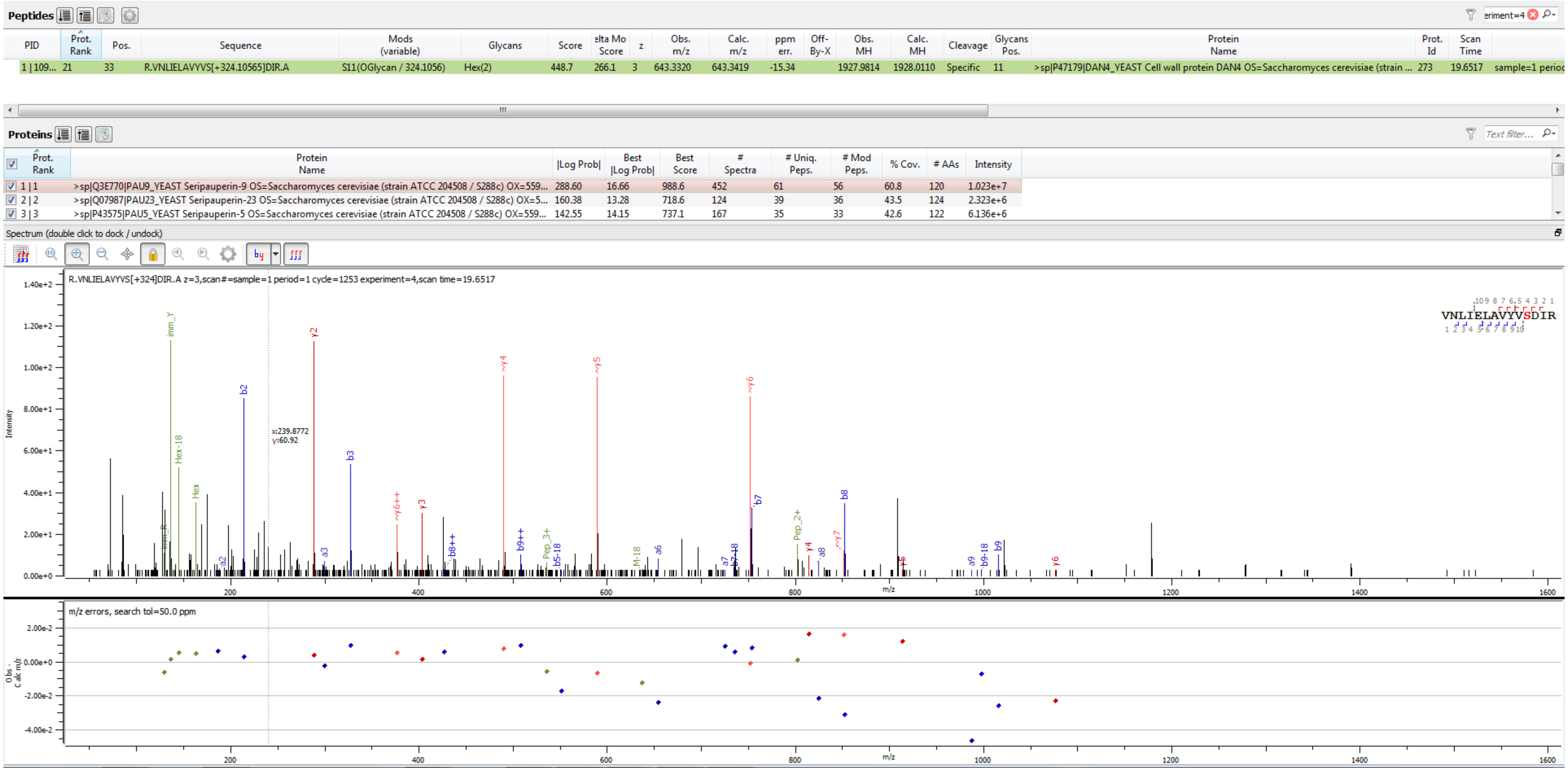


Sample S6 R.VNLIELAVYVS[+324.106]DIR.A

**Peptides**

| PID | Prot. Rank | Pos. | Sequence | Mods (variable) | Glycans | PEP 2D | PEP 1D | \|Log Prob\| | Score | Delta Score | elta Mo Score | z | Obs. m/z | Calc. m/z | ppm err. | Off-By-X | Obs. MH | Calc. MH | Cleavage | Glycans Pos. | Protein Name |
|---|---|---|---|---|---|---|---|---|---|---|---|---|---|---|---|---|---|---|---|---|---|
| 53 \| 21... | 2 | 173 | K.TDEYCCNSGSCN[+203.07937]ATTYSEFFK.T | N12(NGlycan / 203.0794) | HexNAc(1) | 4.2e-19 | 2e-15 | 18.37 | 922.9 | 742.6 | 742.6 | 3 | 858.3154 | 858.3311 | -18.38 | | 2572.9316 | 2572.9789 | Specific | 12 | >XP_002282988.1 PREDICTED: protein P21 [Vitis |

**Proteins**

| Prot. Rank | Protein Name | \|Log Prob\| | Best \|Log Prob\| | Best Score | # Spectra | # Uniq. Peps. | # Mod Peps. | % Cov. | # AAs | Intensity |
|---|---|---|---|---|---|---|---|---|---|---|
| 2 \| 2 | >XP_002282988.1 PREDICTED: protein P21 [Vitis vinifera] | 204.30 | 16.98 | 1052.5 | 416 | 32 | 8 | 52.9 | 225 | 1.215e+7 |
| 3 \| 3 | >NP_001268061.1 VVTL1 precursor [Vitis vinifera] | 171.94 | 18.82 | 1055.1 | 122 | 20 | 3 | 63.1 | 222 | 2.065e+6 |

Spectrum (double click to dock / undock)

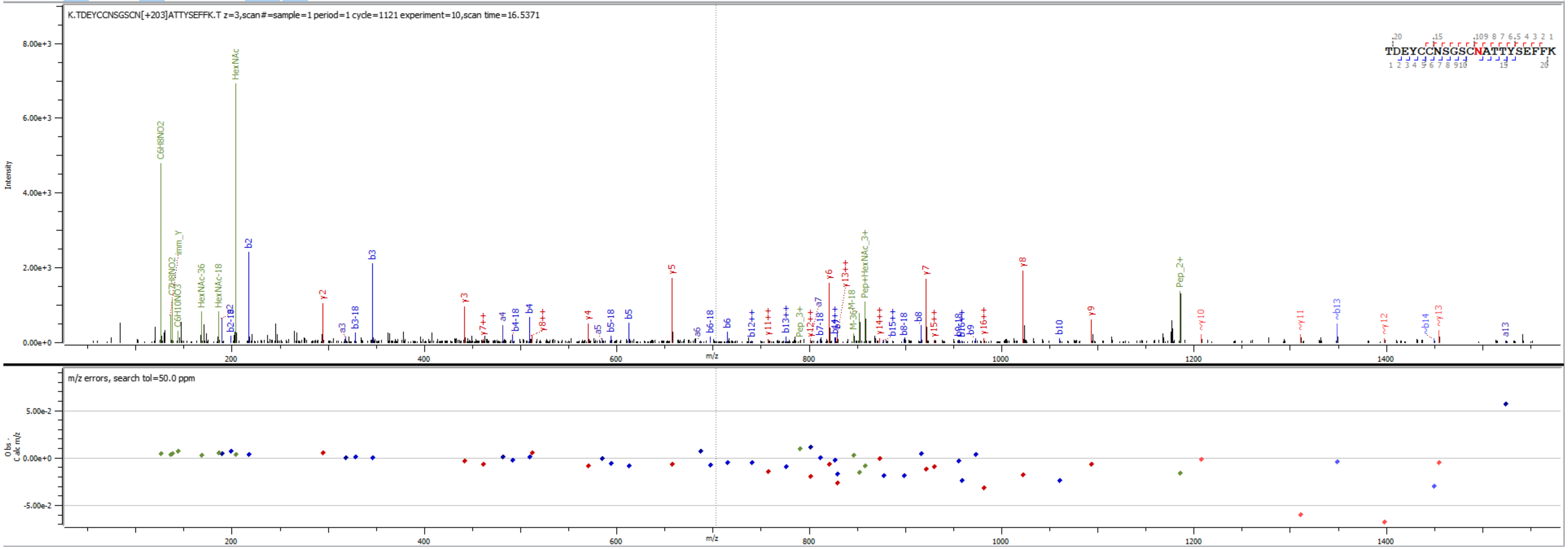


Sample S1 K.TDEYCCNSGSCN[+203.079]ATTYSEFFK.T

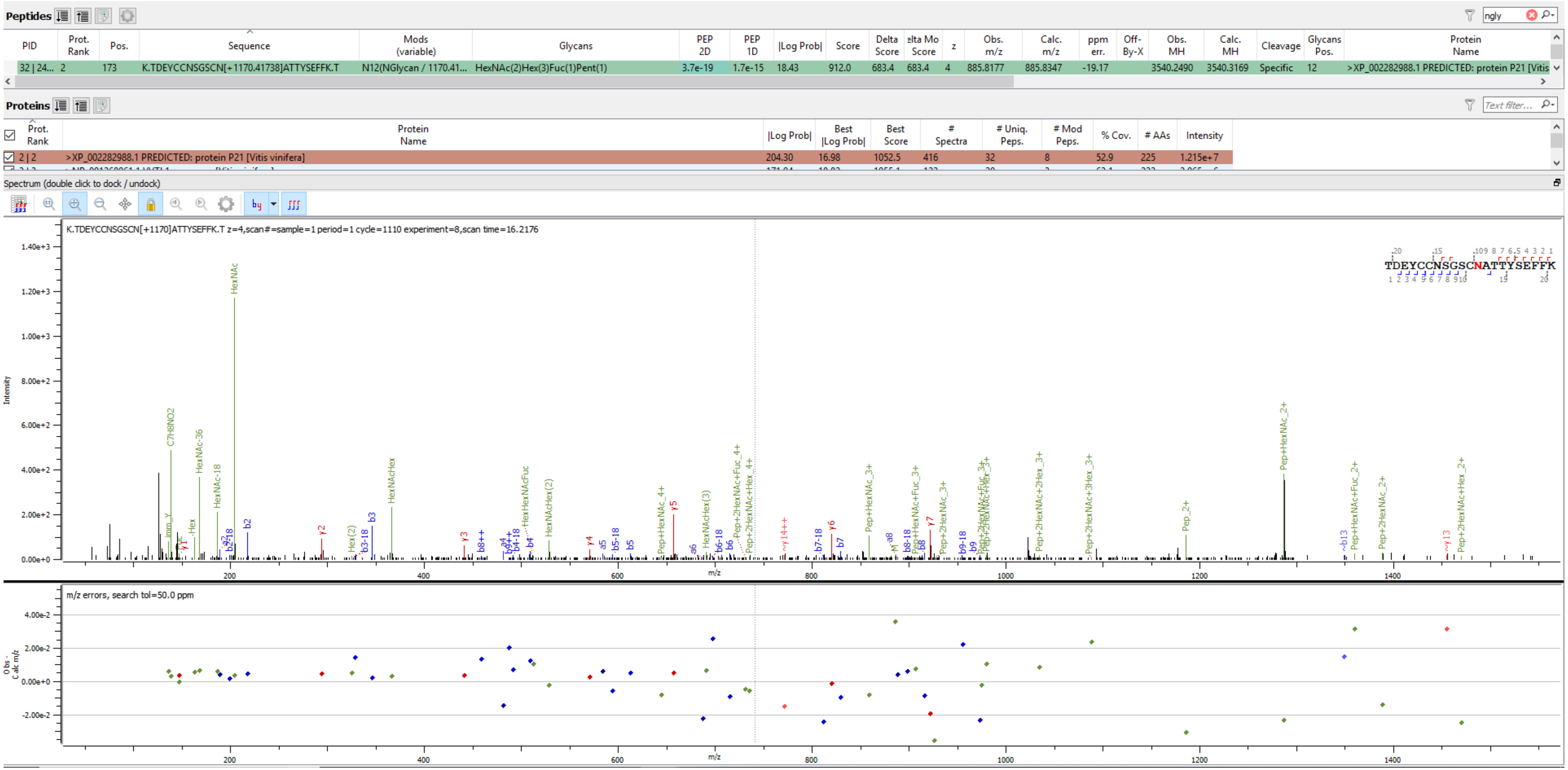


Sample S1 K.TDEYCCNSGSCN[+1170.417]ATTYSEFFK.T

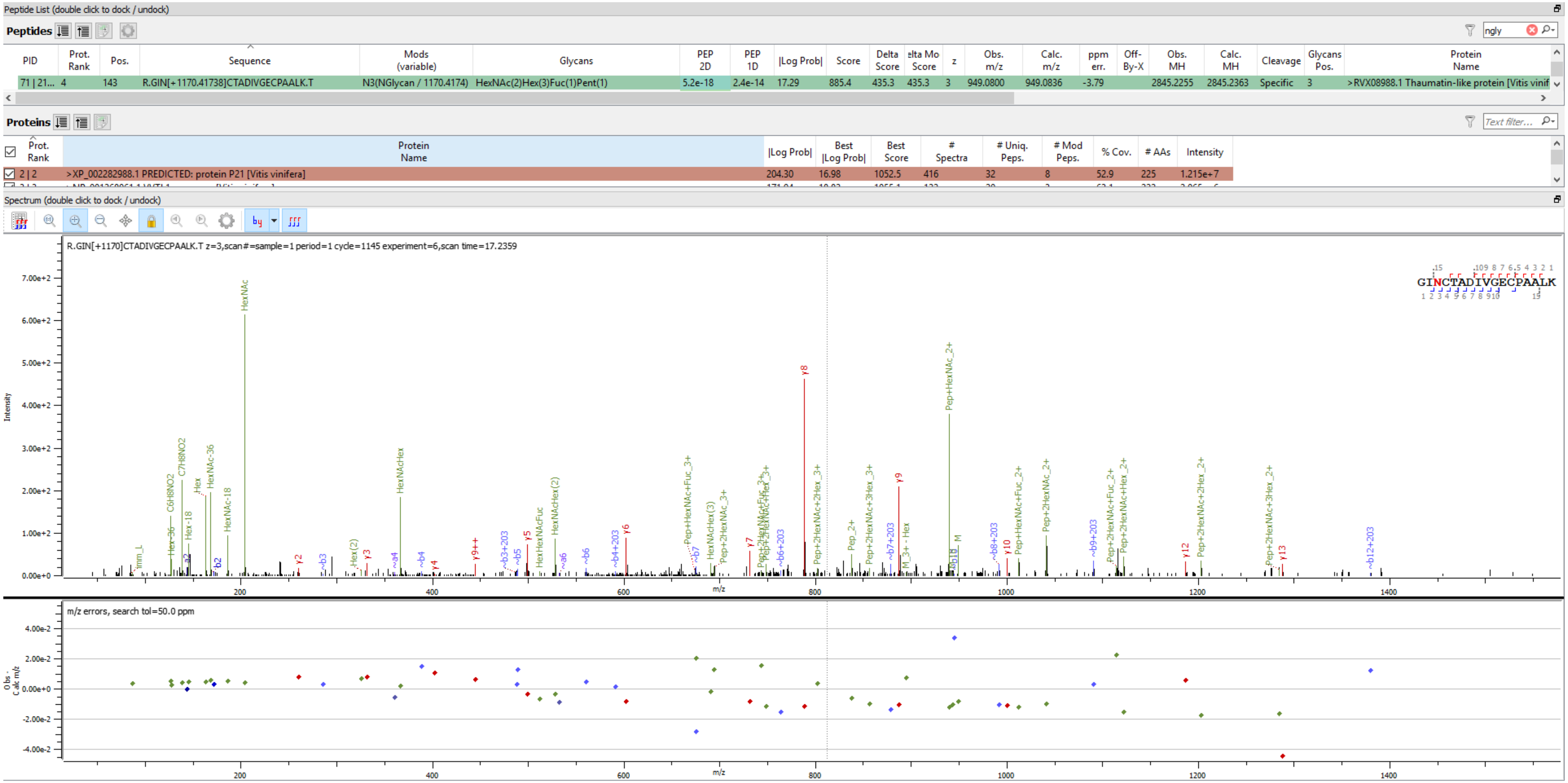


Sample S1 R.GIN[+1170.417]CTADIVGECPAALK.T

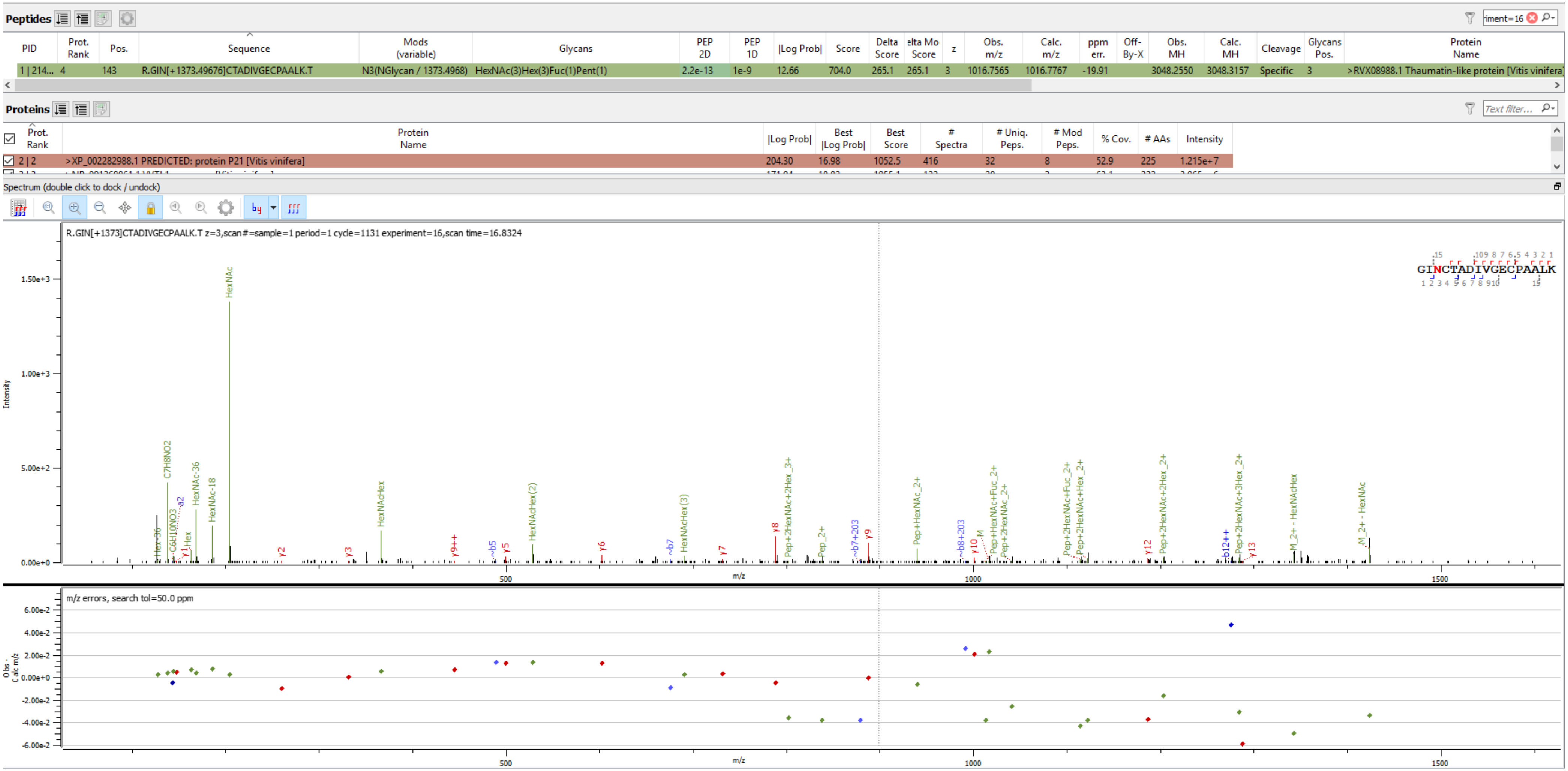


Sample S1 R.GIN[+1373.497]CTADIVGECPAALK.T

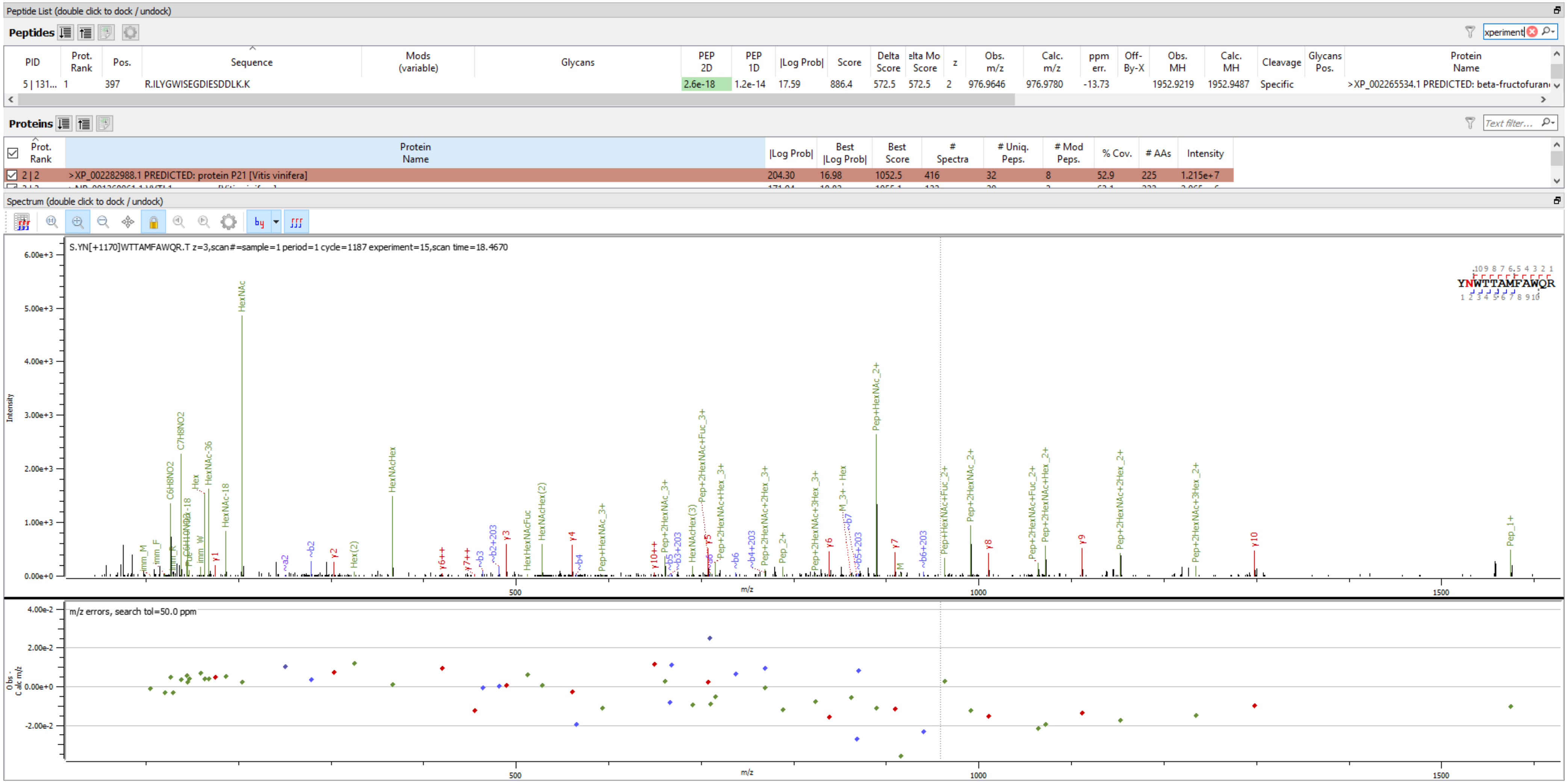


Sample S1 S.YN[+1170.417]WTTAMFAWQR.T

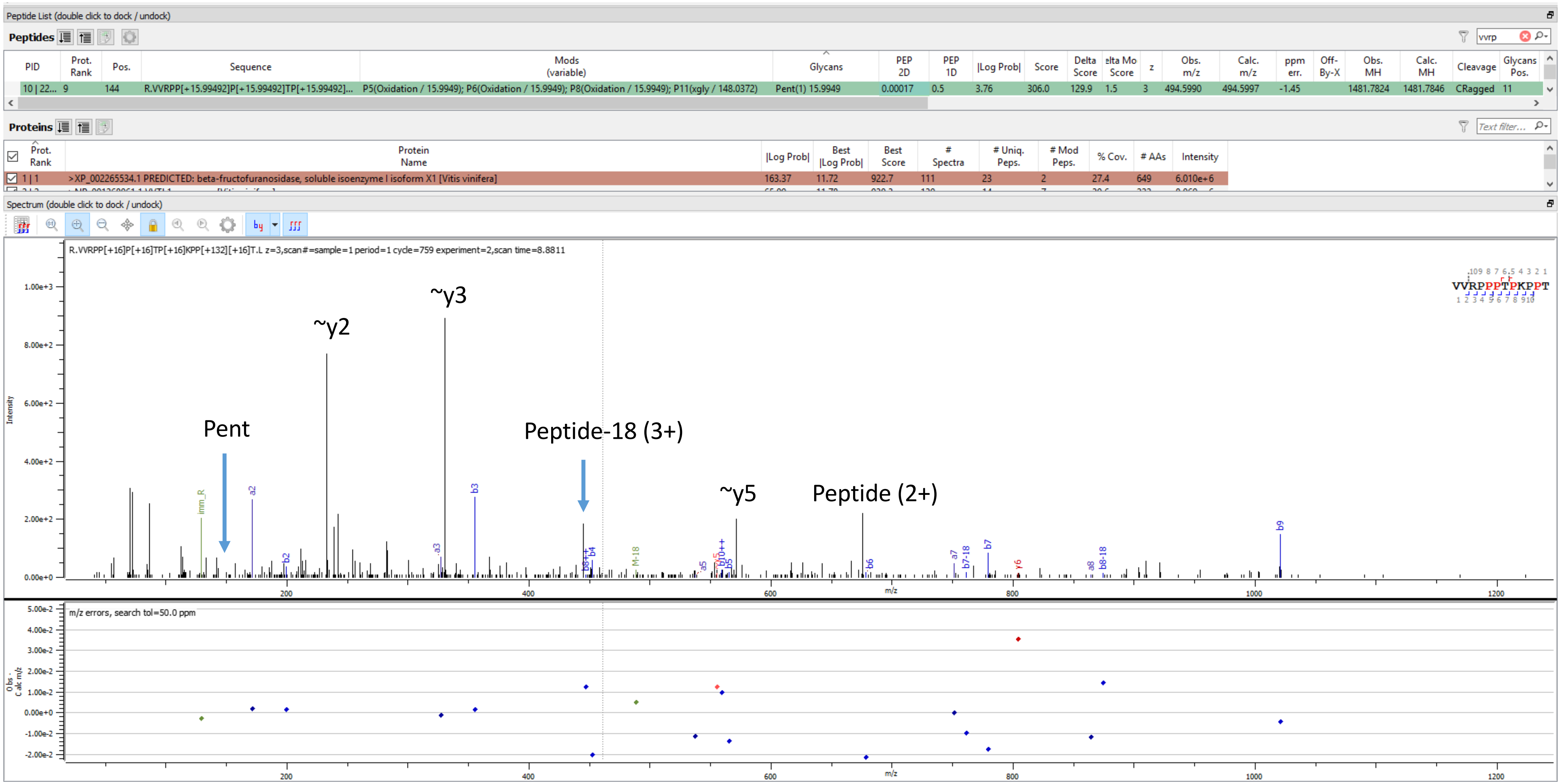


Sample S3 R.VVRPP[+15.995]P[+15.995]TP[+15.995]KPP[+132.042][+15.995]T.L

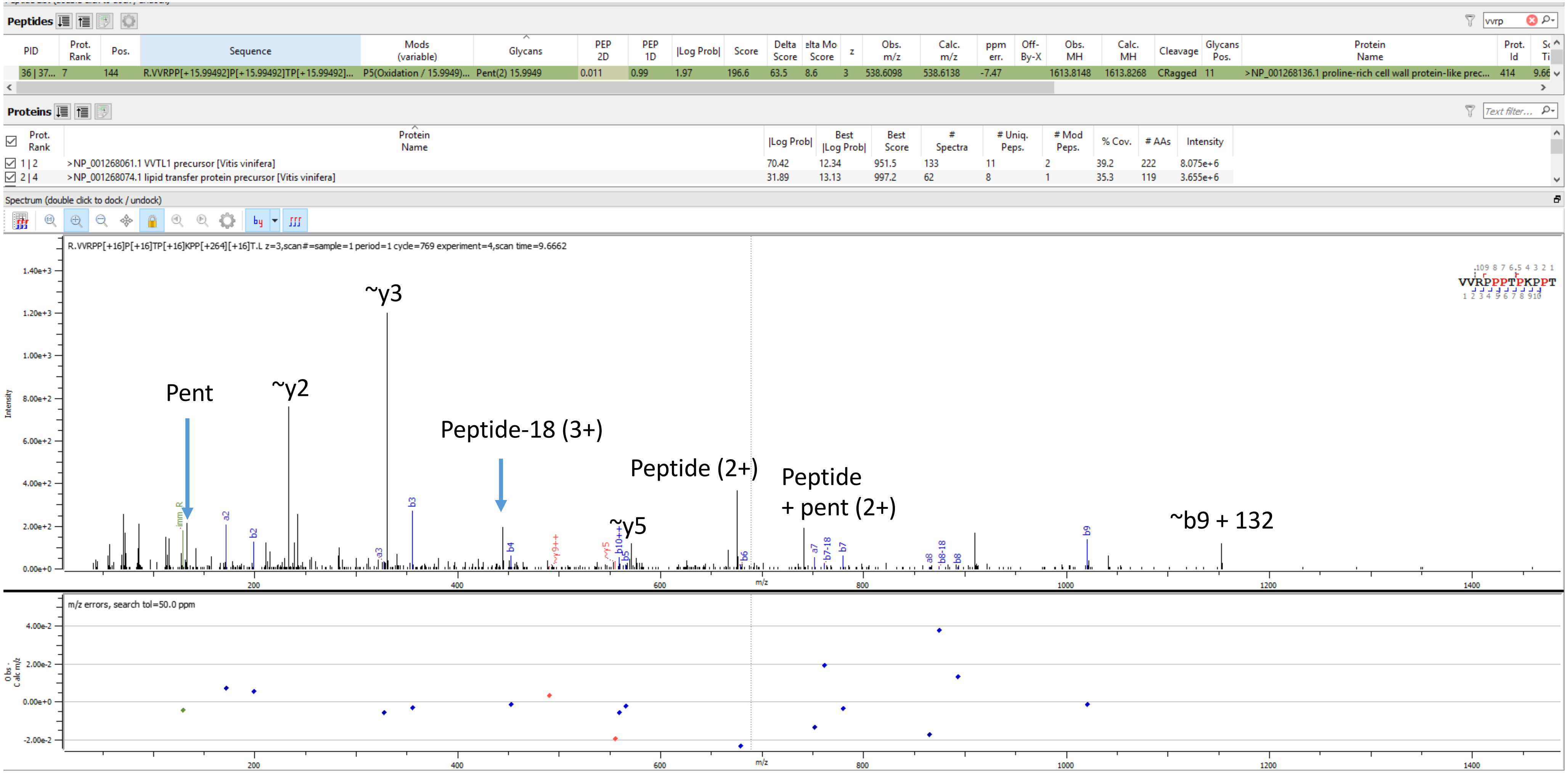


Sample S4 R.VVRPP[+15.995]P[+15.995]TP[+15.995]KPP[+264.085][+15.995]T.L

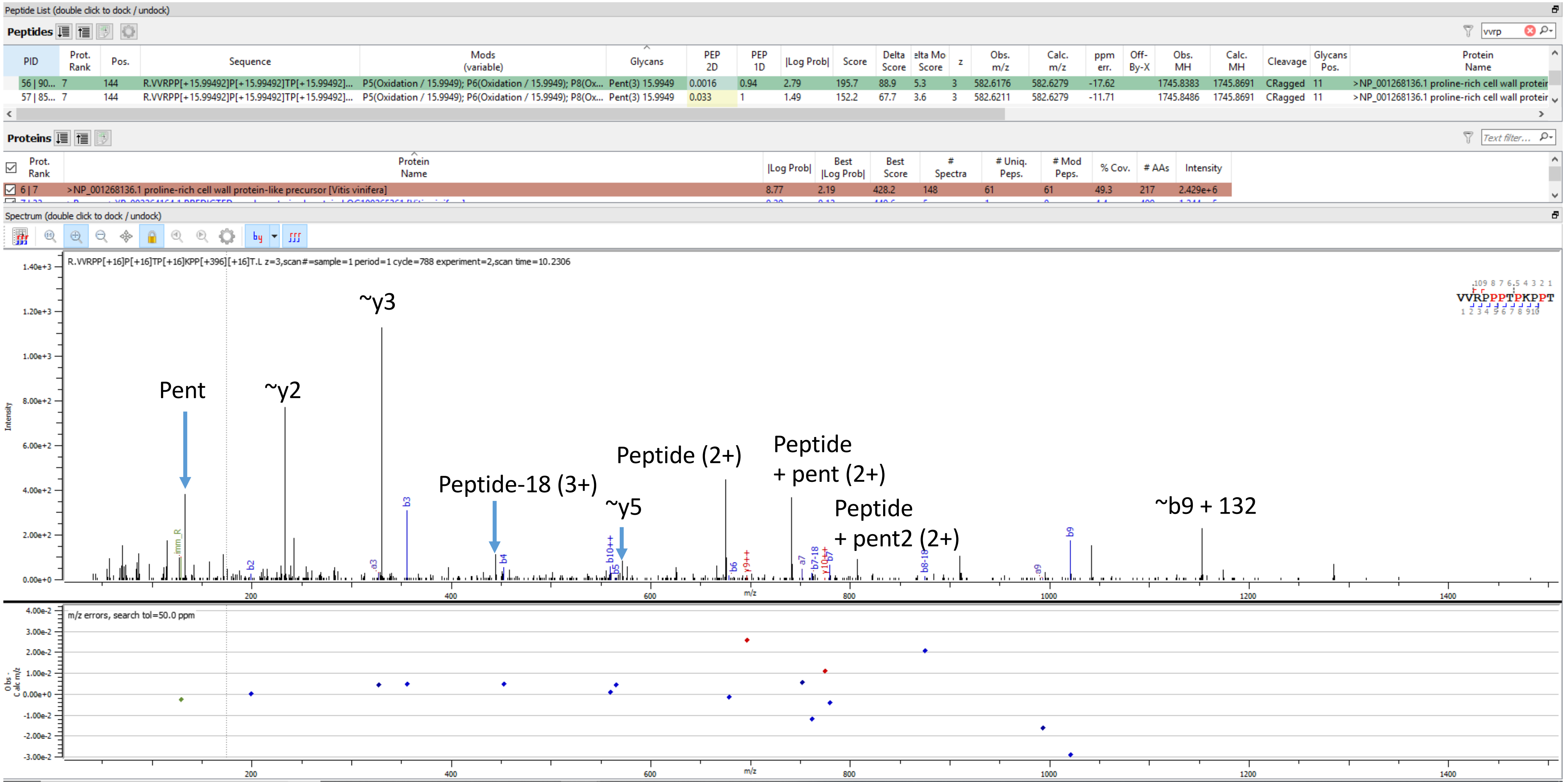


Sample S4 R.VVRPP[+15.995]P[+15.995]TP[+15.995]KPP[+396.127][+15.995]T.L

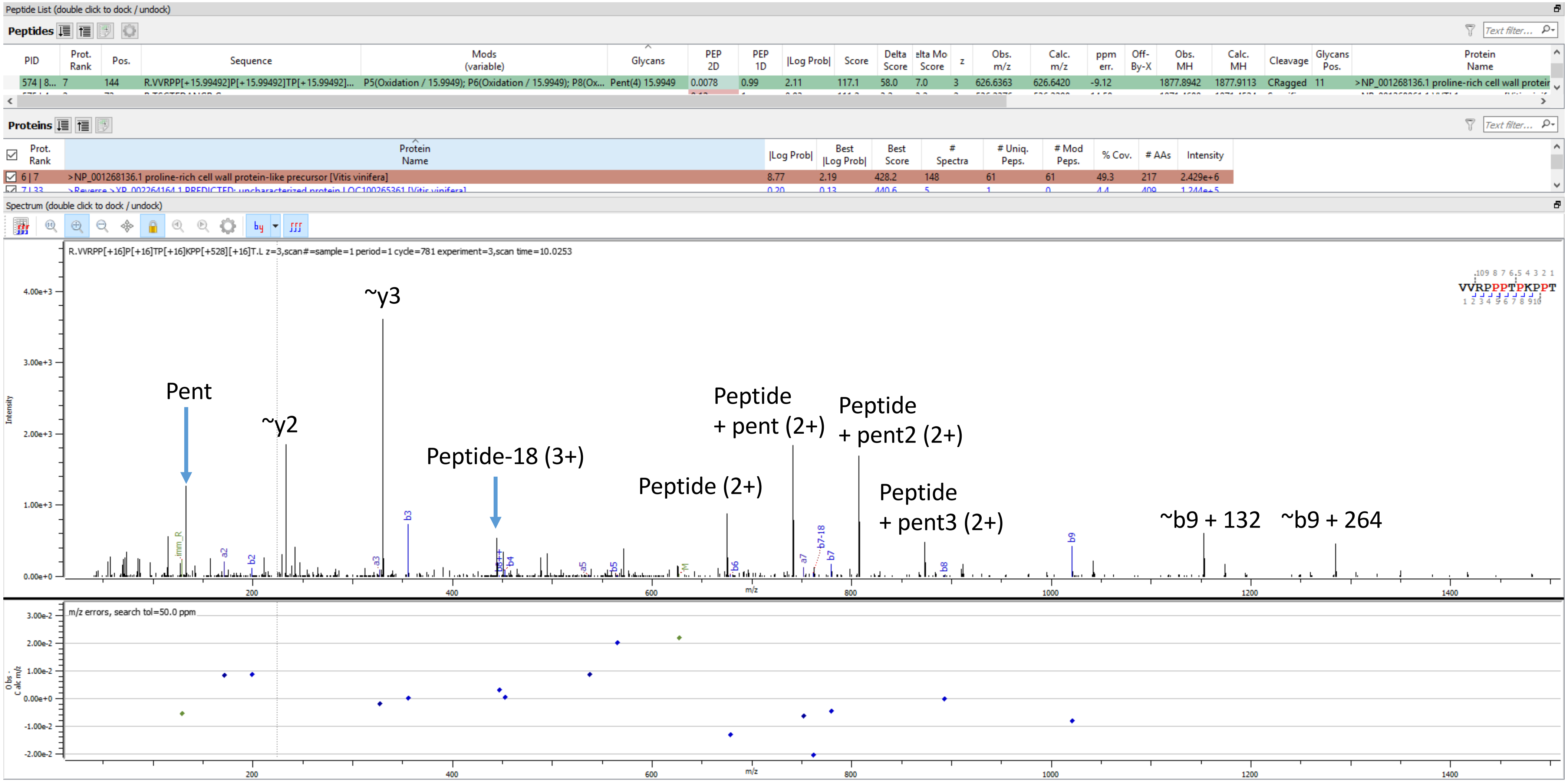


Sample S4 R.VVRPP[+15.995]P[+15.995]TP[+15.995]KPP[+528.169][+15.995]T.L

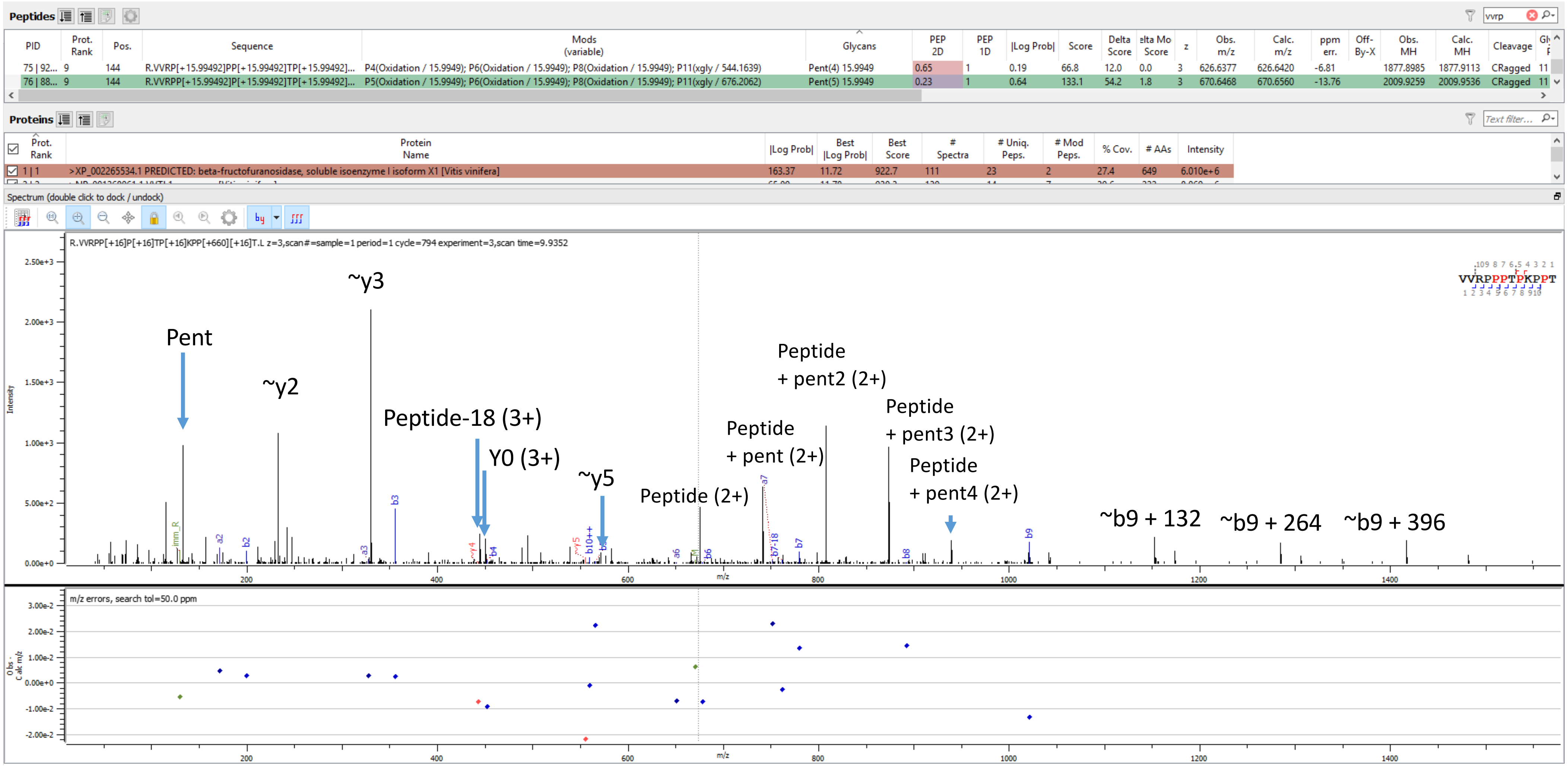


Sample S3 R.VVRPP[+15.995]P[+15.995]TP[+15.995]KPP[+660.211][+15.995]T.L

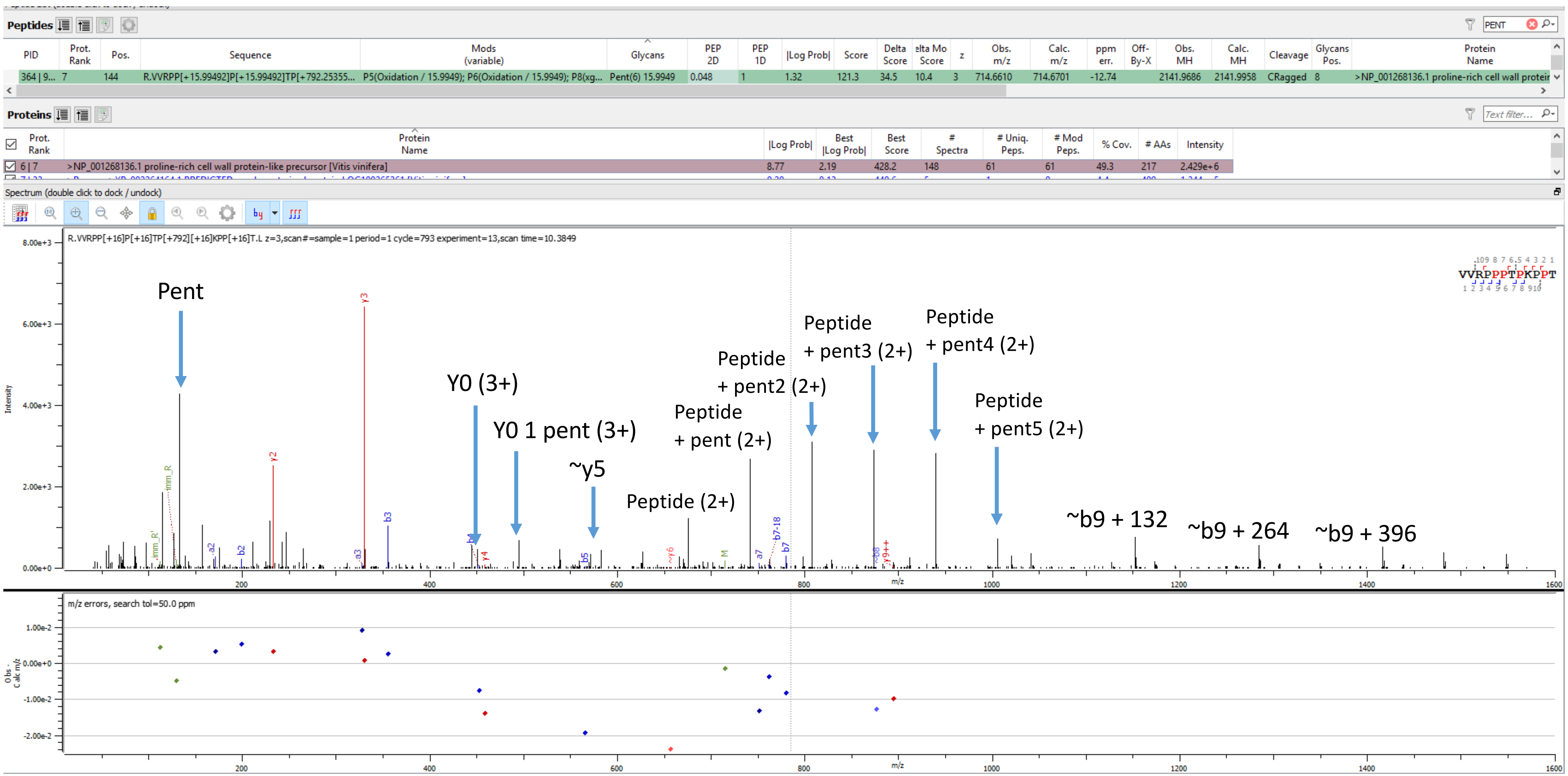


Sample S4 R.VVRPP[+15.995]P[+15.995]TP[+792.254][+15.995]KPP[+15.995]T.L
