## Supplementary Figure S2 for "Quantitative data independent acquisition glycoproteomics of sparkling wine"

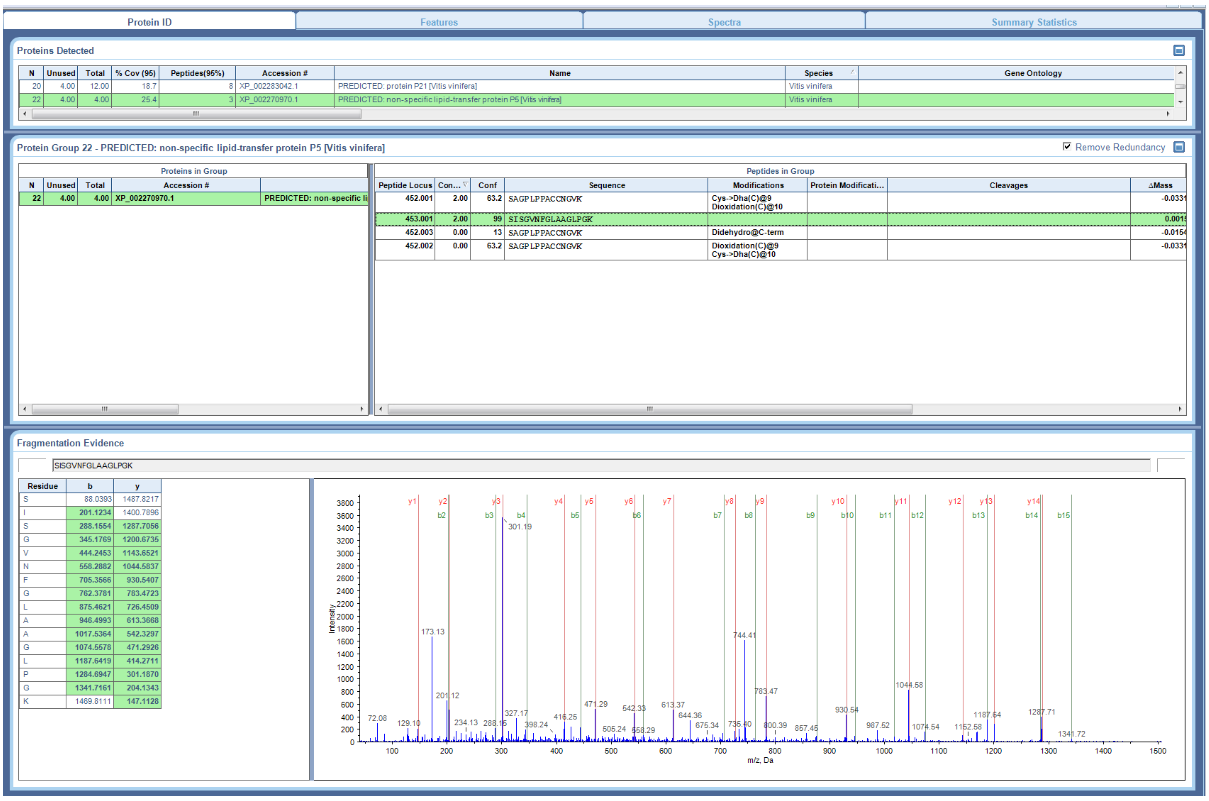


Supplementary Figure S2a. MS/MS of SISGVNFGLAAGLPGK identifying XP_002270970.1 PREDICTED: non-specific lipid-transfer protein P5 [Vitis vinifera].

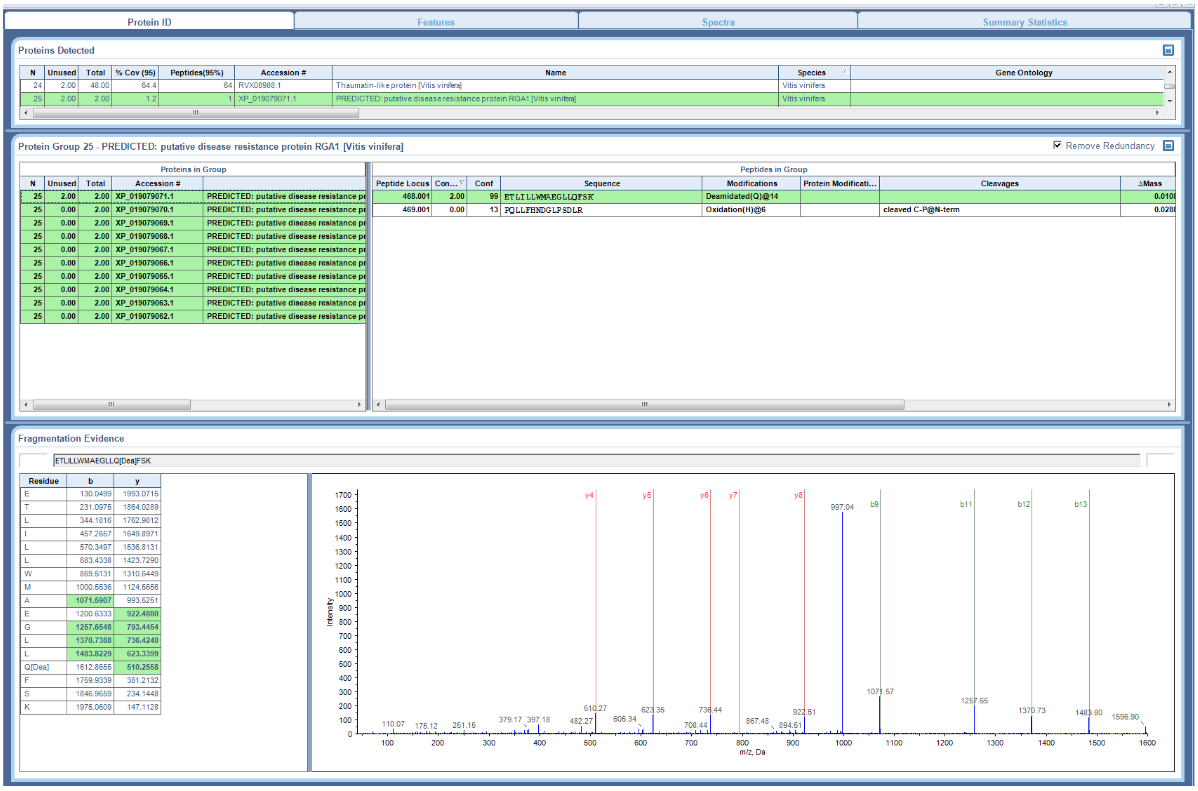


Supplementary Figure S2b. MS/MS of ETLILLWMAEGLLQFSK identifying XP_019079071.1 PREDICTED: putative disease resistance protein RGA1 [Vitis vinifera].

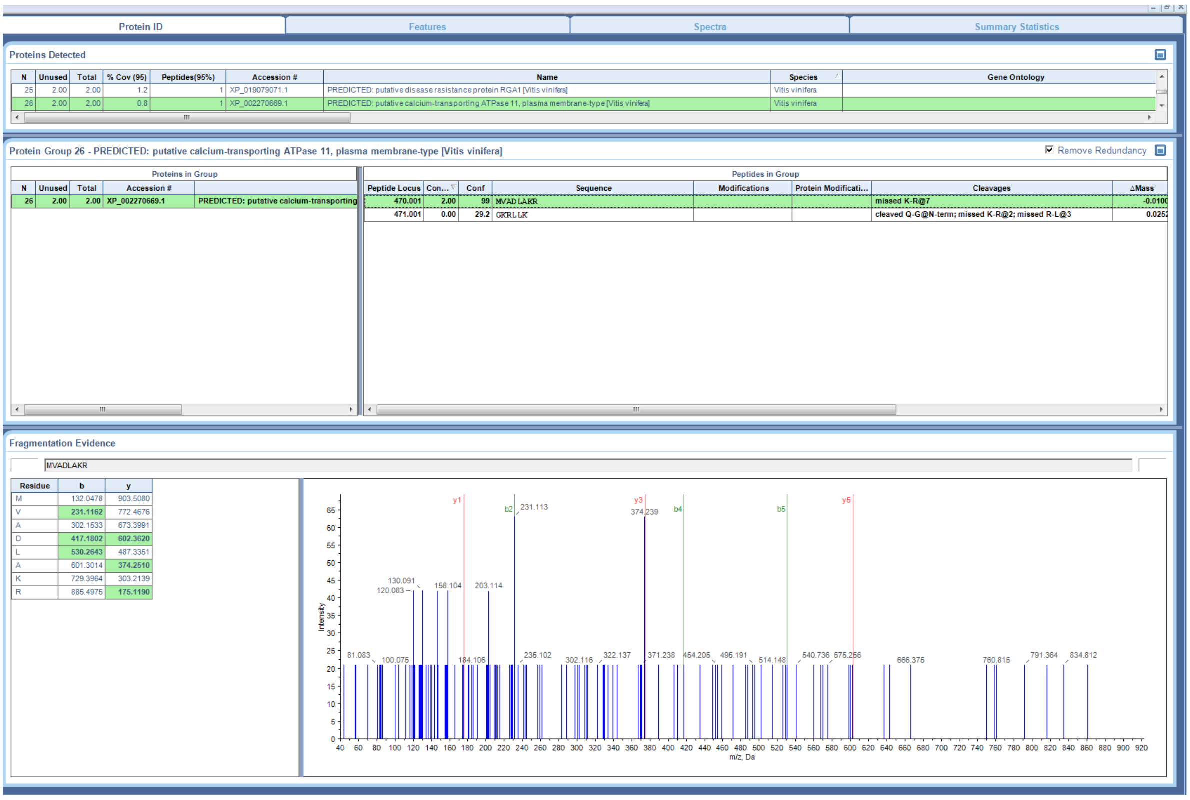


Supplementary Figure S2c. MS/MS of MVADLAKR identifying XP_002270669.1 PREDICTED: putative calcium-transporting ATPase 11, plasma membrane-type [Vitis vinifera].

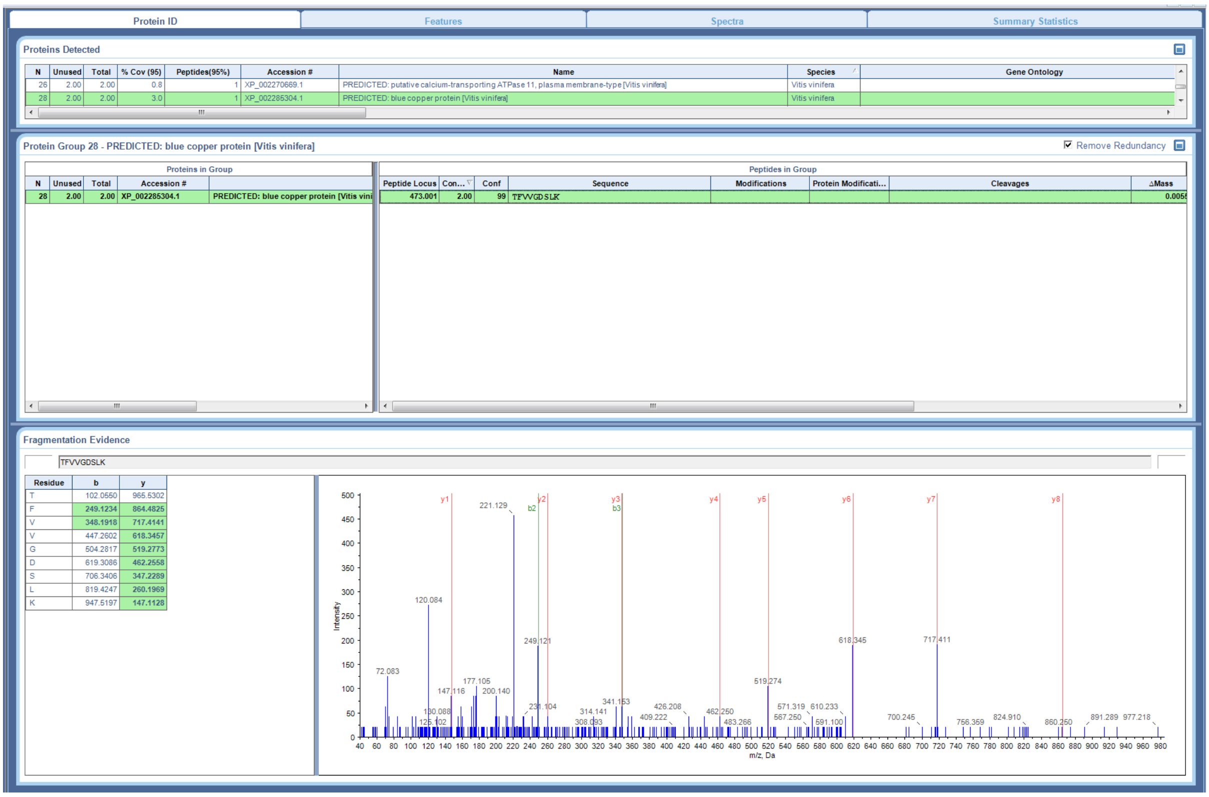


Supplementary Figure S2d. MS/MS of TFVVGDSLK identifying XP_002285304.1 PREDICTED: blue copper protein [Vitis vinifera].

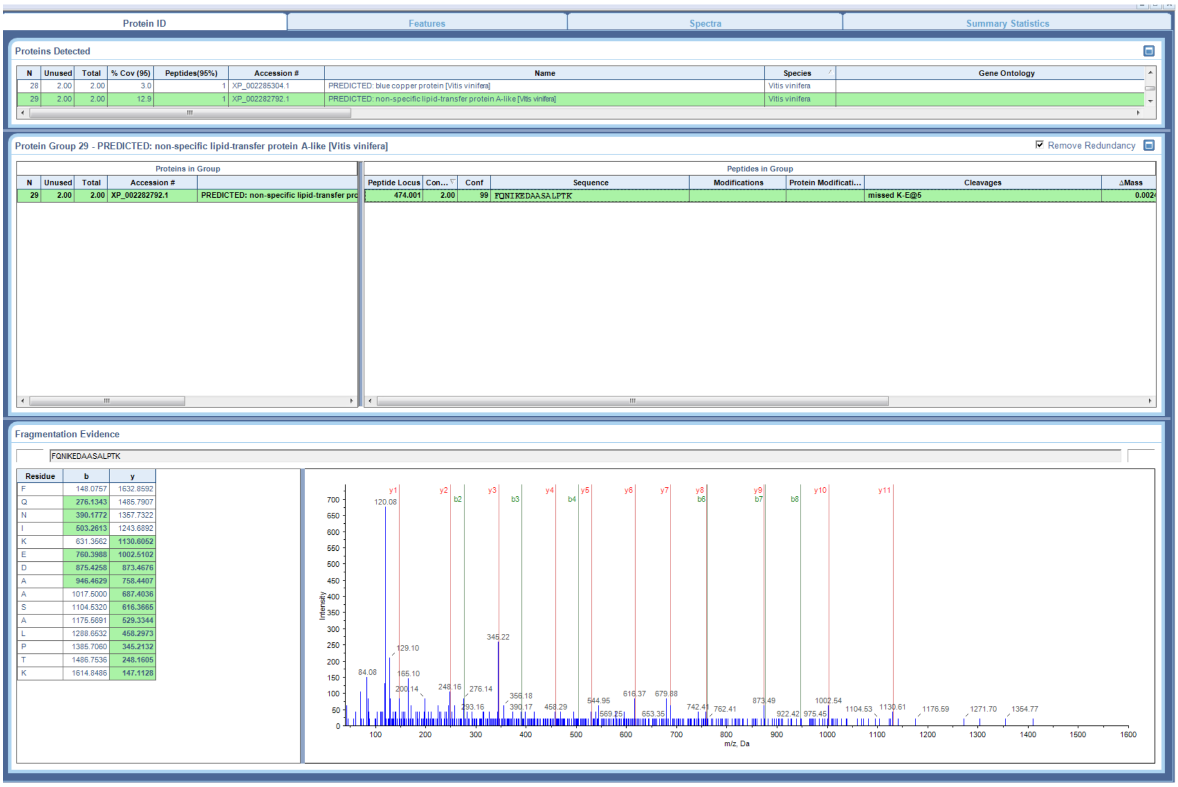


Supplementary Figure S2e. MS/MS of FQNIKEDAASALPTK identifying XP_002282792.1 PREDICTED: non-specific lipid-transfer protein A-like [Vitis vinifera].

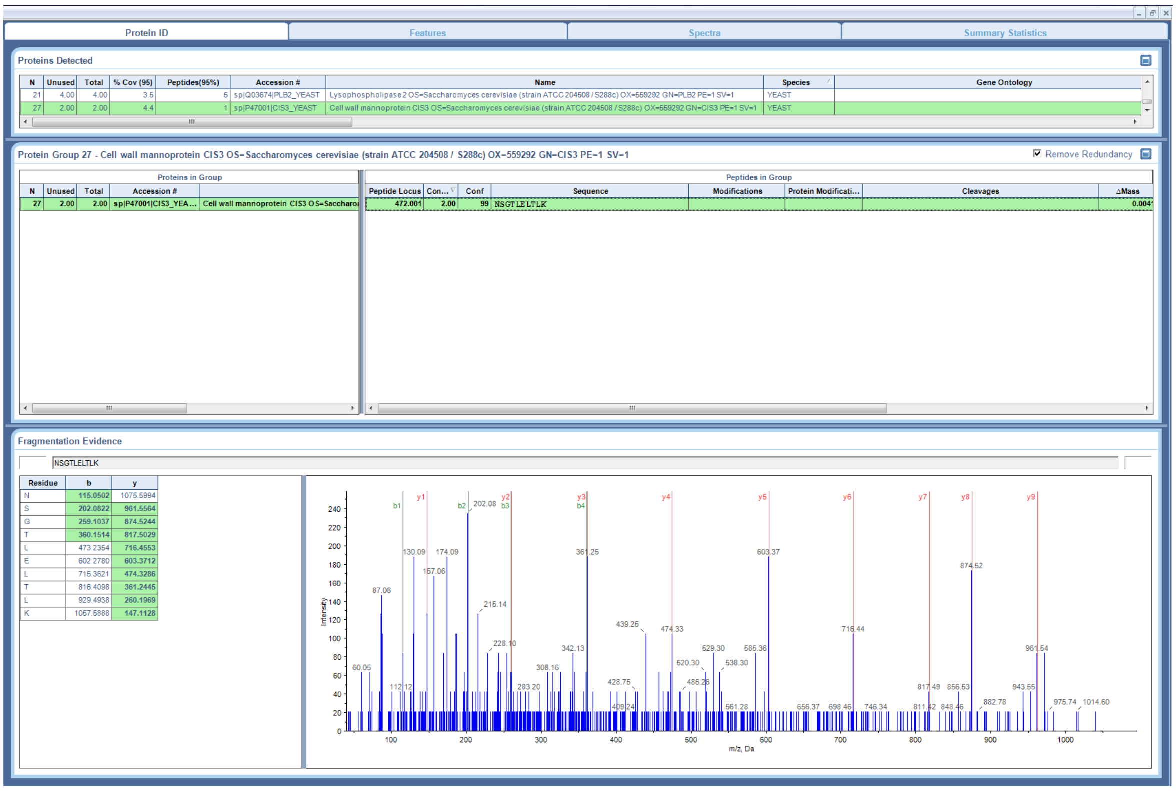


Supplementary Figure S2f. MS/MS of NSGTLELTLK identifying sp|P47001|CIS3_YEAST Cell wall mannoprotein CIS3 OS=Saccharomyces cerevisiae (strain ATCC 204508 / S288c) OX=559292 GN=CIS3 PE=1 SV=1.
