## Supplementary Figure S3 for "Quantitative data independent acquisition glycoproteomics of sparkling wine"

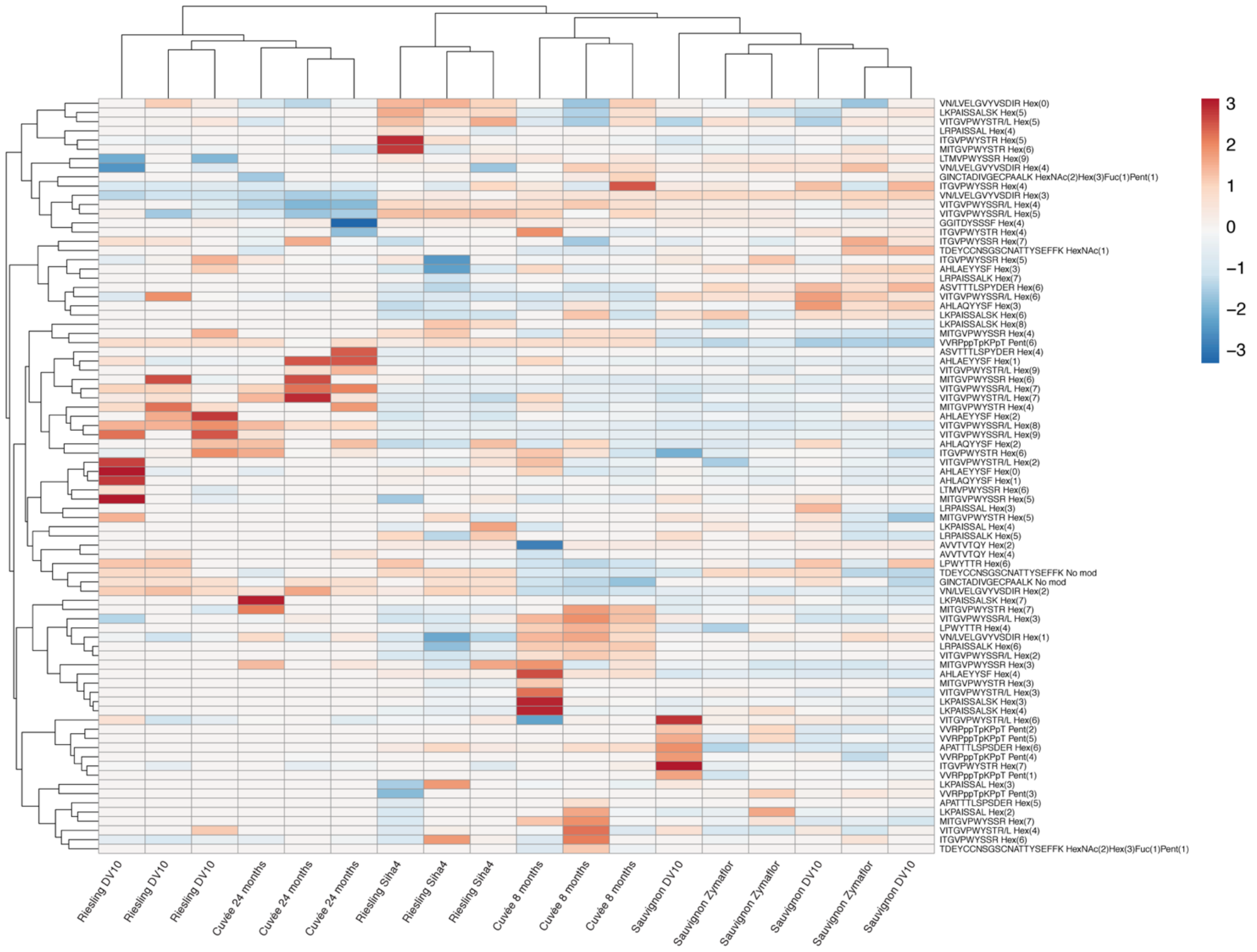


**Supplementary Figure S3.** Clustered heatmap of the relative abundance of glycoforms in all replicates between experimental conditions.. Both rows and columns were clustered using correlation distance and average linkage. The rows are labelled with the glycopeptide glycoforms. The samples are a cuvée aged 8 months or 24 months on lees; a Sauvignon blanc base wine fermented with the yeast strains DV10 or Zymaflore aged 16 months; and a Riesling base wine fermented with the yeast strains DV10 or Siha4 aged 17 months.
